# MCHelper automatically curates transposable element libraries across eukaryotic species

**DOI:** 10.1101/2023.10.17.562682

**Authors:** Simon Orozco-Arias, Pío Sierra, Richard Durbin, Josefa González

## Abstract

The number of species with high quality genome sequences continues to increase, in part due to scaling up of multiple large scale biodiversity sequencing projects. While the need to annotate genic sequences in these genomes is widely acknowledged, the parallel need to annotate transposable element sequences that have been shown to alter genome architecture, rewire gene regulatory networks, and contribute to the evolution of host traits is becoming ever more evident. However, accurate genome-wide annotation of transposable element sequences is still technically challenging. Several *de novo* transposable element identification tools are now available, but manual curation of the libraries produced by these tools is needed to generate high quality genome annotations. Manual curation is time-consuming, and thus impractical for large-scale genomic studies, and lacks reproducibility. In this work, we present the Manual Curator Helper tool MCHelper, which automates the TE library curation process. By leveraging MCHelper’s fully automated mode with the outputs from three de novo transposable element identification tools, RepeatModeler2, EDTA and REPET, in fruit fly, rice, hooded crow, zebrafish, maize, and human, we show a substantial improvement in the quality of the transposable element libraries and genome annotations. MCHelper libraries are less redundant, with up to 65% reduction in the number of consensus sequences, have up to 11.4% fewer false positive sequences, and up to ∼48% fewer “unclassified/unknown” transposable element consensus sequences. Genome-wide transposable element annotations were also improved, including larger unfragmented insertions. Moreover, MCHelper is an easy to install and easy to use tool.

## INTRODUCTION

After two decades of sequencing projects, reference genomes for thousands of eukaryotic species are already available and many more are currently being sequenced as part of large-scale coordinated initiatives (Consortium, 2022; Lewin et al., 2022). Thanks to long-read sequencing technologies, our ability to identify genomic variants has now expanded from the well-characterized single nucleotide polymorphisms and short indels, to structural variants, which have been shown to contribute more diversity at the nucleotide level than any other type of genetic variant (Sedlazeck et al., 2018). Accurate annotation of transposable elements (TEs), a major source of structural variants, is still challenging and so far, has only been performed in a few selected groups of species (Jebb et al., 2020; Osmanski et al., 2023). However, TEs are not only present in virtually all genomes studied to date, but in some groups of species, such as mammals and plants, they are also the largest genome component. Moreover, TEs have been shown to be major contributors to the organization, rearrangement, and regulation of genomes across species (Casacuberta & González, 2013, p. 201; Hayward & Gilbert, 2022). For example, TEs have been recruited to perform diverse biological functions in adaptive immunity, placental development, memory formation and brain development (Dunn-Fletcher et al., 2018; Huang et al., 2016; Pastuzyn et al., 2018), while they have also been associated with autoimmune and neurological diseases, cancer, and aging (De Cecco et al., 2019; Payer & Burns, 2019). Thus, lack of accurate annotations of TE sequences results in extensive undiscovered genetic variation with potentially important implications for genome function, structure and evolution.

While attempts at including TE diversity in the analysis of genome sequences are now common, they are often restricted to the use of homology-based methods, which are based on searching for known elements present in databases of the genome of interest, or that of a closely related species. However, the accuracy of TE identification using only homology-based methods decreases as the phylogenetic distance between the species being annotated and the species where the known elements were described increases (Platt et al., 2016). Combining homology-based and *de novo* TE identification in the species of interest is needed to accurately annotate TEs in genome sequences (Platt et al., 2016; Sotero-Caio et al., 2016). *De novo* methodologies use the repetitive nature and the structural characteristics of TE sequences to generate genome-specific libraries. Currently, there are several automatic tools available that allow *de novo* discovery and annotation of TE insertions *e.g.* RepeatModeler2 (Flynn et al., 2020), REPET (Flutre et al., 2011), and EDTA (Ou et al., 2019) (for a more comprehensive list visit: https://tehub.org/; Elliott et al., 2021). However, libraries produced by these tools lead to low quality and incomplete TE annotations (Goubert et al., 2022; Rodriguez & Makalowski, 2022). This is mainly due to the high diversity of TE sequences present in genomes that precludes the generation of high-quality libraries unless manual curation of the raw libraries produced is carried out. Due to the increasing interest in TE biology, several user-friendly manuals have been recently published (Baril et al., 2024; Goubert et al., 2022; Jamilloux et al., 2016; Platt et al., 2016; J. M. Storer et al., 2021, 2022). Additionally, some specific tools to aid in the manual curation have also been produced (Baril et al., 2024; Goubert et al., 2022; Orozco-Arias et al., 2021, 2022). Still, manual curation of TE libraries is time-consuming and requires acquiring knowledge on the biology of TEs present in the genomes of interest. As an example, the recent annotation of TEs in 248 mammal genome assemblies required between 1 and 19 rounds of detailed curation (Osmanski et al., 2023). Another important limitation of manual curation is the lack of reproducibility, since the curator takes decisions based on the available information but also based on his/her own expertise and experience (Baril et al., 2024). This lack of reproducibility is especially relevant for comparative genomic approaches that require consistent and accurate annotations of a large number of genomes, unlikely to be done by a single researcher.

In this work, we introduce the Manual Curator Helper, MCHelper, a tool that automates all the steps required to curate TE libraries, enabling non-TE experts to generate high quality eukaryotic TE libraries, as well as helping experienced TE curators to do so more efficiently. MCHelper is based on published manual curation protocols developed by experts and allows for faster and reproducible TE annotations. We tested MCHelper on libraries generated by three *de novo* TE tools, RepeatModeler2, EDTA and REPET, which differ in their output formats, and in six species, which differ in TE content and genome size: the fruit fly (*Drosophila melanogaster*), rice (*Oryza sativa*), the hooded crow (*Corvus cornix*), zebrafish (*Danio rerio*), maize (*Zea mays*), and human (*Homo sapiens*).

## RESULTS

### MCHelper integrates protocols to automatically curate transposable element libraries

MCHelper is a computational tool that integrates transposable element (TE) manual curation protocols developed by TE experts to improve the completeness and accuracy of TE libraries generated by automatic *de novo* tools (Baril et al., 2024; Goubert et al., 2022; Jamilloux et al., 2016; Ou & Jiang, 2018; Platt et al., 2016; J. M. Storer et al., 2021, 2022). MCHelper is a flexible tool designed to accept libraries generated by *de novo* TE identification tools that produce a FASTA file output, such as RepeatModeler2 (RM2; Flynn et al., 2020) or EDTA (Ou et al., 2019), as well as the output files produced by the *TEdenovo* pipeline from REPET (Flutre et al., 2011; Figure 1). MCHelper was designed to reduce the most common problems present in libraries generated with automatic tools: redundant, fragmented, false positive, and unclassified/unknown TE sequences. Briefly, MCHelper: (i) reduces consensus redundancy, once at the beginning and once at the end of the pipeline, using either the CD-HIT (Li & Godzik, 2006) or the MeShClust V3 (Girgis, 2022) clustering algorithms; (ii) extends the consensus sequences; (iii) identifies and filters false positive consensus sequences, *i.e.* sequences wrongly classified as TEs (see Methods); (iv) identifies previously known TEs based on homology; (v) performs structural checks on consensus sequences initially labeled by *de novo* TE identification tools as “classified” TEs; and (vi) assigns a classification to consensus sequences initially labeled as “unclassified/unknown” TEs (Figure 1 and Supplemental Fig S1).

**Figure 1.**
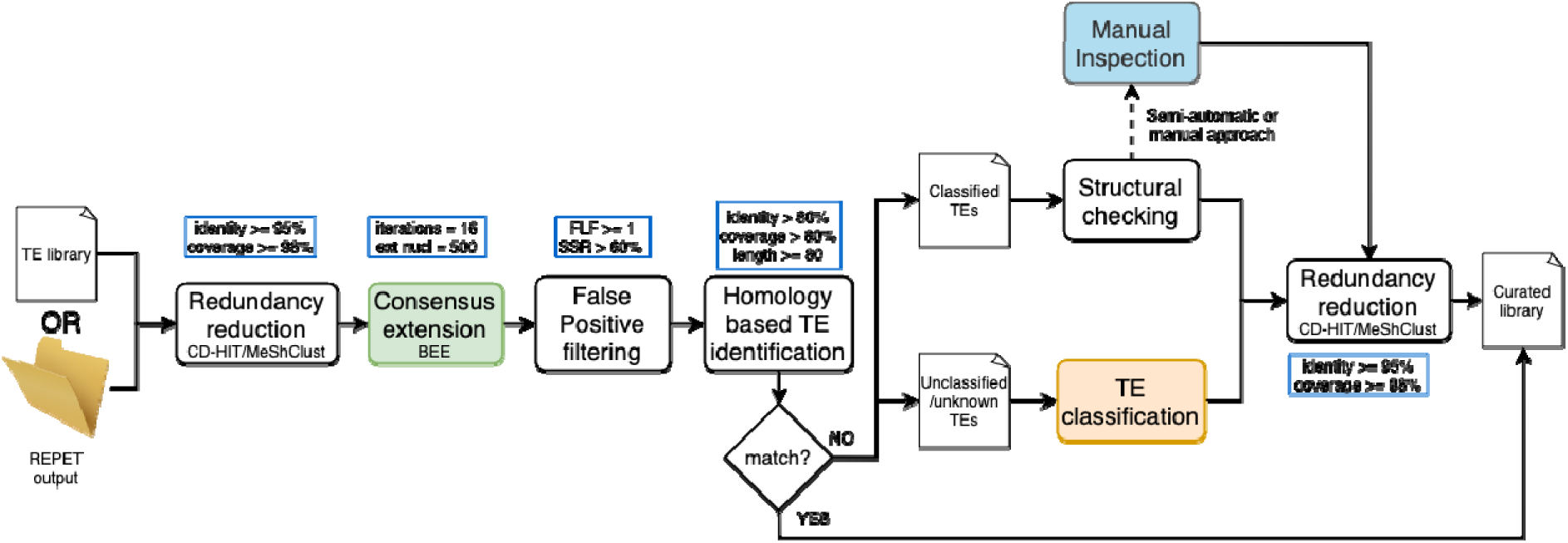
MCHelper’s tool workflow. Inputs, main steps, and default parameters of the MCHelper tool. MCHelper accepts as input a single FASTA file containing consensus sequences produced by a *de novo* TE identification tool or a folder containing files produced by the *TEdenovo* pipeline from REPET (see the MCHelper’s GitHub for details). Colored boxes correspond to modules that can also be executed independently. Workflows of these modules are provided in Supplemental Figure S1.

MCHelper offers three distinct levels of automation: fully-automatic, semi-automatic, and fully-manual, thus enabling users to tailor the software’s functionality to their curation requirements and time availability (Figure 1). In the fully-automatic mode, consensus sequences that pass the structural check (“complete”) and those that fail (“incomplete”) are both kept in the curated library (see Methods). “Incomplete” sequences are kept as they could correspond to older families for which a full-length copy is no longer present in the genome analyzed. MCHelper adds the suffix “_inc” to “incomplete” sequences for easy identification by the user. In the semi-automatic mode, MCHelper automatically retains only “complete” sequences and provides graphical information, such as TE-Aid plots (Goubert et al., 2022), multiple sequence alignment (MSA) plots, and information on TE length and copy number, to facilitate the user’s manual inspection of the “incomplete” sequences. The user is then responsible for deciding whether to keep, remove, or reclassify the “incomplete” sequences. In the fully-manual mode, MCHelper provides all the information needed to manually inspect all the consensus sequences.

### MCHelper improves the number, quality, and length of consensus sequences in transposable element libraries across species

To test the MCHelper tool, we ran the fully-automatic mode on six species that differ in genome size and TE content and that have available reference TE libraries: the fruit fly (*D. melanogaster*), rice (*O. sativa*), the hooded crow (*C. cornix*), zebrafish (*D. rerio*), maize (*Z. mays*), and human (*H. sapiens*). For each species, we ran MCHelper with the raw TE libraries obtained from three different *de novo* tools, RepeatModeler2 (RM2; Flynn et al., 2020), EDTA (Ou et al., 2019), and REPET (Flutre et al., 2011), with the exception of the human T2T genome where a REPET library could not be obtained. We compared the performance of the two clustering algorithms implemented in MCHelper, CD-HIT and MeShClust, and we found that the proportion of overlapping clusters between the algorithms is high and depends on the genome analyzed (Supplemental Fig S2).

We then compared the MCHelper libraries (generated using CD-HIT as the clustering algorithm) with the available reference libraries: BDGP (Kaminker et al., 2002) and MCTE libraries (Rech et al., 2022) for *D. melanogaster*, the manually curated “standard library” provided by Ou et al. (2019), for *O. sativa*, the MClibrary (Weissensteiner et al., 2020) for *C. cornix,* Dfam (J. Storer et al., 2021) and Repbase (Jurka et al., 2005) for *D. rerio,* MTEC curated by Ou et al. (https://github.com/oushujun/MTEC) for *Z. mays*, and the Dfam library for *H. sapiens*.

MCHelper reduced the total number of consensus sequences in the raw libraries produced by RM2, EDTA and REPET in the six species analyzed. Using the RM2 output, MCHelper reduced the number of consensus sequences between 34.3% in *O. sativa* and 54.8%, in *D. melanogaster* (Figure 2A and Supplemental Table S1A). Additionally, MCHelper was able to remove between 3.1% (in *D. melanogaster*) and 11.4% (in *O. sativa*) of false positives in these six species (Supplemental Figure S3 and Table S1B). Using EDTA’s output, MCHelper reduced the number of families between 40.7% (in *C. cornix*) and 65.4% (in *H. sapiens*), and removed 2.4% (in *C. cornix*) and 3.9% (in *D. rerio*) of false positives. Using REPET’s output, MCHelper reduced the number of families between 48.9% (in *D. rerio*) and 60.6% (in *C. cornix*), and removed between 2.8% (in *Z. mays*) and 6.7% (in *O. sativa*) of false positives. Thus overall, MCHelper libraries are more similar to the reference libraries than to the raw libraries, consistent with the findings of Storer et al. (2022), who noted that raw *de novo* libraries are typically twice as large as necessary.

**Figure 2.**
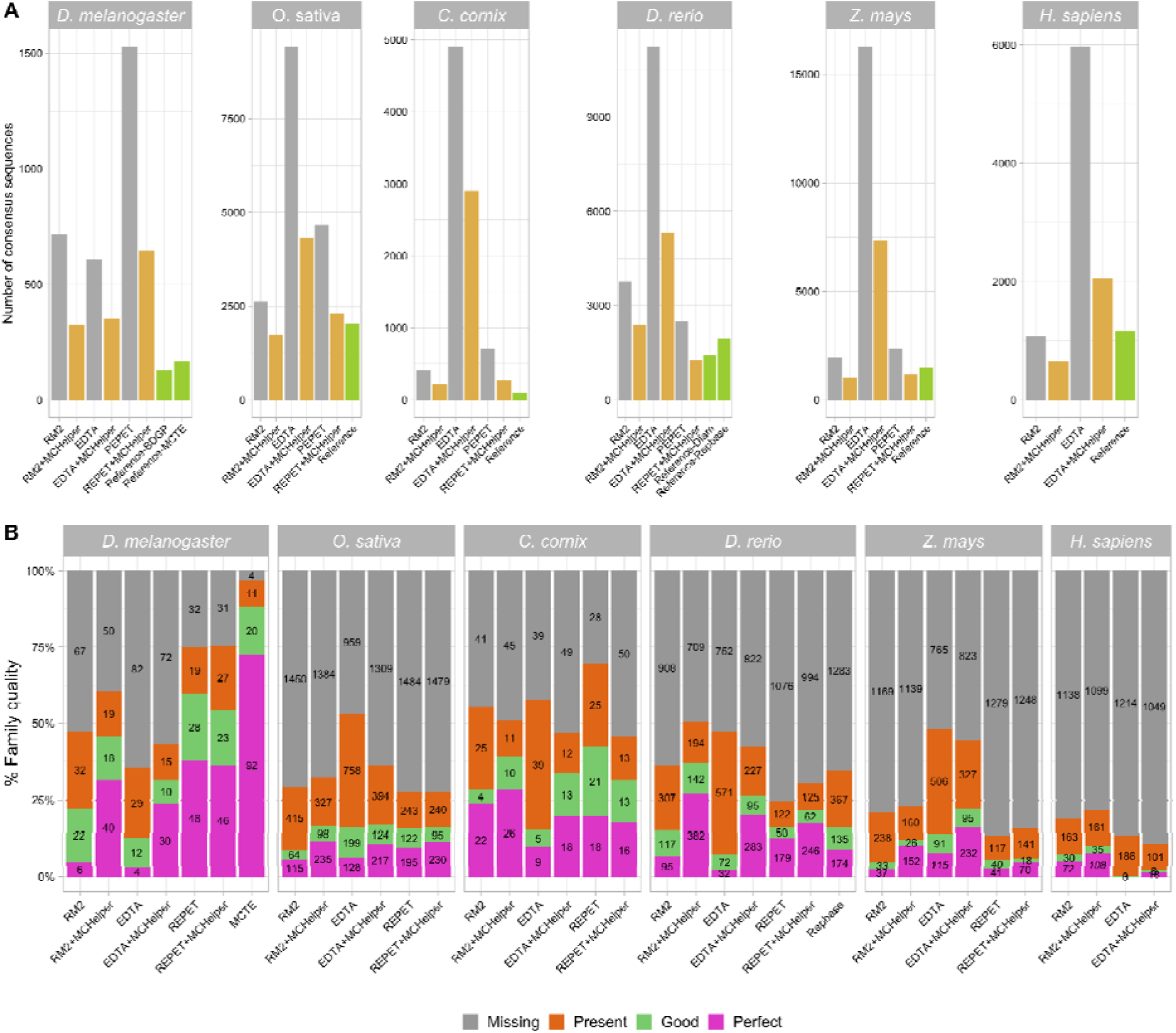
Comparison of MCHelper libraries with raw and reference libraries in the six genomes analyzed. A) Number of consensus sequences contained in the raw library, the MCHelper library and the reference libraries of the six species analyzed. Reference libraries used are: BDGP (Kaminker et al., 2002) and MCTE (Rech et al., 2022) for *D. melanogaster*, standard library (Ou et al., 2019) for *O. sativa*, MClibrary (Weissensteiner et al., 2020) for *C. cornix,* Dfam (J. Storer et al., 2021) and Repbase (Jurka et al., 2005) for *D. rerio,* MTEC curated by Ou (https://github.com/oushujun/MTEC) for *Z. mays*, and Dfam for *H. sapiens*. B) Number of consensus sequences classified as Perfect, Good, Present and Missing families following the methodology proposed by Flynn et al. (2020). Note that for *D. melanogaster* and *D. rerio* the MCTE and the Repbase libraries are also compared to the BDGP and Dfam libraries, respectively.

MCHelper libraries contained up to 8.8 times more perfect families than the raw libraries (Figure 2B and Supplemental Table S1C). Family quality was defined according to the methodology described in Flynn et al. (2020), which classifies each family contained in a reference library as perfect, good, present, or missing in the comparison library (see Methods). We also found that the majority of the MCHelper automatically curated libraries contained fewer or very similar numbers of missing families for all genomes and *de novo* tools (Figure 2B).

Finally, MCHelper consistently generated longer consensus sequences than the raw libraries, thus exhibiting lengths more similar to those reported in the reference libraries (Supplemental Figure S4 and Supplemental Table S1D) across almost all major orders of TEs: LINE, SINE, LTR, and MITEs). We observed that for all the species, except for *C. cornix*, MCHelper also generated, in average, longer consensus sequences for TIR elements in the raw libraries produced by 2/3 of the programs tested (2/2 in *H. sapiens*). In the case of Helitrons, MCHelper generated longer consensus sequences in the raw libraries produced by 1/3 of the programs tested in four species (*C. cornix, D. rerio, Z.mays,* and *H. sapiens*) and in 2/3 in the other two (*D. melanogaster* and *O. sativa*). This observation reinforces that raw libraries often contain fragmented TEs instead of complete consensus sequences, underscoring the importance of the consensus extension step in the curation process (Figure 1, see Methods) (Baril et al., 2024; Goubert et al., 2022; J. M. Storer et al., 2021). However, as noted by Baril et al. (2024), longer consensus sequences do not always equate to a better representation of the TE family. To avoid over-extensions and fuzzy terminations, MCHelper incorporates an approach that detects when the consensus sequence reaches either end (Supplemental Fig S1A). We observed that the percentage of TEs that do not contain their corresponding terminal repeats, or poly(A) tail sequences decreased in MCHelper libraries in comparison with RM2’s raw libraries up to 38% (in *Z. mays*), with EDTA’s raw libraries up to 13% (in *D. melanogaster*), and with REPET’s raw libraries up to 12% (in *Z. mays*) (Supplemental Table S2). Additionally, we tested how often chimeric TE sequences can be found in the raw libraries and in the MCHelper libraries. As a proxy to identify chimeric TE sequences, we did a BLASTN search between each library and the MCHelper’s internal TE database (see Supplementary Methods) to look for those consensus sequences with hits with >80% identity and >50% coverage with consensus sequences from two or more orders. We found that the MCHelper libraries contained less chimeric TEs than the raw libraries in all species, except for *H. sapiens* in which the MCHelper library contains one chimeric sequence while the raw EDTA library does not contain any (Supplemental Table S3).

Finally, we compared the MCHelper tool with SENMAP, a tool specifically designed to manually curate full-length LTR consensus sequences (Orozco-Arias et al., 2021). While SENMAP works better in plant genomes compared with animal genomes, as expected since SENMAP was trained with plant data, MCHelper identifies a higher number of LTR elements across tools as well as a higher number of perfect families (Supplemental Table S4A). If we focus on complete LTRs, while SENMAP identifies more LTRs from *D. melanogaster* REPET libraries compared with MCHelper, the number of perfect families identified by MCHelper is higher (Supplemental Table S4B).

### MCHelper recovers up to ∼48% of the TEs originally labeled as “unclassified/unknown”

Another common issue with raw libraries is the high number of unclassified TEs, especially in non-model organisms due to the lack of TE consensus sequences for these species in the available TE databases. To assign a classification to previously unclassified TEs, MCHelper follows a three-step process: homology search, coding domain presence (*e.g.* transposase and reverse transcriptase), and terminal repeat presence (LTRs and TIRs) (Supplemental Fig S1C; see Methods). Using this approach, MCHelper was able to assign a TE class for up to 48.24% of the TEs initially labeled as “unclassified/unknown” TEs (Figure 3A and Supplemental Table S5A). The majority of TEs in the six genomes analyzed were re-classified in the homology step, with the REPET output showing the bigger improvement compared to RM2, with the exception of *Z. mays* (Figure 3A). Across species and *de novo* TE identification tools, TEs from the LTR and TIR orders were the most commonly recovered sequences, consistent with the relative abundance of these orders in the genomes analyzed, with the exception of *H. sapiens* where LINE was more common (Figure 3B and Supplemental Table S5B).

**Figure 3:**
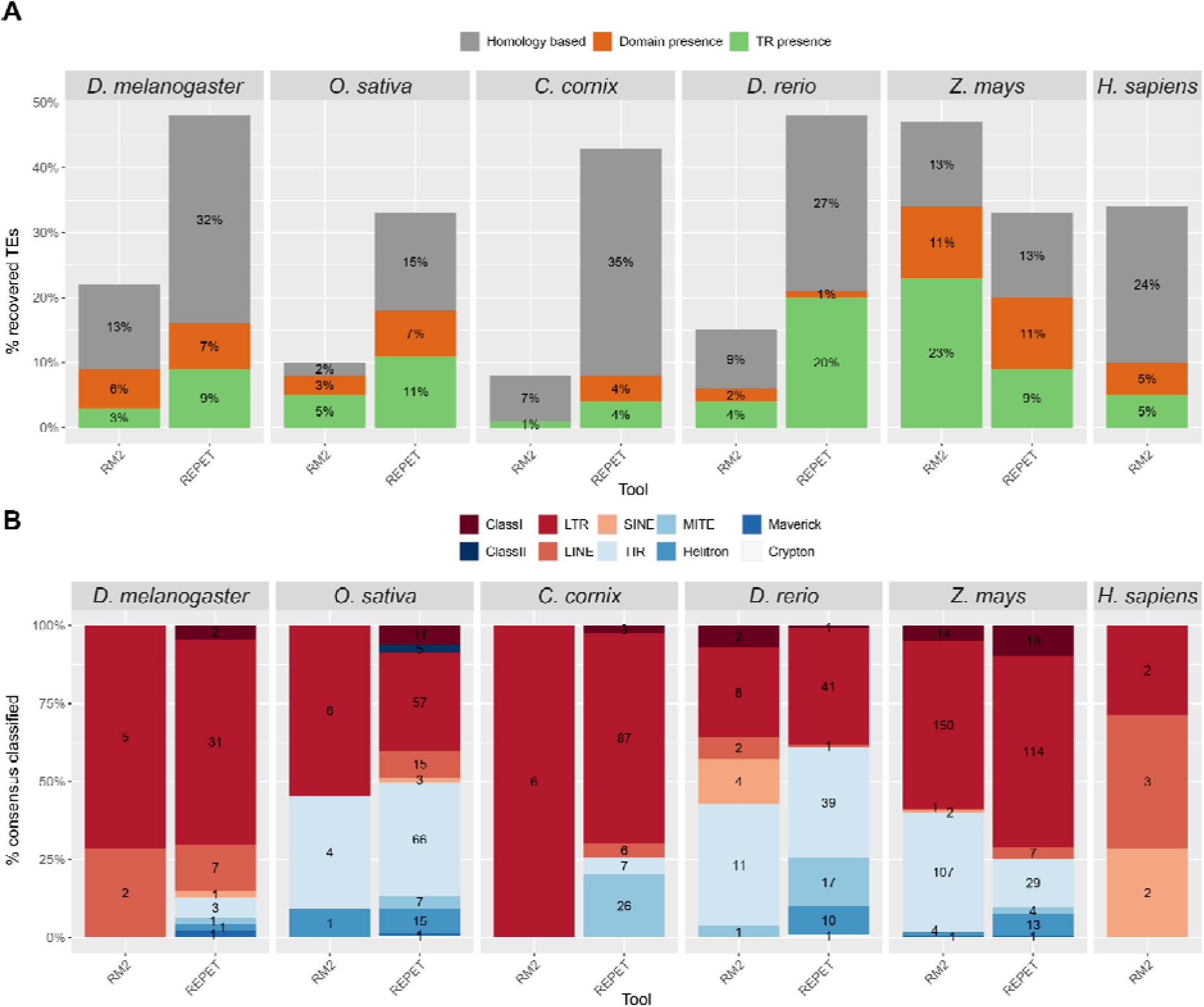
“Unclassified/unknown” consensus sequences in the raw libraries that are classified by MCHelper. A) Proportion of TEs originally labeled as “unclassified/unknown”, which were recovered by the three-step approach of MCHelper. TR: terminal repeat. B) Number of recovered TE sequences by TE order/class. Class I and Class II indicate all the elements that contain one or more retrotransposon-like domains in the first case or DNA-like domains in the latter case, but it was not possible to assign a deeper classification. EDTA is not included in this figure because the software does not report any “unknown/unclassified” elements.

We also used TEClass2 (Bickmann et al., 2023), TERL (da Cruz et al., 2021), and DeepTE (Yan et al., 2020) to classified the “unclassified/unknown” TEs (see Methods) to compare the performance against MCHelper and we found that although the recovering proportion was smaller in MCHelper compared with the other tools, the MCHelper precision was generally higher (Supplemental Table S6).

### Genome-wide TE annotations with MCHelper libraries are more accurate

Genome-wide TE annotations performed with raw TE libraries often lead to overestimation of the number of TE copies, mostly because a single copy is annotated as several fragmented copies (Jamilloux et al., 2016). In addition, raw libraries may have chimeric consensus sequences leading to the annotation of a single TE copy by multiple consensus sequences (Rodriguez & Makalowski, 2022). To evaluate the quality of the annotations obtained with the different TE libraries (raw, MCHelper, and reference), we analyzed the total number of copies annotated, the number of short copies annotated (<100 bp), and the number of overlapping annotations (copies annotated with more than one consensus).

We found that the number of TE copies annotated with the MCHelper library was smaller than the number of copies annotated with the raw libraries for the majority of tools and species analyzed (Figure 4A and Supplemental Table S7A). The reduction in number of copies ranged between 9.9% (in *D. rerio*) and 37.8% (in *Z. mays*) compared with the RM2 libraries, between 24.5% (in *H. sapiens*) and 48.8% (in *Z. mays*) compared with EDTA libraries and between 0.2% (in *O. sativa*) and 16.5% (in *Z. mays*) compared with REPET libraries. The only exceptions were the *C. corvix* genome, in which the number of annotated copies by MCHelper libraries were slightly bigger, and the annotation of the human genome with the raw RM2 library.

**Figure 4.**
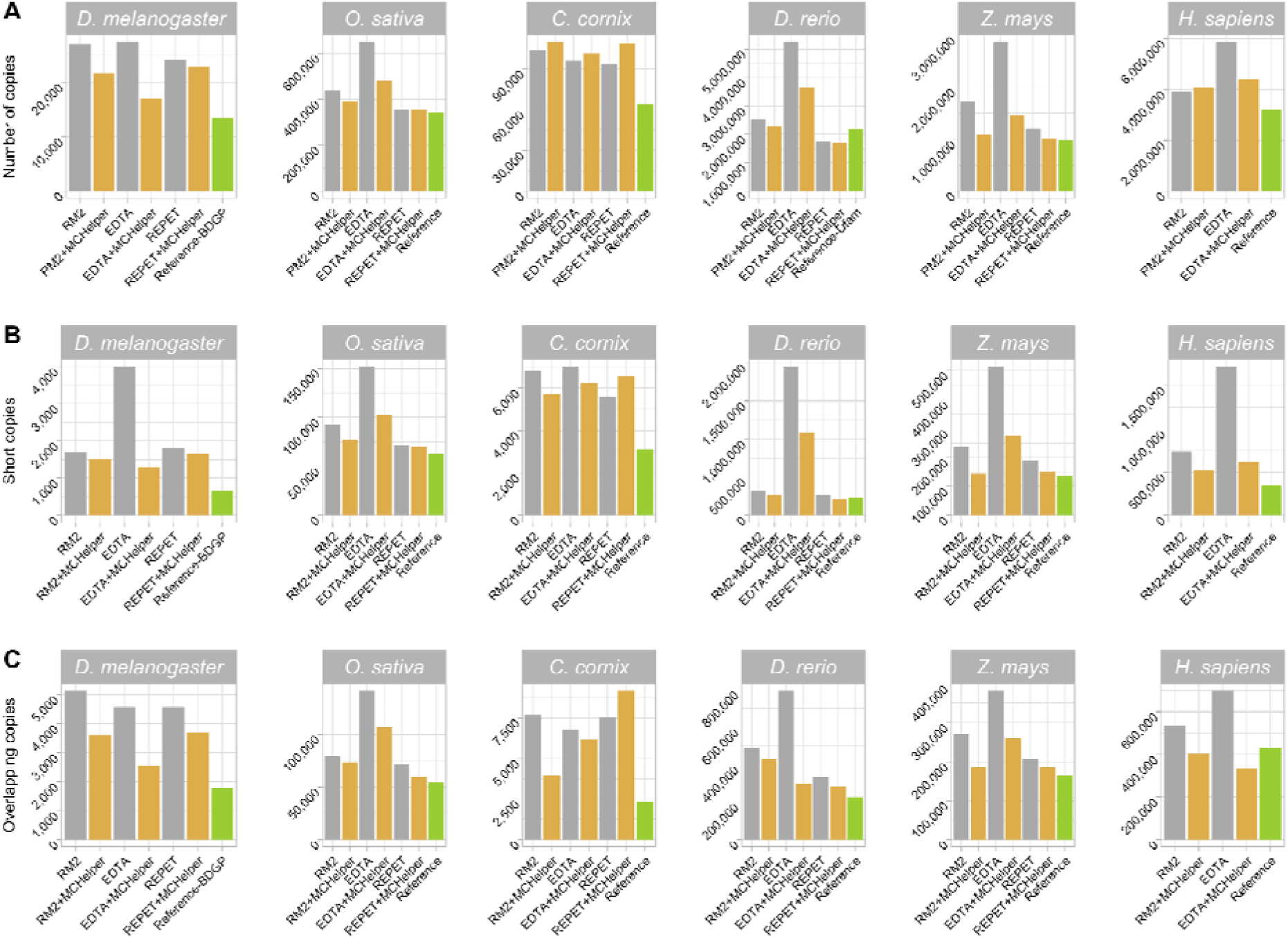
Genome-wide number and type of TE annotations for each of the three libraries analyzed in the six species. The BDGP, EDTA, MClibrary, Dfam-zebrafish, MTEC, and Dfam-Human libraries were used as reference for *D. melanogaster, O. sativa, C. cornix, D. rerio, Z. mays,* and *H. sapiens*, respectively. A) Number of copies annotated by each library. B) Number of copies categorized as short (<100bp). C) Number of copies categorized as overlapping (annotated by two or more consensus sequences).

The number of short copies (< 100 bp) was also reduced in MCHelper annotations, getting closer to the number of short copies in reference annotations in the six species when comparing with the raw RM2 annotations (between 10.8% in *D. melanogaster* and 39% in *Z. mays* fewer short copies) (Figure 4B and Supplemental Table S7B). When comparing MCHelper with EDTA and REPET, the number of short annotated copies was also reduced between 11.2% (in *C. cornix*) and 67.7% (in *D. melanogaster*) for EDTA outputs and between 2.4% (in *O. sativa*) and 20.9% (in *Z. mays*) in REPET outputs. The only exception found was *C. cornix* with the REPET+MCHelper, where the number of short copies increased by 14.5%.

Finally, MCHelper reduces the number of overlapping annotations, *i.e.* those annotated using more than one consensus sequence, again resulting in annotations that were more similar to the reference ones (Figure 4C and Supplemental Table S7C). The MCHelper library led to a decrease of overlapping annotations of 29.4% (RM2), 44% (EDTA) and 19.2% (REPET) in *D. melanogaster*, of 7.8% (RM2), 24.4% (EDTA) and 17.6% (REPET) in *O. sativa*, of 47.6% (RM2), and 9.9% (EDTA) in *C. cornix*, of 11.7.% (RM2), 61.8% (EDTA) and 14.7% (REPET) in *D. rerio*, of 31% (RM2), 31.9% (EDTA) and 9.7% (REPET) in *Z. mays*, and of 25.4% (RM2) and 52.1% (EDTA) in *H. sapiens*. The only exception was found in *C. cornix* where MCHelper annotated 17.9% more overlapping copies compared with REPET. Note that the number of overlapping copies might be an overestimate as the raw libraries, and thus the MCHelper libraries, are only classified to the superfamily level. Thus copies that are detected as annotated with two consensus sequences might actually be annotated with redundant consensus from the same family or subfamily.

### Annotations made by MCHelper libraries are longer and more contiguous than those of raw libraries

To assess the improvements in the genome-wide TE annotation based on the MCHelper libraries compared with the annotation based on the raw libraries, we used as a proxy the NTE50 and the LTE50 metrics, and the proportion of each TE order in the genome annotations. Briefly, the NTE50 is the number of the largest copies needed to annotate 50% of the mobilome. Thus, lower NTE50 values indicate that annotated copies are longer and thus that the annotation includes large unfragmented TE insertions (Jamilloux et al., 2016). The LTE50 is the length such that 50% of the mobilome is annotated by copies longer than that. So, higher LTE50 indicates better annotations (Jamilloux et al., 2016).

Our results showed that MCHelper libraries produced longer annotations than the raw library for the majority of tools and genomes (Figure 5). The NTE50 values were smaller when annotating the genome with the MCHelper library compared with the raw library with only three exceptions: EDTA NTE50 values were smaller in *D. melanogaster* and *C. cornix*, and REPET NTE50 values were smaller in *C. cornix* (Figure 5 and Table 1). Similarly, and as expected if the quality of MCHelper TE annotations is higher than the annotation performed with the raw library, the LTE50 values were higher when the genome was annotated with MCHelper libraries except for the annotation with EDTA libraries in three of the six species analyzed: *D. melanogaster*, *O. sativa* and *C. cornix*.

**Figure 5.**
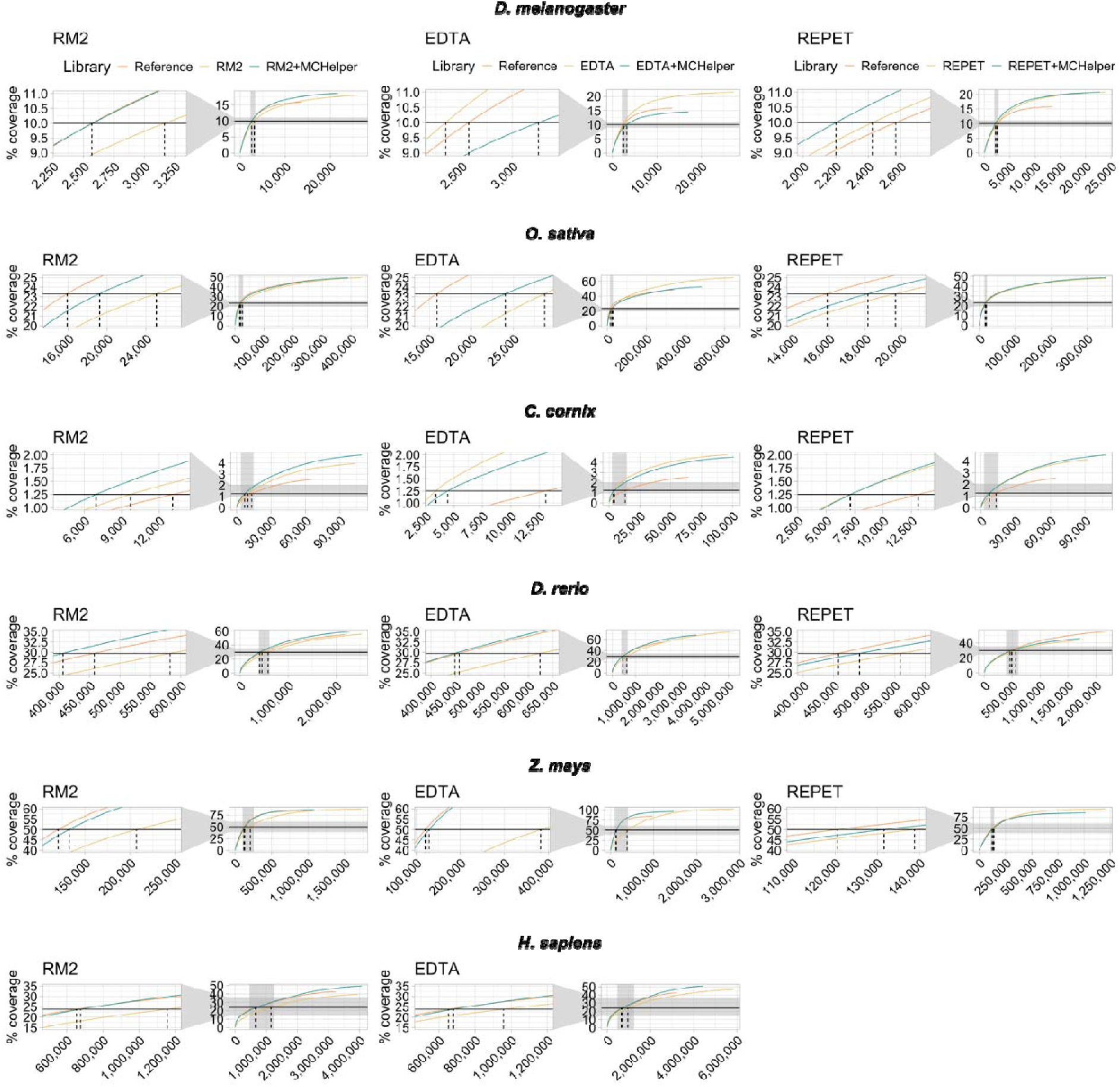
Cumulative length coverage plots for TE annotations in the six species analyzed. The black horizontal line represents half of the mobilome and it was used to calculate the LTE50 values (Table 1). The dash vertical line indicates the NTE50 values. For each graph, a zoom capture is provided on the left to show when the cumulative length plot of each library crosses the black horizontal line (NTE50 values). Mobilome proportions are 20% in *D. melanogaster* (Goubert et al., 2015; Hill 2019; Mérel et al., 2020), 46.64% in *O. sativa* (Ou et al., 2019), 2.5% in *C. cornix,* 59.5% in *D. rerio* (Chang et al., 2022), and 47.68% in *H. sapiens* (Hoyt et al 2022). In *Z. mays* we used 50% of the full genome space to calculate the NTE50 and LTE50 as recommended by Jamilloux et al. (2016). NTE50 and LTE50 values are shown in Table 1.

**Table 1.** NTE50 and LTE50 values of each library. The NTE50 is calculated as the number of the largest copies needed to annotate 50% of the mobilome, while the LTE50 is defined as the length of the shortest TE annotation that annotates 50% of the mobilome (Jamilloux et al., 2016).

| Library | *D. melanogaster* |  | *O. sativa* |  | *C. cornix* |  | *D. rerio* |  | *Z. mays* |  | *H. sapiens* |  |
| --- | --- | --- | --- | --- | --- | --- | --- | --- | --- | --- | --- | --- |
|  | NTE50 | LTE50 | NTE50 | LTE50 | NTE50 | LTE50 | NTE50 | LTE50 | NTE50 | LTE50 | NTE50 | LTE50 |
| **RM2** | 3167 | 2188 | 25075 | 1296 | 9699 | 769 | 582490 | 378 | 207065 | 2564 | 1155606 | 327 |
| **RM2+**<br>**MCHelper** | 2577 | 3007 | 19235 | 1824 | 6953 | 1077 | 406526 | 515 | 132536 | 5046 | 676997 | 464 |
| **EDTA** | 2342 | 3925 | 28674 | 1584 | 3405 | 2164 | 623608 | 340 | 382400 | 1565 | 957177 | 321 |
| **EDTA+**<br>**MCHelper** | 3259 | 1762 | 24031 | 1411 | 4453 | 1371 | 448658 | 468 | 129175 | 5328 | 680093 | 464 |
| **REPET** | 2426 | 3514 | 19675 | 1754 | 7108 | 979 | 559160 | 337 | 138840 | 4897 | - | - |
| **REPET+**<br>**MCHelper** | 2201 | 4082 | 18144 | 2032 | 7150 | 1051 | 492326 | 364 | 131590 | 5074 | - | - |
| **Reference** | 2570 | 3158 | 15900 | 2279 | 13088 | 569 | 457644 | 439 | 120514 | 5453 | 653254 | 412 |

Finally, we also examined the proportions of each TE order in the annotations produced by the raw, MCHelper, and reference libraries. EDTA raw libraries consistently annotated a higher proportion of TE orders compared both to MCHelper and raw libraries across genomes (Figure 6 and Supplemental Table S8). However, in *Z. mays* the number of nucleotides annotated as TEs was equal to the number of nucleotides in the assembly, suggesting that annotations with this library overestimates the number of TEs. For RM2 and REPET, the proportions of TE orders were higher in annotations performed with the MCHelper libraries compared with annotations performed with raw libraries except for REPET in *Z. mays* (Figure 6 and Supplemental Table S8). In *D. melanogaster*, the proportion of TE orders obtained with the MCHelper library was even higher than the one obtained with the reference-BDGP library and more similar to the MCTE library, suggesting that the MCHelper produces a high quality annotation. In *O. sativa*, the TE order proportions were very similar when comparing the MCHelper and the raw RM2 library, while the proportions were smaller for the raw REPET library, consistent with the lower number of copies detected by REPET in this species (Figure 4A) (39.2% REPET vs 48.1% standard library). In *C. corvix*, the manually curated library annotated the lowest proportion of TEs, and only from two orders, LTR and LINE. This result suggests that although the *C. corvix* manually curated library is of high quality, it is probably incomplete. In *D. rerio*, we also found that the raw library from REPET annotated lower TE proportions compared with MCHelper library and the reference library (34.7%, 45.7% and 54.4%, respectively). In *Z. mays,* the MCHelper library processed from REPET annotated virtually the same proportion as the reference (85% REPET+MCHelper vs 84.7% reference).

**Figure 6.**
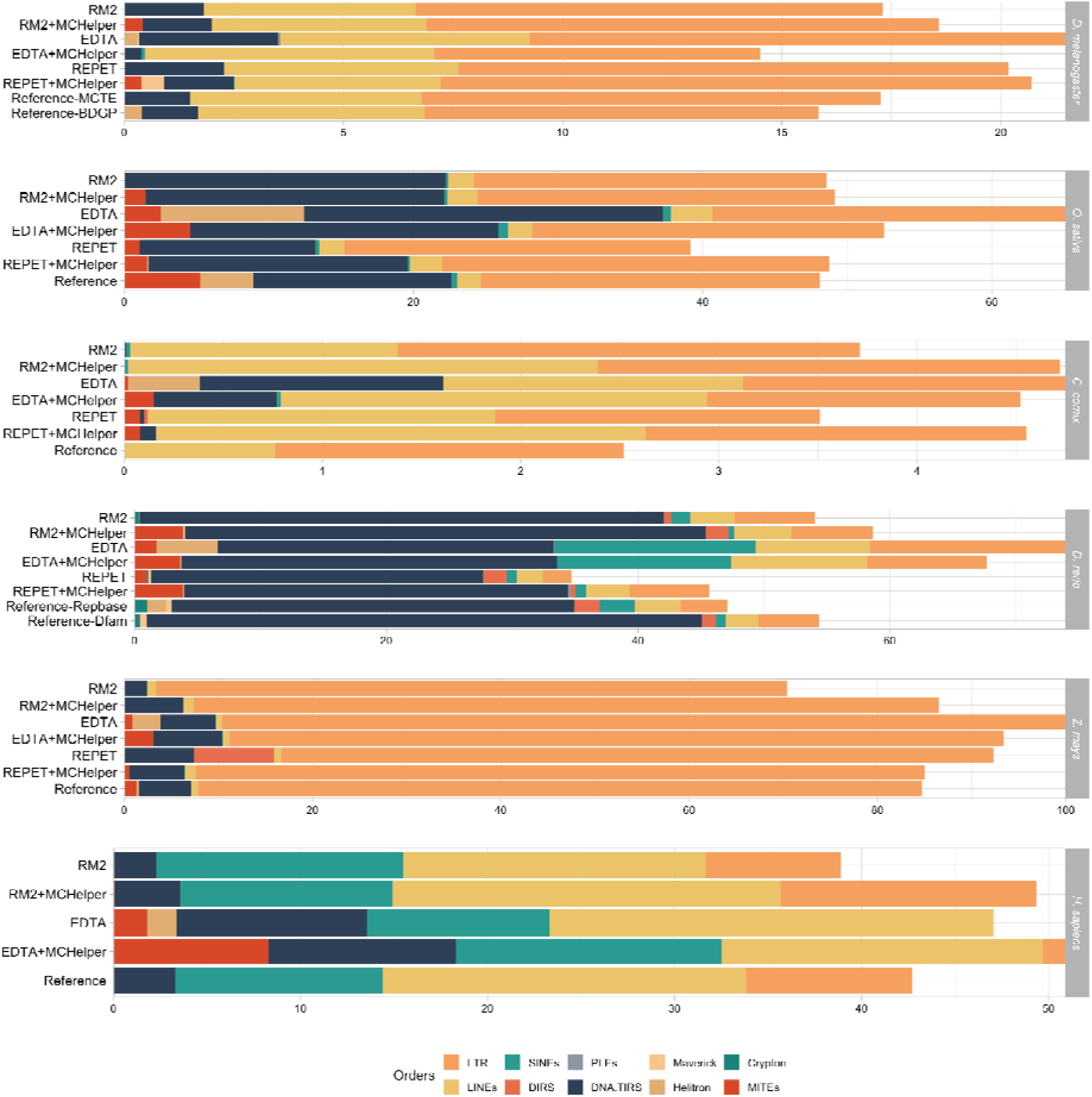
Genomic proportions of TE orders annotated by the raw, MCHelper and reference libraries. Note that the proportion of each genome labeled as “Unclassified/Unknown” TEs is not shown to ease the comparison of TE order proportions across libraries. Annotation with the EDTA library clearly overestimates the amount of TEs in *Z. mays* which could be due to a incomplete defragmentation of TE copies.

Overall, MCHelper libraries produce longer and less fragmented annotations than raw libraries for the majority of tested genomes and *de novo* tools. Importantly, and as expected, although the number of consensus sequences in the MCHelper libraries was smaller compared with the raw libraries (Figure 2A), the proportions of TE orders annotated with the MCHelper libraries were closer to the ones produced with the reference libraries (Figure 6).

### MCHelper curates TE libraries in hours for small genomes and it is easy to install and run

Besides improving the completeness and accuracy of TE libraries, another main goal of MCHelper was to reduce the required time to curate TE sequences. To test the MCHelper runtimes, we executed MCHelper ten times for the raw library of the four species with genomes < 2Gb produced by the three *de novo* tools used in this study, on a server with 48 CPUs. We measured the execution time for ten distinct processes included in the MCHelper tool (see Methods). On average, MCHelper took between 1.10 (*D. melanogaster* - EDTA) and 41.95 (*D. rerio* - EDTA) hours (Table 2). In all cases, the extension step was the most time-consuming: up to 95.91% of the total MCHelper execution time for the *D. rerio* when using the RM2 library. For the two T2T genomes analyzed in this work, the running times of MCHelper ranged between 105,26 and 334,2 hours using 100 cores.

**Table 2.**
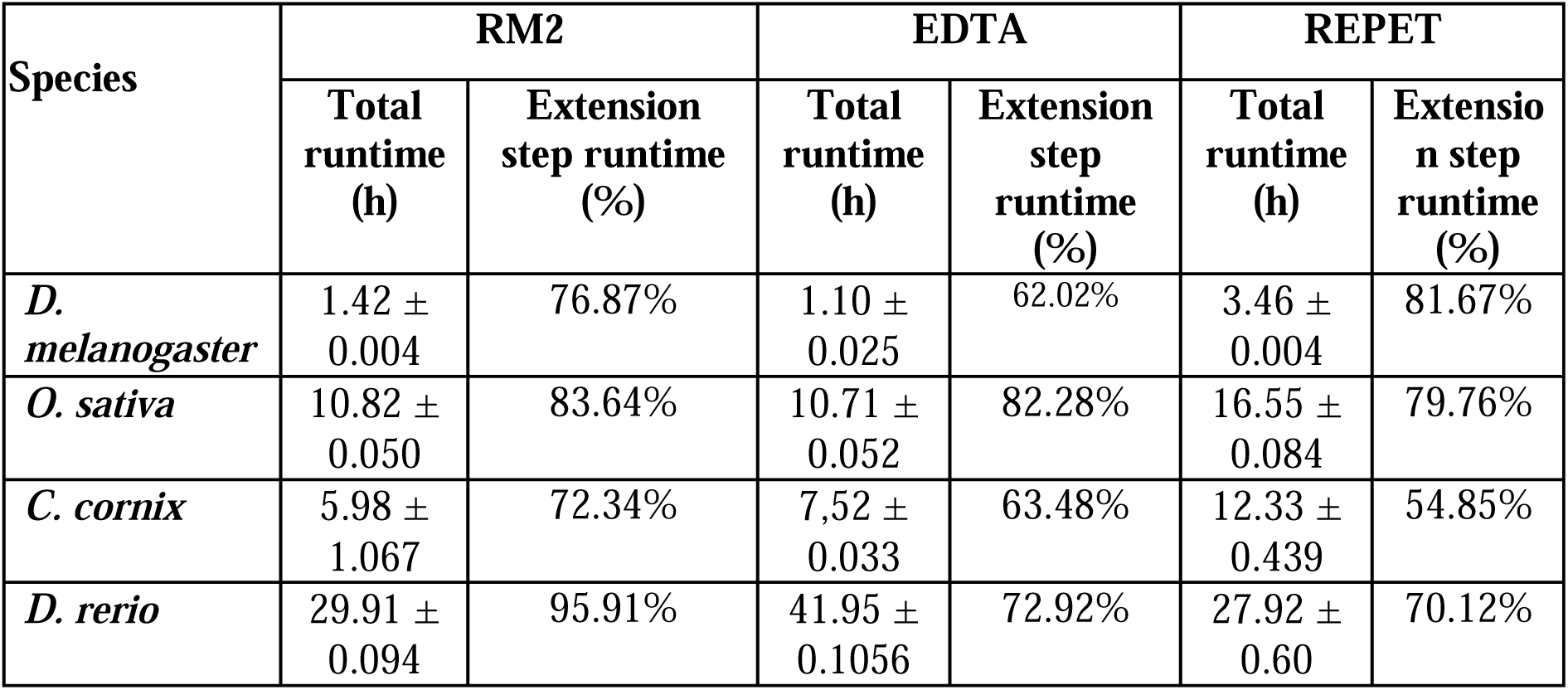
MCHelper average runtimes for the three species analyzed. MCHelper was executed 10 times for each raw library (RM2 EDTA and REPET) and the average and standard variation (STD) was calculated. Total runtimes are given in hours.

MCHelper is an easy tool to set up, as all the dependencies are available in a YML file that allows them all to be installed in an Anaconda environment using a single command. Also, MCHelper is easy to use because it can be run using a single Python command line with only three inputs: the raw library, the genome assembly, and the BUSCO gene set. As such, this tool can be easily integrated into any automatic pipeline to annotate TEs.

## DISCUSSION

We have shown that MCHelper increases TE libraries accuracy and completeness, producing longer and unfragmented TE genome annotations across eukaryotic species, by integrating and automating TE curation protocols (Figure 1). Libraries generated by *de novo* TE identification tools (raw libraries) overestimate the number of TE copies as they contain TE consensus sequences that are redundant, fragmented, and false positive sequences (Rodriguez & Makalowski, 2022; J. M. Storer et al., 2022). Based on several benchmarking metrics, we showed that MCHelper’s automatically curated libraries for *D. melanogaster*, *O. sativa, C. cornix, D, rerio, Z. mays,* and *H. sapiens* are more accurate and complete compared with raw libraries, and thus more similar to high-quality manually curated (reference) libraries (Figure 2). Moreover, MCHelper tool incorporates a module specifically designed to classify TE sequences initially labeled as “unknown/unclassified” by *de novo* TE identification tools, which allows to recover classifications for up to 48% of these sequences (Figure 3). This is highly relevant as the percentage of unclassified elements in raw libraries can be as high as 93% leading to very incomplete and inaccurate TE annotations (Gilbert et al., 2020; Petersen et al., 2019). MCHelper can be executed in parallel in multiple CPUs taking advantage of currently available supercomputing resources. MCHelper allows to curate raw TE libraries in hours for small genomes, with the Consensus extension module being the more time-consuming step (Table 2). Thus, the number of iterations chosen to run the Consensus extension module will affect the total time needed to run MCHelper. Therefore, we recommend the users to test the appropriate value of this parameter to obtain the best possible results in the shortest possible run time, especially when analyzing a large number of genomes.

Most previous TE curation automation efforts consisted of scripts dealing with individual limitations of the raw libraries that needed to be executed independently by the user for each one of the consensus sequences (Goubert et al., 2022; J. M. Storer et al., 2021). Recently, a pipeline to *de novo* annotate TEs included multiple steps to deal with some specific limitations of raw libraries, in particular the presence of redundant and fragmented consensus (Baril et al., 2024). Machine learning approaches have also been implemented to automatically curate particular TE orders in plants (Orozco-Arias et al., 2021, 2022). Thus, previous attempts at automation were partial and still required substantial time investment by the researcher. On the other hand, MCHelper implements multiple manual curation protocols developed by TE experts that tackle the most common limitations of *de novo* TE identification tools across TE orders and species (Goubert et al., 2022; Jamilloux et al., 2016; Platt et al., 2016; J. M. Storer et al., 2021, 2022).

Besides the fully automatic mode that allows the user to improve library accuracy and completeness in a time-effective manner, MCHelper also provides the user with the possibility of performing manual inspection of the generated libraries through a dedicated module (Figure 1 and Supplemental Fig S1B). This module provides the user with information to detect and remove low-quality TE consensus sequences to further increase the overall quality of the TE libraries (Goubert et al., 2022; Tumescheit et al., 2022). We envision that the automatic mode of MCHelper will be highly useful for research projects that need to annotate TEs in a large number of genomes, while the semi-automatic or fully-manual modes should be implemented to generate TE libraries for groups of species for which there are no reference libraries available yet.

MCHelper, in combination with one of the many *de novo* TE identification tools already available, should be instrumental in substantially increasing the number of genomes with accurate TE annotations, which is currently reduced to a very small fraction of the eukaryotic diversity available. Accurate TE annotations in the increasing number of eukaryotic genomes available are the first step towards understanding the dynamics of these important genome components but also the biology of genomes given their important roles in function, structure and evolution.

## METHODS

### MCHelper workflow

First, MCHelper conducts pre-processing analysis, ensuring file completeness and consistency with Wicker’s classification nomenclature (Wicker et al., 2007). For the raw libraries of RM2 and EDTA (or any other FASTA library), MCHelper replaces DNA order names with TIR, and “Unknown” with “Unclassified”. Moreover, it excludes sequences with classifications not contained in the MCHelper nomenclature list (see Supplementary Methods), as well as satellites and RNAs and creates a file with the list of excluded sequences and the reason of exclusion.

#### Redundancy reduction

MCHelper reduces redundancy by clustering all consensus sequences using either CD-HIT v.4.8.1 (Li & Godzik, 2006) or MeShClust V3 (Girgis, 2022), depending on the user decision, with an identity threshold of 95% and coverage of 98% as suggested by Flutre et al. (2011).

#### Consensus Extension

Consensus sequences are processed using *n* iterations of BLAST, Extract, and Extend (BEE) rounds (16 by default; Baril et al., 2024; Platt et al., 2016; J. M. Storer et al., 2021). MCHelper uses NCBI-BLAST v.2.10.1 with parameter “-evalue 1e-20” based on (Goubert et al., 2022). After each BEE round, consensus families are split into subfamilies, if needed, by applying a clustering approach based on the Kimura two parameter distance on the multiple sequence alignment generated (see Supplementary Methods). All consensus sequences are extended until the sequence reaches both TE ends or until the end of the iterations (see Supplementary Methods).

#### False positive filtering

Consensus sequences with (i) very few full-length fragments (FLF) in the genome (by default one FLF but this parameter can be defined by the user; Jamilloux et al., 2016); (ii) having homology with multi-copy genes or rRNAs; and (iii) containing simple sequence repeats (SSR) (by default 60% also adjustable by the user) are considered false positive consensus sequences and filtered out (see Supplementary Methods; Storer et al., 2021, 2022). For genes and rRNAs searching, MCHelper uses hmmscan from HMMER (Eddy, 2011) v.3.3.2 with the following parameters “-E 10 --noali” and the rest by default.

#### Homology based TE identification

Once false positives are filtered out, the remaining sequences are searched against a curated TE database, and those with an 80-80-80 hit (at least 80bp long with 80% sequence identity over 80% of their length; Wicker et al., 2007) are saved in the final curated library (see Supplementary Methods). Those that do not fulfill the 80-80-80 rule are divided into classified and unclassified sequences and follow two distinct paths (Figure 1). MCHelper uses NCBI-BLAST v.2.10.1 to perform this search.

#### Structural Checking

For classified sequences, MCHelper looks for structural information: presence of protein domains, terminal repeats (TRs), and poly(A) tails, to check if the original classification of the consensus sequence matches the expected TE features (see Supplementary Methods; Goubert et al., 2022). In the MCHelper semi-automatic mode, those sequences that present the expected features based on their classification are kept, and the others are manually inspected in the Manual Inspection Module (see Supplementary Methods).

#### TE classification

For “unclassified” elements, MCHelper tries to infer the correct classification using three steps: homology (using 70-70-70 hits), coding domain presence (*e.g.* transposase and reverse transcriptase), and terminal repeats (LTRs and TIRs) (see Supplementary Methods).

#### Manual inspection

MCHelper allows the user to manually visualize useful information to decide if a consensus sequence should be kept, removed or re-classified. All the graphical information is generated by TE+Aid v.1.0 (Goubert et al., 2022) except the multiple sequence alignment plot that is generated by CIAlign v.1.0.18 (Tumescheit et al., 2022). Finally, structural information needed for manual inspection is provided through the terminal (see Supplementary Methods).

### Genome assemblies used

We evaluated the performance of the MCHelper tool on six different species with high-quality manually curated TE libraries: *D. melanogaster*, *O. sativa*, *C. cornix, D. rerio, Z. mays, and H. sapiens*. The assemblies used were GenBank assemblies GCA_002050065.1 for *D. melanogaster*, GCF_000738735.6 for *C. cornix*, and GCA_000002035.4 for *D. rerio*, the RGAP assembly version 7 for *O. sativa* (available at https://phytozome-next.jgi.doe.gov/info/Osativa_v7_0; Ouyang et al., 2007), and the T2T assemblies GCA_022117705.1 for *Z. mays*, and GCF_009914755 for *H. sapiens*. These species were selected based on their varying genome sizes: *D. melanogaster*: 180 Mb (Ashburner et al., 2005; Berlin et al., 2015); *O. sativa*: 466 Mb (Kawahara et al., 2013; Yu et al., 2002); *C. corvix:* 1.1 Gb (Weissensteiner et al., 2020); *D. rerio*: 1.7 Gb (Howe et al., 2013); *Z. mays*: 2.3 Gb (Schnable et al., 2009); and *H. sapiens*: 3.3 Gb (Nurk et al., 2022) respectively), and differences in TE content: ∼20% in *D. melanogaster* (Mérel et al., 2020; Sicat et al., 2022), 46.64% in *O. sativa* (Ou et al., 2019) 59.5% in *D. rerio* (Chang et al., 2022), 85% in *Z. mays* (Schnable et al., 2009), and 47.68% in *H. sapiens* (Hoyt et al., 2022).

### Source of the raw libraries

To obtain the raw TE libraries, we utilized three *de novo* tools: RepeatModeler2 (RM2; Flynn et al., 2020), EDTA (Ou et al., 2019) and *TEdenovo* from REPET (Flutre et al., 2011). For the RM2 raw libraries, we employed those available in its repository (https://github.com/jmf422/TE_annotation/tree/master/benchmark_libraries/RM2) for *D. melanogaster, O. sativa and D. rerio*. For *C. cornix, Z. mays and H. sapiens* the raw libraries were generated using the TEtools Docker image available at https://github.com/Dfam-consortium/TETools, with the assemblies mentioned in the previous section. With respect to the EDTA libraries, we generated them for the six species using the EDTA pipeline v.2.0 with -- anno 0 and the other parameters by default and the same assemblies that were used for the RM2 library generation. Regarding the REPET libraries, we used the same assemblies that were used for the two previous tools. For the *D. melanogaster* genome, we ran the *TEdenovo* pipeline using default parameters and followed the recommended steps in the user’s guideline (https://urgi.versailles.inra.fr/Tools/REPET/TEdenovo-tuto). For the *D. rerio* genome, we created a subset of 300 Mb using the PreProcess.py script available with the REPET package and then we ran *TEdenovo* pipeline using default parameters as for the *D. melanogaster* genome (Jamilloux et al., 2016). The REPET group kindly provided us with the output generated with the *O. sativa* genome available in the RepetDB database (Amselem et al., 2019) and with the libraries for *C. cornix* and *Z. mays*.

### Source of the reference libraries

As the reference libraries, we used the Berkeley Drosophila Genome Project (BDGP) dataset (Kaminker et al., 2002) and the Manual Curated TE library (MCTE, Rech, et al., 2022) for *D. melanogaster*, the library published by Ou et al. (2019) for *O. sativa* (referred to as “standard library” by the authors), the MClibrary available at (Weissensteiner et al., 2020) for *C. cornix,* the manual curated TE models available in Dfam release 3.7 (J. Storer et al., 2021) and in Repbase version 20181026 for *D. rerio*, MTEC curated by Ou et al (https://github.com/oushujun/MTEC) for *Z. mays*, and curated TE models in Dfam 3.7 for *H. sapiens*. For the rice “standard library”, Dfam and Repbase libraries, we unified the LTR sequences with their corresponding internal part before performing further analysis, using in-house scripts.

### Generation of the MCHelper libraries

MCHelper v.1.7.0 was executed with default parameters (to see customizable parameters see GitHub: https://github.com/GonzalezLab/MCHelper). In the False positive filtering step, MCHelper requires a gene set to detect homology between the consensus sequences and multicopy gene families. We used the following BUSCO gene sets: for *D. melanogaster*, we utilized the diptera set (diptera_odb10), for *O. sativa* and *Z. mays*, the viriplantae set (viriplantae_odb10), for *C. cornix*, the aves set (aves_odb10), for *D. rerio*, the actinopterygii set (actinopterygii_odb10), and for *H. sapiens*, the mammalian set (mammalia_odb10). To accommodate MCHelper’s expectation of a single file with all the hidden Markov models (HMMs), we concatenated all the available HMM files into a single file for each species. Note that all the libraries used and generated in this work are available at our institutional repository (http://hdl.handle.net/10261/362092). The genome assemblies and the genome annotations are also available at our institutional repository (http://hdl.handle.net/10261/362092).

### Benchmarking metrics used

At the library level, we used different metrics to benchmark the quality of the consensus models in the raw, MCHelper, and reference libraries used. We divided those metrics into two distinct categories: completeness and accuracy, based on the criteria of a “high quality library” (Goubert et al., 2022; Jamilloux et al., 2016; J. M. Storer et al., 2021, 2022). Metrics used to measure completeness were: the total number of consensus sequences, family quality evaluation, and TE order length distribution. The family quality evaluation consists of defining perfect, good, present and missing families, as follows: Perfect families are those that have a match with one sequence in the analyzed library with >95% nucleotide similarity and >95% length coverage. Good families are those that have multiple overlapping matches in the analyzed library with >95% nucleotide similarity and >95% coverage. Present families are the same as Good Families, but with >80% similarity and >80% coverage. Below those thresholds, the families are considered Missing (Flynn et al., 2020; Rodriguez & Makalowski, 2022). We used the script developed by Flynn et al. (2020) and available at https://github.com/jmf422/TE_annotation/ to classify the consensus sequences.

To measure accuracy, we used the number of false positive sequences as calculated in the False Positive filtering step of MCHelper (see Supplementary Methods). Briefly, we calculated the number of consensus sequences with >60% of their sequence length corresponding to single sequence repeats (SSR), and the number of consensus sequences with hits with BUSCO genes and rRNAs.

To benchmark the quality of the annotations, we ran RepeatMasker v.4.1.2-p1 (Smit et al., 2015) on each of the six genomes with the raw, reference and MCHelper libraries using the -lib parameter, and the following additional parameters: -gff -nolow -no_is -norna. We then used the OneCodeToFindThemAll script (Bailly-Bechet et al., 2014) in each genome annotation to defragment the copies, and we used the “--strict” parameter for *D. melanogaster, D. rerio,* and *C. cornix*. The output of this script was used to estimate the benchmarking metrics. We used the following groups of metrics: quantity of copies, lengths of copies, and genomic proportions occupied by the annotated copies. The metrics used for the quantity of copies were: the total number of annotations, the number of short annotations (< 100 pb), and the number of overlapping annotations (*i.e.,* copies annotated by two or more TE consensus sequences). To define overlapping copies, we considered those that contained any bases annotated with more than one consensus sequence. We used BEDTools merge from the BEDTools suite version 2.30 (https://bedtools.readthedocs.io/en/latest/) with parameters “-d -1 -c 9,9 -o count,collapse” to merge the overlapping copies, and we counted the number of merged copies that came from more than one consensus sequence. To benchmark the length of copies, we used the NTE50 and the LTE50. To calculate these two metrics, we first ordered the copy lengths from longest to shortest and then calculated a cumulative sum of the copy lengths until we reached 50% of the total TE content. The NTE50 is analogous to the N50 metric, used in assembly quality checks, and corresponds to the number of the largest copies needed to annotate 50% of the mobilome. Lower NTE50 values are indicative of longer annotations suggesting that the annotation has captured large unfragmented TE insertions. LTE50 corresponds to the length of the TE annotation at the point in the ordering that 50% of the mobilome is annotated. Higher LTE50 indicates better annotations (Jamilloux et al., 2016). Note that for *Z. mays*, NT50 and LTE50 were calculated for 50% of the full genome space instead of for 50% of the mobilome. Finally, we also calculated the TE order genomic proportions as the percentages of nucleotides that were annotated with TE consensus sequences for each one of the orders analyzed. We obtained the genomic proportions directly from the outputs of the *OneCodeToFindThemAll* script, which sums up all the bases corresponding to each TE order divided by the total assembly size. All scripts used to calculate the metrics described above are available at Supplemental Code File and at https://github.com/GonzalezLab/MCHelper/tree/main/benchmarking_scripts.

### Comparisons with classification tools

In order to compare the MCHelper efficiency when classifying unknown TEs, we processed the “unclassified/unknown” TEs from the RM2 libraries from four of the species analyzed in this study (*D. melanogaster, O. sativa, C. cornix* and *D. rerio*), and then we followed two different approaches to try to assign a classification. First, we manually inspected all the consensus sequences to infer the classification based on the observed structure. Second, we used cross_match with a substitution matrix specially trained for non-coding sequences available in RepeatMasker (named 25p41g.matrix) and against the Repbase library version 20181026 to infer the classification based on homology. Then, we create a subset of the sequences which we inferred the classification based on either structure or homology and we took as ground true the classification inferred. Using this subset as input, we executed TEClass2 (Bickmann et al., 2023), TERL (da Cruz et al., 2021) and DeepTE (Yan et al., 2020) to predict the classification of the sequences. TEClass2 was run using the web GUI (https://www.bioinformatics.uni-muenster.de/tools/teclass2/index.pl?), TERL was executed with the parameter “-m Models/DS3/“ to indicate the model to be used and the rest of parameters by default. Finally, DeepTE was run using the Metazoans_model for *D. melanogaster,* Plans_model for *O. sativa*, and Others_model for *C. cornix* and *D. rerio.* Finally, we calculated three metrics to evaluate the performance of the three tools together with MCHelper: Recovering proportion, overlapping proportion, and precision. We defined the recovering proportion as the proportion of the sequences in the subset recovered by each tool, without taking account the classification predicted. The overlapping proportion was defined as the proportion of the sequences in the subset that was recovered by each tool and for which the classification was correctly predicted at the order level. And the precision was defined as the proportion of the recovered sequences by each tool that was correctly classified.

### Comparison with SENMAP

In order to compare the performance of MCHelper with another curation algorithm, we ran the SENMAP neural network through the script available at: https://github.com/simonorozcoarias/SENMAP. SENMAP was trained only with plant LTRs and its objective is to detect whether an element is complete or not. For this purpose, we used an animal (*D. melanogaster*) and a plant (*O. sativa*) and calculated the number of elements in the final library of both SENMAP (filtering out those that the network detects as incomplete) and MCHelper. In addition, we calculated how many elements were classified as perfect, good, present and missing families.

### Run-time tests

To test the time required to curate TE libraries, we executed MCHelper in the fully automated mode, using 16 extension iterations extending 500 bases at each iteration, 48 CPUs and the default values of other MCHelper parameters. We ran the software ten times to see the variability of the execution times and we calculated the average and the standard deviation of the ten executions. Besides measuring total time, we calculated the execution time of ten subroutines: pre-processing, extension (all the iterations), FLF filtering, False Positive filtering, TE feature calculation, classification assignment for the classified consensus sequences, FLF filtering, TE feature calculation, classification assignment by homology for the unclassified consensus sequences, and order inferred from structural features also for the unclassified consensus sequences. All the executions were performed in a dedicated computing node with 64 CPUs and 240 Gb in RAM.

## SOFTWARE AVAILABILITY

All the in-house scripts used in this manuscript as well as the MCHelper source code is available at the Supplemental code file and at https://github.com/GonzalezLab/MCHelper.

## ACKNOWLEDGEMENTS

We thank the REPET team for providing the *O.sativa, C. cornix* and *Z. mays* REPET libraries. This work was supported by grant PID2020-115874GB-I00 funded by MICIU/AEI /10.13039/501100011033 and by grant 2021 SGR 00417 funded by Departament de Recerca i Universitats, Generalitat de Catalunya awarded to J.G. P.S. is funded by the European Union’s Horizon 2020 research and innovation programme under the Marie Skłodowska-Curie grant agreement No 956229. R.D. acknowledges funding from Wellcome grant 207492.

S. O.-A.: designed research, performed research, analyzed data, contributed analytical tools, wrote the paper. P.S.: contributed analytical tools. R.D.: designed research. J.G.: designed research, analyzed data, wrote the paper. All authors reviewed and commented on the submitted manuscript.

