## Supplemental Material for "MCHelper automatically curates transposable element libraries across eukaryotic species"

**Table of contents:**

**Pages 2-5:** Supplemental Figures

**Pages 6-17:** Supplemental Methods

**Supplemental Figure S1. MCHelper modules that can also be run independently.** MCHelper is composed of three modules that can be executed all in a single run or independently (through the -r option). A) Steps of the “consensus extension” module. B) Steps of the “manual inspection” module. C) Steps and default parameters of the “TE classification” module. When the “TE classification” module is executed independently from the main MCHelper flow, the user's TE library will be extended and filtered at the beginning of the module.

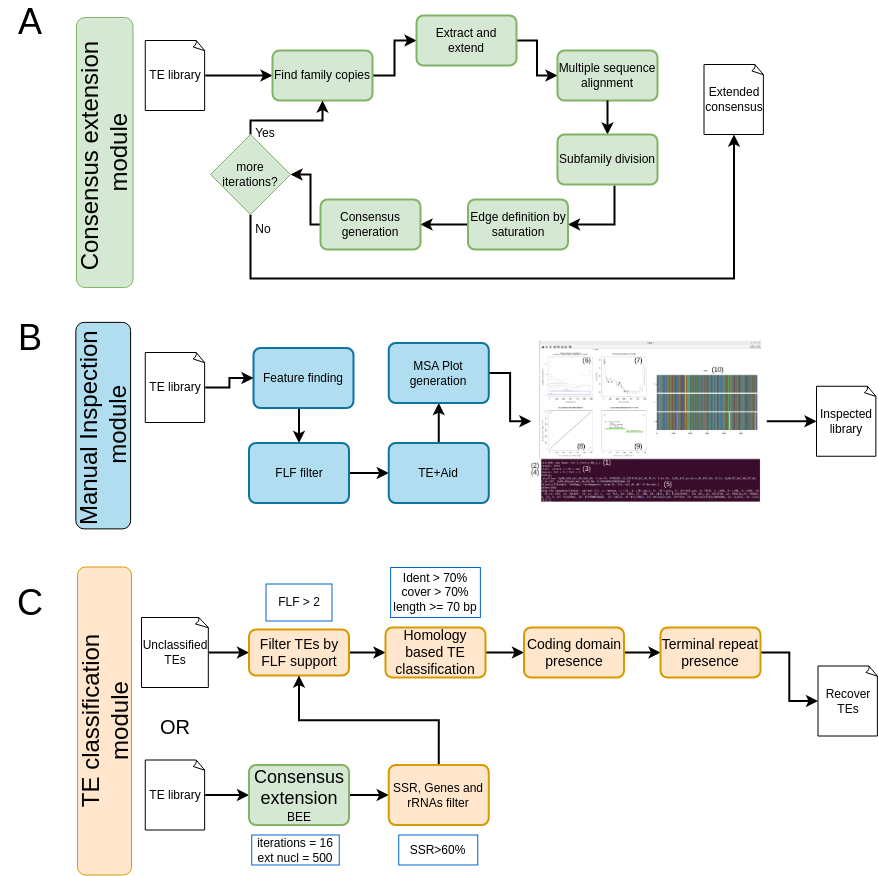

**Supplemental Figure S2. Comparison of the clustering algorithms implemented in MCHelper.**

Number of overlapping consensus sequences for libraries obtained when MCHelper is run using CD-HIT and MeShClust V3 algorithms.

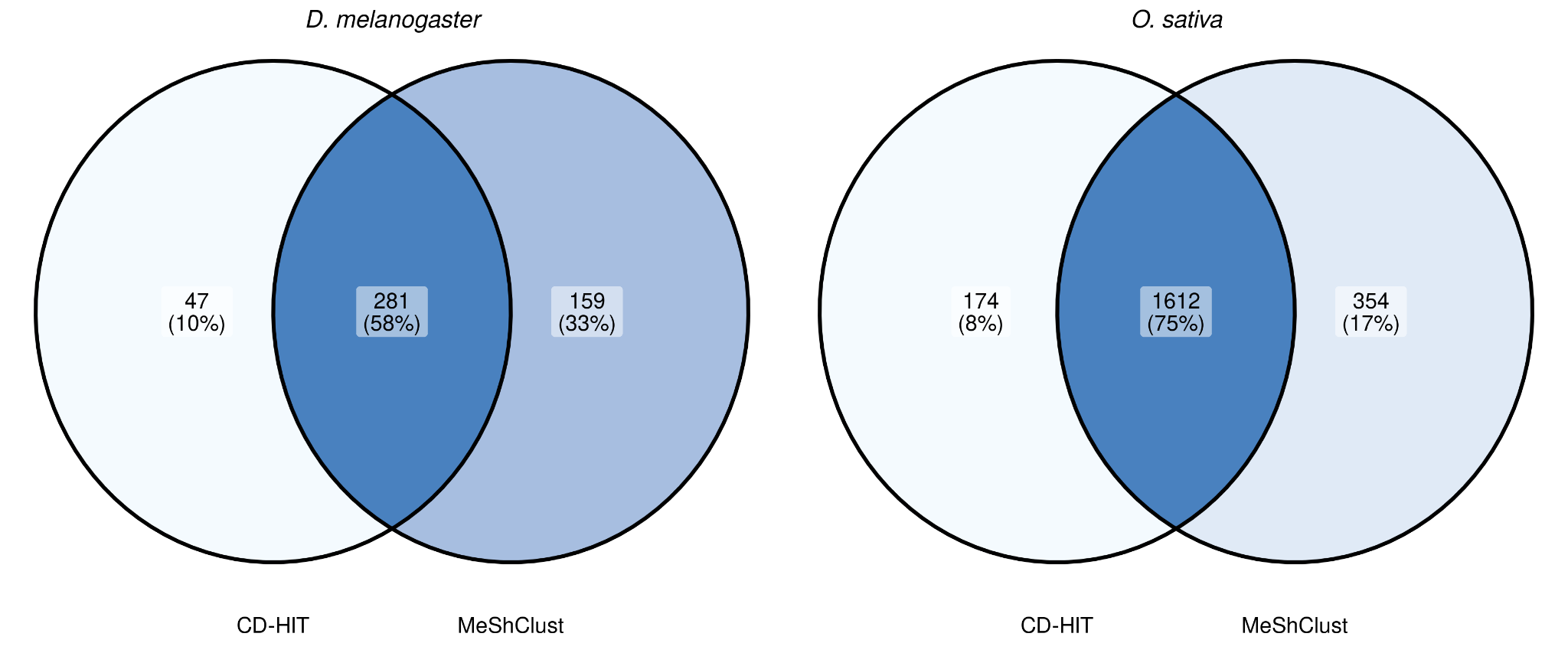

**Supplemental Figure S3.** **Number of false positives consensus sequences found in each library.**

We consider as false positives those consensus sequences that i) contain Simple Sequence Repeats (SSR) in more than 60% of its length; and/or ii) have homology with BUSCO genes; and/or iii) have homology with rRNAs.

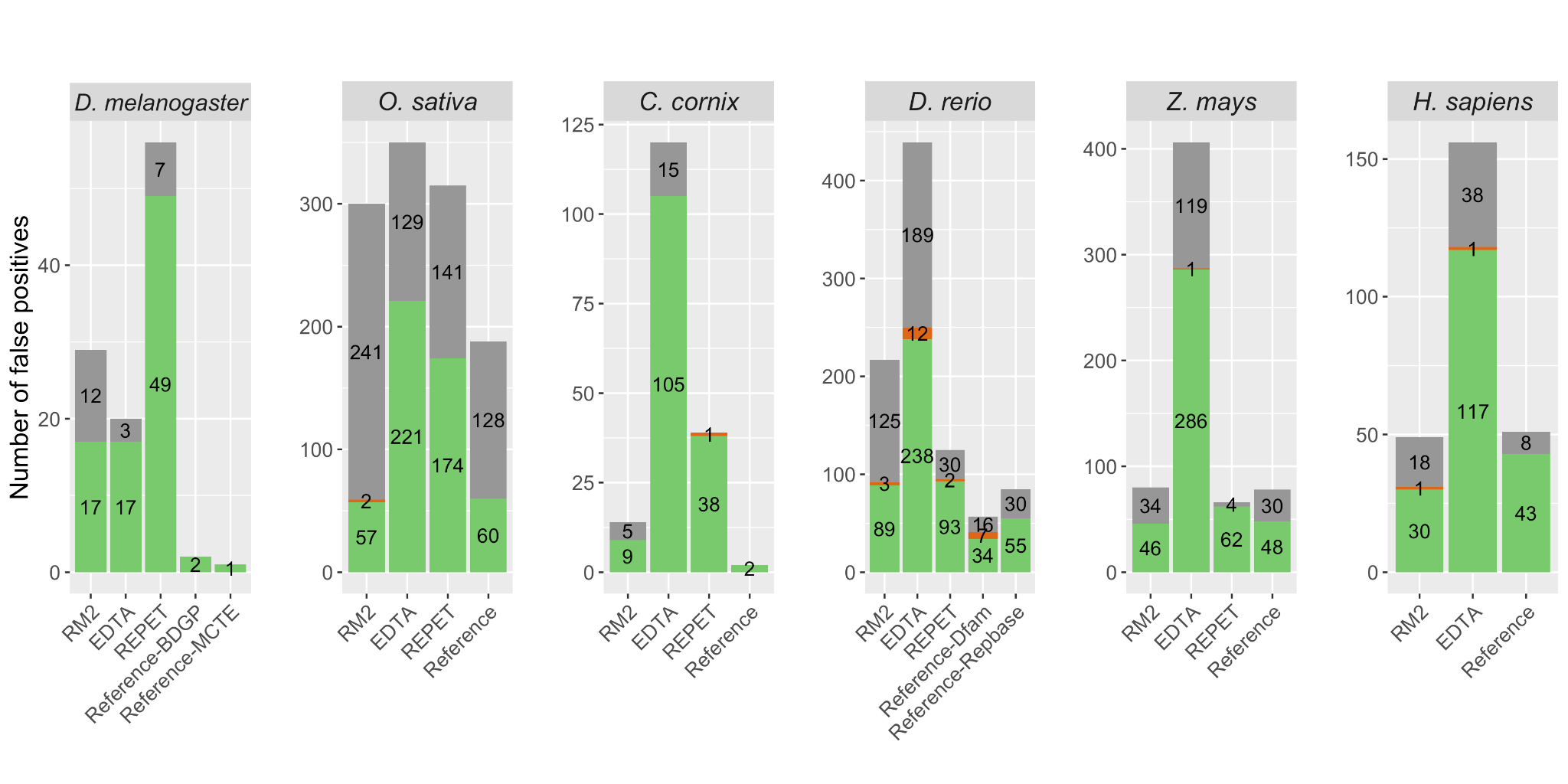

**Supplemental Figure S4. Consensus length distribution of each TE order in the six species analyzed.**

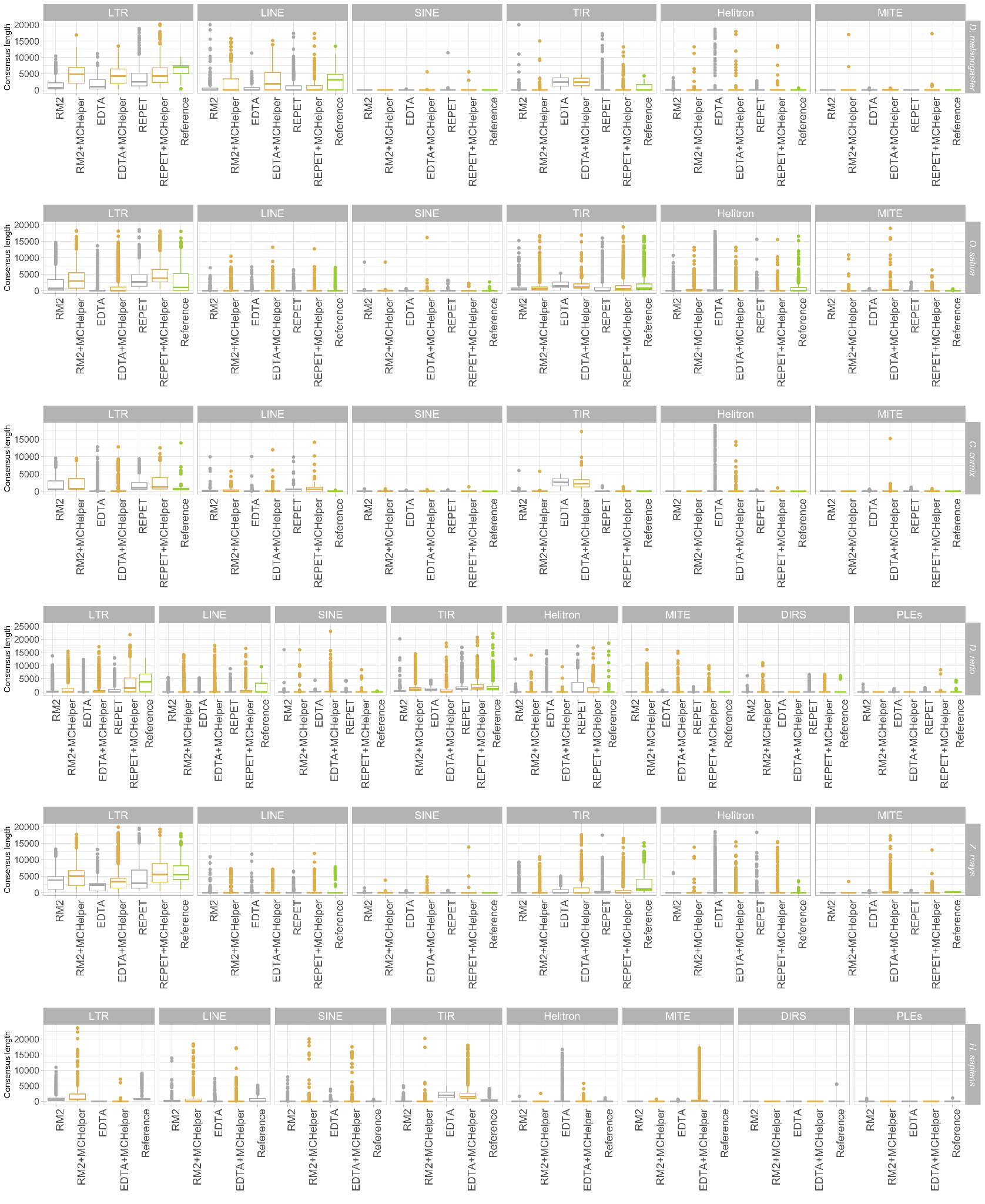

**Supplemental Methods**

**Consensus extension module**

Because one of the most common issues of the *de novo* tools is the fragmentation of the consensus, MCHelper applies a method to extend the consensus sequences based on the iterative process of Blast, Extract and Extend (BEE) (Baril et al., 2022; Goubert et al., 2022; Platt et al., 2016). After each BEE iteration, MCHelper splits multiple sequence alignments (MSA) into sub-groups to generate consensus sequences that better represent the evolutionary history of each TE family/subfamily. To do this, MCHelper applies an automatic clustering approach directly on the MSA using the DBSCAN algorithm (Ester et al., 1996) implemented in scikit-learn v.1.0.2 (Pedregosa et al., 2011) and based on a kimura two parameters distance between the aligned sequences (Kimura, 1980). MCHelper utilizes two parameters to define the clustering: the minimum number of sequences that a cluster can include, and a value ε, which is the minimum distance at which one sequence has to be from another to belong to the same cluster. We select the smallest value of ε that produces the largest number of “stable” clusters. We consider as stable clusters those which remain the same for an interval of ε values of at least 0.06, that is, the number of clusters remains the same for all values of ε on an interval of length ≥ 0.06. This is an empirical value that was the smallest to be able to ignore spurious clusters that can be produced if a very specific value of ε is chosen. This step allows MCHelper to generate multiple sub-alignments and treat each one separately in the following steps.

The next step is to find the 5’ and 3’ ends of the candidate TE, or the closest ends possible to generate a good consensus to keep extending the sequence. Here, MCHelper does not use prior information about the TE order, as this could create a bias in the case of small, partial sequences erroneously assigned to a classification different from the actual one of the full element. Instead, MCHelper determines which are the leftmost and rightmost columns on each alignment that appear to belong to the consensus. This is determined by assessing if the aligned column has a similar number of bases forming the consensus as the rest of columns belonging to it, and, in the case that it is separated from the main group of bases forming the consensus, if it is due to just some insertion on one of the sequences or if it is a false positive Taking this into account the process to decide the limits of the consensus are the following:

i) considering that the bases on the alignment can be separated in two groups, those belonging to the consensus and those not belonging to it, MCHelper calculates the number of sequences that form the plurality on each base position, and select as those putatively belonging to the consensus the ones which have a plurality larger than the average - 2 sigma of the distribution of those bases with a plurality >0.4. This allows us to have a variable cut off point that takes into account the divergence of the sequences in the alignment, which can be larger for older elements.

ii) We used a second heuristic to confirm that the first and last base positions selected previously belong to the actual TE candidate sequence. As we extend some hundreds of bases the sequences, when the number of sequences is small it is expected that some bases achieve randomly the required plurality. For example, for a standard plurality requested on the TE sequence of 0.8, and for 10 sequences, extending 500 bases to each side, in the worst case we still have a 57% of having a false positive. That value is already < 1% for 16 sequences. To remove those false positives we consider that, unlike the real positives of the TE sequence, they will be distributed randomly and require that for low numbers of sequences a variable number of consecutive bases forming a consensus is needed to consider it a valid edge. With this process we avoid extending further away from the end of the putative TE, at the risk of sometimes losing one valid base from the edge.

Once we have obtained the position of the bases where to trim the consensus, the last step is to perform a check to determine whether those ends are the actual end of the putative TE, or just the temporal termination point due to the alignment not capturing the full element. The check requires that the section between the start (or the end) of the consensus and the start (or the end) of the alignment includes a number of columns/bases, in which several sequences (rows) have data that is not aligned (to avoid the distance to be produced by a single outlier insertion).

The algorithm iterates these steps until the edges of the consensus being dealt with have been found, or until a certain number of extensions have been performed (16 by default in MCHelper).

**False positive filtering step**

As part of the MCHelper curation, a false positive discovery and removal step is performed. MCHelper detects three types of false positive sequences: (i) artifacts with very few full-length fragments (FLF) in the genome (by default one FLF but this parameter can be modified by the user (Jamilloux et al., 2016)); (ii) sequences having homology with multi-copy genes or RNAs; and (iii) sequences containing simple sequence repeats (SSRs) (by default 60% also adjustable by the user). To detect the number of FLF, MCHelper blasts (NCBI-blastn v.2.5.0+) each consensus sequence against the genome with an e-value of 10e-^8^ and counts the hits with length > 94% of the consensus length. To discover homology between consensus sequences and multi-copy genes or RNAs, MCHelper uses hmmscan from HMMER v.3.3.2 (Eddy, 2011) (http://hmmer.org/) using the parameters "-E 10 --noali". It searches for hits against reference/BUSCO genes (provided by the user) and against a dataset of rRNAs provided by the REPET development team (URGI) named “rRNA Eukaryota databank”. To identify SSR containing sequences, we used the TRF tool (Benson, 1999), using the parameters "2 3 5 80 10 20 15 -h -d" (as implemented in REPET; Flutre et al., (2011)).

**Homology based TE identification step**

To search if a consensus sequence is an already known TE, we created a curated TE database containing sequences from high quality manual curated datasets:

the MCTE library (Rech et al., 2022), Flybase (Gramates et al., 2022), Dfam v.3.6 (Storer et al., 2021), and repbase version 20181026 (Jurka et al., 2005). For the Dfam sequences we filtered those that were not TEs (different types of RNAs, Satellites and TEs without reported order). For Repbase we only kept the sequences belonging to the following orders: LTR, LINEs, SINEs, DNA, and RC (Figure 1). At the end, all the sequences were clustered using CD-HIT v.4.8.1 (Li & Godzik, 2006) with an identity and coverage threshold of 80% to reduce the redundancy between different sources. The final dataset contained 43,967 TE sequences.

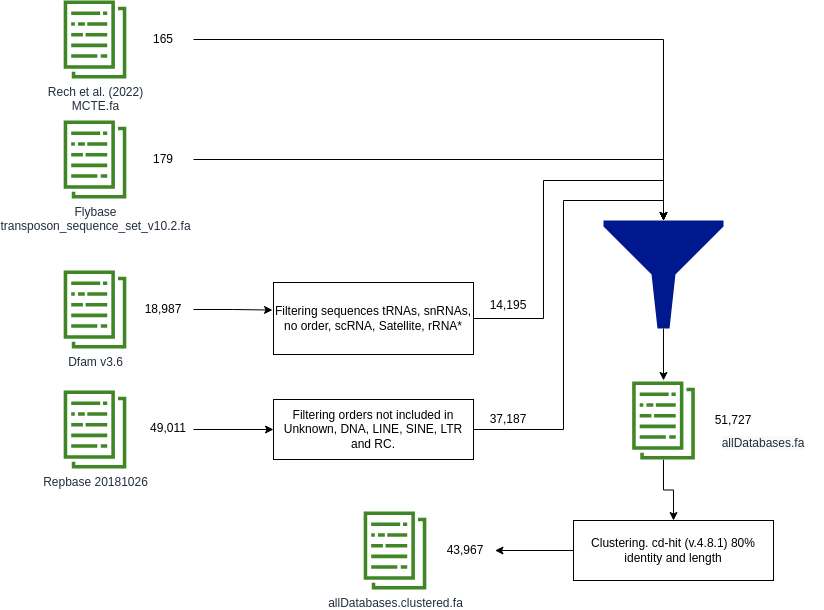

Figure 1. Workflow of the curated TE database used by MCHelper.

Using the curated TE database, MCHelper checks if the consensus sequences from the input library are already reported TEs by blasting (NCBI-blastn v.2.5.0+) them with the following parameters: "-qcov_hsp_perc 80 -perc_identity 80 -max_hsps 1”. MCHelper also checks that the hit is >80 bp to satisfy the 80-80-80 rule (Wicker et al., 2007). Consensus sequences that satisfy the 80-80-80 rule are sent directly to the final TE library produced by MCHelper, while the others continue in the MCHelper workflow.

**Structural checking step**

MCHelper looks for structural information: presence of protein domains, terminal repeats (TRs), and polyA tails. To identify coding domains, MCHelper first extracts the ORFs from the consensus sequences using the getorf tool from EMBOSS v.6.6.0.0 (Rice et al., 2000) with a minimum length of 300 pb (parameter -minsize 300). Then, MCHelper uses hmmscan with parameters “-E 10 --noali” to search for protein HMM profiles from Pfam/GypsyDB databank collected by the REPET development group (available at https://urgi.versailles.inrae.fr/download/repet/profiles/ProfilesBankForREPET_Pfam35.0_GypsyDB.hmm.tar.gz). To identify TRs, MCHelper blasts (NCBI-BLASTN v.2.5.0+) each consensus sequence against itself using the following parameters: -word_size 11 -gapopen 5 -gapextend 2 -reward 2 -penalty -3. Then, it searches for hits larger than 10 bp that are located in the 10% of the two ends (5’ and 3’ ends) of the consensus sequence. MCHelper defines the TRs as LTRs if the two hits (the 5' and the 3') are in the same direction and as TIRs otherwise. If multiple hits satisfy these conditions, the reported TRs are the largest ones. To identify poly(A) tails, MCHelper looks for string patterns with length ≥ 10 pb that match the regular expressions “(.*?)[^A]” or “(.*?)[^T]”, allowing 1 mismatch. Using all these structural features, MCHelper uses decision rules designed for each TE order (Figure 2) to categorize as “complete” sequences those that satisfy all the conditions, and as “incomplete” sequences those that do not match the expected structure according to the initial classification in the raw libraries (Figure 2). To count how many domains are present in a consensus sequence, MCHelper only takes into account those that are supposed to be found based on the initial classification and reports both the number of expected and unexpected domains (Table 1). If at least one unexpected domain is found in the sequence, MCHelper defines it as “incomplete”, because it could be a chimeric consensus sequence. In the case of non-autonomous TEs (SINEs, TRIM, LARD, and MITE), it is also necessary that the consensus sequence has >2 FLF to be considered as “complete”, and will be labeled as “incomplete” otherwise. For “complete” non-autonomous LTR, MCHelper also assigns the TE as a TRIM if the consensus length is < 2,500 bp or as a LARD otherwise. In the fully-automatic mode, MCHelper saves all the sequences in the final curated library, putting the suffix “_inc” to the incomplete ones, while in the semi-automatic, the “incomplete” TEs are shown to the user through the Manual Inspection Module.

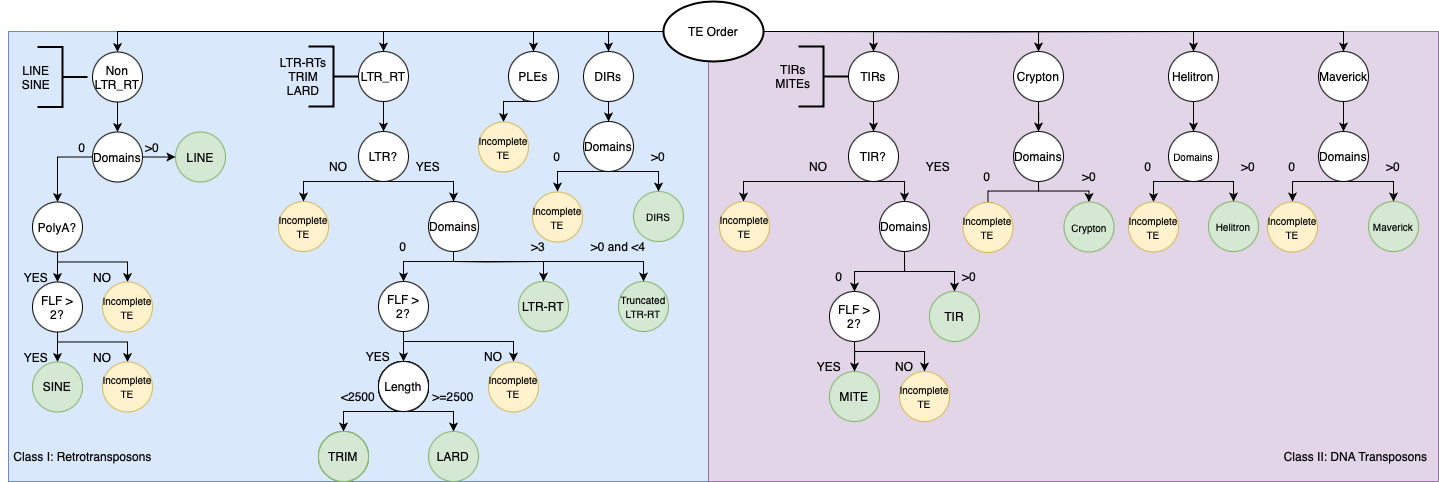

Figure 2. Decision rules to define whether a consensus sequence is a “complete” or “incomplete” TE based on the original classification given by the *de novo* tool, and the features found by MCHelper. PLEs, DIRs and Crypton orders are considered as “incomplete” because of the difficulty to check their structures automatically.

Table 1. Expected domains by TE order. MCHelper searches for the right domains based on the classification and reports as incorrect any other found domain.

| **Order** | **Expected domains** | **References** |
| --- | --- | --- |
| LINE | GAG, RT, EN, RNAseH | (Wicker et al., 2007) |
| LTR | GAG, AP, INT, RT, RNAseH, ENV | (Wicker et al., 2007) |
| PLEs | RT, EN | (Goubert et al., 2022; Mérel et al., 2020; Wicker et al., 2007) |
| DIRs | GAG, RT, RNAseH, YR | (Goubert et al., 2022; Poulter & Butler, 2015; Wicker et al., 2007) |
| TIR | Tase | (Wicker et al., 2007) |
| Crypton | YR | (Poulter & Butler, 2015; Wicker et al., 2007) |
| Helitron | HEL, EN, RPA, REP, OTU, SET | (Thomas & Pritham, 2015; Wicker et al., 2007) |
| Maverick | ATPase, INT*, AP* | (Goubert et al., 2022; Kapitonov & Jurka, 2006; Wicker et al., 2007) |

Abbreviations: GAG: capsid protein; RT: reverse transcriptase; EN: endonuclease; AP: aspartic proteinase; AP*: cysteine protease; INT, integrase; INT*: retroviral integrase; ENV: envelope protein; YR: tyrosine recombinase; Tase: transposase; HEL: helicase; RPA: replication protein A; REP: replicator; OTU: OTU protein; SET: SET auxiliar gene; and ATPase: packing ATPase

**TE classification module**

For the consensus sequences initially labeled as “unknown/unclassified” TEs by the de novo tool, MCHelper applies a three-step process to infer the correct classification: homology, coding domain presence, and terminal repeats. In the homology step, MCHelper first blasts (NCBI-blastn v.2.5.0+) the consensus sequences against the sequences already included in the MCHelper curated library but using a more relaxed threshold of 70% identity, 70% coverage and length >70 bp (parameters -qcov_hsp_perc 70 -perc_identity 70 -max_hsps 1). The remaining sequences are blasted against the curated TE database used in the Homology based TE identification step, using the 70-70-70 rule as well. The consensus sequences with significant hits are assigned the classification of the matched sequence. For the remaining unclassified sequences MCHelper searches for coding domains such as reverse transcriptase, integrase, and transposase to infer the classification in a hierarchical way. First, MCHelper looks if the sequence has domains from Class I or Class II elements, then looks for specific domains inside each order (such as envelope for LTR, transposase for TIRs, or helicase for Helitrons). The remaining sequences are searched for long terminal repeats or terminal inverted repeats to assign either the LTR or the TIR order. Elements re-classified in this last step are marked with the suffix "_unconfirm" since no coding domains were found in the consensus sequences.

**Manual inspection module**

To allow the user to perform visual inspection of the consensus sequences, MCHelper presents the information in two formats through the terminal and through a Graphical User Interface (GUI). In the terminal, the user will find the following information: sequence name, TE length in bp, classification, number of full length fragments in the genome, coding domains and terminal repeats/poly A tail. In the GUI, the user will be able to find the following information: copy divergence to the consensus, TE genomic coverage, TE consensus self-dotplot, TE structure and protein hits, and a multiple Sequences Alignment (MSA) plot. In some cases, MCHelper may display redundant information, such as the number of full-length fragments, terminal repeats, and coding domains. This redundancy is beneficial as it provides additional evidence when one approach fails to provide information. All the graphical information is generated by TE+Aid v.1.0 (Goubert et al., 2022), except by the MSA plot which is generated by CIAlign v.1.0.18 (Tumescheit et al., 2022).

**Classification nomenclature used by MCHelper**

MCHelper uses the initial classification given by the *de novo* tool to check the completeness of the TE. Thus, it is crucial that the raw consensus sequences have TE classification names that correspond to the ones that MCHelper already knows. Nevertheless, there are many classification systems with different nomenclatures that refer to the same or similar groups of elements (Arkhipova, 2017; Kapitonov & Jurka, 2008; Piégu et al., 2015; Wicker et al., 2007). Therefore, MCHelper uses an updated Wicker et al. (2007) nomenclature that includes the following TE classes, orders and superfamilies.

- Class I: Retrotransposons
  - Order: LTR; superfamilies: Copia, Gypsy, Bel-Pao, ERV, TRIM, LARD
  - Order: LINE; superfamilies: CR1, R1, R2, RTE, JOCKEY, L1, L2, LOA, I,
  - Order: SINE
  - Order: PLE
  - Order: DIRS; superfamilies: DIRS, Ngaro, VIPER
- Class II: DNA Transposons
  - Order: TIR; superfamilies: Tc1-Mariner, hAT, Mutator, Merlin, Transib, P, PiggyBac, PIF-Habinger, CACTA, Mule, CMC
  - Order: MITE
  - Order: Crypton
  - Order: Helitron
  - Order: Maverick
- Unclassified
