## Supplemental Code for "MCHelper automatically curates transposable element libraries across eukaryotic species": annot_metrics_paper_v2.sh

```
#!/bin/bash
tool_path=/lustre/home/ibe/sorozco/MCHelper/tools

rm -rf annot_file_results.txt dico.txt

# D melanogaster
work_dir=/lustre/home/ibe/sorozco/Dmel
ref=${work_dir}/reference/RM_ref/dmel-all-chromosome-r6.47.fasta.out
raw=${work_dir}/raw/RM_raw/dmel-all-chromosome-r6.47.fasta.out
cur=${work_dir}/RM_curated_lib/dmel-all-chromosome-r6.47.fasta.out
genome_name=dmel-all-chromosome-r6.47.fasta
genome=${work_dir}/${genome_name}
ref_lib=${work_dir}/reference/D_mel_transposon_sequence_set_v10.2.fa
raw_lib=${work_dir}/raw/Dmel-families.fa
cur_lib=${work_dir}/curation_ite16/curated_sequences_NR.fa

genome_size="137547960"
perc_TEs="20"
threads=48

rm -rf {ref}.* ${ref}_curated.out
rm -rf {raw}.* ${ref}_curated.out
rm -rf {cur}.* ${ref}_curated.out
echo "######## Quantity of copies section ############" >> annot_file_results.txt
########################################## reference #####################################################
# change TIR by DNA to work with OneCodetoFindThemAll as well as Helitron by RC
cat ${ref} | sed 's/Helitron\/HELITRON/RC\/Helitron/g' | sed 's/DNA\/Helitron/RC\/Helitron/g'| sed 's/TIR/DNA/g' | sed 's/Unclassified/Unknown/g' | sed 's/LARD/LTR\/LARD/g' | sed 's/MITE/DNA/g' > ${ref}.format
${tool_path}/Onecodetofindthemall/build_dictionary.pl --rm ${ref}.format > dico.txt
${tool_path}/Onecodetofindthemall/one_code_to_find_them_all.pl --rm ${ref}.format --ltr dico.txt --fasta --strict --unknown
cat ${ref}.format_*.elem_sorted.csv | grep "^###" | sed 's/^###//g' > ${ref}_curated.out

num_cop=`cat ${ref}_curated.out | wc -l`
num_short_cop=`awk '{if($8 < 100){print $0}}' ${ref}_curated.out | wc -l`

awk '{print $5"\tRepeatMasker\tsimilarity\t"$6"\t"$7"\t"$1"\t"$9"\t.\tTarget \"Motif:"$10"#"$11"\" "$12" "$13}' ${ref}_curated.out | bedtools sort | bedtools merge -d -1 -c 9,9 -o count,collapse > ${ref}_curated.out.gff_merge_count.gff
quim=`python3 extract_quimerics_and_non_overlaping.py ${ref}_curated.out.gff_merge_count.gff`

echo "${num_cop}\t${num_short_cop}\t${quim}" >> annot_file_results.txt
########################################## raw #####################################################
cat ${raw} | sed 's/Helitron\/HELITRON/RC\/Helitron/g' | sed 's/DNA\/Helitron/RC\/Helitron/g'   | sed 's/TIR/DNA/g' | sed 's/Unclassified/Unknown/g' | sed 's/LARD/LTR\/LARD/g' | sed 's/MITE/DNA/g' > ${raw}.format
${tool_path}/Onecodetofindthemall/build_dictionary.pl --rm ${raw}.format > dico.txt
${tool_path}/Onecodetofindthemall/one_code_to_find_them_all.pl --rm ${raw}.format --ltr dico.txt --fasta --strict --unknown
cat ${raw}.format_*.elem_sorted.csv | grep "^###" | sed 's/^###//g' > ${raw}_curated.out

num_cop=`cat ${raw}_curated.out | wc -l`
num_short_cop=`awk '{if($8 < 100){print $0}}' ${raw}_curated.out | wc -l`

awk '{print $5"\tRepeatMasker\tsimilarity\t"$6"\t"$7"\t"$1"\t"$9"\t.\tTarget \"Motif:"$10"#"$11"\" "$12" "$13}' ${raw}_curated.out | bedtools sort | bedtools merge -d -1 -c 9,9 -o count,collapse > ${raw}_curated.out.gff_merge_count.gff
quim=`python3 extract_quimerics_and_non_overlaping.py ${raw}_curated.out.gff_merge_count.gff`

echo "${num_cop}\t${num_short_cop}\t${quim}" >> annot_file_results.txt

########################################## curated #####################################################
cat ${cur} | sed 's/CLASSI\///g' | sed 's/CLASSII\///g' | sed 's/HELITRON/RC\/Helitron/g' | sed 's/DNA\/Helitron/RC\/Helitron/g' | sed 's/TIR/DNA/g'| sed 's/Unclassified/Unknown/g' | sed 's/LARD/LTR\/LARD/g' | sed 's/MITE/DNA/g' > ${cur}.format
${tool_path}/Onecodetofindthemall/build_dictionary.pl --rm ${cur}.format > dico.txt
${tool_path}/Onecodetofindthemall/one_code_to_find_them_all.pl --rm ${cur}.format --ltr dico.txt --fasta --strict --unknown
cat ${cur}.format_*.elem_sorted.csv | grep "^###" | sed 's/^###//g' > ${cur}_curated.out

num_cop=`cat ${cur}_curated.out | wc -l`
num_short_cop=`awk '{if($8 < 100){print $0}}' ${cur}_curated.out | wc -l`

awk '{print $5"\tRepeatMasker\tsimilarity\t"$6"\t"$7"\t"$1"\t"$9"\t.\tTarget \"Motif:"$10"#"$11"\" "$12" "$13}' ${cur}_curated.out| bedtools sort | bedtools merge -d -1 -c 9,9 -o count,collapse > ${cur}_curated.out.gff_merge_count.gff
quim=`python3 extract_quimerics_and_non_overlaping.py ${cur}_curated.out.gff_merge_count.gff`

echo "${num_cop}\t${num_short_cop}\t${quim}" >> annot_file_results.txt

echo "######## Length of copies section ############" >> annot_file_results.txt
python3 NTE50_LTE50_FLC_v2.py $genome_size $perc_TEs ${ref}_curated.out ${raw}_curated.out ${cur}_curated.out ${ref_lib} ${raw_lib} ${cur_lib} >> annot_file_results.txt

echo "######## Proportions of copies section ############" >> annot_file_results.txt
LINE=`grep -E "######Type:LINE|######Type:SINE" ${ref}.format_*.copynumber.csv | cut -f7 | awk -v "genome=$genome_size" '{sum=sum+$1}END{print (sum/genome)*100}'`
LTR=`grep "######Type:LTR" ${ref}.format_*.copynumber.csv | cut -f7 | awk -v "genome=$genome_size" '{sum=sum+$1}END{print (sum/genome)*100}'`
Maverick=`grep "###DNA/Maverick" ${ref}.format_*.copynumber.csv | cut -f7 | awk -v "genome=$genome_size" '{sum=sum+$1}END{print (sum/genome)*100}'`
Helitron=`grep "###DNA/RC" ${ref}.format_*.copynumber.csv | cut -f7 | awk -v "genome=$genome_size" '{sum=sum+$1}END{print (sum/genome)*100}'`
Crypton=`grep "###DNA/Crypton" ${ref}.format_*.copynumber.csv | cut -f7 | awk -v "genome=$genome_size" '{sum=sum+$1}END{print (sum/genome)*100}'`
DNA_raw=`grep "######Type:DNA" ${ref}.format_*.copynumber.csv | cut -f7 | awk -v "genome=$genome_size" '{sum=sum+$1}END{print (sum/genome)*100}'`
DNA=`echo "$DNA_raw $Maverick $Helitron" | awk {'print $1-$2-$3'}`
Unclass=`grep "######Type:Unknown" ${ref}.format_*.copynumber.csv | cut -f7 | awk -v "genome=$genome_size" '{sum=sum+$1}END{print (sum/genome)*100}'`
echo "${LTR}\t${LINE}\t0\t0\t${DNA}\t${Maverick}\t${Helitron}\t${Crypton}\t${Unclass}" >> annot_file_results.txt

LINE=`grep -E "######Type:LINE|######Type:SINE" ${raw}.format_*.copynumber.csv | cut -f7 | awk -v "genome=$genome_size" '{sum=sum+$1}END{print (sum/genome)*100}'`
LTR=`grep "######Type:LTR" ${raw}.format_*.copynumber.csv | cut -f7 | awk -v "genome=$genome_size" '{sum=sum+$1}END{print (sum/genome)*100}'`
Maverick=`grep "###DNA/Maverick" ${raw}.format_*.copynumber.csv | cut -f7 | awk -v "genome=$genome_size" '{sum=sum+$1}END{print (sum/genome)*100}'`
Helitron=`grep "###DNA/RC" ${raw}.format_*.copynumber.csv | cut -f7 | awk -v "genome=$genome_size" '{sum=sum+$1}END{print (sum/genome)*100}'`
Crypton=`grep "###DNA/Crypton" ${raw}.format_*.copynumber.csv | cut -f7 | awk -v "genome=$genome_size" '{sum=sum+$1}END{print (sum/genome)*100}'`
DNA_raw=`grep "######Type:DNA" ${raw}.format_*.copynumber.csv | cut -f7 | awk -v "genome=$genome_size" '{sum=sum+$1}END{print (sum/genome)*100}'`
DNA=`echo "$DNA_raw $Maverick $Helitron" | awk {'print $1-$2-$3'}`
Unclass=`grep "######Type:Unknown" ${raw}.format_*.copynumber.csv | cut -f7 | awk -v "genome=$genome_size" '{sum=sum+$1}END{print (sum/genome)*100}'`
echo "${LTR}\t${LINE}\t0\t0\t${DNA}\t${Maverick}\t${Helitron}\t${Crypton}\t${Unclass}" >> annot_file_results.txt

LINE=`grep -E "######Type:LINE|######Type:SINE" ${cur}.format_*.copynumber.csv | cut -f7 | awk -v "genome=$genome_size" '{sum=sum+$1}END{print (sum/genome)*100}'`
LTR=`grep "######Type:LTR" ${cur}.format_*.copynumber.csv | cut -f7 | awk -v "genome=$genome_size" '{sum=sum+$1}END{print (sum/genome)*100}'`
Maverick=`grep "###DNA/Maverick" ${cur}.format_*.copynumber.csv | cut -f7 | awk -v "genome=$genome_size" '{sum=sum+$1}END{print (sum/genome)*100}'`
Helitron=`grep "###DNA/RC" ${cur}.format_*.copynumber.csv | cut -f7 | awk -v "genome=$genome_size" '{sum=sum+$1}END{print (sum/genome)*100}'`
Crypton=`grep "###DNA/Crypton" ${cur}.format_*.copynumber.csv | cut -f7 | awk -v "genome=$genome_size" '{sum=sum+$1}END{print (sum/genome)*100}'`
DNA_raw=`grep "######Type:DNA" ${cur}.format_*.copynumber.csv | cut -f7 | awk -v "genome=$genome_size" '{sum=sum+$1}END{print (sum/genome)*100}'`
DNA=`echo "$DNA_raw $Maverick $Helitron" | awk {'print $1-$2-$3'}`
Unclass=`grep "######Type:Unknown" ${cur}.format_*.copynumber.csv | cut -f7 | awk -v "genome=$genome_size" '{sum=sum+$1}END{print (sum/genome)*100}'`
echo "${LTR}\t${LINE}\t0\t0\t${DNA}\t${Maverick}\t${Helitron}\t${Crypton}\t${Unclass}" >> annot_file_results.txt
```
