## Supplemental Code for "MCHelper automatically curates transposable element libraries across eukaryotic species": asign_families_to_unclass.py

```
import sys
import os
import pandas as pd
from Bio import SeqIO

dicc_orders = {2: "LTR", 3: "Copia", 4: "Gypsy", 5: "BelPao",
               6: "TRIM", 7: "LARD", 8: "LINE", 9: "SINE", 10: "TIR", 11: "MITE", 12: "Helitron", 13: "Maverick",
               14: "PLE", 15: "DIRS", 16: "R2", 17: "RTE", 18: "Jockey", 19: "L1", 20: "I", 21: "Tc1–Mariner",
               22: "hAT", 23: "Mutator", 24: "Merlin", 25: "Transib", 26: "P", 27: "PiggyBac", 28: "PIF–Harbinger",
               29: "CACTA"}

def create_fasta_files(ref_tes, keep_seqs, file_name, outputdir, orders):
    record_curated = []
    create_output_folders(outputdir)

    try:
        for sequence in SeqIO.parse(ref_tes, "fasta"):
            if str(sequence.id) in keep_seqs:
                if len(orders) > 0:
                    pos = keep_seqs.index(str(sequence.id))
                    classification = dicc_orders[orders[pos]]
                    if orders[pos] >= 3 and orders[pos] <= 5:
                        classification = "LTR/" + classification
                    elif orders[pos] >= 16 and orders[pos] <= 20:
                        classification = "LINE/" + classification
                    elif orders[pos] >= 21 and orders[pos] <= 29:
                        classification = "TIR/" + classification
                    sequence.id = sequence.id + "#" + str(classification)
                    sequence.description = ''
                    record_curated.append(sequence)
                else:
                    record_curated.append(sequence)

        SeqIO.write(record_curated, file_name, "fasta")
    except FileNotFoundError:
        print(
            "FATAL ERROR: I couldn't find the files, please check: '" + ref_tes + "' and '" + file_name + "'. Path not found")
        sys.exit(0)
    except PermissionError:
        print(
            "FATAL ERROR: I couldn't access the files, please check: ' " + ref_tes + "' and '" + file_name + "'. I don't have permissions.")
        sys.exit(0)
    except:
        print("FATAL ERROR: There is a unknown problem writing sequences in : '" + outputdir + "'.")
        sys.exit(0)

def create_output_folders(folder_path):
    try:
        if not os.path.exists(folder_path):
            os.mkdir(folder_path)
    except FileNotFoundError:
        print("FATAL ERROR: I couldn't create the folder " + folder_path + ". Path not found")
        sys.exit(0)
    except PermissionError:
        print("FATAL ERROR: I couldn't create the folder " + folder_path + ". I don't have permissions.")
        sys.exit(0)

if __name__ == '__main__':
    proj_name = "Dmel"
    blastn_results1 = "/home/simon/invase_species/100Drosophilas/curation_done_v2/unclassPutative_vs_finalLibV3.blast"
    blastn_results2 = "/home/simon/invase_species/100Drosophilas/curation_done_v2/unclassPutative_vs_allDatabases.blast"
    blastn_results3 = "/home/simon/invase_species/100Drosophilas/curation_done_v2/unclassPutative_vs_finalLibV3.tblastx"
    list1 = "/home/simon/invase_species/100Drosophilas/curation_done_v2/list_blastn_finalLibV3.txt"
    list2 = "/home/simon/invase_species/100Drosophilas/curation_done_v2/list_blastn_allDatabases.txt"
    list3 = "/home/simon/invase_species/100Drosophilas/curation_done_v2/list_tblastx_finalLibV3.txt"

    keep_seqs = []
    orders = []
    openfile = open(list1, "r").readlines()
    seqs_species = [x.replace('\n', '') for x in openfile if proj_name in x]

    blastresult = pd.read_table(blastn_results1, sep='\t', names=['qseqid', 'sseqid', 'pident', 'length',
                                'mismatch', 'gapopen', 'qstart', 'qend', 'sstart', 'send', 'evalue', 'bitscore'])

    for seq in seqs_species:
        lenHit = 0
        order_i = ""
        for i in range(blastresult.shape[0]):
            if blastresult.loc[i, 'qseqid'] == seq:
                if lenHit < blastresult.loc[i, 'length']:
                    lenHit = blastresult.loc[i, 'length']
                    order_i = blastresult.loc[i, 'sseqid'].split("#")[1]
        print(lenHit)
        print(order_i)
        orders.append(order_i)
        keep_seqs.append(seq)

    openfile = open(list2, "r").readlines()
    seqs_species = [x.replace('\n', '') for x in openfile if proj_name in x]
    blastresult = pd.read_table(blastn_results2, sep='\t', names=['qseqid', 'sseqid', 'pident', 'length',
                                                                  'mismatch', 'gapopen', 'qstart', 'qend', 'sstart',
                                                                  'send', 'evalue', 'bitscore'])

    for seq in seqs_species:
        lenHit = 0
        order_i = ""
        for i in range(blastresult.shape[0]):
            if blastresult.loc[i, 'qseqid'] == seq:
                if lenHit < blastresult.loc[i, 'length']:
                    lenHit = blastresult.loc[i, 'length']
                    order_i = blastresult.loc[i, 'sseqid'].split("#")[1]
        print(lenHit)
        print(order_i)
        orders.append(order_i)
        keep_seqs.append(seq)

    openfile = open(list3, "r").readlines()
    seqs_species = [x.replace('\n', '') for x in openfile if proj_name in x]
    blastresult = pd.read_table(blastn_results3, sep='\t', names=['qseqid', 'sseqid', 'pident', 'length',
                                                                  'mismatch', 'gapopen', 'qstart', 'qend', 'sstart',
                                                                  'send', 'evalue', 'bitscore'])

    for seq in seqs_species:
        lenHit = 0
        order_i = ""
        for i in range(blastresult.shape[0]):
            if blastresult.loc[i, 'qseqid'] == seq:
                if lenHit < blastresult.loc[i, 'length']:
                    lenHit = blastresult.loc[i, 'length']
                    order_i = blastresult.loc[i, 'sseqid'].split("#")[1]
        print(lenHit)
        print(order_i)
        orders.append(order_i)
        keep_seqs.append(seq)

    orders = [x.replace("-", "/") for x in orders]
    print(orders)
    order_number = []
    orders_superfamilies = [x for x in dicc_orders.values()]
    for ord in range(len(orders)):
        if "/" in orders[ord]:
            order_given = orders[ord].split("/")[0]
            superfamily = orders[ord].split("/")[1]
        else:
            order_given = orders[ord]
            superfamily = "NA"

        if superfamily in orders_superfamilies:
            # superfamily found !!!
            order_number.append(orders_superfamilies.index(superfamily) + 2)
            print(orders_superfamilies.index(superfamily) + 2)
        elif order_given in orders_superfamilies:
            # Well, superfamily didn't find, but I found the order
            order_number.append(orders_superfamilies.index(order_given) + 2)
            print(orders_superfamilies.index(order_given) + 2)
        else:
            print("WARNING: Neither order and superfamily found: " + orders[ord])
            keep_seqs.remove(keep_seqs[ord])


    create_fasta_files("/home/simon/invase_species/100Drosophilas/curation_done_v2/unclassified_putative_tes_allSpecies.fa_nobuscoGenes", keep_seqs, "/home/simon/invase_species/100Drosophilas/curation_done_v2/"+proj_name+"/curation_raw/round7/round3/kept_seqs_unclass.fa", "/home/simon/invase_species/100Drosophilas/curation_done_v2/"+proj_name+"/curation_raw/round7/round3/", order_number)
```
