## Supplemental Code for "MCHelper automatically curates transposable element libraries across eukaryotic species": busco_like_metrics.py

```
from Bio import SeqIO
import pandas as pd
import os
import subprocess
import sys

ref_lib = sys.argv[1]
test_lib = sys.argv[2]
perc_ident = 80
len_tolerance = 1.10
cores = 32

busco_like_stats = {}

for te in SeqIO.parse(ref_lib, "fasta"):
    busco_like_stats[te.id] = [len(str(te.seq)), 0, 0]   # TE length, complete, fragment

if not os.path.exists(ref_lib + ".nhr"):
    output = subprocess.run(
        ['makeblastdb', '-in', ref_lib, '-dbtype', 'nucl'], stdout=subprocess.DEVNULL, stderr=subprocess.DEVNULL, text=True)

if not os.path.exists(test_lib + ".blast"):
    output = subprocess.run(
        ['blastn', '-query', test_lib, '-db', ref_lib, '-out', test_lib + ".blast", '-num_threads', str(cores),
         "-outfmt", "6", "-qcov_hsp_perc", "80", "-perc_identity", "80"],
        stdout=subprocess.PIPE, text=True)


blast_results = pd.read_table(test_lib + ".blast", sep='\t',
                                names=['qseqid', 'sseqid', 'pident', 'length', 'mismatch', 'gapopen', 'qstart',
                                       'qend', 'sstart', 'send', 'evalue', 'bitscore'])

for i in range(blast_results.shape[0]):
    te_id = blast_results.at[i, 'sseqid']
    te_values = busco_like_stats[te_id]
    hit_len = int(blast_results.at[i, 'length'])
    if hit_len <= te_values[0] * len_tolerance:
        if (hit_len * 100) / te_values[0] >= perc_ident:
            te_values[1] += 1  # a complete model
        else:
            te_values[2] += 1  # a fragment model
    #else:  # It's a model bigger than the actual
        #print("The model: "+blast_results.at[i, 'qseqid']+" representing to "+blast_results.at[i, 'sseqid']+" (len "+str(te_values[0])+") is "+str(blast_results.at[i, 'length'])+ " length")


single_copy = 0
duplic_copy = 0
fragme_copy = 0
missing = 0
for key in busco_like_stats.keys():
    if busco_like_stats[key][1] == 1:  # single and complete models
        single_copy += 1
    elif busco_like_stats[key][1] > 1:  # duplicated and complete models
        duplic_copy += 1
    elif busco_like_stats[key][2] > 0:  # fragmented models
        fragme_copy += 1
    elif busco_like_stats[key][1] + busco_like_stats[key][2] == 0:
        missing += 1

#print(str(single_copy)+"\t"+str(duplic_copy)+"\t"+str(fragme_copy)+"\t"+str(missing))
print(str(duplic_copy))
```
