## Supplemental Code for "MCHelper automatically curates transposable element libraries across eukaryotic species": count_FP.py

```
import pandas as pd
from Bio import SeqIO
import subprocess
import os
import sys

# Search for single repeats in TEs
new_ref_tes = sys.argv[1]
busco_library = sys.argv[2]
RNAs_library = sys.argv[3]+"/db/rRNA_Eukaryota.hmm"
tools_path = sys.argv[3]+"/tools/"
perc_ssr = 60
cores = 46

try:
    output = subprocess.run(
        [tools_path + '/trf409.linux64', new_ref_tes, '2', '3', '5', '80', '10', '20', '15', '-h', '-d'],
        stdout=subprocess.PIPE, stderr=subprocess.PIPE, text=True)
except Exception as exp:
    print("FATAL ERROR: I couldn't execute properly the TRF program. Please check the error: " + exp.args)

trf_file = os.path.basename(new_ref_tes) + ".2.3.5.80.10.20.15.dat"

kept_seqs = []
deleted_seqs = []
dicc_sr_pos = {}
with open(trf_file, 'r') as inFile:
    nbInSeq = 0
    for line in inFile:
        row = line.split(" ")
        if len(row) > 1 and "Sequence:" in row[0]:
            nbInSeq += 1
            seqName = row[1].replace('\n', '')
        if len(row) >= 14 and "Sequence:" not in row[0]:
            start = row[0]
            end = row[1]
            if seqName in dicc_sr_pos.keys():
                #dicc_sr[seqName] += int(end) - int(start) + 1
                dicc_sr_pos[seqName].append([int(start), int(end)])
            else:
                #dicc_sr[seqName] = int(end) - int(start) + 1
                dicc_sr_pos[seqName] = [[int(start), int(end)]]

# collapse all overlapping satellite matches
for sat in dicc_sr_pos.keys():
    pos_list = dicc_sr_pos[sat]
    new_list = [pos_list[0]]
    for i in range(1, len(pos_list)):
        # if there is an overlap
        overlap = False
        for j in range(len(new_list)):
            if (new_list[j][0] <= pos_list[i][0] <= new_list[j][1] or new_list[j][0] <= pos_list[i][1] <= new_list[j][1])\
            or (pos_list[i][0] <= new_list[j][0] <= pos_list[i][1] or pos_list[i][0] <= new_list[j][1] <= pos_list[i][1]):
                new_start = new_list[j][0] if new_list[j][0] < pos_list[i][0] else pos_list[i][0]
                new_end = new_list[j][1] if new_list[j][1] > pos_list[i][1] else pos_list[i][1]
                new_list[j] = [new_start, new_end]
                overlap = True
        if not overlap:
            new_list.append([pos_list[i][0], pos_list[i][1]])
    dicc_sr_pos[sat] = new_list

# count how many bases are satellites
dicc_sr = {}
for sat in dicc_sr_pos.keys():
    sum_bp = 0
    pos_list = dicc_sr_pos[sat]
    for i in range(len(pos_list)):
        sum_bp += pos_list[i][1] - pos_list[i][0] + 1
    dicc_sr[sat] = sum_bp

# Search for matches with References/BUSCO genes
if not os.path.exists(busco_library+".h3m"):
    output = subprocess.run(['hmmpress', busco_library],
                            stdout=subprocess.PIPE, stderr=subprocess.PIPE, text=True)

output = subprocess.run(
    ['hmmscan', '--tblout', "tes_vs_genes.hmm", '-E', '10', '--noali', '--cpu',
     str(cores), busco_library, new_ref_tes],
    stdout=subprocess.PIPE, stderr=subprocess.PIPE, text=True)

command = 'grep -v "^#" ' + 'tes_vs_genes.hmm | awk \'{print $1"\t"$2"\t"$3"\t"$4"\t"$5"\t"$6"\t"$7"\t"$8"\t"$9"\t"$10}\' > ' + "tes_vs_genes.hmm_formatted"
process = subprocess.Popen(command, shell=True)
process.communicate()
hmm_results = pd.read_table("tes_vs_genes.hmm_formatted", sep='\t',
                            names=['target', 'acc', 'query', 'acc_query', 'evalue', 'score', 'A', 'B', 'C',
                                   'D'])

tes_with_matches = []
for x in range(hmm_results.shape[0]):
    if hmm_results.loc[x, "query"] not in tes_with_matches:
        tes_with_matches.append(hmm_results.loc[x, "query"])

# Search for matches with tRNAs or rRNAs
if not os.path.exists(RNAs_library+".h3m"):
    output = subprocess.run(['hmmpress', RNAs_library],
                            stdout=subprocess.PIPE, stderr=subprocess.PIPE, text=True)

output = subprocess.run(
    ['hmmscan', '--tblout', new_ref_tes + "_tes_vs_rnas.hmm", '-E', '10', '--noali', '--cpu',
     str(cores), RNAs_library, new_ref_tes],
    stdout=subprocess.PIPE, stderr=subprocess.PIPE, text=True)

command = 'grep -v "^#" ' + new_ref_tes + '_tes_vs_rnas.hmm | awk \'{print $1"\t"$2"\t"$3"\t"$4"\t"$5"\t"$6"\t"$7"\t"$8"\t"$9"\t"$10}\' > ' + new_ref_tes + '_tes_vs_rnas.hmm_formatted'
process = subprocess.Popen(command, shell=True)
process.communicate()
hmm_results = pd.read_table(new_ref_tes + '_tes_vs_rnas.hmm_formatted', sep='\t',
                            names=['target', 'acc', 'query', 'acc_query', 'evalue', 'score', 'A', 'B', 'C',
                                   'D'])

tes_with_rnas = []
for x in range(hmm_results.shape[0]):
    if hmm_results.loc[x, "query"] not in tes_with_rnas:
        tes_with_rnas.append(hmm_results.loc[x, "query"])

# Remove those TEs with SSR > 60% or those that match with genes
ssr_filter = 0
rna_filter = 0
busco_filter = 0
for te in SeqIO.parse(new_ref_tes, "fasta"):
    if te.id not in tes_with_matches:
        if te.id not in tes_with_rnas: # It hasn't a match with Reference/BUSCO genes neither with RNAs
            if te.id in dicc_sr.keys():
                te_len = len(str(te.seq))
                lenSR = dicc_sr[te.id]
                if (lenSR * 100) / te_len < perc_ssr:  # It hasn't less than 60% of SSR in its sequence
                    kept_seqs.append(te.id)
                else:
                    ssr_filter += 1
                    deleted_seqs.append(te.id)
            else:  # It hasn't any SSR
                #print("Holi, "+te.id+" is nt in the dicc :/")
                kept_seqs.append(te.id)
        else:
            rna_filter += 1
            deleted_seqs.append(te.id)
    else:
        busco_filter += 1
        deleted_seqs.append(te.id)


"""kept_seqs_objets = [te for te in SeqIO.parse(new_ref_tes, "fasta") if te.id in kept_seqs]
SeqIO.write(kept_seqs_objets, new_ref_tes+"_good_candidates", "fasta")

output = subprocess.run(
        ['makeblastdb', '-in', curated_tes, '-dbtype', 'nucl'], stdout=subprocess.PIPE, text=True)

output = subprocess.run(
    ['blastn', '-query', new_ref_tes+"_good_candidates", '-db', curated_tes, '-out',
     new_ref_tes + "_unclasstes_vs_classtes.blast", '-num_threads', str(cores), "-outfmt", "6",
     "-qcov_hsp_perc", "70", "-perc_identity", "70"], stdout=subprocess.PIPE, text=True)

blastresult = pd.read_table(new_ref_tes + "_unclasstes_vs_classtes.blast", sep='\t',
                                names=['qseqid', 'sseqid', 'pident', 'length', 'mismatch', 'gapopen', 'qstart',
                                       'qend', 'sstart', 'send', 'evalue', 'bitscore'])
tes_with_hits = []
for x in range(blastresult.shape[0]):
    if blastresult.loc[x, "qseqid"] not in tes_with_hits:
        tes_with_hits.append(blastresult.loc[x, "qseqid"])"""

print(str(ssr_filter)+";"+str(busco_filter)+";"+str(rna_filter)+";"+str(ssr_filter+busco_filter+rna_filter))
```
