## Supplemental Code for "MCHelper automatically curates transposable element libraries across eukaryotic species": extract_quimerics_and_non_overlaping.py

```
# to have the chromatin "merge-count" file, run:
# awk '{print $1"\t"$2"\t"$3"\t"$4"\t"$5"\t"$6"\t"$7"\t"$8"\t"$10}' RM_BDGP/dmel-all-chromosome-r6.47.fasta.out.gff.euchr | bedtools sort | bedtools merge -c 9,9 -o count,collapse > lib_level_comparisons/BDGP_euchr_merge_count.gff

import sys

gff_file = sys.argv[1]

fileopen = open(gff_file).readlines()
outputfile = open(gff_file+"_quimeric", "w")
quim = 0
for line in fileopen:
    columns = line.split("\t")
    if int(columns[3]) > 1:
        families = columns[4].replace('\n', '').replace('"', '').split(",")
        diffFam = all(element == families[0] for element in families)
        if not diffFam:
            quim += 1
            outputfile.write(line)
print(quim)

outputfile = open(gff_file+"_non_overlapping", "w")
for line in fileopen:
    columns = line.split("\t")
    if int(columns[3]) > 1:
        families = columns[4].replace('\n', '').replace('"', '').split(",")
        diffFam = all(element == families[0] for element in families)
        if diffFam:
            outputfile.write(line)
    else:
    	outputfile.write(line)
```
