## Supplemental Code for "MCHelper automatically curates transposable element libraries across eukaryotic species": generate_length_dataframe.py

```
import pandas as pd

dataframes = []
real_names = {"Dmel": "D. melanogaster", "Osativa": "O. sativa", "Ccornix": "C. cornix", "Drerio": "D. rerio", "Zmays": "Z. mays", "Hsapiens": "H. sapiens"}
for spe in ["Dmel", "Osativa", "Ccornix", "Drerio", "Zmays", "Hsapiens"]:
	for program in ["RM2", "EDTA", "REPET"]:
		# for adding the MCHelper libraries
		path = "../"+spe+"/"+program+"/curation_ite16/curated_sequences_NR.fa_length.csv"
		try:
			df = pd.read_csv(path, sep=';') 
			df.fillna(0, inplace=True)
			df = df.drop(columns=['Unnamed: 0'])
			new_df = pd.DataFrame({'Species': [real_names[spe]]*df.shape[1], 'Library': [program+"+MCHelper"]*df.shape[1], 'LTR': df.iloc[0, :], 'DIRS': df.iloc[1, :], 'PLEs': df.iloc[2, :], 'LINE': df.iloc[3, :], 'SINE': df.iloc[4, :], 'TIR': df.iloc[5, :], 'Crypton': df.iloc[6, :], 'Helitron': df.iloc[7, :], 'Maverick': df.iloc[8, :], 'MITE': df.iloc[9, :] })
			dataframes.append(new_df)
		except:
			print("File '"+path+"' not found :P")	

	# for adding the Reference libraries
	path = "../"+spe+"/Reference/Reference.fa_length.csv"
	df = pd.read_csv(path, sep=';') 
	df.fillna(0, inplace=True)
	df = df.drop(columns=['Unnamed: 0'])
	new_df = pd.DataFrame({'Species': [real_names[spe]]*df.shape[1], 'Library': ["Reference"]*df.shape[1], 'LTR': df.iloc[0, :], 'DIRS': df.iloc[1, :], 'PLEs': df.iloc[2, :], 'LINE': df.iloc[3, :], 'SINE': df.iloc[4, :], 'TIR': df.iloc[5, :], 'Crypton': df.iloc[6, :], 'Helitron': df.iloc[7, :], 'Maverick': df.iloc[8, :], 'MITE': df.iloc[9, :] })
	dataframes.append(new_df)

	# for adding the RM2 libraries
	path = "../"+spe+"/RM2/"+spe+"-families.fa_length.csv"
	df = pd.read_csv(path, sep=';') 
	df.fillna(0, inplace=True)
	df = df.drop(columns=['Unnamed: 0'])
	new_df = pd.DataFrame({'Species': [real_names[spe]]*df.shape[1], 'Library': ["RM2"]*df.shape[1], 'LTR': df.iloc[0, :], 'DIRS': df.iloc[1, :], 'PLEs': df.iloc[2, :], 'LINE': df.iloc[3, :], 'SINE': df.iloc[4, :], 'TIR': df.iloc[5, :], 'Crypton': df.iloc[6, :], 'Helitron': df.iloc[7, :], 'Maverick': df.iloc[8, :], 'MITE': df.iloc[9, :] })
	dataframes.append(new_df)

	# for adding the REPET libraries
	try:
		path = "../"+spe+"/REPET/"+spe+"_refTEs.fa_short_length.csv"
		df = pd.read_csv(path, sep=';') 
		df.fillna(0, inplace=True)
		df = df.drop(columns=['Unnamed: 0'])
		new_df = pd.DataFrame({'Species': [real_names[spe]]*df.shape[1], 'Library': ["REPET"]*df.shape[1], 'LTR': df.iloc[0, :], 'DIRS': df.iloc[1, :], 'PLEs': df.iloc[2, :], 'LINE': df.iloc[3, :], 'SINE': df.iloc[4, :], 'TIR': df.iloc[5, :], 'Crypton': df.iloc[6, :], 'Helitron': df.iloc[7, :], 'Maverick': df.iloc[8, :], 'MITE': df.iloc[9, :] })
		dataframes.append(new_df)
	except:
		print("File '"+path+"' not found :P")

	# for adding the EDTA libraries
	try:
		path = "../"+spe+"/EDTA/EDTA.TElib.fa_length.csv"
		df = pd.read_csv(path, sep=';') 
		df.fillna(0, inplace=True)
		df = df.drop(columns=['Unnamed: 0'])
		new_df = pd.DataFrame({'Species': [real_names[spe]]*df.shape[1], 'Library': ["EDTA"]*df.shape[1], 'LTR': df.iloc[0, :], 'DIRS': df.iloc[1, :], 'PLEs': df.iloc[2, :], 'LINE': df.iloc[3, :], 'SINE': df.iloc[4, :], 'TIR': df.iloc[5, :], 'Crypton': df.iloc[6, :], 'Helitron': df.iloc[7, :], 'Maverick': df.iloc[8, :], 'MITE': df.iloc[9, :] })
		dataframes.append(new_df)
	except:
		print("File '"+path+"' not found :P")


# Concatenar todos los dataframes en uno solo
dataframe_final = pd.concat(dataframes, ignore_index=True)

dataframe_final.to_csv("FigureS2.csv", sep=';', index=False)
```
