## Supplemental Code for "MCHelper automatically curates transposable element libraries across eukaryotic species": genomic_proportions.py

```
import pandas as pd
from Bio import SeqIO
import subprocess
import os
import numpy as np
import matplotlib.pyplot as plt
import sys

genomic_size = int(sys.argv[1])
annot = sys.argv[2]

annot_RM_out = pd.read_table(annot, sep='\t', names=['chr', 'source', 'strategy', 'init', 'end',
                                                  'divergence', 'strand', 'dot', 'classification'])

dicc_orders = {'LTR': 0, 'LINEs': 0, 'DIRS': 0, 'PLEs': 0, 'TIR': 0, 'Maverick': 0, 'Helitron': 0, 'Cryton': 0, 'Unclassified': 0}

for i in range(annot_RM_out.shape[0]):
    length_copy = int(annot_RM_out.at[i, 'end']) - int(annot_RM_out.at[i, 'init'])
    if 'LTR' in annot_RM_out.at[i, 'classification'].split("#")[1].upper():
        dicc_orders['LTR'] += length_copy
    elif 'LINE' in annot_RM_out.at[i, 'classification'].split("#")[1].upper():
        dicc_orders['LINEs'] += length_copy
    elif 'DIRS' in annot_RM_out.at[i, 'classification'].split("#")[1].upper():
        dicc_orders['DIRS'] += length_copy
    elif 'PLES' in annot_RM_out.at[i, 'classification'].split("#")[1].upper():
        dicc_orders['PLEs'] += length_copy
    elif 'HELITRON' in annot_RM_out.at[i, 'classification'].split("#")[1].upper() or 'DNA/RC' in annot_RM_out.at[i, 'classification'].split("#")[1].upper():
        dicc_orders['Helitron'] += length_copy
    elif 'MAVERICK' in annot_RM_out.at[i, 'classification'].split("#")[1].upper():
        dicc_orders['Maverick'] += length_copy
    elif 'CRYPTON' in annot_RM_out.at[i, 'classification'].split("#")[1].upper():
        dicc_orders['Cryton'] += length_copy
    elif 'TIR' in annot_RM_out.at[i, 'classification'].split("#")[1].upper() or 'DNA' in annot_RM_out.at[i, 'classification'].split("#")[1].upper():
        dicc_orders['TIR'] += length_copy
    elif 'UNCLASSIFIED' in annot_RM_out.at[i, 'classification'].split("#")[1].upper() or 'UNKNOWN' in annot_RM_out.at[i, 'classification'].split("#")[1].upper():
        dicc_orders['Unclassified'] += length_copy

ltr = (dicc_orders['LTR'] * 100) / genomic_size
lines = (dicc_orders['LINEs'] * 100) / genomic_size
dirs = (dicc_orders['DIRS'] * 100) / genomic_size
ples = (dicc_orders['PLEs'] * 100) / genomic_size
tirs = (dicc_orders['TIR'] * 100) / genomic_size
Maverick = (dicc_orders['Maverick'] * 100) / genomic_size
Helitron = (dicc_orders['Helitron'] * 100) / genomic_size
Cryton = (dicc_orders['Cryton'] * 100) / genomic_size
Unclassified = (dicc_orders['Unclassified'] * 100) / genomic_size

print(str(ltr) + "\t" + str(lines) + "\t" + str(dirs) + "\t" + str(ples)
      + "\t" + str(tirs) + "\t" + str(Maverick) + "\t" + str(Helitron)
      + "\t" + str(Cryton) + "\t" + str(Unclassified))
```
