## Supplemental Code for "MCHelper automatically curates transposable element libraries across eukaryotic species": get_family_summary_paper.sh

```
#! /bin/bash
# sh get_family_summary_paper.sh <library.fasta.out>

# Jullien Flynn
#

# This script takes a RepeatMasker.out file that was produced by masking a benchmarked library (-lib parameter) against a test library you want to evaluate.
# For best results, use the -nolow parameter with RepeatMasker, e.g.:
# RepeatMasker -lib dmel_curated.fasta -nolow -pa 4 dmel_RepeatModeler2_lib.fasta

# It will calculate the number of sequences that are perfect, good, and total present. See Flynn, Hubley et al. RepeatModeler2 paper for details.


#date
d1=$(date +%s)
echo $1

# process the file, need to divide between forward sequence matches and reverse complement matches.

cat $1 | sed -e 1,3d | sed -e 's/^[ \t]*//' | tr -s " " | sed 's| |\t|g' | sed 's/(//g' | sed 's/)//g' > file_work_with.txt

cat file_work_with.txt | awk '$9=="C" {print $0}' | awk -v OFS="\t" '{print $10, $11, $12+$13, $13-$14, $5, $7+$8, $7-$6, $2}' > file2.c.txt

cat file2.c.txt | awk -v OFS="\t" '{print $1, $2, $3, $4/$3, $5, $6, $7/$6, $8}' > file3.c.txt

cat file_work_with.txt | awk '$9=="+" {print $0}' | awk -v OFS="\t" '{print $10, $11, $14+$13, $13-$12, $5, $7+$8, $7-$6, $2}' > file2.f.txt

cat file2.f.txt | awk -v OFS="\t" '{print $1, $2, $3, $4/$3, $5, $6, $7/$6, $8}' > file3.f.txt

# Filter the sequences

cat file3.f.txt file3.c.txt | awk '$7>0.1 {print $0}' > file_final.0.1.txt

#perfect 1:1 match: these are "perfect families"
echo "perfect 1:1 match:"
cat file_final.0.1.txt | awk '{if($4>0.95 && $7>0.95 && $8<5.0) {print $0}}' | cut -f 1 | sort -u > perfect.families
wc -l perfect.families


# next build a script to see its total percent coverage across all contigs
# do for c and f separately
# loop through each repbase family found:
# print out family, print out start and end of family for that alignment, print family sequence length, print out contig name, print out divergence
# for each family, use bedtools to get the coverage of the family.

cat file_work_with.txt | cut -f 10 | sort -u > repbase.families

# this is the number of present families
while read f
do
	cat file_work_with.txt | awk '$9=="C" {print $0}' | awk -v f="$f" '$10==f {print $0}' | awk -v OFS="\t" '{print $10, $12+$13, $14, $13, $5, $2}' > $f.c.summary
	cat file_work_with.txt | awk '$9=="+" {print $0}' | awk -v f="$f" '$10==f {print $0}' | awk -v OFS="\t" '{print $10, $14+$13, $12, $13, $5, $2}' > $f.f.summary
	# cambiar aqquiiiiiiiiiiiii. Original:  $6<20.0
	cat $f.c.summary $f.f.summary | awk '$6<20.0 {print $0}' > $f.summary # check only for things that are within 10% divergence
	#make into bed file
	cat $f.summary | awk -v OFS="\t" '{print $1,$3,$4}' | sort -k2,2n > $f.bed
	bases=`bedtools merge -i $f.bed -d 10 | awk -v OFS="\t" '{print $1, $2, $3, $3-$2}' | awk '{ sum += $4 } END { print sum }'`
	length=`cat $f.summary | cut -f 2 | head -n 1`
	printf "%s\t%i\t%i\n" $f $length $bases >> repbase.families.summary80
done < repbase.families

less repbase.families.summary80 | awk '$2>0 {print $0}' | awk -v OFS="\t" '{print $1, $2, $3/$2}' > repbase.families.covsumm80	

# cambiar aqquiiiiiiiiiiiii. Original: $3>0.8 Coberturaaaaa
cat repbase.families.covsumm80 | awk '$3>0.8 {print $0}' | cut -f 1 | sort -u > present.all.families

rm *.bed

mkdir summary_files
mv *.summary summary_files

# this will get the number of good families -  do the same thing with different thresholds
while read f
do
	cat file_work_with.txt | awk '$9=="C" {print $0}' | awk -v f="$f" '$10==f {print $0}' | awk -v OFS="\t" '{print $10, $12+$13, $14, $13, $5, $2}' > $f.c.summary
	cat file_work_with.txt | awk '$9=="+" {print $0}' | awk -v f="$f" '$10==f {print $0}' | awk -v OFS="\t" '{print $10, $14+$13, $12, $13, $5, $2}' > $f.f.summary
	cat $f.c.summary $f.f.summary | awk '$6<5.0 {print $0}' > $f.summary # check only for things that are within 10% divergence
	# get the max and min and compare to the length
	cat $f.summary | awk -v OFS="\t" '{print $1,$3,$4}' | sort -k2,2n > $f.bed
	bases=`bedtools merge -i $f.bed -d 10 | awk -v OFS="\t" '{print $1, $2, $3, $3-$2}' | awk '{ sum += $4 } END { print sum }'`
	length=`cat $f.summary | cut -f 2 | head -n 1`
	printf "%s\t%i\t%i\n" $f $length $bases >> repbase.families.summary95
done < repbase.families

cat repbase.families.summary95 | awk '$2>0 {print $0}'  | awk -v OFS="\t" '{print $1, $2, $3/$2}' > repbase.families.covsumm95
cat repbase.families.covsumm95 | awk '$3>0.95 {print $0}' | cut -f 1 | sort -u > good.perfect.families

comm -23 good.perfect.families perfect.families > good.families
echo "number of good families:"
wc -l good.families

echo " total present families:"
wc -l present.all.families

rm *.bed


mv *.summary summary_files


#date
d2=$(date +%s)
sec=$(( ( $d2 - $d1 ) ))
hour=$(echo - | awk '{ print '$sec'/3600}')
echo Runtime: $hour hours \($sec\s\)
```
