## Supplemental Code for "MCHelper automatically curates transposable element libraries across eukaryotic species": lib_metrics_paper_v2.sh

```
#!/bin/bash
export LC_ALL=C
Script_path=`echo "$(cd "$(dirname "$0")" && pwd -P)"`
threads=40
mchelper_path=/lustre/home/ibe/sorozco/MCHelper

# D melanogaster
work_dir=/lustre/home/ibe/sorozco/Dmel_repet/
ref_lib=/lustre/home/ibe/sorozco/Dmel/reference/D_mel_transposon_sequence_set_v10.2.fa
raw_lib_dir=${work_dir}/
raw_lib_name=Dmel_refTEs_class.fa
raw_lib=`echo "${raw_lib_dir}/${raw_lib_name}"`
cur_lib_dir=${work_dir}/curation_ite16
cur_lib_name=curated_sequences_NR.fa
cur_lib=`echo "${cur_lib_dir}/${cur_lib_name}"`
busco=/lustre/home/ibe/sorozco/MCHelper/db/diptera_odb10.hmm


rm -f lib_metrics.csv
rm -rf summary_files good.families good.perfect.families repbase.* file_work_with.txt file2.c.txt file3.c.txt file2.f.txt
########################################## reference #####################################################
result_ref=`grep -c ">" $ref_lib`
lengths_ref=`python3 ${Script_path}/te_order_len.py $ref_lib $work_dir "reference_len.png"`
false_positive=`python3 ${Script_path}/count_FP.py $ref_lib $busco $mchelper_path`
echo "$result_ref;0;0;0;0;$lengths_ref;$false_positive;0;0" >> lib_metrics.csv

########################################## raw  ##########################################################
rm -f ${raw_lib}.blast
result_raw=`grep -c ">" ${raw_lib}`
RepeatMasker -lib ${ref_lib} -nolow -pa ${threads} -dir ${raw_lib_dir}/RM_ref ${raw_lib}
metrics_output=`sh ${Script_path}/get_family_summary_paper.sh ${raw_lib_dir}/RM_ref/${raw_lib_name}.out`
echo $metrics_output
perfect=`cat perfect.families | wc -l`
good=`cat good.families | wc -l`
present_tmp=`cat present.all.families | wc -l`
good_perfect=`cat good.perfect.families | wc -l`
present=$(( $present_tmp - $good_perfect ))
missing=$(( $result_ref - $perfect - $good - $present ))

lengths_ref=`python3 ${Script_path}/te_order_len.py $raw_lib $work_dir "raw_len.png"`

false_positive=`python3 ${Script_path}/count_FP.py $raw_lib $busco $mchelper_path`

avg_models=`cut -f2 ${raw_lib}.blast | sort | uniq -c | awk '{sum=sum+$1; con=con+1}END{print sum/con}'`

echo "$result_raw;$perfect;$good;$present;$missing;$lengths_ref;$false_positive;$avg_models" >> lib_metrics.csv

rm -rf summary_files good.families good.perfect.families repbase.* file_work_with.txt file2.c.txt file3.c.txt file2.f.txt
########################################## Curated  ######################################################
rm -f ${cur_lib}.blast
result_cur=`grep -c ">" ${cur_lib}`

RepeatMasker -lib ${ref_lib} -nolow -pa ${threads} -dir ${cur_lib_dir}/RM_ref ${cur_lib}
rm -rf summary_files
metrics_output=`sh ${Script_path}/get_family_summary_paper.sh ${cur_lib_dir}/RM_ref/${cur_lib_name}.out`
perfect=`cat perfect.families | wc -l`
good=`cat good.families | wc -l`
present_tmp=`cat present.all.families | wc -l`
good_perfect=`cat good.perfect.families | wc -l`
present=$(( $present_tmp - $good_perfect ))
missing=$(( $result_ref - $perfect - $good - $present ))

lengths_ref=`python3 ${Script_path}/te_order_len.py ${cur_lib} $work_dir "curated_len.png"`

false_positive=`python3 ${Script_path}/count_FP.py ${cur_lib} $busco $mchelper_path`

avg_models=`cut -f2 ${cur_lib}.blast | sort | uniq -c | awk '{sum=sum+$1; con=con+1}END{print sum/con}'`

echo "$result_cur;$perfect;$good;$present;$missing;$lengths_ref;$false_positive;;$avg_models" >> lib_metrics.csv
rm -rf summary_files good.families good.perfect.families repbase.* file_work_with.txt file2.c.txt file3.c.txt file2.f.txt
```
