## Supplemental Code for "MCHelper automatically curates transposable element libraries across eukaryotic species": mcc.py

```
import sys
import math

TP = int(sys.argv[1])
FN = int(sys.argv[2])
TN = int(sys.argv[3])
FP = int(sys.argv[4])

mcc = (TP * TN - FP * FN) / math.sqrt((TP + FP) * (TP + FN) * (TN + FP) * (TN + FN))
print(mcc)
```
