## Supplemental Code for "MCHelper automatically curates transposable element libraries across eukaryotic species": NTE50_LTE50_FLC_v2.py

```
import pandas as pd
from Bio import SeqIO
import subprocess
import os
import numpy as np
#import matplotlib.pyplot as plt
import sys

# NTE50
genome_size = int(sys.argv[1])
perc_TEs = float(sys.argv[2])
per_base_tes = genome_size * (perc_TEs/100)
cores = 48
ref_annot = sys.argv[3]
raw_annot = sys.argv[4]
ref_lib = sys.argv[5]
raw_lib = sys.argv[6]


########################### reference annots ########################################
annot_gff = pd.read_table(ref_annot, sep='\t',
                         names=['score', 'divergence', 'deletion', 'insertion', 'query', 'start', 'end', 'length',
                                'sense', 'family', 'classification', 'start_TE', 'end_TE', 'TE_left', 'ID',
                                'num_assembled', '%_ref'])
annot_lengths = []
ref_len = {}

for te in SeqIO.parse(ref_lib, "fasta"):
    ref_len[str(te.id).split("#")[0]] = len(str(te.seq))

FLF = 0
FLC = 0
for x in range(annot_gff.shape[0]):
    annot_lengths.append(int(annot_gff.loc[x, 'length']))
    real_name = annot_gff.loc[x, 'family'].replace('-int', '').replace('-LTR', '')
    if real_name in ref_len.keys():
        if annot_gff.loc[x, '%_ref'] != "No_ref_available":
            if float(annot_gff.loc[x, '%_ref']) > 0.94:
                if int(annot_gff.loc[x, 'num_assembled']) == 1:
                    FLF += 1
                else:
                    FLC += 1
        elif (int(annot_gff.loc[x, 'length'])*100)/ref_len[real_name] > 94:  # Full length element
            if int(annot_gff.loc[x, 'num_assembled']) == 1:
                FLF += 1
            else:
                FLC += 1

annot_lengths_sorted = sorted(annot_lengths, reverse=True)
perc_cov_ref = []
acum_sum_bp_ref = 0
acum_sum_bp = 0
medium_pos = -1
for i in range(len(annot_lengths_sorted)):
    acum_sum_bp += annot_lengths_sorted[i]
    perc_cov_ref.append((acum_sum_bp*100)/genome_size)
    if acum_sum_bp >= (per_base_tes / 2) and medium_pos == -1:
        medium_pos = i
        acum_sum_bp_ref = acum_sum_bp

print(str(medium_pos) + "\t" + str(annot_lengths_sorted[medium_pos]) + "\t" + str(FLF) + "\t" + str(FLC) + "\t" + str(sum(annot_lengths_sorted)/genome_size))
annot_lengths_ref_sorted = annot_lengths_sorted

########################### Raw annots ########################################
annot_gff = pd.read_table(raw_annot, sep='\t',
                         names=['score', 'divergence', 'deletion', 'insertion', 'query', 'start', 'end', 'length',
                                'sense', 'family', 'classification', 'start_TE', 'end_TE', 'TE_left', 'ID',
                                'num_assembled', '%_ref'])

annot_lengths = []
ref_len = {}

for te in SeqIO.parse(raw_lib, "fasta"):
    ref_len[str(te.id).split("#")[0]] = len(str(te.seq))

FLF = 0
FLC = 0
for x in range(annot_gff.shape[0]):
    annot_lengths.append(int(annot_gff.loc[x, 'length']))
    real_name = annot_gff.loc[x, 'family'].replace('-int', '')
    if real_name in ref_len.keys():
        if annot_gff.loc[x, '%_ref'] != "No_ref_available":
            if float(annot_gff.loc[x, '%_ref']) > 0.94:
                if int(annot_gff.loc[x, 'num_assembled']) == 1:
                    FLF += 1
                else:
                    FLC += 1
        elif (int(annot_gff.loc[x, 'length']) * 100) / ref_len[real_name] > 94:  # Full length element
            if int(annot_gff.loc[x, 'num_assembled']) == 1:
                FLF += 1
            else:
                FLC += 1

annot_lengths_sorted = sorted(annot_lengths, reverse=True)
perc_cov_raw = []
medium_pos = -1
acum_sum_bp_raw = 0
acum_sum_bp = 0
for i in range(len(annot_lengths_sorted)):
    acum_sum_bp += annot_lengths_sorted[i]
    perc_cov_raw.append((acum_sum_bp * 100) / genome_size)
    if acum_sum_bp >= (per_base_tes / 2) and medium_pos == -1:
        medium_pos = i
        acum_sum_bp_raw = acum_sum_bp

print(str(medium_pos)+ "\t" + str(annot_lengths_sorted[medium_pos]) + "\t" + str(FLF) + "\t" + str(FLC) + "\t" + str(sum(annot_lengths_sorted)/genome_size))
annot_lengths_raw_sorted = annot_lengths_sorted


"""
cumulative_ref = np.flip(annot_lengths_ref_sorted, 0)
np.cumsum(cumulative_ref, out=cumulative_ref)
cumulative_raw = np.flip(annot_lengths_raw_sorted, 0)
np.cumsum(cumulative_raw, out=cumulative_raw)
cumulative_cur = np.flip(annot_lengths_cur_sorted, 0)
np.cumsum(cumulative_cur, out=cumulative_cur)

############################# PLOT ##########################################
# plot the cumulative function
#plt.plot([x for x in range(len(perc_cov_fly))], perc_cov_fly, c='black')
plt.plot([x for x in range(len(perc_cov_ref))], perc_cov_ref, c='blue')
plt.plot([x for x in range(len(perc_cov_raw))], perc_cov_raw, c='green')
plt.plot([x for x in range(len(perc_cov_cur))], perc_cov_cur, c='red')
plt.legend(["Reference", "Raw", "Curated"])
plt.ylim([0, 60])
plt.show()"""
species = sys.argv[7]
library = sys.argv[8]

if library == "Reference":
    df_export = pd.DataFrame({'Species': [species]*len(perc_cov_ref), 'Library': [library]*len(perc_cov_ref), 'Value': perc_cov_ref})
else:
    df_export = pd.DataFrame({'Species': [species]*len(perc_cov_raw), 'Library': [library]*len(perc_cov_raw), 'Value': perc_cov_raw})
#df_export = pd.concat([df_export, pd.DataFrame({'Species': [species]*len(perc_cov_raw), 'Library': [library]*len(perc_cov_raw), 'Value': perc_cov_raw})])
#df_export = pd.concat([df_export, pd.DataFrame({'Species': [species]*len(perc_cov_cur), 'Program': [program]*len(perc_cov_cur), 'Library': ['MCHelper']*len(perc_cov_cur), 'Value': perc_cov_cur})])

file_name = species.replace(". ", "")+"_"+library+".csv"
df_export.to_csv(file_name, sep=",", index=False)
"""
df_export = pd.read_csv("Dmelanogaster_RM2.csv", sep=",", index_col=False, header=0)
df_export = pd.concat([df_export, pd.read_csv("Dmelanogaster_REPET.csv", sep=",", index_col=False, header=0)], ignore_index=True)
df_export = pd.concat([df_export, pd.read_csv("Osativa_RM2.csv", sep=",", index_col=False, header=0)], ignore_index=True)
df_export = pd.concat([df_export, pd.read_csv("Osativa_REPET.csv", sep=",", index_col=False, header=0)], ignore_index=True)
df_export = pd.concat([df_export, pd.read_csv("Drerio_RM2.csv", sep=",", index_col=False, header=0)], ignore_index=True)
df_export = pd.concat([df_export, pd.read_csv("Drerio_REPET.csv", sep=",", index_col=False, header=0)], ignore_index=True)
print(df_export)
df_export.to_csv("NTE50.csv", sep=",", index=False)"""
```
