## Supplemental Code for "MCHelper automatically curates transposable element libraries across eukaryotic species": te_order_len.py

```
# Import libraries
#import matplotlib.pyplot as plt
import numpy as np
from Bio import SeqIO
import pandas as pd
import sys
from statistics import mean


#fig = plt.figure(figsize=(20, 10))

# Creating dataset
fasta_file = sys.argv[1]
output_path = sys.argv[2]
output_file = sys.argv[3]


data_ltr = []
data_dirs = []
data_ple = []
data_line = []
data_sine = []
data_tir = []
data_crypton = []
data_helitron = []
data_maverick = []
data_mite = []
for te in SeqIO.parse(fasta_file, "fasta"):
    if "#" in str(te.id):
        order = str(te.id).split("#")[1].split(" ")[0]
        if "DIRS" in order.upper():
            data_dirs.append(len(str(te.seq)))
        elif "PLE" in order.upper() or "PENELOPE" in order.upper():
            data_ple.append(len(str(te.seq)))
        elif "LTR" in order.upper():
            data_ltr.append(len(str(te.seq)))
        elif "LINE" in order.upper():
            data_line.append(len(str(te.seq)))
        elif "SINE" in order.upper():
            data_sine.append(len(str(te.seq)))
        elif "HELITRON" in order.upper():
            data_helitron.append(len(str(te.seq)))
        elif "MAVERICK" in order.upper():
            data_helitron.append(len(str(te.seq)))
        elif "MITE" in order.upper():
            data_mite.append(len(str(te.seq)))
        elif "TIR" in order.upper() or "DNA" in order.upper():
            data_tir.append(len(str(te.seq)))


meanltr = 0
meandirs = 0
meanple = 0
meanline = 0
meansine = 0
meantir = 0
meancrypton = 0
meanhelitron = 0
meanmaverick = 0
meanmite = 0
if len(data_ltr) > 0:
    meanltr = mean(data_ltr)
if len(data_dirs) > 0:
    meandirs = mean(data_dirs)
if len(data_ple) > 0:
    meanple = mean(data_ple)
if len(data_line) > 0:
    meanline = mean(data_line)
if len(data_sine) > 0:
    meansine = mean(data_sine)
if len(data_tir) > 0:
    meantir = mean(data_tir)
if len(data_crypton) > 0:
    meancrypton = mean(data_crypton)
if len(data_helitron) > 0:
    meanhelitron = mean(data_helitron)
if len(data_maverick) > 0:
    meanmaverick = mean(data_maverick)
if len(data_mite) > 0:
    meanmite = mean(data_mite)

print(str(meanltr) + ";" + str(meandirs) + ";" + str(meanple) + ";" + str(meanline) + ";" + str(meansine) + ";" + str(meantir) + ";" + str(meancrypton) + ";" + str(meanhelitron) + ";" + str(meanmaverick) + ";" + str(meanmite))

dataframe = pd.DataFrame([data_ltr, data_dirs, data_ple, data_line, data_sine, data_tir, data_crypton, data_helitron, data_maverick, data_mite])
dataframe.to_csv(fasta_file+"_length.csv", sep=";")

# Creating plot
#data = [data_1, data_2, data_3, data_4]
#fig, ax = plt.subplots()
# Creating axes instance
#bp = ax.boxplot(data)
#ax.set_ylim(0, 25000)
# x-axis labels
#ax.set_xticklabels(['LTR', 'LINE', 'TIR', 'Helitron'])
#plt.show()
#plt.savefig(output_path+'/'+output_file)
```
