## Supplemental Code for "MCHelper automatically curates transposable element libraries across eukaryotic species": MCHelper.py

```
"""

Manual Curation Helper Version 1.7.0
Novelties:
  * Added MeShClust as an alternative clustering algorithm (-k parameter)
  * Changed cd-hit output to keep the entire sequence IDs
  * Removed the Pandas' Future Warnings
  * Check sequence IDs in the genome to not containing only numbers
  * Removed TE sequences with lengths < 100 pb
  * Added WebAgg GUI backend to work with WSL version 1 (Experimental)
  * TE+Aid plots are now kept in PDF due to CentOS incompatibilities

"""

import sys
import os
import io
import re
import argparse
import pandas as pd
import multiprocessing
from pdf2image import convert_from_path
from Bio import SeqIO
from Bio.Seq import Seq
from Bio.SeqRecord import SeqRecord
import subprocess
import cv2
from matplotlib import pyplot as plt
#import matplotlib
#matplotlib.use('WebAgg')
import psutil
import glob
import shutil
import time
import numpy as np
import math
from sklearn.cluster import DBSCAN
import warnings
warnings.simplefilter(action='ignore', category=FutureWarning)

dicc_orders = {2: "LTR", 3: "COPIA", 4: "GYPSY", 5: "BELPAO", 6: "ERV", 7: "TRIM", 8: "LARD", 9: "LINE", 10: "SINE",
               11: "R2", 12: "RTE", 13: "JOCKEY", 14: "L1", 15: "I", 16: "R1", 17: "CR1", 18: "LOA", 19: "L2",  20: "PLE",
               21: "DIRS", 22: "NGARO", 23: "VIPER", 24: "TIR", 25: "MITE", 26: "TC1MARINER",  27: "HAT", 28: "MUTATOR",
               29: "MERLIN", 30: "TRANSIB", 31: "P", 32: "PIGGYBAC", 33: "PIFHARBINGER",  34: "CACTA", 35: "MULE",
               36: "CMC", 37: "HELITRON", 38: "MAVERICK", 39: "CRYPTON", 40: "UNCLASSIFIED", 41: "CLASSI", 42: "CLASSII"}

orders_superfamilies = [x for x in dicc_orders.values()]


def write_sequences_file(sequences, filename):
    try:
        SeqIO.write(sequences, filename, "fasta")
    except FileNotFoundError:
        print("FATAL ERROR: I couldn't find the file, please check: '" + filename + "'. Path not found")
        sys.exit(0)
    except PermissionError:
        print("FATAL ERROR: I couldn't access the files, please check: '" + filename + "'. I don't have permissions.")
        sys.exit(0)
    except Exception as exp :
        print("FATAL ERROR: There is a unknown problem writing sequences in : '" + filename + "'.")
        print(exp)
        sys.exit(0)


def create_output_folders(folder_path):
    try:
        if not os.path.exists(folder_path):
            os.mkdir(folder_path)
    except FileNotFoundError:
        print("FATAL ERROR: I couldn't create the folder " + folder_path + ". Path not found")
        sys.exit(0)
    except PermissionError:
        print("FATAL ERROR: I couldn't create the folder " + folder_path + ". I don't have permissions.")
        sys.exit(0)


def delete_files(file_path):
    if os.path.exists(file_path):
        try:
            os.remove(file_path)
        except PermissionError:
            print("WARNING: The file " + file_path + " couldn't be removed because I don't have permissions.")


def filter_flf(ref_tes, flf_file, minFullLenFragments, outputdir):
    flf_lines = open(flf_file, 'r').readlines()[1:]
    seqsID_with_flf = [s.split("\t")[0] for s in flf_lines]
    seqs_with_flf = []
    numCopies = {}
    for te in SeqIO.parse(ref_tes, "fasta"):
        seq_name = str(te.id).split("#")[0]
        if seq_name in seqsID_with_flf:
            numfullCopies = int([s.split("\t")[6] for s in flf_lines if s.split("\t")[0] == seq_name][0])
            numfullFrags = int([s.split("\t")[4] for s in flf_lines if s.split("\t")[0] == seq_name][0])
            if numfullFrags >= minFullLenFragments:  # number of copies greater or equal than threshold?
                seqs_with_flf.append(te)
                numCopies[seq_name] = [numfullFrags, numfullCopies]

    write_sequences_file(seqs_with_flf, outputdir + "/cons_flf.fa")
    return outputdir + "/cons_flf.fa", numCopies


def processing_gff(seqname, gff_files, pre):
    posBlan = []
    postBlax = []
    posBlax = []
    posTR = []

    gff_blastn = open(gff_files + '/' + pre + '_TE_BLRn.gff3', 'r').readlines()
    for anno in gff_blastn:
        columns = anno.split('\t')
        # if columns[0] == seqname and columns[2] == 'TE_BLRn':
        if columns[0] == seqname and columns[2] == 'exon':
            posBlan.append((int(columns[3]), int(columns[4]) - int(columns[3])))

    gff_tblastx = open(gff_files + '/' + pre + '_TE_BLRtx.gff3', 'r').readlines()
    for anno in gff_tblastx:
        columns = anno.split('\t')
        # if columns[0] == seqname and columns[2] == 'TE_BLRtx':
        if columns[0] == seqname and columns[2] == 'exon':
            postBlax.append((int(columns[3]), int(columns[4]) - int(columns[3])))

    gff_blastx = open(gff_files + '/' + pre + '_TE_BLRx.gff3', 'r').readlines()
    for anno in gff_blastx:
        columns = anno.split('\t')
        # if columns[0] == seqname and columns[2] == 'TE_BLRx':
        if columns[0] == seqname and columns[2] == 'exon':
            posBlax.append((int(columns[3]), int(columns[4]) - int(columns[3])))

    gff_tr = open(gff_files + '/' + pre + '_TR.gff3', 'r').readlines()
    for anno in gff_tr:
        columns = anno.split('\t')
        if columns[0] == seqname and columns[2] == 'exon':
            posTR.append((int(columns[3]), int(columns[4]) - int(columns[3])))

    return posBlan, postBlax, posBlax, posTR


def count_flf_fasta(ref_tes, genome, cores, outputdir):

    if not os.path.exists(outputdir + "/fullLengthFrag.txt"):
        create_output_folders(outputdir)

        flf_df = pd.DataFrame(columns=['TE', 'length', 'covg', 'frags', 'fullLgthFrags', 'copies', 'fullLgthCopies'])

        if not os.path.exists(genome + ".nhr"):
            output = subprocess.run(
                ['makeblastdb', '-in', genome, '-dbtype', 'nucl'], stdout=subprocess.PIPE, text=True)

        output = subprocess.run(
            ['blastn', '-query', ref_tes, '-db', genome, '-out', outputdir + "/TEs_vs_genome.blast", '-num_threads',
             str(cores), "-outfmt", "6 qseqid length", "-evalue", "10e-8"], stdout=subprocess.PIPE, text=True)

        blastresult = pd.read_table(outputdir + "/TEs_vs_genome.blast", sep='\t', names=['qseqid', 'length'])

        for te in SeqIO.parse(ref_tes, "fasta"):
            seq_name = str(te.id).split("#")[0]
            te_hits = blastresult[blastresult.qseqid == str(te.id)].reset_index()
            count_frag = 0
            count_flf = 0
            te_len = len(str(te.seq))
            for i in range(te_hits.shape[0]):
                len_hit = te_hits.loc[i, 'length']
                if len_hit >= (te_len * 0.94):  # it's a full-length fragment
                    count_flf += 1
                else:
                    count_frag += 1
            flf_df = pd.concat([flf_df, pd.DataFrame({'TE': seq_name, 'length': [te_len], 'covg': [0], 'frags': [count_frag], 'fullLgthFrags': [count_flf],
                 'copies': [count_frag], 'fullLgthCopies': [count_flf]})], ignore_index=True)

        flf_df.to_csv(outputdir + "/fullLengthFrag.txt", header=True, sep='\t', index=False)
        delete_files(outputdir + "/TEs_vs_genome.blast")
    else:
        print("MESSAGE: Full-length file was already created, please verify or change the output directory")
    return outputdir + "/fullLengthFrag.txt"


def checkChrNames(genome_file):
    Ids = [x.id.isdigit() for x in SeqIO.parse(genome_file, "fasta")]
    if True in Ids:
        return False
    else:
        return True


def check_classification_Userlibrary(user_library, outputdir):
    ids_no_contained = []
    reasons = []
    fine_tes = []
    for te in SeqIO.parse(user_library, "fasta"):
        if "#" not in str(te.id):  # it doesn't have the correct format
            ids_no_contained.append(te.id)
            reasons.append(
                "The sequence ID doesn't have the character '#' needed to separate the ID to the classification")
        elif len(str(te.id).split("#")[1].split("/")) > 3:
            ids_no_contained.append(te.id)
            reasons.append(
                "The sequence ID has more than 3 levels of classification. I can work with the following formats:\n   1) Class/Order/Superfamily\n   2) Order/Superfamily\n   Class/Order\n   3) order or superfamily")
        else:
            classification = str(te.id).split("#")[1].upper().replace("-", "")
            if len(classification.split("/")) == 3:
                # The most complete case Class / order / superfamily
                order_given = classification.split("/")[1]
                superfamily = classification.split("/")[2]
            elif len(classification.split("/")) == 2:
                # The most common case: Class / Order or  order / superfamily
                order_given = classification.split("/")[0]
                superfamily = classification.split("/")[1]
            elif len(classification.split("/")) == 1:
                # the most incomplete but still fine to work with: only the order or superfamily is provided
                order_given = classification
                superfamily = "NA"

            if superfamily not in orders_superfamilies and order_given not in orders_superfamilies:
                # Maybe is it Repbase/Dfam taxonomy?
                if "UNKNOWN" in classification:
                    te.id = te.id.split("#")[0] + "#Unclassified"
                    te.description = ""
                    fine_tes.append(te)
                elif "DNA" in classification:
                    classification = classification.replace("DNA", "TIR")
                    te.id = te.id.split("#")[0] + "#" + classification
                    te.description = ""
                    fine_tes.append(te)
                else:
                    ids_no_contained.append(te.id)
                    reasons.append(
                        "The classification wasn't find in my classification system. Remember that I use the Wicker nomenclature.")
            else:
                fine_tes.append(te)

    # Checking that there is no duplicated sequences
    duplicates = False
    if len(fine_tes) != len(list(set([x.id for x in fine_tes]))):
        freq = {}
        for TE in fine_tes:
            freq[TE.id] = freq.get(TE.id, 0) + 1
            if freq[TE.id] > 1:
                ids_no_contained.append(TE.id)
                reasons.append("Duplicated sequence ID")
                duplicates = True
    write_sequences_file(fine_tes, outputdir + "/candidate_tes.fa")
    if len(ids_no_contained) == 0:  # Congrats!! everything looks great!
        return 0
    else:
        output_file = open(outputdir + "/sequences_with_problems.txt", "w")
        output_file.write("Sequence ID\tProblem\n")
        for i in range(len(ids_no_contained)):
            output_file.write(ids_no_contained[i] + "\t" + reasons[i] + "\n")
        if duplicates:
            return -2
        return -1


def check_repet_input_folder(repet_input_dir, proj_name):
    ref_tes = repet_input_dir + "/" + proj_name + "_refTEs.fa"
    # flf_file = repet_input_dir + "/TEannot/" + proj_name + "_chr_allTEs_nr_noSSR_join_path.annotStatsPerTE_FullLengthFrag.txt"
    features_table = repet_input_dir + "/" + proj_name + "_denovoLibTEs_PC.classif"
    plots_dir = repet_input_dir + "/plotCoverage"
    gff_files = repet_input_dir + "/gff_reversed"

    if not os.path.exists(repet_input_dir):
        valid = False
        reason = "FATAL ERROR: "+repet_input_dir+" does not exist. Please check the path and re-run the software"
    elif not os.path.exists(ref_tes):
        valid = False
        reason = "FATAL ERROR: "+ref_tes+" does not exist. Please check the path and re-run the software"
    elif os.path.getsize(ref_tes) == 0:
        valid = False
        reason = "FATAL ERROR: "+ref_tes+" is empty. Please check the file and re-run the software"
    #elif not os.path.exists(flf_file):
    #    valid = False
    #    reason = "FATAL ERROR: "+flf_file+" does not exist. Please check the path and re-run the software"
    #elif os.path.getsize(flf_file) == 0:
    #    valid = False
    #    reason = "FATAL ERROR: "+flf_file+" is empty. Please check the file and re-run the software"
    elif not os.path.exists(features_table):
        valid = False
        reason = "FATAL ERROR: "+features_table+" does not exist. Please check the path and re-run the software"
    elif os.path.getsize(features_table) == 0:
        valid = False
        reason = "FATAL ERROR: "+features_table+" is empty. Please check the file and re-run the software"
    elif not os.path.exists(plots_dir):
        valid = False
        reason = "FATAL ERROR: "+plots_dir+" does not exist. Please check the path and re-run the software"
    elif not os.path.exists(gff_files):
        valid = False
        reason = "FATAL ERROR: "+gff_files+" does not exist. Please check the path and re-run the software"
    else:
        # In this case, everything is ok to run the curation process !!
        valid = True
        reason = ""

    return valid, reason


def find_TRs2(te, outputdir, minLTR, minTIR, minpolyA, cores):
    lenLTR = 0
    lenTIR = 0
    lenPolyA = 0

    mismatches = minpolyA // 8
    if te.seq[:minpolyA + mismatches].upper().count("A") >= minpolyA:
        lenPolyA = max(lenPolyA, te.seq[:minpolyA + mismatches].upper().count("A") + len(
            re.match("(.*?)[^A]", str(te.seq)[minpolyA:]).group()) - 2)
    if te.seq[:minpolyA + mismatches].upper().count("T") >= minpolyA:
        lenPolyA = max(lenPolyA, te.seq[:minpolyA + mismatches].upper().count("T") + len(
            re.match("(.*?)[^T]", str(te.seq)[minpolyA:]).group()) - 2)
    if te.seq[-(minpolyA + mismatches):].upper().count("A") >= minpolyA:
        lenPolyA = max(lenPolyA, te.seq[-(minpolyA + mismatches):].upper().count("A") + len(
            re.match("(.*?)[^A]", str(te.seq)[:len(te.seq) - minpolyA][::-1]).group()) - 2)
    if te.seq[-(minpolyA + mismatches):].upper().count("T") >= minpolyA:
        lenPolyA = max(lenPolyA, te.seq[-(minpolyA + mismatches):].upper().count("T") + len(
            re.match("(.*?)[^T]", str(te.seq)[:len(te.seq) - minpolyA][::-1]).group()) - 2)

    seq_name = str(te.id).split("#")[0]
    write_sequences_file(te, outputdir + "/temp/" + seq_name + ".fa")
    output = subprocess.run(['makeblastdb', '-in',  outputdir + "/temp/" + seq_name + ".fa", '-dbtype', 'nucl'], stdout=subprocess.PIPE, text=True)
    output = subprocess.run(["blastn -query " + outputdir + "/temp/" + seq_name + ".fa -db " + outputdir + "/temp/" + seq_name + ".fa -num_threads " + str(cores) + " -outfmt \"6 qseqid qstart qend sstart send\" -word_size 11 -gapopen 5 -gapextend 2 -reward 2 -penalty -3 | sed 's/#/-/g' > " + outputdir + "/temp/" + str(seq_name) + ".blast"], shell=True)
    blastresult = pd.read_table(outputdir + "/temp/" + str(seq_name) + ".blast", sep='\t', names=['qseqid', 'qstart', 'qend', 'sstart', 'send'],  dtype={'qseqid': str, 'qstart': int, 'qend': int, 'sstart': int, 'send': int} )
    blastHits = blastresult.shape[0]
    if blastresult.shape[0] > 1:
        blastresult = blastresult[1:]
        blastresult["len"] = abs(blastresult["qend"] - blastresult["qstart"])
        ltr_check = blastresult[(((blastresult.qend-blastresult.qstart) * (blastresult.send-blastresult.sstart)) > 0)
                                & (blastresult[['qstart', 'qend', 'sstart', 'send']].min(axis=1) < len(te) * 0.1)
                                & (blastresult[['qstart', 'qend', 'sstart', 'send']].max(axis=1) > len(te) * 0.9)]
        try:
            lenLTR = max(ltr_check[ltr_check.len >= minLTR].len)
        except:
            lenLTR = 0
        tir_check = blastresult[(((blastresult.qend-blastresult.qstart) * (blastresult.send-blastresult.sstart)) < 0)
                                & (blastresult[['qstart', 'qend', 'sstart', 'send']].min(axis=1) < len(te) * 0.1)
                                & (blastresult[['qstart', 'qend', 'sstart', 'send']].max(axis=1) > len(te) * 0.9)]
        try:
            lenTIR = max(tir_check[tir_check.len >= minTIR].len)
        except:
            lenTIR = 0

    return lenLTR, lenTIR, lenPolyA, blastHits


def find_profiles(te, outputdir, ref_profiles):
    # domains:      LTR retrotransposons/LINE                                                DIRS/Crypton
    domains_dict = {'_GAG_': 0, '_AP_': 0, '_INT_': 0, '_RT_': 0, '_RNaseH_': 0, '_ENV_': 0, '_PhageINT_': 0,
                    # PLE       TIRs         Helitron
                    '_EN_': 0, '_Tase_': 0, '_HEL_': 0, '_RPA_': 0, '_REP_': 0, '_OTU_': 0, '_SET_': 0,
                    # Maverick
                    '_Prp': 0, '_ATPase_': 0}
    seq_name = str(te.id).split("#")[0]
    write_sequences_file(te, outputdir + "/" + seq_name + "_putative_te.fa")

    output = subprocess.run(
        ['getorf', '-sequence', outputdir + "/" + seq_name + "_putative_te.fa", '-minsize', '300', '-outseq',
         outputdir + "/" + seq_name + "_putative_te_orf.fa"],
        stdout=subprocess.DEVNULL, stderr=subprocess.DEVNULL, text=True)

    output = subprocess.run(
        ['hmmscan', '--tblout', outputdir + "/" + seq_name + "_profiles_found.hmm", '-E', '10', '--noali', '--cpu',
         '1', ref_profiles, outputdir + "/" + seq_name + "_putative_te_orf.fa"],
         stdout=subprocess.DEVNULL, stderr=subprocess.DEVNULL, text=True)

    result = ""
    struc_domains = ""

    if os.path.exists(outputdir + "/" + seq_name + "_profiles_found.hmm"):
        command = 'grep -v "^#" ' + outputdir + '/' + seq_name + '_profiles_found.hmm | wc -l'
        process = subprocess.Popen(command, shell=True, stdout=subprocess.PIPE)
        hits = int(process.stdout.read())
        if hits > 0:
            command = 'grep -v "^#" ' + outputdir + '/' + seq_name + '_profiles_found.hmm | sed "s/_RVT_/_RT_/g" | awk \'{print $1"\t"$2"\t"$3"\t"$4"\t"$5"\t"$6"\t"$7"\t"$8"\t"$9"\t"$10}\' > ' + outputdir + "/" + seq_name + "_profiles_found.hmm_formatted"
            process = subprocess.Popen(command, shell=True)
            process.communicate()

            hmm_results = pd.read_table(outputdir + "/" + seq_name + "_profiles_found.hmm_formatted", sep='\t',
                                        names=['domain', 'acc', 'orf', 'acc_orf', 'evalue', 'score', 'A', 'B', 'C',
                                               'D'])
            hasDomains = False
            for domain in domains_dict.keys():
                best_hit = hmm_results[hmm_results.domain.str.contains(domain)].reset_index()
                if best_hit.shape[0] > 0:
                    hasDomains = True
                    domains_dict[domain] = best_hit.loc[0, 'evalue']
                    result += best_hit.loc[0, 'domain'] + ": " + str(best_hit.loc[0, 'evalue']) + ", "
                    struc_domains += domain.replace("_", "") + " "

            if hasDomains:
                result = "profiles: " + result[:-2]
            delete_files(outputdir + "/" + seq_name + "_profiles_found.hmm_formatted")
        delete_files(outputdir + "/" + seq_name + "_profiles_found.hmm")

    delete_files(outputdir + "/" + seq_name + "_putative_te_orf.fa")
    delete_files(outputdir + "/" + seq_name + "_putative_te.fa")

    return result, struc_domains


def find_blast_hits(te, outputdir, blastx_db, blastn_db):
    seq_name = str(te.id).split("#")[0]
    write_sequences_file(te, outputdir + "/" + seq_name + "_putative_te.fa")
    output = subprocess.run(
        ['blastx', '-query', outputdir + "/" + seq_name + "_putative_te.fa", '-db', blastx_db, '-out',
         outputdir + "/" + seq_name + "_putative_te.fa.blastx", '-num_threads', '1', "-outfmt",
         "6 sseqid pident evalue bitscore"],
        stdout=subprocess.PIPE, text=True)

    blastresult = pd.read_table(outputdir + "/" + seq_name + "_putative_te.fa.blastx", sep='\t',
                                names=['sseqid', 'pident', 'evalue', 'bitscore'])
    blastx_result = ""
    i = 0
    if blastresult.shape[0] > 0:
        blastx_result = "TE_BLRx: "
        while i < 3 and i < blastresult.shape[0]:
            blastx_result += blastresult.loc[0, "sseqid"] + ": " + str(blastresult.loc[0, "pident"]) + "%, "
            i += 1
        blastx_result = blastx_result[:-2]

    output = subprocess.run(
        ['tblastx', '-query', outputdir + "/" + seq_name + "_putative_te.fa", '-db', blastn_db, '-out',
         outputdir + "/" + seq_name + "_putative_te.fa.tblastx", '-num_threads', '1', "-outfmt",
         "6 sseqid pident evalue bitscore"],
        stdout=subprocess.PIPE, text=True)

    blastresult = pd.read_table(outputdir + "/" + seq_name + "_putative_te.fa.tblastx", sep='\t',
                                names=['sseqid', 'pident', 'evalue', 'bitscore'])
    blasttx_result = ""
    i = 0
    if blastresult.shape[0] > 0:
        blasttx_result = "TE_BLRtx: "
        while i < 3 and i < blastresult.shape[0]:
            blasttx_result += blastresult.loc[0, "sseqid"] + ": " + str(blastresult.loc[0, "pident"]) + "%, "
            i += 1
        blasttx_result = blasttx_result[:-2]

    delete_files(outputdir + "/" + seq_name + "_putative_te.fa")
    delete_files(outputdir + "/" + seq_name + "_putative_te.fa.tblastx")
    delete_files(outputdir + "/" + seq_name + "_putative_te.fa.blastx")
    return blastx_result, blasttx_result


def build_class_table_parallel(ref_tes, cores, outputdir, blastn_db, blastx_db, ref_profiles, do_blast):
    if not os.path.exists(outputdir + "/denovoLibTEs_PC.classif"):
        tes = [te for te in SeqIO.parse(ref_tes, "fasta")]
        n = len(tes)
        seqs_per_procs = int(n / cores)
        remain = n % cores
        ini_per_thread = []
        end_per_thread = []

        # indexing the Pfam database if needed
        if not os.path.exists(ref_profiles+".h3m"):
            output = subprocess.run(['hmmpress', ref_profiles],
            stdout=subprocess.DEVNULL, stderr=subprocess.DEVNULL, text=True)

        for p in range(cores):
            if p < remain:
                init = p * (seqs_per_procs + 1)
                end = n if init + seqs_per_procs + 1 > n else init + seqs_per_procs + 1
            else:
                init = p * seqs_per_procs + remain
                end = n if init + seqs_per_procs > n else init + seqs_per_procs
            ini_per_thread.append(init)
            end_per_thread.append(end)

        # Run in parallel the checking
        pool = multiprocessing.Pool(processes=cores)
        create_output_folders(outputdir + "/temp")
        localresults = [pool.apply_async(build_class_table,
                                         args=[tes[ini_per_thread[x]:end_per_thread[x]], ref_profiles, outputdir, blastn_db,
                                               blastx_db, do_blast, cores]) for x in range(cores)]

        local_dfs = [p.get() for p in localresults]
        class_df = pd.DataFrame(
            columns=['Seq_name', 'length', 'strand', 'confused', 'class', 'order', 'Wcode', 'sFamily', 'CI', 'coding',
                     'struct', 'other'])
        for i in range(len(local_dfs)):
            class_df = pd.concat([class_df, local_dfs[i]], ignore_index=True)
        pool.close()

        class_df.to_csv(outputdir + "/denovoLibTEs_PC.classif", header=True, sep='\t', index=False)

        new_tes_user = []
        for te in SeqIO.parse(ref_tes, "fasta"):
            te.id = str(te.id).split("#")[0]
            te.description = ""
            new_tes_user.append(te)

        os.system("rm -r " + outputdir + "/temp")
        write_sequences_file(new_tes_user, outputdir + "/new_user_lib.fa")
    else:
        print("MESSAGE: TE Feature table was already created, please verify or change the output directory")


def build_class_table(ref_tes, ref_profiles, outputdir, blastn_db, blastx_db, do_blast, cores):
    class_df = pd.DataFrame(
        columns=['Seq_name', 'length', 'strand', 'confused', 'class', 'order', 'Wcode', 'sFamily', 'CI', 'coding',
                 'struct', 'other'])
    orders = ['LTR', 'TRIM', 'LARD', 'LINE', 'SINE', 'PLE', 'DIRS', 'TIR', 'MITE', 'HELITRON', 'MAVERICK', 'CRYPTON']
    superfamilies = ["COPIA", "GYPSY", "BELPAO", "ERV", "PLE", "DIRS", "NGARO", "VIPER", "R2", "RTE", "JOCKEY", "L1",
                     "I", "R1", "CR1", "LOA", "L2", "SINE", "TC1MARINER", "HAT", "MUTATOR", "MERLIN", "TRANSIB", "P",
                     "PIGGYBAC", "PIFHARBINGER", "CACTA", "MULE", "CMC", "MITE", "HELITRON", "MAVERICK", "CRYPTON"]
    classes = ["CLASSI", "CLASSII", "RETROTRANSPOSON", "TRANSPOSON"]
    for te in ref_tes:
        seq_name = str(te.id).split("#")[0]
        classification = str(te.id).split("#")[1].upper().replace("-", "")
        length = len(str(te.seq))
        class_given = ""
        if len(classification.split("/")) == 3:
            # The most complete case Class / order / superfamily
            class_given = classification.split("/")[0]
            order_given = classification.split("/")[1]
            superfamily = classification.split("/")[2]
        elif len(classification.split("/")) == 2:
            # The most common case Class / Order or order / superfamily
            order_given = classification.split("/")[0]
            superfamily = classification.split("/")[1]

            if superfamily in orders and order_given in classes:  # it's the form Class / order
                class_given = order_given
                order_given = superfamily
                superfamily = "NA"

        elif len(classification.split("/")) == 1:
            # the most incomplete but still fine to work with
            order_given = classification

            if order_given in superfamilies:  # it's a superfamily
                superfamily = order_given
            elif order_given in classes:  # it's a class
                class_given = order_given
                order_given = "NA"
                superfamily = "NA"
            else:  # it's actually an order
                superfamily = "NA"


        if superfamily in superfamilies:
            # superfamily found !!!
            sFamily = superfamily
            order = ""
            sFamily_index = superfamilies.index(superfamily)
            if sFamily_index <= 3:
                order = "LTR"
            elif sFamily_index == 4:
                order = "PLE"
            elif sFamily_index >= 5 and sFamily_index <= 7:
                order = "DIRS"
            elif sFamily_index >= 8 and sFamily_index <= 16:
                order = "LINE"
            elif sFamily_index == 17:
                order = "SINE"
            elif sFamily_index >= 18 and sFamily_index <= 28:
                order = "TIR"
            elif sFamily_index == 29:
                order = "MITE"
            elif sFamily_index == 30:
                order = "Helitron"
            elif sFamily_index == 31:
                order = "Maverick"
            elif sFamily_index == 32:
                order = "Crypton"

            if sFamily_index <= 17:
                classTE = "CLASSI"
            else:
                classTE = "CLASSII"
        elif order_given in orders:
            # Well, superfamily didn't find, but I found the order
            sFamily = "NA"
            order = order_given
            orden_index = orders.index(order)
            if orden_index <= 6:
                classTE = "CLASSI"
            else:
                classTE = "CLASSII"
        elif class_given != "":
            classTE = class_given
            order = "NA"
            sFamily = "NA"
        else:
            #print("WARNING: I didn't find the superfamily, neither the order: "+order_given+"/"+superfamily)
            classTE = "Unclassified"
            order = "Unclassified"
            sFamily = "Unclassified"

        lenLTR, lenTIR, lenPolyA, blastHits = find_TRs2(te, outputdir, 10, 10, 10, cores)
        profiles, struc_dom = find_profiles(te, outputdir, ref_profiles)
        if do_blast:
            blastx, blasttx = find_blast_hits(te, outputdir, blastx_db, blastn_db)
        else:
            blastx, blasttx = "", ""

        terminals = ''
        terminals_list = []
        if lenLTR > 0:
            terminals_list.append("termLTR: "+str(lenLTR))
        if lenTIR > 0:
            terminals_list.append("termTIR: "+str(lenTIR))
        if lenPolyA > 0:
            terminals_list.append("polyAtail: "+str(lenPolyA))
        if len(terminals_list) > 0:
            terminals = 'TermRepeats: ' + ' '.join(terminals_list) + ';'

        final_coding_string = ""
        if blasttx != "":
            final_coding_string = blasttx + "; "
        if blastx != '':
            final_coding_string += blastx + ";"
        if profiles != "":
            final_coding_string += profiles

        if blastx == "" and blasttx == "" and profiles == "":
            final_coding_string = "NA"

        ### Heuristcs to purge simple repeats and chimeric elements
        other = 'other=NA'
        if blastHits > 40:  ### MAGIC NUMBER; INCLUDE AS OPTIONAL PARAMETHER
            other = 'other=Satellites/Simple Repeats found'
        elif lenLTR > length * 0.35:  ### MAGIC NUMBER; INCLUDE AS OPTIONAL PARAMETHER
            other = 'other=Chimeric evidence found'

        class_df = pd.concat([class_df, pd.DataFrame(
            {'Seq_name': seq_name, 'length': [length], 'strand': '+', 'confused': 'False', 'class': classTE,
             'order': order, 'Wcode': 'NA', 'sFamily': sFamily, 'CI': [0],
             'coding': 'coding=(' + final_coding_string + ')',
             'struct': 'struct=(TElength: ' + str(length) + 'bps; ' + terminals + ' ' + struc_dom + ')',
             'other': other})], ignore_index=True)

    return class_df


def count_domains_by_order(profiles, order):
    right_doms = 0
    other_doms = 0
    if order == "LINE":
        right_doms = len(
            [x for x in profiles.split(",") if '_RT_' in x or '_EN_' in x or '_RNaseH_' in x or '_GAG_' in x])
        other_doms = len([x for x in profiles.split(",") if '_AP_' in x or '_INT_' in x or '_ENV_' in x or '_Tase_' in x
                          or '_HEL_' in x or '_RPA_' in x or '_REP_' in x or '_OTU_' in x or '_SET_' in x or '_Prp' in x
                          or '_ATPase_' in x or '_PhageINT_' in x])
    elif order == "LTR":
        right_doms = len([x for x in profiles.split(",") if
                        '_GAG_' in x or '_AP_' in x or '_INT_' in x or '_RT_' in x or '_RNaseH_' in x or '_ENV_' in x])
        other_doms = len([x for x in profiles.split(",") if '_EN_' in x or '_Tase_' in x or '_HEL_' in x or '_RPA_' in x
                          or '_REP_' in x or '_OTU_' in x or '_SET_' in x or '_Prp' in x or '_ATPase_' in x or '_PhageINT_' in x])

    elif order == "DIRS":
        right_doms = len([x for x in profiles.split(",") if
                        '_GAG_' in x or '_RT_' in x or '_RNaseH_' in x or '_PhageINT_' in x])
        other_doms = len([x for x in profiles.split(",") if '_AP_' in x or '_INT_' in x or '_ENV_' in x or '_EN_' in x
                          or '_Tase_' in x or '_HEL_' in x or '_RPA_' in x or '_REP_' in x or '_OTU_' in x or '_SET_'
                          in x or '_Prp' in x or '_ATPase_' in x])

    elif order == "TIR":
        right_doms = len([x for x in profiles.split(",") if '_Tase_' in x])
        other_doms = len([x for x in profiles.split(",") if '_GAG_' in x or '_AP_' in x or '_INT_' in x or '_RT_' in x
                          or '_RNaseH_' in x or '_ENV_' in x or '_EN_' in x or '_HEL_' in x or '_RPA_' in x or '_REP_'
                          in x or '_OTU_' in x or '_SET_' in x or '_Prp' in x or '_ATPase_' in x or '_PhageINT_' in x])

    elif order == "HELITRON":
        right_doms = len([x for x in profiles.split(",") if '_HEL_' in x or '_EN_' in x or '_RPA_' in x or '_REP_' in x
                          or '_OTU_' in x or '_SET_' in x])
        other_doms = len([x for x in profiles.split(",") if
                          '_GAG_' in x or '_AP_' in x or '_INT_' in x or '_RT_' in x or '_RNaseH_' in x or '_ENV_' in x
                          or '_Tase_' in x or '_Prp' in x or '_ATPase_' in x or '_PhageINT_' in x])

    elif order == "MAVERICK":
        right_doms = len([x for x in profiles.split(",") if '_Prp' in x or '_ATPase_' in x or '_INT_' in x or '_AP_' in x])
        other_doms = len([x for x in profiles.split(",") if
                          '_GAG_' in x or '_AP_' in x or '_INT_' in x or '_RT_' in x or '_RNaseH_' in x or '_ENV_' in x
                          or '_EN_' in x or '_Tase_' in x or '_HEL_' in x or '_RPA_' in x or '_REP_' in x or '_OTU_' in x
                          or '_SET_' in x or '_PhageINT_' in x])

    elif order == "CRYPTON":
        right_doms = len([x for x in profiles.split(",") if '_PhageINT_' in x])
        other_doms = len([x for x in profiles.split(",") if '_GAG_' in x or '_AP_' in x or '_INT_' in x or '_RT_' in x
                          or '_RNaseH_' in x or '_ENV_' in x or '_EN_' in x or '_HEL_' in x or '_RPA_' in x or '_REP_'
                          in x or '_OTU_' in x or '_SET_' in x or '_Prp' in x or '_ATPase_' in x or '_Tase_' in x])

    return right_doms, other_doms


def inferring_domains(input_profiles):
    inferred = False
    new_class = 0

    if len(input_profiles) > 0:
        profiles = input_profiles[0]
        # Class I or II ?
        class2_doms = len(
            [x for x in profiles.split(",") if '_Tase_' in x or '_HEL_' in x or '_RPA_' in x or '_REP_' in x or '_OTU_'
             in x or '_SET_' in x or '_Prp' in x or '_ATPase_' in x  or '_PhageINT_' in x])
        class1_doms = len([x for x in profiles.split(",") if '_GAG_' in x or '_AP_' in x or '_INT_' in x or '_RT_' in x or
                          '_RNaseH_' in x or '_ENV_' in x or '_EN_' in x])

        if class1_doms > 0 and class2_doms == 0:
            # Retrotransposon !
            inferred = True
            new_class = orders_superfamilies.index("CLASSI") + 2

            # which order?
            count_orders = 0
            for ord in ['LTR', 'LINE', 'DIRS']:
                good_doms, other_doms = count_domains_by_order(profiles, ord)
                if good_doms > 0 and other_doms == 0:
                    new_class = orders_superfamilies.index(ord) + 2
                    count_orders += 1

            if count_orders > 1:  # It wasn't possible to distinguish between Class I's orders
                new_class = orders_superfamilies.index("CLASSI") + 2

        elif class1_doms == 0 and class2_doms > 0:
            # Transposon !
            inferred = True
            new_class = orders_superfamilies.index("CLASSII") + 2

            # which order?
            count_orders = 0
            for ord in ['TIR', 'HELITRON', 'MAVERICK', 'CRYPTON']:
                good_doms, other_doms = count_domains_by_order(profiles, ord)
                if good_doms > 0 and other_doms == 0:
                    new_class = orders_superfamilies.index(ord) + 2
                    count_orders += 1

            if count_orders > 1:  # It wasn't possible to distinguish between Class II's orders
                new_class = orders_superfamilies.index("CLASSII") + 2

    return inferred, new_class


def decision_tree_rules(struc_table, profiles, i, keep_seqs, minDomLTR, num_copies, orders, minFLNA, kept_seqs_record, ref_tes, automatic):
    status = -1  # 0 = delete, 1 = keep, -1 = Manual Inspection classified module, -3 = unclassified module
    reason = ""
    superFamily = str(struc_table.at[i, "sFamily"])

    #### Class 1 Retrotransposons
    if str(struc_table.at[i, "order"]).upper() == 'LINE' or str(struc_table.at[i, "order"]).upper() == 'SINE':
        if len(profiles) > 0 and len(profiles[0].split(",")) >= 1:
            rigth_doms, other_doms = count_domains_by_order(profiles[0], "LINE")
            if rigth_doms > 0 and other_doms == 0:
                keep_seqs.append(struc_table.at[i, "Seq_name"])
                kept_seqs_record.append([x for x in SeqIO.parse(ref_tes, "fasta") if
                                         str(x.id).split("#")[0] == struc_table.at[i, "Seq_name"]][0])
                if superFamily in orders_superfamilies:
                    orders.append(orders_superfamilies.index(superFamily) + 2)
                else:
                    orders.append(orders_superfamilies.index("LINE") + 2)
                reason = "It's a LINE !!! Kept for having at least one domain !"
                status = 1
            else:
                status = -1
                if automatic != 'F':
                    reason = "Sent to manual inspection because having non-LINE domains !"
                else:
                    reason = "Marked as incomplete TE for having non-LINE domains"
        elif len(profiles) == 0 and "polyAtail" in struc_table.at[i, "struct"] and (
                num_copies[struc_table.at[i, "Seq_name"]][0] >= minFLNA or num_copies[struc_table.at[i, "Seq_name"]][1]
                >= minFLNA):
            keep_seqs.append(struc_table.at[i, "Seq_name"])
            orders.append(orders_superfamilies.index("SINE") + 2)
            kept_seqs_record.append([x for x in SeqIO.parse(ref_tes, "fasta") if
                                     str(x.id).split("#")[0] == struc_table.at[i, "Seq_name"]][0])
            reason = "It's a SINE !!! Kept for having polyA tail, no domains and at least " + str(
                minFLNA) + " full-length fragments or copies !"
            status = 1
        else:
            if automatic != 'F':
                reason = "Sent to manual inspection for don't having any domains or PolyA"
            else:
                reason = "Marked as incomplete TE for don't having any domains or PolyA"
            status = -1
    elif str(struc_table.at[i, "order"]).upper() == 'LTR' or str(struc_table.at[i, "order"]).upper() == 'LARD' or \
            str(struc_table.at[i, "order"]).upper() == 'TRIM':
        status_ltr = False
        if "termLTR: " in struc_table.at[i, "struct"]:
            if len(profiles) > 0 and len(profiles[0].split(",")) >= minDomLTR:
                rigth_doms, other_doms = count_domains_by_order(profiles[0], "LTR")
                if rigth_doms > minDomLTR and other_doms == 0:
                    reason = "It's a LTR-RT !!! Kept for having at least " + str(minDomLTR) + " domains and LTRs !"
                    status_ltr = True
                    status = 1
                else:
                    status = -1
                    if automatic != 'F':
                        reason = "Sent to manual inspection for having non-LTR-RTs domains !"
                    else:
                        reason = "Marked as incomplete TE for having non-LTR-RTs domains !"
            elif len(profiles) > 0 and len(profiles[0].split(",")) > 0 and struc_table.at[i, "order"] == 'LTR':
                rigth_doms, other_doms = count_domains_by_order(profiles[0], "LTR")
                if rigth_doms > 0 and other_doms == 0:
                    reason = "It's a truncated LTR-RT !!! Kept for having " + str(
                        len(profiles[0].split(","))) + " domains and LTRs !"
                    status_ltr = True
                    status = 1
                else:
                    status = -1
                    if automatic != 'F':
                        reason = "Sent to manual inspection for don't having non-LTR-RTs domains !"
                    else:
                        reason = "Marked as incomplete TE for don't having non-LTR-RTs domains !"
            elif len(profiles) == 0 and (num_copies[struc_table.at[i, "Seq_name"]][0] >= minFLNA or
                                         num_copies[struc_table.at[i, "Seq_name"]][1] >= minFLNA):
                keep_seqs.append(struc_table.at[i, "Seq_name"])
                kept_seqs_record.append([x for x in SeqIO.parse(ref_tes, "fasta") if
                                         str(x.id).split("#")[0] == struc_table.at[i, "Seq_name"]][0])
                if int(struc_table.at[i, "length"]) <= 2500:
                    name = "TRIM"
                else:
                    name = "LARD"
                orders.append(orders_superfamilies.index(name) + 2)
                reason = "It's a " + name + " !!! Kept for having LTRs, no domains and at least " + str(
                    minFLNA) + " full-length fragments or copies !"
                status = 1
        else:
            # no profiles and no LTRs
            if len(profiles) == 0:
                if automatic != 'F':
                    reason = "Sent to manual inspection for don't having any domains neither LTRs!"
                else:
                    reason = "Marked as incomplete TE for don't having any domains neither LTRs!"
            else:
                if automatic != 'F':
                    reason = "Sent to manual inspection for don't having at least " + str(minDomLTR) + " domains and LTRs!"
                else:
                    reason = "Marked as incomplete TE for don't having at least " + str(minDomLTR) + " domains and LTRs!"
            status = -1

        if status_ltr:
            if superFamily.upper().replace("-", "") in orders_superfamilies:
                orders.append(orders_superfamilies.index(superFamily.upper().replace("-", "")) + 2)
            else:
                orders.append(orders_superfamilies.index("LTR") + 2)
            keep_seqs.append(struc_table.at[i, "Seq_name"])
            kept_seqs_record.append([x for x in SeqIO.parse(ref_tes, "fasta") if
                                     str(x.id).split("#")[0] == struc_table.at[i, "Seq_name"]][0])
    elif str(struc_table.at[i, "order"]).upper() == 'DIRS':
        # manual inspection in classified module
        if len(profiles) > 0 and len(profiles[0].split(",")) >= 1:
            rigth_doms, other_doms = count_domains_by_order(profiles[0], "DIRS")
            if rigth_doms > 0 and other_doms == 0:
                keep_seqs.append(struc_table.at[i, "Seq_name"])
                kept_seqs_record.append([x for x in SeqIO.parse(ref_tes, "fasta") if
                                         str(x.id).split("#")[0] == struc_table.at[i, "Seq_name"]][0])
                orders.append(orders_superfamilies.index("DIRS") + 2)
                reason = "It's a DIRS !!! Kept for having at least one domain !"
                status = 1
            else:
                status = -1
                if automatic != 'F':
                    reason = "Sent to manual inspection for having non-DIRS domains !"
                else:
                    reason = "Marked as incomplete TE for having non-DIRS domains !"
        else:
            status = -1
            if automatic != 'F':
                reason = "Sent to manual inspection for don't having domains !"
            else:
                reason = "Marked as incomplete TE for don't having domains !"
    elif str(struc_table.at[i, "order"]).upper() == 'PLE':
        # manual inspection in classified module
        if automatic != 'F':
            reason = "Sent to manual inspection"
        else:
            reason = "Marked as incomplete TE"
        status = -1

    #### Class 2 DNA Transposons
    elif str(struc_table.at[i, "order"]).upper() == 'TIR' or str(struc_table.at[i, "order"]).upper() == 'MITE':
        if "termTIR: " in struc_table.at[i, "struct"]:
            if len(profiles) > 0 and len(profiles[0].split(",")) > 0:
                rigth_doms, other_doms = count_domains_by_order(profiles[0], "TIR")
                if rigth_doms > 0 and other_doms == 0:
                    keep_seqs.append(struc_table.at[i, "Seq_name"])
                    kept_seqs_record.append([x for x in SeqIO.parse(ref_tes, "fasta") if
                                             str(x.id).split("#")[0] == struc_table.at[i, "Seq_name"]][0])

                    if superFamily.upper().replace("-", "") in orders_superfamilies:
                        orders.append(orders_superfamilies.index(superFamily.upper().replace("-", "")) + 2)
                    else:
                        orders.append(orders_superfamilies.index("TIR") + 2)
                    reason = "It's a TIR !!! Kept for having at least one domain !"
                    status = 1
                else:
                    status = -1
                    if automatic != 'F':
                        reason = "Sent to manual inspection because having non-TIR domains !"
                    else:
                        reason = "Marked as incomplete TE because having non-TIR domains !"
            else:
                if num_copies[struc_table.at[i, "Seq_name"]][0] >= minFLNA or num_copies[struc_table.at[i, "Seq_name"]][
                1] >= minFLNA:
                    keep_seqs.append(struc_table.at[i, "Seq_name"])
                    kept_seqs_record.append([x for x in SeqIO.parse(ref_tes, "fasta") if
                                             str(x.id).split("#")[0] == struc_table.at[i, "Seq_name"]][0])
                    orders.append(orders_superfamilies.index("MITE") + 2)
                    reason = "It's a MITE !!! Kept for having TIRs, no domains and at least " + str(
                        minFLNA) + " full-length fragments or copies !"
                    status = 1
                else:
                    status = -1
                    if automatic != 'F':
                        reason = "Sent to manual inspection because having TIRs, no domains but less than " + str(
                        minFLNA) + " full-length fragments or copies !"
                    else:
                        reason = "Marked as incomplete TE because having TIRs, no domains but less than " + str(
                        minFLNA) + " full-length fragments or copies !"
        else:
            # no profiles and no TIRs
            status = -1
            if automatic != 'F':
                reason = "Sent to manual inspection for don't having TIRs neither domains !"
            else:
                reason = "Marked as incomplete TE for don't having TIRs neither domains !"

    elif str(struc_table.at[i, "order"]).upper() == 'CRYPTON':
        if len(profiles) > 0 and len(profiles[0].split(",")) >= 1:
            rigth_doms, other_doms = count_domains_by_order(profiles[0], "CRYPTON")
            if rigth_doms > 0 and other_doms == 0:
                keep_seqs.append(struc_table.at[i, "Seq_name"])
                kept_seqs_record.append([x for x in SeqIO.parse(ref_tes, "fasta") if
                                         str(x.id).split("#")[0] == struc_table.at[i, "Seq_name"]][0])
                orders.append(orders_superfamilies.index("CRYPTON") + 2)
                reason = "It's a CRYPTON !!! Kept for having at least one domain !"
                status = 1
            else:
                status = -1
                if automatic != 'F':
                    reason = "Sent to manual inspection for having non-CRYPTON domains !"
                else:
                    reason = "Marked as incomplete TE for having non-CRYPTON domains !"
        else:
            status = -1
            if automatic != 'F':
                reason = "Sent to manual inspection for don't having domains !"
            else:
                reason = "Marked as incomplete TE for don't having domains !"
    elif str(struc_table.at[i, "order"]).upper() == 'HELITRON':
        if len(profiles) > 0:
            rigth_doms, other_doms = count_domains_by_order(profiles[0], "HELITRON")
            if rigth_doms > 0 and other_doms == 0:
                keep_seqs.append(struc_table.at[i, "Seq_name"])
                kept_seqs_record.append([x for x in SeqIO.parse(ref_tes, "fasta") if
                                         str(x.id).split("#")[0] == struc_table.at[i, "Seq_name"]][0])
                orders.append(orders_superfamilies.index("HELITRON") + 2)
                reason = "It's a Helitron !!! Kept for having Helicase !"
                status = 1
            else:
                status = -1
                if automatic != 'F':
                    reason = "Sent to manual inspection for having non-Helitron domains"
                else:
                    reason = "Marked as incomplete TE for having non-Helitron domains"
        elif "helitronExtremities: " in struc_table.at[i, "struct"] :
            keep_seqs.append(struc_table.at[i, "Seq_name"])
            kept_seqs_record.append([x for x in SeqIO.parse(ref_tes, "fasta") if
                                     str(x.id).split("#")[0] == struc_table.at[i, "Seq_name"]][0])
            orders.append(orders_superfamilies.index("HELITRON") + 2)
            reason = "It's a Helitron !!! Kept for having helitron Extremities !"
            status = 1
        else:
            status = -1
            if automatic != 'F':
                reason = "Sent to manual inspection for don't having Helitron Extremities neither helicase !"
            else:
                reason = "Marked as incomplete TE for don't having Helitron Extremities neither helicase !"
    elif str(struc_table.at[i, "order"]).upper() == 'MAVERICK':
        if len(profiles) > 0 and len(profiles[0].split(",")) >= 1:
            rigth_doms, other_doms = count_domains_by_order(profiles[0], "MAVERICK")
            if rigth_doms > 0 and other_doms == 0:
                keep_seqs.append(struc_table.at[i, "Seq_name"])
                kept_seqs_record.append([x for x in SeqIO.parse(ref_tes, "fasta") if
                                         str(x.id).split("#")[0] == struc_table.at[i, "Seq_name"]][0])
                orders.append(orders_superfamilies.index("MAVERICK") + 2)
                reason = "It's a Maverick !!! Kept for having at least one domain !"
                status = 1
            else:
                status = -1
                if automatic != 'F':
                    reason = "Sent to manual inspection for having non-Maverick domains !"
                else:
                    reason = "Marked as incomplete TE for having non-Maverick domains !"
        else:
            status = -1
            if automatic != 'F':
                reason = "Sent to manual inspection for don't having domains !"
            else:
                reason = "Marked as incomplete TE for don't having domains !"
    else:
        status = -3
        reason = "Sent to unclassified module due to its order is unknown !"
    return status, reason


def manual_inspection(genome, outputdir, te_library, seqs_to_mi, seqID_list, struc_table, te_aid, repet, automatic, pre, plots_dir,
                      gff_files, min_perc_model, seqs_to_module3, keep_seqs, orders, kept_seqs_record, non_curated,
                      num_copies):
    if te_aid == 'Y':
        run_te_aid_parallel(tools_path + "/TE-Aid-master/", genome, te_library, outputdir, cores, min_perc_model)

    seqs_manu = 0
    ele_number = 0
    orders_incomplete = []
    for i in range(struc_table.shape[0]):
        if (automatic == 'M' or struc_table.at[i, "Seq_name"] in seqs_to_mi) and struc_table.at[i, "Seq_name"] in seqID_list:
            if automatic != 'F':
                codings = struc_table.at[i, "coding"].replace("coding=(", " ").replace(")", "")
                codigs = codings.split(";")
                print("[" + str(ele_number + 1) + "/" + str(struc_table.shape[0]) + "] Seq Name: " + str(struc_table.at[i, "Seq_name"]))
                print("Length: " + str(struc_table.at[i, "length"]))
                print("Class: " + str(struc_table.at[i, "class"]) + " / " + str(
                    struc_table.at[i, "order"]) + " / " + str(struc_table.at[i, "sFamily"]))
                print("Copies: FLF = " + str(num_copies[struc_table.at[i, "Seq_name"]][0]) + " / FLC = " + str(
                    num_copies[struc_table.at[i, "Seq_name"]][1]))
                print("Coding: ")
                for cod in codigs:
                    print(cod)
                print(struc_table.at[i, "struct"])
                print(struc_table.at[i, "other"])

                if te_aid == 'N':
                    fig = plt.figure(figsize=(20, 10))
                    # setting values to rows and column variables
                    rows = 1
                    columns = 3

                    # reading images
                    try:
                        Image1 = cv2.imread(plots_dir + '/' + pre + '_' + struc_table.at[i, "Seq_name"] + '.png')

                        # Adds a subplot at the 1st position
                        fig.add_subplot(rows, columns, 1)

                        # showing image
                        plt.imshow(Image1)
                        plt.axis('off')
                        plt.title("REPET")

                        # obtaining the positions of Blastn, tBlastn y blastx from GFF:
                        posBltn, postBlasx, posBlastx, posTR = processing_gff(struc_table.at[i, "Seq_name"], gff_files, pre)

                        # ploting gff files
                        ax = fig.add_subplot(rows, columns, 2)
                        ax.broken_barh([(0, int(struc_table.at[i, "length"]))], (0, 2), facecolors='black')
                        ax.broken_barh(posBltn, (3, 2), facecolors='blue', alpha=0.2)
                        ax.broken_barh(postBlasx, (6, 2), facecolors='orange', alpha=0.2)
                        ax.broken_barh(posBlastx, (9, 2), facecolors='red', alpha=0.2)
                        ax.broken_barh(posTR, (12, 2), facecolors='green', alpha=0.2)
                        ax.set_ylim(0, 14)
                        ax.set_xlim(0, int(struc_table.at[i, "length"]))
                        ax.set_yticks([1, 4, 7, 10, 13])
                        ax.set_yticklabels(['TE seq', 'BLASTn', 'tBLASTx', 'BLASTx', 'TRs'])
                        ax.set_xlabel('TE consensus')
                        ax.set_title('REPET GFF files')

                        if os.path.exists(outputdir + '/MSA_plots/' + struc_table.at[i, "Seq_name"] + '.copies.cialign_output.png'):
                            Image2 = cv2.imread(outputdir + '/MSA_plots/' + struc_table.at[i, "Seq_name"] + '.copies.cialign_output.png')
                            fig.add_subplot(rows, columns, 3)
                            # showing image
                            plt.imshow(Image2)
                            plt.axis('off')
                            plt.title("MSA")
                        else:
                            ax = fig.add_subplot(rows, columns, 3)
                            ax.set_title('MSA')
                            ax.axis([0, 10, 0, 10])
                            ax.text(4, 5, 'Not enough TE copies', style='italic', fontsize=14, fontweight='bold')

                        plt.tight_layout()
                        plt.draw()
                        plt.pause(1)
                    except Exception as ex:
                        print("WARNING: The plot from " + struc_table.at[i, "Seq_name"] + " has a problem:")
                        print(ex)

                else:
                    # create figure
                    fig = plt.figure(figsize=(20, 10))

                    # setting values to rows and column variables
                    rows = 1
                    if repet:
                        columns = 3 + 1
                    else:
                        columns = 1 + 1
                    """if automatic != 'M':
                        columns += 1"""
                    try:
                        pos_te_aid = 1
                        if repet:
                            # reading images
                            Image1 = cv2.imread(plots_dir + '/' + pre + '_' + struc_table.at[i, "Seq_name"] + '.png')

                            # Adds a subplot at the 1st position
                            fig.add_subplot(rows, columns, 1)

                            # showing image
                            plt.imshow(Image1)

                            plt.axis('off')
                            plt.title("REPET")

                            # obtaining the positions of Blastn, tBlastn y blastx from GFF:
                            posBltn, postBlasx, posBlastx, posTR = processing_gff(struc_table.at[i, "Seq_name"],
                                                                                  gff_files, pre)
                            # ploting gff files
                            ax = fig.add_subplot(rows, columns, 2)
                            ax.broken_barh([(0, int(struc_table.at[i, "length"]))], (0, 2), facecolors='black')
                            ax.broken_barh(posBltn, (3, 2), facecolors='blue', alpha=0.2)
                            ax.broken_barh(postBlasx, (6, 2), facecolors='orange', alpha=0.2)
                            ax.broken_barh(posBlastx, (9, 2), facecolors='red', alpha=0.2)
                            ax.broken_barh(posTR, (12, 2), facecolors='green', alpha=0.2)
                            ax.set_ylim(0, 14)
                            ax.set_xlim(0, int(struc_table.at[i, "length"]))
                            ax.set_yticks([1, 4, 7, 10, 13])
                            ax.set_yticklabels(['TE seq', 'BLASTn', 'tBLASTx', 'BLASTx', 'TRs'])
                            ax.set_xlabel('TE consensus')
                            ax.set_title('REPET GFF files')
                            pos_te_aid = 3

                        try:
                            pages = convert_from_path(
                                outputdir + '/te_aid/' + struc_table.at[i, "Seq_name"] + '.fa.c2g.pdf')

                            # Saving pages in jpeg format
                            for page in pages:
                                page.save(outputdir + '/te_aid/' + struc_table.at[
                                        i, "Seq_name"] + '.fa.c2g.jpeg', 'JPEG')

                            Image2 = cv2.imread(outputdir + '/te_aid/' + struc_table.at[
                                    i, "Seq_name"] + '.fa.c2g.jpeg')

                            # Adds a subplot at the 2nd position
                            fig.add_subplot(rows, columns, pos_te_aid)

                            # showing image
                            plt.imshow(Image2)
                            plt.axis('off')
                            plt.title("TE-aid")

                            # if automatic != 'M':
                            if os.path.exists(outputdir + '/MSA_plots/' + struc_table.at[
                                i, "Seq_name"] + '.copies.cialign_output.png'):
                                Image2 = cv2.imread(
                                    outputdir + '/MSA_plots/' + struc_table.at[i, "Seq_name"] + '.copies.cialign_output.png')
                                fig.add_subplot(rows, columns, pos_te_aid + 1)
                                # showing image
                                plt.imshow(Image2)
                                plt.axis('off')
                                plt.title("MSA")
                            else:
                                ax = fig.add_subplot(rows, columns, pos_te_aid + 1)
                                ax.set_title('MSA')
                                plt.axis('off')
                                ax.axis([0, 10, 0, 10])
                                ax.text(4, 5, 'Not enough TE copies', style='italic', fontsize=14, fontweight='bold')

                            plt.tight_layout()
                            plt.draw()
                            plt.pause(1)
                        except Exception as ex:
                            print("WARNING: The plot from " + struc_table.at[i, "Seq_name"] + " has a problem:")
                            print(ex)

                        delete_files(outputdir + '/te_aid/' + struc_table.at[i, "Seq_name"] + '.fa.c2g.jpeg')
                    except FileNotFoundError:
                        print("WARNING: Element " + struc_table.at[i, "Seq_name"] + "couldn't be processed")

                try:
                    if str(struc_table.at[i, "sFamily"]) in orders_superfamilies:
                        current_clas = orders_superfamilies.index(str(struc_table.at[i, "sFamily"])) + 2
                    elif str(struc_table.at[i, "order"]) in orders_superfamilies:
                        current_clas = orders_superfamilies.index(str(struc_table.at[i, "order"])) + 2
                    elif str(struc_table.at[i, "class"]) in orders_superfamilies:
                        current_clas = orders_superfamilies.index(str(struc_table.at[i, "class"])) + 2
                    else:
                        current_clas = orders_superfamilies.index("UNCLASSIFIED") + 2
                    keep = input(
                            "Keep the sequence?[enter: current ("+str(current_clas)+"), 1: remove, 2: LTR, 3: COPIA, 4: "
                    "GYPSY, 5: BELPAO, 6: ERV, 7: TRIM, 8: LARD, 9: LINE, 10: SINE, 11: R2, 12: RTE, 13: JOCKEY, 14: L1,"
                    " 15: I, 16: R1, 17: CR1, 18: LOA, 19: L2, 20: PLE, 21: DIRS, 22: NGARO, 23: VIPER, 24: TIR, 25: MITE"
                    ", 26: TC1MARINER,  27: HAT, 28: MUTATOR, 29: MERLIN, 30: TRANSIB, 31: P, 32: PIGGYBAC, 33: PIFHARBINGER,"
                    "  34: CACTA, 35: MULE, 36: CMC, 37: HELITRON, 38: MAVERICK, 39: CRYPTON, 40: UNCLASSIFIED, 41: "
                    "CLASSI, 42: CLASSII] ") or current_clas
                    keep = int(keep)
                except ValueError:
                    keep = -1

                while keep < 1 or keep > 43:
                    try:
                        print('You must indicate a number between 1 and 43.')
                        if str(struc_table.at[i, "sFamily"]) in orders_superfamilies:
                            current_clas = orders_superfamilies.index(str(struc_table.at[i, "sFamily"])) + 2
                        elif str(struc_table.at[i, "order"]) in orders_superfamilies:
                            current_clas = orders_superfamilies.index(str(struc_table.at[i, "order"])) + 2
                        elif str(struc_table.at[i, "class"]) in orders_superfamilies:
                            current_clas = orders_superfamilies.index(str(struc_table.at[i, "class"])) + 2
                        else:
                            current_clas = orders_superfamilies.index("UNCLASSIFIED") + 2
                        keep = input(
                                "Keep the sequence?[enter: current ("+str(current_clas)+"), 1: remove, 2: LTR, 3: COPIA, 4: "
                    "GYPSY, 5: BELPAO, 6: ERV, 7: TRIM, 8: LARD, 9: LINE, 10: SINE, 11: R2, 12: RTE, 13: JOCKEY, 14: L1,"
                    " 15: I, 16: R1, 17: CR1, 18: LOA, 19: L2, 20: PLE, 21: DIRS, 22: NGARO, 23: VIPER, 24: TIR, 25: MITE"
                    ", 26: TC1MARINER,  27: HAT, 28: MUTATOR, 29: MERLIN, 30: TRANSIB, 31: P, 32: PIGGYBAC, 33: PIFHARBINGER,"
                    "  34: CACTA, 35: MULE, 36: CMC, 37: HELITRON, 38: MAVERICK, 39: CRYPTON, 40: UNCLASSIFIED, 41: "
                    "CLASSI, 42: CLASSII] ") or current_clas
                        keep = int(keep)
                    except ValueError:
                        keep = -1
                if keep == 41:
                    seqs_to_module3.append(struc_table.at[i, "Seq_name"])
                elif keep > 1:
                    keep_seqs.append(struc_table.at[i, "Seq_name"])
                    orders.append(keep)
                    kept_seqs_record.append([x for x in SeqIO.parse(te_library, "fasta") if
                                             str(x.id).split("#")[0] == struc_table.at[i, "Seq_name"]][0])

                # increase the number of sequences manually analyzed
                seqs_manu += 1
                print("---------------------------------------------------------------\n")
            else:
                te = [x for x in SeqIO.parse(te_library, "fasta") if
                      x.id.split("#")[0] == struc_table.at[i, "Seq_name"]][0]
                superFamilyLib = ""
                if str(struc_table.at[i, "sFamily"]).upper() in [x for x in dicc_orders.values()]:
                    superFamilyLib = "/" + str(struc_table.at[i, "sFamily"]).upper()
                    orders_incomplete.append(orders_superfamilies.index(str(struc_table.at[i, "sFamily"]).upper()) + 2)
                else:
                    orders_incomplete.append(orders_superfamilies.index(str(struc_table.at[i, "order"]).upper()) + 2)
                te.id = str(struc_table.at[i, "Seq_name"]) + "_inc#" + str(struc_table.at[i, "order"]) + superFamilyLib
                te.description = ""
                non_curated.append(te)
        ele_number += 1

    print("")
    print("Total stats :")
    print("Sequences kept: " + str(len(keep_seqs) + len(non_curated)))
    print("-------------------------------------------")
    print("TEs detected as complete: " + str(len(kept_seqs_record)))
    print("TEs detected as incomplete: " + str(len(non_curated)))
    print("TEs manually analyzed : " + str(seqs_manu))
    print("TEs sent to Unclassified module : " + str(len(seqs_to_module3)))
    print("-------------------------------------------")
    print("")
    return seqs_to_module3, keep_seqs, orders, kept_seqs_record, non_curated, orders_incomplete


def new_module1(plots_dir, ref_tes, gff_files, outputdir, pre, te_aid, automatic, minDomLTR, num_copies, minFLNA,
                verbose, repet, blastn_db, blastx_db, tools_path, ref_profiles, library_path, min_perc_model):
    kept_seqs_record = []
    seqs_to_module3 = []
    non_curated = []
    seqs_module3 = 0
    keep_auto = 0
    ele_number = 0

    # obtain seqIDs of consensi with at least one full length fragment
    seqID_list = [str(te.id).split("#")[0] for te in SeqIO.parse(ref_tes, "fasta")]

    seqs_to_mi = []
    orders_seqs_mi = []
    ref_tes_bee = ref_tes
    if automatic != 'M':
        start_time = time.time()
        # Step1 create feature table
        if not os.path.exists(ref_profiles):
            print("FATAL ERROR: " + ref_profiles + " does not exist. Please check that the file 'Pfam35.0.hmm' is located at db folder and re-run the software")
            sys.exit(0)

        build_class_table_parallel(ref_tes_bee, cores, outputdir+'/classifiedModule/', blastn_db, blastx_db,
                                   ref_profiles, False)
        struc_table = pd.read_csv(outputdir + "/classifiedModule/denovoLibTEs_PC.classif", sep='\t')
        end_time = time.time()
        if verbose:
            print("MESSAGE: TE Feature table was created [" + str(end_time - start_time) + " seconds]")

        # Step2 BLASTn with ref library
        start_time = time.time()
        keep_seqs, orders = run_blast(library_path, ref_tes_bee, cores, 80, 80, 80)
        keep_auto += len(keep_seqs)
        for seq_id in keep_seqs:
            kept_seqs_record.append([x for x in SeqIO.parse(ref_tes_bee, "fasta") if x.id.split("#")[0] == seq_id][0])
        end_time = time.time()
        if verbose:
            print("MESSAGE: Total TEs kept because of hits with library: " + str(len(keep_seqs)) + " [" + str(end_time - start_time) + " seconds]")

        # Step 3 Structural Check
        totalTEs = struc_table.shape[0]
        struc_table.loc[:, "Reason"] = [""]*totalTEs
        for i in range(struc_table.shape[0]):
            status = -1  # 0 = delete, 1 = keep, -1 = Manual Inspection, -2 = Removed
            if int(struc_table.at[i, "length"]) < 100:
                codings = struc_table.at[i, "coding"].replace("coding=(", " ").replace(")", "")
                codigs = codings.split(";")
                reason = "Removed because it is shorter than 100 bp !"
                if struc_table.at[i, "Seq_name"] in keep_seqs:  # removing the element that was kept by homology
                    index_to_rem = keep_seqs.index(struc_table.at[i, "Seq_name"])
                    keep_seqs.remove(struc_table.at[i, "Seq_name"])
                    orders.pop(index_to_rem)
                    kept_seqs_record = [teLong for teLong in kept_seqs_record if teLong.id.split("#")[0] != struc_table.at[i, "Seq_name"]]
                status = -2
                if verbose:
                    print("[" + str(ele_number + 1) + "/" + str(totalTEs) + "] Seq Name: " + str(
                        struc_table.at[i, "Seq_name"]))
                    print("Length: " + str(struc_table.at[i, "length"]))
                    print("Class: " + str(struc_table.at[i, "class"]) + " / " + str(
                        struc_table.at[i, "order"]) + " / " + str(struc_table.at[i, "sFamily"]))
                    print("Copies: FLF = " + str(num_copies[struc_table.at[i, "Seq_name"]][0]) + " / FLC = " + str(
                        num_copies[struc_table.at[i, "Seq_name"]][1]))
                    print("Coding: ")
                    for cod in codigs:
                        print(cod)
                    print(struc_table.at[i, "struct"])
                    print(struc_table.at[i, "other"])
                    print(reason)
                    print("---------------------------------------------------------------\n")
            elif struc_table.at[i, "Seq_name"] not in keep_seqs and struc_table.at[i, "Seq_name"] in seqID_list:
                codings = struc_table.at[i, "coding"].replace("coding=(", " ").replace(")", "")
                codigs = codings.split(";")

                # Structural checks
                if str(struc_table.at[i, "class"]).upper() in ['UNCLASSIFIED', 'NA', 'NAN'] or str(struc_table.at[i, "order"]).upper() in ['UNCLASSIFIED', 'NA', 'NAN']:
                    seqs_to_module3.append(struc_table.at[i, "Seq_name"])
                    status = -3
                    seqs_module3 += 1
                    reason = "Sent to unclassified module due to its order is unclassified or unknown !"
                else:
                    profiles = [cod for cod in codigs if "profiles:" in cod]
                    status, reason = decision_tree_rules(struc_table, profiles, i, keep_seqs, minDomLTR, num_copies, orders, minFLNA, kept_seqs_record, ref_tes_bee, automatic)

                if status == 1:
                    keep_auto += 1
                elif status == -1:
                    all_class = [x.replace("–", "").upper() for x in dicc_orders.values()]
                    seqs_to_mi.append(struc_table.at[i, "Seq_name"])

                    if str(struc_table.at[i, "sFamily"]).upper() != "NAN":
                        class_mod2 = all_class.index(str(struc_table.at[i, "sFamily"]).upper()) + 2
                    else:
                        class_mod2 = all_class.index(str(struc_table.at[i, "order"]).upper()) + 2
                    orders_seqs_mi.append(class_mod2)

                if verbose:
                    print("[" + str(ele_number + 1) + "/" + str(totalTEs) + "] Seq Name: " + str(
                        struc_table.at[i, "Seq_name"]))
                    print("Length: " + str(struc_table.at[i, "length"]))
                    print("Class: " + str(struc_table.at[i, "class"]) + " / " + str(
                        struc_table.at[i, "order"]) + " / " + str(struc_table.at[i, "sFamily"]))
                    print("Copies: FLF = " + str(num_copies[struc_table.at[i, "Seq_name"]][0]) + " / FLC = " + str(
                        num_copies[struc_table.at[i, "Seq_name"]][1]))
                    print("Coding: ")
                    for cod in codigs:
                        print(cod)
                    print(struc_table.at[i, "struct"])
                    print(struc_table.at[i, "other"])
                    print(reason)
                    print("---------------------------------------------------------------\n")
            else:
                reason = "TE Kept because it has homology with our databases !"
                if verbose:
                    codings = struc_table.at[i, "coding"].replace("coding=(", " ").replace(")", "")
                    codigs = codings.split(";")
                    print("[" + str(ele_number + 1) + "/" + str(totalTEs) + "] Seq Name: " + str(
                        struc_table.at[i, "Seq_name"]))
                    print("Length: " + str(struc_table.at[i, "length"]))
                    print("Class: " + str(struc_table.at[i, "class"]) + " / " + str(
                        struc_table.at[i, "order"]) + " / " + str(struc_table.at[i, "sFamily"]))
                    print("Copies: FLF = " + str(num_copies[struc_table.at[i, "Seq_name"]][0]) + " / FLC = " + str(
                        num_copies[struc_table.at[i, "Seq_name"]][1]))
                    print("Coding: ")
                    for cod in codigs:
                        print(cod)
                    print(struc_table.at[i, "struct"])
                    print(struc_table.at[i, "other"])
                    print(reason)
                    print("---------------------------------------------------------------\n")
            ele_number += 1
            struc_table.loc[i, "Reason"] = reason
    else:
        ref_tes_bee = ref_tes
        start_time = time.time()
        build_class_table_parallel(ref_tes_bee, cores, outputdir + '/classifiedModule/', blastn_db, blastx_db,
                                   ref_profiles, False)
        struc_table = pd.read_csv(outputdir + "/classifiedModule/denovoLibTEs_PC.classif", sep='\t')
        end_time = time.time()
        if verbose:
            print("MESSAGE: TE Feature table was created [" + str(end_time - start_time) + " seconds]")
        keep_seqs, orders = [], []

    seqs_to_module3, keep_seqs, orders, kept_seqs_record, non_curated, orders_incomplete = manual_inspection(genome,
    outputdir+"/classifiedModule", ref_tes_bee, seqs_to_mi, seqID_list, struc_table, te_aid, repet, automatic, pre, plots_dir, gff_files,
    min_perc_model, seqs_to_module3, keep_seqs, orders, kept_seqs_record, non_curated, num_copies)

    for index in range(len(kept_seqs_record)):
        # put the order having the superfamily
        classification = dicc_orders[orders[index]]
        if orders[index] >= 3 and orders[index] <= 8:
            classification = "LTR/" + classification
        elif orders[index] >= 11 and orders[index] <= 19:
            classification = "LINE/" + classification
        elif orders[index] >= 21 and orders[index] <= 23:
            classification = "DIRS/" + classification
        elif orders[index] >= 27 and orders[index] <= 36:
            classification = "TIR/" + classification
        elif orders[index] == 40:
            classification = "UNCLASSIFIED"

        # put the class having the order/superfamily
        if orders[index] >= 2 and orders[index] <= 23:
            classification = "CLASSI/" + classification
        elif orders[index] >= 24 and orders[index] <= 39:
            classification = "CLASSII/" + classification

        new_name = keep_seqs[index] + "#" + classification
        kept_seqs_record[index].id = new_name
        kept_seqs_record[index].description = ""

    for index in range(len(non_curated)):
        # put the order having the superfamily
        classification = dicc_orders[orders_incomplete[index]]
        if orders_incomplete[index] >= 3 and orders_incomplete[index] <= 8:
            classification = "LTR/" + classification
        elif orders_incomplete[index] >= 11 and orders_incomplete[index] <= 19:
            classification = "LINE/" + classification
        elif orders_incomplete[index] >= 21 and orders_incomplete[index] <= 23:
            classification = "DIRS/" + classification
        elif orders_incomplete[index] >= 27 and orders_incomplete[index] <= 36:
            classification = "TIR/" + classification
        elif orders_incomplete[index] == 40:
            classification = "UNCLASSIFIED"

        # put the class having the order/superfamily
        if orders_incomplete[index] >= 2 and orders_incomplete[index] <= 23:
            classification = "CLASSI/" + classification
        elif orders_incomplete[index] >= 24 and orders_incomplete[index] <= 39:
            classification = "CLASSII/" + classification

        new_name = str(non_curated[index].id).split("#")[0] + "#" + classification
        non_curated[index].id = new_name
        non_curated[index].description = ""


    seqs_to_module3_record = [te for te in SeqIO.parse(ref_tes_bee, "fasta") if
                              str(te.id).split("#")[0] in seqs_to_module3]
    write_sequences_file(kept_seqs_record, outputdir + "/classifiedModule/kept_seqs_classified_module_curated.fa")
    write_sequences_file(non_curated, outputdir + "/classifiedModule/kept_seqs_classified_module_non_curated.fa")
    write_sequences_file(seqs_to_module3_record, outputdir + "/classifiedModule/input_to_unclassified_module_seqs.fa")
    struc_table.to_csv(outputdir + "/classifiedModule/denovoLibTEs_PC.classif", header=True, sep='\t', index=False)

    delete_files(outputdir + "/classifiedModule/extended_cons.fa")
    delete_files(outputdir + "/classifiedModule/putative_TEs.fa")
    delete_files(outputdir + "/classifiedModule/new_user_lib.fa")


def K2Pdistance(maf):
    """
    Kimura 2-Parameter distance = -0.5 log( (1 - 2p -q) * sqrt( 1 - 2q ) )
    where:
    p = transition frequency
    q = transversion frequency
    """
    start = time.time()
    distance_matrix = np.zeros((maf.shape[0], maf.shape[0]), dtype=float)
    transitions = ["AG", "GA", "CT", "TC"]
    transversions = ["AC", "CA", "AT", "TA", "GC", "CG", "GT", "TG"]
    for i in range(maf.shape[0]-1):
        seq1 = "".join(maf.iloc[i, 1:]).upper()
        for j in range(i + 1, maf.shape[0]):
            seq2 = "".join(maf.iloc[j, 1:]).upper()
            # collect ungapped pairs
            pairs = [x for x in zip(seq1, seq2) if '-' not in x]
            length = len(pairs)
            ts_count = len([(x, y) for (x, y) in pairs if x + y in transitions])
            tv_count = len([(x, y) for (x, y) in pairs if x + y in transversions])
            try:
                p = float(ts_count) / length
                q = float(tv_count) / length
                d = -0.5 * math.log((1 - 2 * p - q) * math.sqrt(1 - 2 * q))
            except:
                d = 1
            distance_matrix[j][i] = distance_matrix[i][j] = d
    end = time.time()
    #print(maf.iloc[0, 0] + ": " + str(end - start))
    return distance_matrix


def read_maf(maf_file):
    """
    This function takes a fasta file path as input and returns a pandas DataFrame
    with the sequence id as the key and the sequence as the value.
    """
    name = os.path.basename(maf_file).split(".")[0]

    sequences = {}
    with open(maf_file) as f:
        sequence = ''
        sequence_id = ''
        for line in f:
            if line.startswith('>'):
                if sequence_id:
                    sequences[sequence_id] = sequence
                    sequence = ''
                sequence_id = line.strip().lstrip('>')
            else:
                sequence += line.strip()
        sequences[sequence_id] = sequence.lower()
    # Read sequences from MAF file.
    data = []
    for sequence_id, sequence in sequences.items():
     row = list(sequence)
     row.insert(0, name)
     data.append(row)
    headers = ['id'] + list(range(len(data[0]) - 1))
    df = pd.DataFrame(data, columns=headers)
    # Return data table with the name of the maf as name of the first column to pass it between functions.
    return df


def seq_clus(maf, min_cluster, max_sequences, cluster_factor, group_outliers, num_subfamilies, max_num_subfamilies):
    if maf is None:
        return None
    elif num_subfamilies > max_num_subfamilies:
        return [maf]
    else:
        name = maf.iloc[0, 0]
        num_seqs = maf.shape[0]
        if maf.shape[0] > min_cluster:  # Don't try to cluster if less than min_cluster sequences
            if maf.shape[0] > max_sequences:  # If more than max_sequences get a sample of them
                maf = maf.sample(n=max_sequences, axis=0)

            d = K2Pdistance(maf)
            # Calculating mp value for DBSCAN based on the clustering factor and the minimum size of cluster
            mp = num_seqs // cluster_factor
            mp = max(mp, min_cluster // 2)

            # Calculating the eps for DBSCAN as the one that produces more "stable" clusters.
            # We define "stable" as clusters that remain for a distance of at least 0.04 (3 eps "steps")
            # The idea is to ignore clusters that are happened just at specific values as outliers.
            points = []
            # Convert NAs to maximum distance.
            d[d == -0.] = 0

            # min eps granularity of 0.2. Could be made a parameter.
            for eps in np.arange(0.02, 1.00, 0.02):
                candidate = len(np.unique(DBSCAN(eps=eps, min_samples=mp, metric='precomputed').fit_predict(d)))
                points.append(candidate)
            # 3 means the cluster must extend at least over 3 eps candidate steps ("stable cluster")
            # Could be made a parameter.
            best_points = []
            for point in set(points):
                if points.count(point) > 3:
                    best_points.append(point)
            nclusters = max(best_points)
            if not group_outliers:
                eps = (points.index(nclusters)) * 0.02 - 0.02
            else:  # If this version is used the 0 clusters are probably just junk and should be
                eps = (50 - (points[::-1].index(nclusters))) * 0.02 - 0.02
            dbscan = DBSCAN(eps=eps, min_samples=mp, metric='precomputed')
            labels = dbscan.fit_predict(d)
            clusters = labels + 1  # add 1 to start clusters at 1 instead of -1 for noise points
            mafs = []
            uniq_clusters = [x for x in set(clusters) if x != 0]
            if len(uniq_clusters) == 1:  # No subfamily split
                clus = maf[clusters == uniq_clusters[0]].reset_index(drop=True)
                clus.iloc[:, 0] = name
                mafs.append(clus)
            else:  # Subfamily divided so we need to add postfixes
                for i in set(clusters):
                    if i != 0:
                        clus = maf[clusters == i].reset_index(drop=True)
                        clus.iloc[:, 0] = name + '_s_' + str(i)
                        mafs.append(clus)
            if (0 in set(clusters)) and (sum(clusters == 0) > mp-1) and (max(clusters) > 1) or (eps == 0.02):
                clus = maf[clusters == 0].reset_index(drop=True)
                clus.iloc[:, 0] = name + '_s_0'
                mafs.append(clus)
        else:
            mafs = [maf]
        return mafs


def process_maf(r, min_plurality, fasta_table_ite, te_class, end_threshold, num_subfamilies):
    if r is None:
        return None
    else:
        name = r.iloc[0, 0]
        r = r.T
        headers = r.iloc[0, :]
        r = r.iloc[1:, :]
        r.columns = headers
        seqs = r.shape[1]
        # Calculate number of empty places and create an index of rows (bases)
        r["res"] = r.isin(["-"]).sum(axis=1)
        r["ID"] = np.arange(len(r))
        # remove fully empty positions (can happen after clustering)
        r = r[r["res"] != seqs]
        r["resA"] = (r == "a").sum(axis=1)
        r["resC"] = (r == "c").sum(axis=1)
        r["resG"] = (r == "g").sum(axis=1)
        r["resT"] = (r == "t").sum(axis=1)
        r["mb"] = r[["resA", "resC", "resG", "resT"]].max(axis=1)  # Majority base
        # "Saturation" is the ratio of bases forming the consensus compared to the total
        # number of sequences.
        r["satur"] = r["mb"] / seqs
        # Remove empty positions and positions with just one base
        r = r[r["res"] + 1 < seqs]
        # Get majority base.
        r["base"] = ""
        r.loc[r["resA"] == r["mb"], "base"] = "a"
        r.loc[r["resC"] == r["mb"], "base"] = "c"
        r.loc[r["resG"] == r["mb"], "base"] = "g"
        r.loc[r["resT"] == r["mb"], "base"] = "t"
        r = r.reset_index(drop=True)
        # Define edges
        # lm and rm are equivalent to saturation, but requiring that the previous (lm) or
        # next (rm) position forms the consensus too.
        r['lm'] = 0
        for i in range(1, r.shape[0]):
            r.at[i, 'lm'] = np.nansum((r.iloc[i, :seqs] == r.at[i, 'base']) & (r.iloc[i - 1, :seqs] == r.at[i - 1, 'base'])) / seqs
        r['rm'] = r['lm'].shift(-1)
        r.loc[r.shape[0]-1, 'rm'] = 0

        m1 = r.loc[r['lm'] > 0.5]['lm'].mean()
        sd1 = r.loc[r['lm'] > 0.5]['lm'].std()
        cutpoint1 = m1 - sd1 * 2
        # Select edges (lm and rm could be unified as the max of both)
        min_r = r[(r['lm'] >= cutpoint1) | (r['rm'] >= cutpoint1)]
        # Correct edges for low number of seqs (need more confidence -> more consecutive bases)
        # Take the edges that match to this rule
        confidence_needed = 20//seqs + 1

        # To calculate where is the 5' edge to cut
        v = min_r.iloc[0:confidence_needed]['rm'] >= cutpoint1
        v = v.tolist()
        stop = False
        while sum(v) < confidence_needed and not stop:
            if False in v:
                min_r = min_r[(confidence_needed - (len(v) - v[::-1].index(False)) + 1):]
                v = min_r[0:confidence_needed]['rm'] >= cutpoint1
                v = v.tolist()
                if len(v) == 0:
                    stop = True
            else:
                stop = True

        # To calculate where is the 3' edge to cut
        v = min_r[(len(min_r) - confidence_needed):(len(min_r))]['lm'] >= cutpoint1
        v = v.tolist()
        while sum(v) < confidence_needed and not stop:
            if False in v:
                min_r = min_r[0:(len(min_r) - confidence_needed + v.index(False))]
                v = min_r[(len(min_r) - confidence_needed):(len(min_r))]['lm'] >= cutpoint1
                v = v.tolist()
                if len(v) == 0:
                    stop = True
            else:
                stop = True

        if not stop:
            # Get the final set of bases that we will use to build the consensus.
            minmin_r = r[(r['ID'] >= min(min_r['ID'])) & (r['ID'] <= max(min_r['ID']))]

            # Build the consensus with a simple plurality model. It can be made as sophisticated as needed.
            minmin_r = minmin_r[minmin_r['satur'] > (min_plurality / 100)].reset_index(drop=True)
            s = ''.join(minmin_r['base'])

            min_l = minmin_r.shape[0]
            end_l_te = r[r['ID'] < minmin_r.loc[0, 'ID']].shape[0] - np.sum(
                (r[r['ID'] < minmin_r.loc[0, 'ID']]['res'] + r[r['ID'] < minmin_r.loc[0, 'ID']]['mb']) == seqs) - (
                                  np.sum(r[r['ID'] < minmin_r.loc[0, 'ID']]['res']) / seqs) > end_threshold
            end_r_te = r[r['ID'] > minmin_r.loc[min_l - 1, 'ID']].shape[0] - np.sum((r[r['ID'] > minmin_r.loc[
                min_l - 1, 'ID']]['res'] + r[r['ID'] > minmin_r.loc[min_l - 1, 'ID']]['mb']) == seqs) - (
                                  np.sum(r[r['ID'] > minmin_r.loc[min_l - 1, 'ID']]['res']) / seqs) > end_threshold

            # add the consensus to the fasta_table
            fasta_table_ite = pd.concat([fasta_table_ite, pd.DataFrame({"seq": name, "cons": s, "cons_size": [len(s)], "class": te_class, "subfamilies": [num_subfamilies], "end_l": end_l_te, "end_r":  end_r_te})], ignore_index=True)
        else:
            # the MSA doesn't have the enough contiguity so, I won't extend it
            print("WARNING: Sequence "+name+" couldn't be processed because the MSA didn't have enough contiguity. I will discard it")
    return fasta_table_ite


def run_extension_by_saturation_parallel(genome, ref_tes, exe_nucl, num_ite, outputdir, minBlastHits, cores, min_perc_model, min_cluster, max_sequences, cluster_factor, group_outliers, min_plurality, end_threshold, max_num_subfamilies):
    create_output_folders(outputdir)

    print('MESSAGE: Starting with BEE step ...')

    # Check if the genome is already formatted to be a BLAST db
    if not os.path.exists(genome + ".n*"):  # Run makeblastdb
        output = subprocess.run(
            ['makeblastdb', '-in', genome, '-dbtype', 'nucl'], stdout=subprocess.PIPE, text=True)

    if os.path.getsize(genome) == 0:
        print("FATAL ERROR: "+genome+" is empty. Please check the file and re-run the software")
        sys.exit(0)
    elif os.path.getsize(ref_tes) == 0:
        print("FATAL ERROR: "+ref_tes+" is empty. Please check the file and re-run the software")
        sys.exit(0)

    ite = 0
    all_finished = False
    tes = [te for te in SeqIO.parse(ref_tes, "fasta")]
    # Fasta table for the sequences will be extended
    fasta_table = pd.DataFrame(columns=["seq", "cons_size", "class", "subfamilies", "end_l", "end_r"])
    for i in range(len(tes)):
        fasta_table = pd.concat([fasta_table, pd.DataFrame(
            {"seq": tes[i].id.split("#")[0], "cons": str(tes[i].seq).lower(),
             "cons_size": [len(tes[i].seq)], "class": str(tes[i].id).split("#")[1], "subfamilies": 0, "end_l": False,
             "end_r": False})], ignore_index=True)
    while ite < num_ite and not all_finished:
        start_time = time.time()
        n = fasta_table.shape[0]
        seqs_per_procs = int(n / cores)
        remain = n % cores
        ini_per_thread = []
        end_per_thread = []
        for p in range(cores):
            if p < remain:
                init = p * (seqs_per_procs + 1)
                end = n if init + seqs_per_procs + 1 > n else init + seqs_per_procs + 1
            else:
                init = p * seqs_per_procs + remain
                end = n if init + seqs_per_procs > n else init + seqs_per_procs
            ini_per_thread.append(init)
            end_per_thread.append(end)

        # Run in parallel the checking
        pool = multiprocessing.Pool(processes=cores)
        localresults = [pool.apply_async(extension_by_saturation,
                                         args=[genome, fasta_table.loc[ini_per_thread[x]:end_per_thread[x]-1, :], exe_nucl,
                                               outputdir, min_perc_model, min_cluster, max_sequences,
                                               cluster_factor, group_outliers, min_plurality, end_threshold,
                                               max_num_subfamilies, minBlastHits]) for x in range(cores)]

        localChecks = [p.get() for p in localresults]
        final_seqs = pd.DataFrame(columns=["seq", "cons_size", "class", "subfamilies", "end_l", "end_r"])
        for i in range(len(localChecks)):
            final_seqs = pd.concat([final_seqs, localChecks[i]], ignore_index=True)
        pool.close()

        if verbose:
            end_time = time.time()
            print("MESSAGE: iteration  " + str(ite + 1) + "/" + str(num_ite) + " done ["+str(end_time-start_time)+" seconds]. Generated " + str(final_seqs.shape[0]) + " sequences.")
        ite += 1
        fasta_table = final_seqs
        if fasta_table[~((fasta_table['end_l'] == True) & (fasta_table['end_r'] == True))].shape[0] == 0:
            all_finished = True

    merged_results = []
    for index in range(fasta_table.shape[0]):
        final_seq = SeqRecord(Seq(fasta_table.loc[index, "cons"]), id=fasta_table.loc[index, "seq"]+"#"+fasta_table.loc[index, "class"])
        merged_results.append(final_seq)

    # refine TE extension, removing over-extended regions
    n = len(merged_results)
    seqs_per_procs = int(n / cores)
    remain = n % cores
    ini_per_thread = []
    end_per_thread = []
    for p in range(cores):
        if p < remain:
            init = p * (seqs_per_procs + 1)
            end = n if init + seqs_per_procs + 1 > n else init + seqs_per_procs + 1
        else:
            init = p * seqs_per_procs + remain
            end = n if init + seqs_per_procs > n else init + seqs_per_procs
        ini_per_thread.append(init)
        end_per_thread.append(end)

    # Run in parallel the checking
    pool = multiprocessing.Pool(processes=cores)
    localresults = [pool.apply_async(refine_extension, args=[merged_results[ini_per_thread[x]:end_per_thread[x]],
                                                             genome, outputdir, x]) for x in range(cores)]

    localChecks = [p.get() for p in localresults]
    final_results = []
    for i in range(len(localChecks)):
        final_results.extend(localChecks[i])
    pool.close()

    write_sequences_file(final_results, outputdir + "/extended_cons.fa")


def extension_by_saturation(genome, fasta_table, exe_nucl, outputdir, min_perc_model, min_cluster, max_sequences, cluster_factor, group_outliers, min_plurality, end_threshold, max_num_subfamilies, minBlastHits):
    fasta_table_ite = pd.DataFrame(columns=["seq", "cons_size", "class", "end_l", "end_r"])
    if fasta_table.shape[0] > 0:
        fasta_table = fasta_table.reset_index()
        for index in range(fasta_table.shape[0]):
            if not fasta_table.loc[index, "end_l"] or not fasta_table.loc[index, "end_r"]:
                seq_name = fasta_table.loc[index, "seq"]
                te_class = fasta_table.loc[index, "class"]
                te = SeqRecord(Seq(fasta_table.loc[index, "cons"]), id=fasta_table.loc[index, "seq"])
                write_sequences_file([te], outputdir + "/" + str(seq_name) + ".consensus.fasta")

                # First step: BLASTn
                # parameters from Anna Protasio github
                output = subprocess.run(
                    ['blastn', '-query', outputdir + "/" + str(seq_name) + ".consensus.fasta", '-db',
                     genome, '-out', outputdir + "/" + str(seq_name) + ".blast", '-num_threads', "1",
                     "-outfmt", "6 sseqid sstart send length pident bitscore", "-evalue", "1e-20"], stdout=subprocess.PIPE, text=True)
                delete_files(outputdir + "/" + str(seq_name) + ".consensus.fasta")

                # Second step EXTRACT and EXTEND
                blastresult = pd.read_table(outputdir + "/" + str(seq_name) + ".blast", sep='\t',
                                        names=['sseqid', 'sstart', 'send', 'length', 'pident', 'bitscore'], nrows=200)
                blastresult = blastresult.sort_values(by=['bitscore', 'pident', 'length'],
                                                      ascending=[False, False, False])
                delete_files(outputdir + "/" + str(seq_name) + ".blast")
                result_file = open(outputdir + "/" + str(seq_name) + ".copies.fa", "w")
                hit = 0
                copies_to_use = 0
                while hit < blastresult.shape[0] and hit < 200:
                    # Adding 300 on completed ends to allow the method to find the edges again.
                    if fasta_table.loc[index, "end_l"]:
                        exe_nucl_5prime = 300
                    else:
                        exe_nucl_5prime = exe_nucl
                    if fasta_table.loc[index, "end_r"]:
                        exe_nucl_3prime = 300
                    else:
                        exe_nucl_3prime = exe_nucl
                    reverse = False
                    subject_seq = blastresult.at[hit, 'sseqid']
                    ini_hit = int(blastresult.at[hit, 'sstart'])
                    end_hit = int(blastresult.at[hit, 'send'])
                    if float(blastresult.at[hit, 'length']) > float(fasta_table.loc[index, "cons_size"]) * min_perc_model:
                        scaff_seq = [x.seq for x in SeqIO.parse(genome, 'fasta') if
                                     str(x.id).split(" ")[0] == subject_seq]
                        if len(scaff_seq) > 0:
                            scaff_seq = str(scaff_seq[0])
                            # for reversed hits
                            if end_hit < ini_hit:
                                tem = ini_hit
                                ini_hit = end_hit
                                end_hit = tem
                                tem = exe_nucl_5prime
                                exe_nucl_5prime = exe_nucl_3prime
                                exe_nucl_3prime = tem
                                reverse = True

                            if ini_hit - exe_nucl_5prime < 0:
                                ini_hit = 0
                            else:
                                ini_hit -= exe_nucl_5prime
                            if end_hit + exe_nucl_3prime > len(scaff_seq):
                                end_hit = len(scaff_seq)
                            else:
                                end_hit += exe_nucl_3prime

                            if reverse is False:
                                result_file.write(
                                    ">copy_" + str(subject_seq) + "_" + str(ini_hit) + "_" + str(
                                        end_hit) + "_" + str(hit) + "\n" + scaff_seq[ini_hit:end_hit].lower() + "\n")
                            else:
                                result_file.write(
                                    ">copy_" + str(subject_seq) + "_" + str(ini_hit) + "_" + str(end_hit) + "_" + str(hit)
                                    + "_reversed\n" + str(Seq(scaff_seq[ini_hit:end_hit].lower()).reverse_complement()) + "\n")
                        else:
                            print(
                                'WARNING: A subject id (sseqid in blast output file) was not found in the genome, please check:')
                            print(subject_seq)
                        copies_to_use += 1
                    hit += 1
                result_file.close()

                if copies_to_use >= minBlastHits:  # if there is at least minBlastHits of the consensus and the genome
                    # parameters from Anna Protasio's gitHub
                    output = subprocess.run(['mafft', '--quiet', '--thread', '1', '--reorder',
                                             outputdir + "/" + str(seq_name) + '.copies.fa'],
                                            stdout=subprocess.PIPE, stderr=subprocess.DEVNULL, text=True)

                    mafft_comm_output = str(output.stdout)

                    if len(mafft_comm_output) > 0:
                        mafft_output = open(outputdir + "/" + str(seq_name) + ".copies.mafft", "w")
                        mafft_output.write(mafft_comm_output)
                        mafft_output.close()
                        del mafft_comm_output   # clean the variable

                        mafft_df = read_maf(outputdir + "/" + str(seq_name) + ".copies.mafft")
                        delete_files(outputdir + "/" + str(seq_name) + ".copies.mafft")
                        maffts_df_list = seq_clus(mafft_df, min_cluster, max_sequences, cluster_factor, group_outliers, fasta_table.loc[index, "subfamilies"], max_num_subfamilies)
                        if len(maffts_df_list) > 1:
                            fasta_table.loc[index, "subfamilies"] += len(maffts_df_list)  # increase the number of created subfamilies

                        for mafft in maffts_df_list:
                            fasta_table_ite = process_maf(mafft, min_plurality, fasta_table_ite, te_class, end_threshold, fasta_table.loc[index, "subfamilies"])
                else:
                    # Model doesn't have enough copies to be extended
                    fasta_table_ite = pd.concat([fasta_table_ite, pd.DataFrame({"seq": fasta_table.loc[index, "seq"], "cons": fasta_table.loc[index, "cons"], "cons_size": [fasta_table.loc[index, "cons_size"]], "class": fasta_table.loc[index, "class"], "subfamilies": fasta_table.loc[index, "subfamilies"], "end_l": True, "end_r": True})], axis=0, ignore_index=True)
                delete_files(outputdir + "/" + str(seq_name) + '.copies.fa')
            else:
                # Model is complete
                fasta_table_ite = pd.concat([fasta_table_ite, pd.DataFrame({"seq": fasta_table.loc[index, "seq"], "cons": fasta_table.loc[index, "cons"], "cons_size": [fasta_table.loc[index, "cons_size"]], "class": fasta_table.loc[index, "class"], "subfamilies": fasta_table.loc[index, "subfamilies"], "end_l": True, "end_r": True})], axis=0, ignore_index=True)

    return fasta_table_ite


def refine_extension(tes_to_refine, genome, outputdir, id_thread):
    te_refined = []
    for te in tes_to_refine:
        # set values for query, db, and evalue variables
        write_sequences_file([te], outputdir+"/torefine.fa_"+str(id_thread))
        query = outputdir+"/torefine.fa_"+str(id_thread)
        cons_len = len(str(te.seq))

        # run blastn command and read output into pandas dataframe
        command = f"blastn -query {query} -db {genome} -evalue 10e-8 -num_threads 1 -outfmt '6 qstart qend'"
        output = subprocess.check_output(command, shell=True).decode()

        if len(output) > 0:
            blast = pd.read_csv(io.StringIO(output), sep='\t', header=None)
            # create matrix of zeros with dimensions based on length of blast dataframe and cons_len variable
            coverage = np.zeros((len(blast), cons_len), dtype=np.bool_)

            # iterate over rows of blast dataframe and set corresponding values in coverage matrix
            for i, row in blast.iterrows():
                start = int(row[0]) - 1
                end = int(row[1]) - 1
                if start <= end:
                    coverage[i][start:end + 1] = 1
                else:
                    coverage[i][end:start + 1] = 1

            # calculate column sums of coverage matrix and write to file
            coverage_sum = np.sum(coverage, axis=0).T

            # remove the over-extended regions based on coverage
            # 5' end
            index = 0
            while coverage_sum[index] == 0 and index < coverage_sum.shape[0]:
                index += 1
            new_start = index

            # 3' end
            index = coverage_sum.shape[0] - 1
            while coverage_sum[index] == 0 and index >= 0:
                index -= 1
            new_end = index

            te.seq = Seq(str(te.seq)[new_start:new_end])
            te.description = ""

        delete_files(query)
        te_refined.append(te)
    return te_refined


def filter_bad_candidates(new_ref_tes, perc_ssr, outputdir, tools_path, busco_library, RNAs_library, cores):
    if os.path.getsize(new_ref_tes) == 0:
        print("FATAL ERROR: " + new_ref_tes + " is empty. Please check the file and re-run the software")
        sys.exit(0)
    elif os.path.getsize(busco_library) == 0:
        print("FATAL ERROR: "+busco_library+" is empty. Please check the file and re-run the software")
        sys.exit(0)

    deleted_seqs = []
    kept_seqs = []

    current_path = os.getcwd()
    os.chdir(outputdir)
    try:
        output = subprocess.run(
            [tools_path + '/trf409.linux64', new_ref_tes, '2', '3', '5', '80', '10', '20', '15', '-h', '-d'],
            stdout=subprocess.PIPE, stderr=subprocess.PIPE, text=True)
    except Exception as exp:
        print("FATAL ERROR: I couldn't execute properly the TRF program. Please check the error: "+exp.args)

    os.chdir(current_path)
    # Search for single repeats in TEs
    dicc_sr_pos = {}
    with open(outputdir + "/"+os.path.basename(new_ref_tes)+".2.3.5.80.10.20.15.dat", 'r') as inFile:
        nbInSeq = 0
        for line in inFile:
            row = line.split(" ")
            if len(row) > 1 and "Sequence:" in row[0]:
                nbInSeq += 1
                seqName = row[1].replace('\n', '')
            if len(row) >= 14 and "Sequence:" not in row[0]:
                start = row[0]
                end = row[1]
                if seqName in dicc_sr_pos.keys():
                    dicc_sr_pos[seqName].append([int(start), int(end)])
                else:
                    dicc_sr_pos[seqName] = [[int(start), int(end)]]

    # collapse all overlapping satellite matches
    for sat in dicc_sr_pos.keys():
        pos_list = dicc_sr_pos[sat]
        new_list = [pos_list[0]]
        for i in range(1, len(pos_list)):
            # if there is an overlap
            overlap = False
            for j in range(len(new_list)):
                if (new_list[j][0] <= pos_list[i][0] <= new_list[j][1] or new_list[j][0] <= pos_list[i][1] <=
                    new_list[j][1]) \
                        or (pos_list[i][0] <= new_list[j][0] <= pos_list[i][1] or pos_list[i][0] <= new_list[j][1] <=
                            pos_list[i][1]):
                    new_start = new_list[j][0] if new_list[j][0] < pos_list[i][0] else pos_list[i][0]
                    new_end = new_list[j][1] if new_list[j][1] > pos_list[i][1] else pos_list[i][1]
                    new_list[j] = [new_start, new_end]
                    overlap = True
            if not overlap:
                new_list.append([pos_list[i][0], pos_list[i][1]])
        dicc_sr_pos[sat] = new_list

    # count how many bases are satellites
    dicc_sr = {}
    for sat in dicc_sr_pos.keys():
        sum_bp = 0
        pos_list = dicc_sr_pos[sat]
        for i in range(len(pos_list)):
            sum_bp += pos_list[i][1] - pos_list[i][0] + 1
        dicc_sr[sat] = sum_bp

    # Search for matches with References/BUSCO genes
    if not os.path.exists(busco_library+".h3m"):
        output = subprocess.run(['hmmpress', busco_library],
                                stdout=subprocess.DEVNULL, stderr=subprocess.DEVNULL, text=True)

    output = subprocess.run(
        ['hmmscan', '--tblout', outputdir + "tes_vs_genes.hmm", '-E', '10', '--noali', '--cpu',
         str(cores), busco_library, new_ref_tes],
        stdout=subprocess.DEVNULL, stderr=subprocess.DEVNULL, text=True)

    command = 'grep -v "^#" ' + outputdir + 'tes_vs_genes.hmm | awk \'{print $1"\t"$2"\t"$3"\t"$4"\t"$5"\t"$6"\t"$7"\t"$8"\t"$9"\t"$10}\' > ' + outputdir + "tes_vs_genes.hmm_formatted"
    process = subprocess.Popen(command, shell=True)
    process.communicate()
    hmm_results = pd.read_table(outputdir + "tes_vs_genes.hmm_formatted", sep='\t',
                                names=['target', 'acc', 'query', 'acc_query', 'evalue', 'score', 'A', 'B', 'C',
                                       'D'])

    tes_with_matches = []
    for x in range(hmm_results.shape[0]):
        if hmm_results.loc[x, "query"] not in tes_with_matches:
            tes_with_matches.append(hmm_results.loc[x, "query"])

    # Search for matches with tRNAs or rRNAs
    if not os.path.exists(RNAs_library + ".h3m"):
        output = subprocess.run(['hmmpress', RNAs_library],
                                stdout=subprocess.PIPE, stderr=subprocess.PIPE, text=True)

    output = subprocess.run(
        ['hmmscan', '--tblout', outputdir + "/tes_vs_rnas.hmm", '-E', '10', '--noali', '--cpu',
         str(cores), RNAs_library, new_ref_tes],
        stdout=subprocess.PIPE, stderr=subprocess.PIPE, text=True)

    command = 'grep -v "^#" ' + outputdir + '/tes_vs_rnas.hmm | awk \'{print $1"\t"$2"\t"$3"\t"$4"\t"$5"\t"$6"\t"$7"\t"$8"\t"$9"\t"$10}\' > ' + outputdir + '/tes_vs_rnas.hmm_formatted'
    process = subprocess.Popen(command, shell=True)
    process.communicate()
    hmm_results = pd.read_table(outputdir + '/tes_vs_rnas.hmm_formatted', sep='\t',
                                names=['target', 'acc', 'query', 'acc_query', 'evalue', 'score', 'A', 'B', 'C', 'D'])

    tes_with_rnas = []
    for x in range(hmm_results.shape[0]):
        if hmm_results.loc[x, "query"] not in tes_with_rnas:
            tes_with_rnas.append(hmm_results.loc[x, "query"])

    # Remove those TEs with SSR > given threshold or those that match with genes
    for te in SeqIO.parse(new_ref_tes, "fasta"):
        if te.id not in tes_with_matches and te.id not in tes_with_rnas:  # It hasn't a match with Reference/BUSCO genes neither with RNAs
            if te.id in dicc_sr.keys():
                te_len = len(str(te.seq))
                lenSR = dicc_sr[te.id]
                if ((lenSR * 100) / te_len) < perc_ssr:  # It hasn't less than the given threshold of SSR in its sequence
                    kept_seqs.append(te)
                else:
                    deleted_seqs.append(te)
            else:  # It hasn't any SSR
                kept_seqs.append(te)
        else:
            deleted_seqs.append(te)

    write_sequences_file(kept_seqs, outputdir + "/putative_TEs.fa")
    delete_files(outputdir + "/" + os.path.basename(new_ref_tes) + ".2.3.5.80.10.20.15.dat")
    delete_files(outputdir + "/tes_vs_genes.hmm")
    delete_files(outputdir + "/tes_vs_genes.hmm_formatted")
    delete_files(outputdir + "/tes_vs_rnas.hmm")
    delete_files(outputdir + "/tes_vs_rnas.hmm_formatted")
    delete_files(busco_library + ".h3f")
    delete_files(busco_library + ".h3i")
    delete_files(busco_library + ".h3m")
    delete_files(busco_library + ".h3p")

    return outputdir + "/putative_TEs.fa", deleted_seqs


def module3(ref_tes, library_path, cores, outputdir, perc_ident, perc_cover, min_len_unc, internal_library):
    # BLASTn with already curated TEs
    start_time = time.time()
    keep_seqs_records = []
    keep_seqs, orders = run_blast(library_path, ref_tes, cores, perc_ident, perc_cover, min_len_unc)

    # BLASTn with internal reference TE database
    keep_seqs_internal, orders_internal = run_blast(internal_library, ref_tes, cores, perc_ident, perc_cover, min_len_unc)

    for te_index in range(len(keep_seqs_internal)):
        if keep_seqs_internal[te_index] not in keep_seqs:
            keep_seqs.append(keep_seqs_internal[te_index])
            orders.append(orders_internal[te_index])

    for i in range(len(keep_seqs)):
        te_selected = [x for x in SeqIO.parse(ref_tes, "fasta") if x.id.split("#")[0] == keep_seqs[i]]
        te_selected[0].id = te_selected[0].id.split("#")[0]
        te_selected[0].description = ""
        keep_seqs_records.append(te_selected[0])

    end_time = time.time()
    print("MESSAGE: BLASTn successfully run [" + str(end_time - start_time) + " seconds]")

    # Structural checks
    start_time = time.time()
    struc_table = pd.read_csv(outputdir + "/unclassifiedModule/denovoLibTEs_PC.classif", sep='\t')
    num_infered_tes = 0
    num_structural_tes = 0
    for i in range(struc_table.shape[0]):
        if struc_table.at[i, "Seq_name"] not in keep_seqs:
            codings = struc_table.at[i, "coding"].replace("coding=(", " ").replace(")", "")
            codigs = codings.split(";")
            profiles = [cod for cod in codigs if "profiles:" in cod]
            inferred, new_class = inferring_domains(profiles)
            if inferred:
                te_selected = [x for x in SeqIO.parse(ref_tes, "fasta") if x.id.split("#")[0] == struc_table.at[i, "Seq_name"]]
                te_selected[0].id = te_selected[0].id.split("#")[0]
                te_selected[0].description = ""
                keep_seqs_records.append(te_selected[0])
                orders.append(new_class)
                num_infered_tes += 1
            else:
                if "termLTR: " in struc_table.at[i, "struct"]:
                    te_selected = [x for x in SeqIO.parse(ref_tes, "fasta") if x.id.split("#")[0] == struc_table.at[i, "Seq_name"]]
                    te_selected[0].id = te_selected[0].id.split("#")[0] + "_unconfirmed"
                    te_selected[0].description = ""
                    keep_seqs_records.append(te_selected[0])
                    orders.append(orders_superfamilies.index("LTR") + 2)
                    num_structural_tes += 1
                elif "termTIR: " in struc_table.at[i, "struct"]:
                    te_selected = [x for x in SeqIO.parse(ref_tes, "fasta") if x.id.split("#")[0] == struc_table.at[i, "Seq_name"]]
                    te_selected[0].id = te_selected[0].id.split("#")[0] + "_unconfirmed"
                    te_selected[0].description = ""
                    keep_seqs_records.append(te_selected[0])
                    orders.append(orders_superfamilies.index("TIR") + 2)
                    num_structural_tes += 1

    for index in range(len(keep_seqs_records)):
        # put the order having the superfamily
        classification = dicc_orders[orders[index]]
        if orders[index] >= 3 and orders[index] <= 8:
            classification = "LTR/" + classification
        elif orders[index] >= 11 and orders[index] <= 19:
            classification = "LINE/" + classification
        elif orders[index] >= 21 and orders[index] <= 23:
            classification = "DIRS/" + classification
        elif orders[index] >= 26 and orders[index] <= 36:
            classification = "TIR/" + classification
        elif orders[index] == 40:
            classification = "UNCLASSIFIED"

        # put the class having the order/superfamily
        if orders[index] >= 2 and orders[index] <= 23:
            classification = "CLASSI/" + classification
        elif orders[index] >= 24 and orders[index] <= 39:
            classification = "CLASSII/" + classification

        new_name = keep_seqs_records[index].id + "#" + classification
        keep_seqs_records[index].id = new_name
        keep_seqs_records[index].description = ""

    end_time = time.time()
    print("MESSAGE: Order inferring from structural features done [" + str(end_time - start_time) + " seconds]")

    if len(keep_seqs_records) > 0:
        write_sequences_file(keep_seqs_records, outputdir + "/unclassifiedModule/kept_seqs_unclassified_module.fa")
        print("")
        print("Total stats :")
        print("Unclassified elements recovered: " + str(len(keep_seqs_records)))
        print("-------------------------------------------")
        print("TEs recovered by homology: " + str(len(keep_seqs)))
        print("TEs recovered by inference of coding domains: " + str(num_infered_tes))
        print("TEs recovered by structural features: " + str(num_structural_tes))
        print("-------------------------------------------")
        print("")
    else:
        print("WARNING: unclassified module couldn't find any TE !")


def run_te_aid_parallel(te_aid_path, genome, ref_tes, outputdir, cores, min_perc_model):
    if not os.path.exists(outputdir + "/te_aid"):

        if not os.path.exists(genome + ".nhr"):
            output = subprocess.run(
                ['makeblastdb', '-in', genome, '-dbtype', 'nucl'], stdout=subprocess.PIPE, text=True)

        tes = [te for te in SeqIO.parse(ref_tes, "fasta")]
        n = len(tes)
        seqs_per_procs = int(n / cores)
        remain = n % cores
        ini_per_thread = []
        end_per_thread = []
        for p in range(cores):
            if p < remain:
                init = p * (seqs_per_procs + 1)
                end = n if init + seqs_per_procs + 1 > n else init + seqs_per_procs + 1
            else:
                init = p * seqs_per_procs + remain
                end = n if init + seqs_per_procs > n else init + seqs_per_procs
            ini_per_thread.append(init)
            end_per_thread.append(end)

        # Run in parallel the checking
        pool = multiprocessing.Pool(processes=cores)
        localresults = [pool.apply_async(run_te_aid,
                                         args=[te_aid_path, genome, outputdir + "/te_aid_" + str(x),
                                               tes[ini_per_thread[x]:end_per_thread[x]], min_perc_model]) for x in
                        range(cores)]

        create_output_folders(outputdir + "/te_aid")
        create_output_folders(outputdir + "/MSA_plots/")
        create_output_folders(outputdir + "/MSA_seeds/")

        localChecks = [p.get() for p in localresults]
        for i in range(len(localChecks)):
            if localChecks[i] == 0:
                print("FATAL ERROR: TE+Aid in Parallel didn't execute well. Please re run.")
                sys.exit(0)
            else:
                pattern = "*.pdf"
                files = glob.glob(outputdir + "/te_aid_" + str(i) + "/" + pattern)

                try:
                    for file in files:
                        # this is getting errors in CentOS
                        """# extract file name form file path
                        file_name = os.path.basename(file)
                        pages = convert_from_path(file)
                        # Saving pages in jpeg format
                        for page in pages:
                            page.save(outputdir + "/te_aid/" + file_name + '.jpeg', 'JPEG', quality=85)"""
                        # extract file name form file path
                        file_name = os.path.basename(file)
                        shutil.move(file, outputdir + "/te_aid/" + file_name)
                except:
                    print("WARNING: The file "+file+" did not have any pages. Skipping TE+Aid plot ...")
                pattern = "*.copies.cialign_output.png"
                files = glob.glob(outputdir + "/te_aid_" + str(i) + "/" + pattern)
                for file in files:
                    file_name = os.path.basename(file)
                    shutil.move(file, outputdir + "/MSA_plots/" + file_name)

                pattern = "*.copies.mafft"
                files = glob.glob(outputdir + "/te_aid_" + str(i) + "/" + pattern)
                for file in files:
                    file_name = os.path.basename(file)
                    shutil.move(file, outputdir + "/MSA_seeds/" + file_name)

                shutil.rmtree(outputdir + "/te_aid_" + str(i))
        pool.close()

    else:
        print("TE+aid already run!")


def run_te_aid(te_aid_path, genome, outputdir, tes, min_perc_model):
    status = -1
    create_output_folders(outputdir)
    for sequence in tes:
        seq_name = str(sequence.id).split("#")[0]
        write_sequences_file([sequence], outputdir + "/" + str(seq_name) + ".fa")

        try:
            output = subprocess.run(
                [te_aid_path + '/TE-Aid', '-q', outputdir + "/" + str(seq_name) + ".fa", '-g', genome, '-o', outputdir],
                 stdout=subprocess.PIPE, stderr=subprocess.PIPE, text=True)

            status = 1
        except:
            status = 0

        # Create the MSA seeds
        # First step: BLASTn
        # parameters from Anna Protasio github
        output = subprocess.run(
            ['blastn', '-query', outputdir + "/" + str(seq_name) + ".fa", '-db',
             genome, '-out', outputdir + "/" + str(seq_name) + ".blast", '-num_threads', "1",
             "-outfmt", "6 sseqid sstart send length", "-evalue", "1e-20"], stdout=subprocess.PIPE, text=True)
        delete_files(outputdir + "/" + str(seq_name) + ".consensus.fasta")

        # Second step EXTRACT and EXTEND
        blastresult = pd.read_table(outputdir + "/" + str(seq_name) + ".blast", sep='\t',
                                    names=['sseqid', 'sstart', 'send', 'length'], nrows=200)

        result_file = open(outputdir + "/" + str(seq_name) + ".copies.fa", "w")
        hit = 0
        copies_to_use = 0
        while hit < blastresult.shape[0] and hit < 200:
            reverse = False
            subject_seq = blastresult.at[hit, 'sseqid']
            ini_hit = int(blastresult.at[hit, 'sstart'])
            end_hit = int(blastresult.at[hit, 'send'])
            if float(blastresult.at[hit, 'length']) > float(len(sequence.seq)) * min_perc_model:
                scaff_seq = [x.seq for x in SeqIO.parse(genome, 'fasta') if
                             str(x.id).split(" ")[0] == subject_seq]
                if len(scaff_seq) > 0:
                    scaff_seq = str(scaff_seq[0])
                    # for reversed hits
                    if end_hit < ini_hit:
                        tem = ini_hit
                        ini_hit = end_hit
                        end_hit = tem
                        reverse = True

                    if reverse is False:
                        result_file.write(
                            ">copy_" + str(subject_seq) + "_" + str(ini_hit) + "_" + str(
                                end_hit) + "_" + str(hit) + "\n" + scaff_seq[ini_hit:end_hit].lower() + "\n")
                    else:
                        result_file.write(
                            ">copy_" + str(subject_seq) + "_" + str(ini_hit) + "_" + str(end_hit) + "_" + str(hit)
                            + "_reversed\n" + str(Seq(scaff_seq[ini_hit:end_hit].lower()).reverse_complement()) + "\n")
                else:
                    print(
                        'WARNING: A subject id (sseqid in blast output file) was not found in the genome, please check:')
                    print(subject_seq)
                copies_to_use += 1
            hit += 1
        result_file.close()

        if copies_to_use > 1:  # if there is at least two hits of the consensus and the genome
            # parameters from Anna Protasio's gitHub
            output = subprocess.run(['mafft', '--quiet', '--thread', '1', '--reorder',
                                     outputdir + "/" + str(seq_name) + '.copies.fa'],
                                    stdout=subprocess.PIPE, stderr=subprocess.DEVNULL, text=True)

            mafft_comm_output = str(output.stdout)

            if len(mafft_comm_output) > 0:
                mafft_output = open(outputdir + "/" + str(seq_name) + ".copies.mafft", "w")
                mafft_output.write(mafft_comm_output)
                mafft_output.close()

                # plot the MSA seeds
                if os.path.exists(outputdir + "/" + str(seq_name) + ".copies.mafft"):
                    output = subprocess.run(
                        ['CIAlign', '--infile', outputdir + "/" + str(seq_name) + ".copies.mafft",
                         '--out',
                         outputdir + "/" + str(seq_name) + ".copies.cialign", '--plot_output',
                         '--silent'])
                    delete_files(outputdir + "/" + str(seq_name) + ".copies.cialign_removed.txt")
                    delete_files(outputdir + "/" + str(seq_name) + ".copies.cialign_cleaned.fasta")
                    delete_files(outputdir + "/" + str(seq_name) + ".copies.cialign_log.txt")

        delete_files(outputdir + "/" + str(seq_name) + ".fa")
    return status


def run_blast(library_path, ref_tes, cores, perc_identity, perc_cov, min_len):
    keep_seqs = []
    orders = []

    if not os.path.exists(library_path + ".nhr"):
        output = subprocess.run(
            ['makeblastdb', '-in', library_path, '-dbtype', 'nucl'], stdout=subprocess.DEVNULL, stderr=subprocess.DEVNULL, text=True)

    if not os.path.exists(ref_tes + ".blast"):
        output = subprocess.run(
            ['blastn', '-query', ref_tes, '-db', library_path, '-out', ref_tes + ".blast", '-num_threads', str(cores),
             "-outfmt", "6 qseqid sseqid length", "-qcov_hsp_perc", str(perc_cov), "-perc_identity", str(perc_identity), "-max_hsps", "1"],
            stdout=subprocess.PIPE, text=True)
    else:
        print("WARNING: Blast output already exists, skipping BLASTn....")

    # BLAST tabular optimized format: qseqid sseqid length
    blastresult = open(ref_tes + ".blast", "r").readlines()

    for hit in blastresult:
        if hit.split("\t")[0].split("#")[0] not in keep_seqs and int(hit.split("\t")[2]) >= min_len:
            blasted_order = hit.replace("?", "").split("\t")[1].split("#")[1]
            if "/" in blasted_order:
                order_given = blasted_order.split("/")[0].upper()
                superfamily = blasted_order.split("/")[1].upper()
            else:
                order_given = blasted_order.upper()
                superfamily = "NA"

            if superfamily.replace('-', '').upper() in orders_superfamilies:
                # superfamily found !!!
                orders.append(orders_superfamilies.index(superfamily.replace('-', '').upper()) + 2)
                keep_seqs.append(hit.split("\t")[0].split("#")[0])
            elif order_given.replace('-', '').upper() in orders_superfamilies:
                # Well, superfamily didn't find, but I found the order
                orders.append(orders_superfamilies.index(order_given.replace('-', '').upper()) + 2)
                keep_seqs.append(hit.split("\t")[0].split("#")[0])
            else:
                print("WARNING: Neither order and superfamily found: " + blasted_order)
                orders.append(31)
                keep_seqs.append(hit.split("\t")[0].split("#")[0])

    delete_files(ref_tes + ".blast")
    return keep_seqs, orders


def run_cdhit(ref_tes, outputdir, cores, identity, coverage):
    create_output_folders(outputdir)

    if not os.path.exists(outputdir + "/non_redundant_lib.fa"):
        output = subprocess.run(
            ['cd-hit-est', '-i', ref_tes, '-o', outputdir + '/non_redundant_lib.fa', '-c', str(identity), '-aS', str(coverage), '-G', '0', '-g' , '1', '-b', '500', '-T', str(cores), '-M', '0', '-d', '0'],
            stdout=subprocess.PIPE, text=True)
    else:
        print("WARNING: cd-hit-est output already exists, skipping cd-hit-est....")

    return outputdir + "/non_redundant_lib.fa"


def run_meshclust(ref_tes, outputdir, cores, identity, coverage):
    create_output_folders(outputdir)

    if not os.path.exists(outputdir + "/non_redundant_lib.fa"):
        output = subprocess.run(
            ['meshclust', '-d', ref_tes, '-o', outputdir + '/non_redundant_lib.fa.clust', '-t', str(identity), '-c', str(cores), '-a', 'no'],
            stdout=subprocess.DEVNULL, text=True)
    else:
        print("WARNING: meshclust output already exists, skipping cd-hit-est....")

    fileopen = open(outputdir + '/non_redundant_lib.fa.clust', "r").readlines()
    members = []
    for line in fileopen:
        if line != "\n":  # it's a member of the cluster
            if line.replace("\n", "").split("\t")[3] == "C":
                member_name = line.split("\t")[1].split(" ")[0].replace(">", "")
                members.append(member_name)

    sequences = [te for te in SeqIO.parse(ref_tes, "fasta") if te.id in members]
    write_sequences_file(sequences, outputdir + '/non_redundant_lib.fa')

    return outputdir + "/non_redundant_lib.fa"


if __name__ == '__main__':

    Installation_path = os.path.dirname(os.path.realpath(__file__))
    ####################################################################################################################
    # Needed by Classified module
    ####################################################################################################################
    tools_path = Installation_path + "/tools/"
    library_path = Installation_path + "/db/allDatabases.clustered_rename.fa"
    ref_profiles = Installation_path + "/db/Pfam35.0.hmm"
    blastn_db = Installation_path + "/db/Dfam_3.7_curatedonly.fa"
    blastx_db = Installation_path + "/db/Dfam_3.7_curatedonly_aa.fasta"
    rnas_db = Installation_path + "/db/rRNA_Eukaryota.hmm"
    minDomLTR = 3  # minimum number of domains to automatically kept an LTR retrotransposon in classified module
    minFLNA = 2  # minimum number of full length fragments of copies to automatically kept a non-autonomous element

    ####################################################################################################################
    # Needed by BEE method to extension
    ####################################################################################################################
    max_nns = 10  # threshold of the percentage of Ns allowed in the final extended consensus to keep it (1-100)
    # Parameters for subfamily clustering
    min_perc_model = 0.9
    min_cluster = 8
    max_sequences = 200
    cluster_factor = 10
    group_outliers = True
    max_num_subfamilies = 4

    # Parameter for calculating the saturation
    min_plurality = 40
    end_threshold = 100

    ####################################################################################################################
    # Needed by Unclassified module
    ####################################################################################################################
    FLF_UNCLASS = 2  # minimum number of full length copies to consider a TE in unclassified module
    perc_ident = 70  # Percentage of identity to a known element to keep a TE in unclassified module
    perc_cover = 70  # Percentage of the hit coverage in the TE sequence to keep it in unclassified module
    min_len_unc = 70  # Minimum length of a hit with a TE sequence to keep it in unclassified module
    ####################################################################################################################

    print("\n#########################################################################")
    print("#                                                                       #")
    print("#                             MCHelper                                  #")
    print("#   Instructions:  Execute the program, press q to close the graph and  #")
    print("#     press a number 2-"+str(len(orders_superfamilies)+1)+" if you want to keep the TE, or 1 otherwise.   #")
    print("#                                                                       #")
    print("#########################################################################\n")

    ### read parameters
    parser = argparse.ArgumentParser()
    parser.add_argument('-r', '--module', required=False, dest='module_user',
                        help='module of curation [A, C, U, T, E, M]. Default=A')
    parser.add_argument('-i', '--input', required=False, dest='input_dir',
                        help='Directory with the files required to do the curation (REPET output directory). Required*')
    parser.add_argument('-g', '--genome', required=False, dest='genome',
                        help='Genome used to detect the TEs. Required*')
    parser.add_argument('-o', '--output', required=False, dest='outputdir',
                        help='Path to the output directory. Required*')
    parser.add_argument('--te_aid', required=False, dest='te_aid', default='Y',
                        help='Do you want to use TE-aid? [Y or N]. Default=Y')
    parser.add_argument('-a', required=False, dest='automatic', default='F',
                        help='Level of automation: F: fully automated, S: semi-automated, M: fully manual?. Default=F')
    parser.add_argument('-n', required=False, dest='proj_name',
                        help='REPET project name. Required*')
    parser.add_argument('-t', required=False, dest='cores', default=-1,
                        help='cores to execute some steps in parallel')
    parser.add_argument('-m', required=False, dest='ref_library_module3',
                        help='Path to the sequences to be used as references in the unclassified module')
    parser.add_argument('-v', required=False, dest='verbose', default='N',
                        help='Verbose? [Y or N]. Default=N')
    parser.add_argument('--input_type', required=False, dest='input_type',
                        help='Input type: fasta or REPET.')
    parser.add_argument('-l', required=False, dest='user_library',
                        help='User defined library to be used with input type fasta.')
    parser.add_argument('-b', required=False, dest='busco_library',
                        help='Reference/BUSCO genes to filter out TEs (HMMs expected). ')
    parser.add_argument('-z', required=False, dest='minBlastHits', default=2,
                        help='Minimum number of blast hits to process an element.')
    parser.add_argument('-c', required=False, dest='minFullLenFragments', default=1,
                        help='Minimum number of full-length fragments to process an element.')
    parser.add_argument('-s', required=False, dest='perc_ssr',
                        help='Maximum length covered by single repetitions (in percentage between 0-100) allowed for a TE not to be removed')
    parser.add_argument('-e', required=False, dest='ext_nucl',
                        help='Number of nucleotides to extend each size of the element. Default=500')
    parser.add_argument('-x', required=False, dest='num_ite',
                        help='Number of iterations to extend the elements. Default=16')
    parser.add_argument('-k', required=False, dest='clustering_alg',
                        help='Clustering algorithm: cd-hit or meshclust. Default=cd-hit')
    parser.add_argument('--version', action='version', version='MCHelper version 1.7.0')


    options = parser.parse_args()
    module_user = options.module_user
    input_dir = options.input_dir
    proj_name = options.proj_name
    genome = options.genome
    outputdir = options.outputdir
    te_aid = options.te_aid
    automatic = options.automatic
    cores = options.cores
    ref_library_module3 = options.ref_library_module3
    verbose = options.verbose
    input_type = options.input_type
    user_library = options.user_library
    busco_library = options.busco_library
    minFullLenFragments = int(options.minFullLenFragments)
    minBlastHits = int(options.minBlastHits)
    perc_ssr = options.perc_ssr
    ext_nucl = options.ext_nucl
    clustering_alg = options.clustering_alg
    num_ite = options.num_ite

    ####################################################################################################################
    # Parameter validation
    ####################################################################################################################
    module = 0
    if module_user is None:
        module_user = 'A'
        module = 123
        print("MESSAGE: Missing module (-r) parameter, using by default: " + module_user)
    elif module_user.upper() not in ['A', 'C', 'U', 'E', 'T', 'M', '3333']:  # 3333 for debugging only
        print('FATAL ERROR: module (-r parameter) must be A (all steps), C (only classified TEs), U (TE classification module), E (Consensus extension module), T (TE_Aid in parallel) or M (Manual Inspection module)')
        sys.exit(0)
    else:
        if module_user.upper() == 'A':
            module = 123
        elif module_user.upper() == 'C':
            module = 1
        elif module_user.upper() == 'U':
            module = 3
        elif module_user.upper() == 'E':
            module = 2
        elif module_user.upper() == 'T':
            module = 4
        elif module_user.upper() == 'M':
            module = 5
        elif module_user.upper() == '3333':
            module = 3333

    if outputdir is None:
        outputdir = os.getcwd()
        print('MESSAGE: Output directory will be '+outputdir)
    elif not os.path.exists(outputdir):
        create_output_folders(outputdir)
        outputdir = os.path.abspath(outputdir)
        print('MESSAGE: Output folder ' + outputdir + " created.")
    else:
        outputdir = os.path.abspath(outputdir)

    if verbose is None:
        verbose = False
        print('MESSAGE: Verbose is not activated')
    elif verbose.upper() not in ['Y', 'N']:
        print('FATAL ERROR: verbose (-v) must be Y or N')
        sys.exit(0)
    else:
        if verbose == 'Y':
            verbose = True
        else:
            verbose = False

    if cores is None or cores == -1:
        cores = int(psutil.cpu_count())
        print("MESSAGE: Missing threads parameter, using by default: " + str(cores))
    else:
        cores = int(cores)

    if perc_ssr is None:
        perc_ssr = 60
        print("MESSAGE: Missing -s parameter, using by default: " + str(perc_ssr))
    elif not perc_ssr.isnumeric():
        print('FATAL ERROR: -s must be numeric, but I received: '+str(perc_ssr))
        sys.exit(0)
    elif int(perc_ssr) < 0 or int(perc_ssr) > 100:
        print('FATAL ERROR: -s must be between 0 and 100')
        sys.exit(0)
    else:
        perc_ssr = int(perc_ssr)

    if ext_nucl is None:
        ext_nucl = 500
        print("MESSAGE: Missing -e parameter, using by default: " + str(ext_nucl))
    elif not ext_nucl.isnumeric():
        print('FATAL ERROR: -e must be numeric, but I received: '+str(ext_nucl))
        sys.exit(0)
    elif int(ext_nucl) < 1 or int(ext_nucl) > 5000:
        print('FATAL ERROR: -e must be between 1 and 5000')
        sys.exit(0)
    else:
        ext_nucl = int(ext_nucl)

    if num_ite is None:
        num_ite = 16
        print("MESSAGE: Missing -x parameter, using by default: " + str(num_ite))
    elif not num_ite.isnumeric():
        print('FATAL ERROR: -x must be numeric, but I received: '+str(num_ite))
        sys.exit(0)
    elif int(num_ite) < 0 or int(num_ite) > 100:
        print('FATAL ERROR: -x must be between 0 and 100')
        sys.exit(0)
    else:
        num_ite = int(num_ite)

    if genome is None:
        print('FATAL ERROR: -g parameter must be specified')
        sys.exit(0)
    if not os.path.exists(genome):
        print("FATAL ERROR: Genome file " + genome + " doesn't exist.")
        sys.exit(0)

    if clustering_alg is None:
        clustering_alg = "cd-hit"
    elif clustering_alg.lower() not in ['cd-hit', 'meshclust']:
        print('FATAL ERROR: unknown clustering algorithm defined in -k parameter. Options are cd-hit or meshclust')
        sys.exit(0)
    elif clustering_alg.lower() == 'meshclust':
        path = shutil.which("meshclust")
        if path is None:
            print('FATAL ERROR: meshclust was not find in your environment. Please add the meshclust path to your PATH variable.')
            sys.exit(0)
        else:
            print("MESSAGE: Using " + str(clustering_alg) + " as clustering algorithm")
    elif clustering_alg.lower() == 'cd-hit':
        path = shutil.which("cd-hit")
        if path is None:
            print('FATAL ERROR: cd-hit was not find in your environment. Please install cd-hit in your conda environment.')
            sys.exit(0)
        else:
            print("MESSAGE: Using " + str(clustering_alg) + " as clustering algorithm")

    ####################################################################################################################
    # Classified module
    ####################################################################################################################
    if module in [1, 123]:
        if te_aid is None:
            te_aid = 'Y'
            print('MESSAGE: Using by default te_aid = Y')
        elif te_aid.upper() not in ['Y', 'N']:
            print('FATAL ERROR: unknown value of --te_aid parameter: ' + te_aid + '. This parameter must be Y or N')
            sys.exit(0)
        else:
            te_aid = te_aid.upper()
        if automatic is not None:
            if automatic.upper() not in ['F', 'S', 'M']:
                print('FATAL ERROR: automation level (-a) must be F, S or M (see help.)')
                sys.exit(0)
            else:
                automatic = automatic.upper()
        if input_type == None:
            print(
                'FATAL ERROR: You need to specify the input type parameter (--input_type). Values can be fasta or repet')
            sys.exit(0)
        elif input_type.upper() not in ['FASTA', 'REPET']:
            print(
                'FATAL ERROR: Unknown value in the input type parameter (--input_type). Values must be fasta or repet')
            sys.exit(0)
        else:
            input_type = input_type.lower()
        if (module == 1 or module == 123) and busco_library is None:
            print('FATAL ERROR: -b parameter must be specified for the unclassified module')
            sys.exit(0)
        elif (module == 1 or module == 123) and not os.path.exists(busco_library):
            print("FATAL ERROR: Reference/BUSCO genes file " + busco_library + " doesn't exist.")
            sys.exit(0)

        ################################################################################################################
        # classified module's Pre-validations :
        ################################################################################################################
        print("MESSAGE: Starting with Classified module...")
        use_repet = True
        if input_type == 'repet':
            if input_dir is None:
                print('FATAL ERROR: -i parameter must be specified in for the classified module and input type REPET')
                sys.exit(0)
            if proj_name is None:
                print('FATAL ERROR: -n parameter must be specified in for the classified module and input type REPET')
                sys.exit(0)

            start_time = time.time()
            input_valid, reason_valid = check_repet_input_folder(input_dir, proj_name)
            end_time = time.time()
            if verbose:
                print("MESSAGE: REPET Input checking done: [" + str(end_time - start_time) + " seconds]")

            if input_valid:
                ref_tes = input_dir + "/" + proj_name + "_refTEs.fa"
                features_table = input_dir + "/" + proj_name + "_denovoLibTEs_PC.classif"
                plots_dir = input_dir + "/plotCoverage"
                gff_files = input_dir + "/gff_reversed"

                if checkChrNames(genome) is False:
                    print("FATAL ERROR: The provided genome ("+genome+") has sequences IDs containing only numbers. Please renamed them to contain letters and numbers (e.g. >Chr1 instead of >1).")
                    sys.exit(0)
            else:
                print(reason_valid)
                sys.exit(0)
        elif input_type == 'fasta':
            use_repet = False
            if user_library is None:
                print('FATAL ERROR: -l parameter must be specified in for the classified module using input type fasta')
                sys.exit(0)
            if not os.path.exists(user_library):
                print("FATAL ERROR: TE library file " + user_library + " doesn't exist.")
                sys.exit(0)

            start_time = time.time()
            check = check_classification_Userlibrary(user_library, outputdir)
            if check == 0:
                if not os.path.exists(outputdir + "/classifiedModule/denovoLibTEs_PC.classif") and not os.path.exists(
                        outputdir + "/classifiedModule/new_user_lib.fa"):
                    if automatic == 'M':
                        do_blast = True
                    else:
                        do_blast = False
                features_table = outputdir + "/classifiedModule/denovoLibTEs_PC.classif"

                if checkChrNames(genome) is False:
                    print("FATAL ERROR: The provided genome ("+genome+") has sequences IDs containing only numbers. Please renamed them to contain letters and numbers (e.g. >Chr1 instead of >1).")
                    sys.exit(0)
            elif check == -2:
                print('FATAL ERROR: There are sequences with duplicated IDs. Please check them in the file: ' + outputdir + '/sequences_with_problems.txt')
                sys.exit(0)
            else:
                print(
                    'WARNING: There are some sequences with problems in your library and MCHelper cannot process them. Please check them in the file: ' + outputdir + '/sequences_with_problems.txt')

            gff_files = ""
            plots_dir = ""
            ref_tes = outputdir + "/candidate_tes.fa"
            end_time = time.time()
            if verbose:
                print("MESSAGE: Fasta pre-processing done: [" + str(end_time - start_time) + " seconds]")

        if not os.path.exists(outputdir + "/classifiedModule/kept_seqs_classified_module_curated.fa"):

            ############################################################################################################
            # First step: Reduce redundancy
            ############################################################################################################
            start_time = time.time()
            # If repet, we need to include the classification info into the library
            if use_repet:
                rename_tes = []
                struc_table = pd.read_csv(features_table, sep='\t')
                for i in range(struc_table.shape[0]):
                    is_in_lib = [x for x in SeqIO.parse(ref_tes, "fasta") if
                                 str(x.id).split("#")[0] == struc_table.at[i, "Seq_name"]]
                    if len(is_in_lib) > 0:
                        bad_named_te = is_in_lib[0]
                        classif = ""
                        if str(struc_table.at[i, "class"]).upper() in ["UNCLASSIFIED", "UNKNOWN", "NA", "NAN"]:
                            classif = "UNCLASSIFIED"
                        else:
                            classif = "CLASS" + str(struc_table.at[i, "class"])
                            if str(struc_table.at[i, "order"]).upper() not in ["LARD", "TRIM"]:
                                classif += "/" + str(struc_table.at[i, "order"])
                                if str(struc_table.at[i, "sFamily"]).upper() != ["NA"]:
                                    classif += "/" + str(struc_table.at[i, "sFamily"])
                            else:
                                classif += "/LTR/" + str(struc_table.at[i, "order"])

                        bad_named_te.id = str(bad_named_te.id).split("#")[0] + "#" + classif
                        bad_named_te.description = ""
                        rename_tes.append(bad_named_te)

                write_sequences_file(rename_tes, ref_tes + "_tmp")
                delete_files(ref_tes)
                shutil.move(ref_tes + "_tmp", ref_tes)
            if clustering_alg == "cd-hit":
                ref_tes_non_redundat = run_cdhit(ref_tes, outputdir, cores, 0.95, 0.98)
            else:
                ref_tes_non_redundat = run_meshclust(ref_tes, outputdir, cores, 0.95, 0.98)
            delete_files(outputdir + "/non_redundant_lib.fa.clstr")

            ############################################################################################################
            # Second step: Extend all the consensus
            ############################################################################################################
            if not os.path.exists(outputdir + "/classifiedModule/extended_cons.fa"):
                start_time = time.time()
                run_extension_by_saturation_parallel(genome, ref_tes_non_redundat, ext_nucl, num_ite, outputdir+ "/classifiedModule/", minBlastHits, cores,
                                                         min_perc_model, min_cluster, max_sequences, cluster_factor,
                                                         group_outliers, min_plurality, end_threshold, max_num_subfamilies)
                delete_files(outputdir + "/non_redundant_lib.fa")
                ref_tes = outputdir + "/classifiedModule/extended_cons.fa"
                end_time = time.time()
                if verbose:
                    print("MESSAGE: The sequences were extended successfully [" + str(end_time - start_time) + " seconds]")
            else:
                delete_files(outputdir + "/non_redundant_lib.fa")
                ref_tes = outputdir + "/classifiedModule/extended_cons.fa"
                if verbose:
                    print("MESSAGE: The extension was already run, please verify or change the output directory")

            ############################################################################################################
            # Third step: extract seqs with at least one full length fragment and minimum minFullLenFragments
            ############################################################################################################
            start_time = time.time()
            flf_file = count_flf_fasta(ref_tes, genome, cores, outputdir + "/classifiedModule")
            new_ref_tes, num_copies = filter_flf(ref_tes, flf_file, minFullLenFragments, outputdir+"/classifiedModule/")
            end_time = time.time()
            if verbose:
                print("MESSAGE: The library was reduced to " + str(len(list(SeqIO.parse(new_ref_tes, 'fasta')))) + " after FLF filtering [" + str(end_time - start_time) + " seconds]")

            if len(list(SeqIO.parse(new_ref_tes, 'fasta'))) == 0:
                print("FATAL ERROR: There is no TEs with at least "+str(minFullLenFragments)+" full length copies in the genome.")
                sys.exit(0)

            ############################################################################################################
            # Fourth step: SSR, BUSCO genes, tRNA and rRNA filters
            ############################################################################################################
            if not os.path.exists(outputdir + "/classifiedModule/putative_TEs.fa"):
                start_time = time.time()
                new_ref_tes, deleted_seqs = filter_bad_candidates(new_ref_tes, perc_ssr, outputdir+"/classifiedModule/", tools_path, busco_library, rnas_db, cores)
                end_time = time.time()
                if verbose:
                    print("MESSAGE: The library was reduced to " + str(len(list(SeqIO.parse(new_ref_tes, 'fasta')))) + " after SSR, genes and RNA filtering [" + str(end_time - start_time) + " seconds]")

                if len(list(SeqIO.parse(new_ref_tes, 'fasta'))) == 0:
                    print("FATAL ERROR: After filtering false positive TEs, there is no elements in the library.")
                    sys.exit(0)
            else:
                if verbose:
                    print("MESSAGE: The False Positive filtering was already run, please verify or change the output directory")

            ############################################################################################################
            # Fifth step: Structural checks, BEE method, and Visualize plots
            ############################################################################################################
            start_time = time.time()
            new_module1(plots_dir, new_ref_tes, gff_files, outputdir, proj_name, te_aid, automatic, minDomLTR,
                        num_copies, minFLNA, verbose, use_repet, blastn_db, blastx_db, tools_path,
                        ref_profiles, library_path, min_perc_model)

        else:
            print("MESSAGE: classified module was already run, please verify or change the output directory")

        if module == 123:
            new_ref_library = []
            if os.path.exists(outputdir + "/classifiedModule/kept_seqs_classified_module_curated.fa"):
                for te in SeqIO.parse(outputdir + "/classifiedModule/kept_seqs_classified_module_curated.fa", "fasta"):
                    new_ref_library.append(te)

            if os.path.exists(outputdir + "/classifiedModule/kept_seqs_classified_module_non_curated.fa"):
                for te in SeqIO.parse(outputdir + "/classifiedModule/kept_seqs_classified_module_non_curated.fa", "fasta"):
                    new_ref_library.append(te)

            if len(new_ref_library) > 0:
                write_sequences_file(new_ref_library, outputdir + "/classifiedModule/kept_seqs_classified_module.fa")
                ref_library_module3 = outputdir + "/classifiedModule/kept_seqs_classified_module.fa"
            else:
                print("WARNING: Classified module didn't find any TE")
                ref_library_module3 = ""
            module3_seqs_file = outputdir + "/classifiedModule/input_to_unclassified_module_seqs.fa"
        delete_files(outputdir + "/candidate_tes.fa")

    ####################################################################################################################
    # Unclassified module
    ####################################################################################################################
    if module in [3, 123]:
        if module == 3 and user_library is None:
            print('FATAL ERROR: -l parameter must be specified for the Classification Module')
            sys.exit(0)
        if module == 3 and ref_library_module3 is None:
            print('FATAL ERROR: -m parameter must be specified for the unclassified module')
            sys.exit(0)
        if module == 3 and genome is None:
            print('FATAL ERROR: -g parameter must be specified for the unclassified module')
            sys.exit(0)
        if module == 3 and busco_library is None:
            print('FATAL ERROR: -b parameter must be specified for the unclassified module')
            sys.exit(0)
        elif module == 3 and not os.path.exists(busco_library):
            print("FATAL ERROR: Reference/BUSCO genes file " + busco_library + " doesn't exist.")
            sys.exit(0)
        if module == 123 and (os.path.getsize(module3_seqs_file) == 0 or module3_seqs_file == ""):
            print("MESSAGE: There is no sequences to unclassified Module, skipping ...")
        else:
            if ref_library_module3 != "":
                create_output_folders(outputdir + "/unclassifiedModule")
                print("MESSAGE: Starting with Unclassified module...")

                ########################################################################################################
                # First step: run BEE in parallel
                ########################################################################################################
                if module == 3:
                    start_time = time.time()
                    check = check_classification_Userlibrary(user_library, outputdir)
                    if check == -2:
                        print(
                            'FATAL ERROR: There are sequences with duplicated IDs. Please check them in the file: ' + outputdir + '/sequences_with_problems.txt')
                        sys.exit(0)
                    elif check != 0:
                        print(
                            'WARNING: There are some sequences with problems in your library and MCHelper cannot process them. Please check them in the file: ' + outputdir + '/sequences_with_problems.txt')

                    module3_seqs_file = outputdir + "/candidate_tes.fa"
                    run_extension_by_saturation_parallel(genome, module3_seqs_file, ext_nucl, num_ite,
                                                         outputdir + "/unclassifiedModule/", minBlastHits, cores,
                                                         min_perc_model, min_cluster, max_sequences, cluster_factor,
                                                         group_outliers, min_plurality, end_threshold, max_num_subfamilies)
                    end_time = time.time()
                    if verbose:
                        print("MESSAGE: The sequences were extended successfully [" + str(
                            end_time - start_time) + " seconds]")

                    start_time = time.time()
                    extendedSeqs = outputdir + "/unclassifiedModule/extended_cons.fa"

                    ########################################################################################################
                    # Second step: Filter elements with not enough FLF copies in the genome
                    ########################################################################################################
                    flf_file = count_flf_fasta(extendedSeqs, genome, cores, outputdir + "/unclassifiedModule")
                    module3_seqs_file, num_copies = filter_flf(extendedSeqs, flf_file, FLF_UNCLASS, outputdir + "/unclassifiedModule/")
                    end_time = time.time()
                    if verbose:
                        print("MESSAGE: The library was reduced to " + str(
                            len(list(SeqIO.parse(module3_seqs_file, 'fasta')))) + " after FLF filtering [" + str(
                            end_time - start_time) + " seconds]")

                    ############################################################################################################
                    # Third step: SSR, BUSCO genes, tRNA and rRNA filters
                    ############################################################################################################
                    start_time = time.time()
                    module3_seqs_file, deleted_seqs = filter_bad_candidates(module3_seqs_file, perc_ssr,
                                                                      outputdir + "/unclassifiedModule/", tools_path,
                                                                      busco_library, rnas_db, cores)
                    end_time = time.time()
                    if verbose:
                        print("MESSAGE: The library was reduced to " + str(len(list(
                            SeqIO.parse(module3_seqs_file, 'fasta')))) + " after SSR, genes and RNA filtering [" + str(
                            end_time - start_time) + " seconds]")

                    if len(list(SeqIO.parse(module3_seqs_file, 'fasta'))) == 0:
                        print(
                            "FATAL ERROR: After filtering false positive TEs, there is no elements in the library.")
                        sys.exit(0)


                else:  # do not run BEE because it was run in Classified module
                    ########################################################################################################
                    # Second step: Filter elements with not enough FLF copies in the genome
                    ########################################################################################################
                    start_time = time.time()
                    module3_seqs_file, num_copies = filter_flf(module3_seqs_file, flf_file, FLF_UNCLASS,
                                                               outputdir + "/unclassifiedModule/")
                    extendedSeqs = module3_seqs_file
                    end_time = time.time()
                    if verbose:
                        print("MESSAGE: The library was reduced to " + str(
                            len(list(SeqIO.parse(module3_seqs_file, 'fasta')))) + " after FLF filtering [" + str(
                            end_time - start_time) + " seconds]")

                ########################################################################################################
                # Third step: find structural features
                ########################################################################################################
                start_time = time.time()
                build_class_table_parallel(extendedSeqs, cores, outputdir + '/unclassifiedModule/',
                                           blastn_db, blastx_db, ref_profiles, False)
                end_time = time.time()
                if verbose:
                    print("MESSAGE: TE Feature table was created [" + str(end_time - start_time) + " seconds]")

                ########################################################################################################
                # Fourth step: process the sequences, BLASTn againts a given curated library, inferred classifications,
                # and create outputs
                ########################################################################################################
                module3(extendedSeqs, ref_library_module3, cores, outputdir, perc_ident, perc_cover, min_len_unc, library_path)
            else:
                if module == 123:
                    print("WARNING: Unclassified module didn't run because classified module didn't find any TE")
                else:
                    print("WARNING: "+ref_library_module3+" is empty")

    ####################################################################################################################
    # BEE module
    ####################################################################################################################
    if module == 2:
        if user_library is None:
            print('FATAL ERROR: -l parameter must be specified for the BEE extension module using input type fasta')
            sys.exit(0)
        if not os.path.exists(user_library):
            print("FATAL ERROR: TE library file " + user_library + " doesn't exist.")
            sys.exit(0)
        if genome is None:
            print('FATAL ERROR: -g parameter must be specified in for the BEE extension module')
            sys.exit(0)
        if not os.path.exists(genome):
            print("FATAL ERROR: Genome file " + genome + " doesn't exist.")
            sys.exit(0)

        ########################################################################################################
        # First step: run BEE in parallel
        ########################################################################################################
        create_output_folders(outputdir + "/bee_module/")
        run_extension_by_saturation_parallel(genome, user_library, ext_nucl, num_ite,
                                             outputdir + "/bee_module/", minBlastHits, cores,
                                             min_perc_model, min_cluster, max_sequences, cluster_factor,
                                             group_outliers, min_plurality, end_threshold, max_num_subfamilies)

    ####################################################################################################################
    # TE+aid in Parallel
    ####################################################################################################################
    if module == 4:
        if user_library is None:
            print('FATAL ERROR: -l parameter must be specified for the BEE extension module using input type fasta')
            sys.exit(0)
        if not os.path.exists(user_library):
            print("FATAL ERROR: TE library file " + user_library + " doesn't exist.")
            sys.exit(0)
        if genome is None:
            print('FATAL ERROR: -g parameter must be specified in for the BEE extension module')
            sys.exit(0)
        if not os.path.exists(genome):
            print("FATAL ERROR: Genome file " + genome + " doesn't exist.")
            sys.exit(0)

        ########################################################################################################
        # First step: run TE+Aid in parallel
        ########################################################################################################
        run_te_aid_parallel(tools_path + "/TE-Aid-master/", genome, user_library, outputdir + "/", cores,
                            min_perc_model)

    ####################################################################################################################
    # Manual Inspection module
    ####################################################################################################################
    if module == 5:
        if user_library is None:
            print('FATAL ERROR: -l parameter must be specified for the BEE extension module using input type fasta')
            sys.exit(0)
        if not os.path.exists(user_library):
            print("FATAL ERROR: TE library file " + user_library + " doesn't exist.")
            sys.exit(0)
        if genome is None:
            print('FATAL ERROR: -g parameter must be specified in for the BEE extension module')
            sys.exit(0)
        if not os.path.exists(genome):
            print("FATAL ERROR: Genome file " + genome + " doesn't exist.")
            sys.exit(0)
        if te_aid is None:
            te_aid = 'Y'
            print('MESSAGE: Using by default te_aid = Y')
        elif te_aid.upper() not in ['Y', 'N']:
            print('FATAL ERROR: unknown value of --te_aid parameter: ' + te_aid + '. This parameter must be Y or N')
            sys.exit(0)
        else:
            te_aid = te_aid.upper()
        if input_type == None:
            print(
                'FATAL ERROR: You need to specify the input type parameter (--input_type). Values can be fasta or repet')
            sys.exit(0)
        elif input_type.upper() not in ['FASTA', 'REPET']:
            print(
                'FATAL ERROR: Unknown value ('+input_type+') in the input type parameter (--input_type). Values must be fasta or repet')
            sys.exit(0)
        else:
            input_type = input_type.lower()

        ########################################################################################################
        # Checking that input is fine
        ########################################################################################################
        use_repet = True
        if input_type == 'repet':
            if input_dir is None:
                print('FATAL ERROR: -i parameter must be specified in for the classified module and input type REPET')
                sys.exit(0)
            if proj_name is None:
                print('FATAL ERROR: -n parameter must be specified in for the classified module and input type REPET')
                sys.exit(0)

            start_time = time.time()
            input_valid, reason_valid = check_repet_input_folder(input_dir, proj_name)
            end_time = time.time()
            if verbose:
                print("MESSAGE: REPET Input checking done: [" + str(end_time - start_time) + " seconds]")

            if input_valid:
                ref_tes = input_dir + "/" + proj_name + "_refTEs.fa"
                features_table = input_dir + "/" + proj_name + "_denovoLibTEs_PC.classif"
                plots_dir = input_dir + "/plotCoverage"
                gff_files = input_dir + "/gff_reversed"
            else:
                print(reason_valid)
                sys.exit(0)
        elif input_type == 'fasta':
            use_repet = False
            if user_library is None:
                print('FATAL ERROR: -l parameter must be specified in for the classified module using input type fasta')
                sys.exit(0)
            if not os.path.exists(user_library):
                print("FATAL ERROR: TE library file " + user_library + " doesn't exist.")
                sys.exit(0)

            start_time = time.time()
            if check_classification_Userlibrary(user_library, outputdir) == 0:
                if not os.path.exists(outputdir + "/classifiedModule/denovoLibTEs_PC.classif") and not os.path.exists(
                        outputdir + "/classifiedModule/new_user_lib.fa"):
                    if automatic == 'M':
                        do_blast = True
                    else:
                        do_blast = False
                features_table = outputdir + "/classifiedModule/denovoLibTEs_PC.classif"

            else:
                print(
                    'WARNING: There are some sequences with problems in your library and MCHelper cannot process them. Please check them in the file: ' + outputdir + '/sequences_with_problems.txt')

            gff_files = ""
            plots_dir = ""
            user_library = outputdir + "/candidate_tes.fa"
            end_time = time.time()
            if verbose:
                print("MESSAGE: Fasta pre-processing done: [" + str(end_time - start_time) + " seconds]")

        ########################################################################################################
        # First step: Build necessary files
        ########################################################################################################
        seqID_list = [str(x.id).split("#")[0] for x in SeqIO.parse(user_library, "fasta")]
        build_class_table_parallel(user_library, cores, outputdir,
                                   blastn_db, blastx_db, ref_profiles, False)
        struc_table = pd.read_csv(outputdir + "/denovoLibTEs_PC.classif", sep='\t')

        ########################################################################################################
        # Second step: Filter elements with not enough FLF copies in the genome
        ########################################################################################################
        flf_file = count_flf_fasta(user_library, genome, cores, outputdir)
        user_library_2, num_copies = filter_flf(user_library, flf_file, 0, outputdir)

        ########################################################################################################
        # Second step: Manual Inspection of all the sequences
        ########################################################################################################
        seqs_to_module3, keep_seqs, orders, kept_seqs_record, non_curated, orders_incomplete = manual_inspection(genome,
        outputdir, user_library, user_library, seqID_list, struc_table, te_aid, use_repet, "M", proj_name, plots_dir,
        gff_files, min_perc_model, [], [], [], [], [], num_copies)

        ########################################################################################################
        # Third step: Save results
        ########################################################################################################
        for index in range(len(kept_seqs_record)):
            # put the order having the superfamily
            classification = dicc_orders[orders[index]]
            if orders[index] >= 3 and orders[index] <= 8:
                classification = "LTR/" + classification
            elif orders[index] >= 11 and orders[index] <= 19:
                classification = "LINE/" + classification
            elif orders[index] >= 21 and orders[index] <= 23:
                classification = "DIRS/" + classification
            elif orders[index] >= 26 and orders[index] <= 36:
                classification = "TIR/" + classification
            elif orders[index] == 40:
                classification = "UNCLASSIFIED"

            # put the class having the order/superfamily
            if orders[index] >= 2 and orders[index] <= 23:
                classification = "CLASSI/" + classification
            elif orders[index] >= 24 and orders[index] <= 39:
                classification = "CLASSII/" + classification

            new_name = keep_seqs[index] + "#" + classification
            kept_seqs_record[index].id = new_name
            kept_seqs_record[index].description = ""

        for index in range(len(non_curated)):
            # put the order having the superfamily
            classification = dicc_orders[orders_incomplete[index]]
            if orders_incomplete[index] >= 3 and orders_incomplete[index] <= 8:
                classification = "LTR/" + classification
            elif orders_incomplete[index] >= 11 and orders_incomplete[index] <= 19:
                classification = "LINE/" + classification
            elif orders_incomplete[index] >= 21 and orders_incomplete[index] <= 23:
                classification = "DIRS/" + classification
            elif orders_incomplete[index] >= 26 and orders_incomplete[index] <= 36:
                classification = "TIR/" + classification
            elif orders_incomplete[index] == 40:
                classification = "UNCLASSIFIED"

            # put the class having the order/superfamily
            if orders_incomplete[index] >= 2 and orders_incomplete[index] <= 23:
                classification = "CLASSI/" + classification
            elif orders_incomplete[index] >= 24 and orders_incomplete[index] <= 39:
                classification = "CLASSII/" + classification

            new_name = str(non_curated[index].id).split("#")[0] + "#" + classification
            non_curated[index].id = new_name
            non_curated[index].description = ""

        seqs_to_module3_record = [te for te in SeqIO.parse(user_library, "fasta") if
                                  str(te.id).split("#")[0] in seqs_to_module3]
        write_sequences_file(kept_seqs_record, outputdir + "/kept_seqs_curated.fa")
        write_sequences_file(seqs_to_module3_record,
                             outputdir + "/unclassified_seqs.fa")

        delete_files(outputdir + "/extended_cons.fa")
        delete_files(outputdir + "/putative_TEs.fa")
        delete_files(outputdir + "/new_user_lib.fa")

    ####################################################################################################################
    # Debugging only !!!!
    ####################################################################################################################
    if module in [3333]:
        print("Debugging....")
        """build_class_table_parallel(user_library, cores, outputdir,
                                   blastn_db, blastx_db, ref_profiles, False)"""
        run_te_aid_parallel(tools_path + "/TE-Aid-master/", genome, user_library, outputdir + "/", cores, min_perc_model)


    ####################################################################################################################
    # Writing the final results
    final_seqs = []
    if module in [1, 12, 123]:
        try:
            for te in SeqIO.parse(outputdir + "/classifiedModule/kept_seqs_classified_module_curated.fa", "fasta"):
                final_seqs.append(te)
        except FileNotFoundError:
            print("WARNING: the file " + outputdir + "/classifiedModule/kept_seqs_classified_module_curated.fa was not created. Please check.")
        try:
            for te in SeqIO.parse(outputdir + "/classifiedModule/kept_seqs_classified_module_non_curated.fa", "fasta"):
                final_seqs.append(te)
        except FileNotFoundError:
            print("WARNING: the file " + outputdir + "/classifiedModule/kept_seqs_classified_module_non_curated.fa was not created. Please check.")
        delete_files(outputdir + "/classifiedModule/non_redundant_lib.fa")
        delete_files(outputdir + "/classifiedModule/non_redundant_lib.fa.clstr")
    if module in [3, 123]:
        try:
            for te in SeqIO.parse(outputdir + "/unclassifiedModule/kept_seqs_unclassified_module.fa", "fasta"):
                final_seqs.append(te)
        except FileNotFoundError:
            print("WARNING: the file " + outputdir + "/unclassifiedModule/kept_seqs_unclassified_module.fa was not created. Please check.")

    # Reduce the final redundancy
    if len(final_seqs) > 0:
        write_sequences_file(final_seqs, outputdir + "/curated_sequences_R.fa")
        if clustering_alg == "cd-hit":
            non_redundat = run_cdhit(outputdir + "/curated_sequences_R.fa", outputdir, cores, 0.95, 0.98)
        else:
            non_redundat = run_meshclust(outputdir + "/curated_sequences_R.fa", outputdir, cores, 0.95, 0.98)

        shutil.move(non_redundat, outputdir + "/curated_sequences_NR.fa")
```
