## Supplemental Code for "MCHelper automatically curates transposable element libraries across eukaryotic species": MCHelper.yml

```
name: MCHelper
channels:
  - conda-forge
  - bioconda
  - defaults
dependencies:
  - _libgcc_mutex=0.1=main
  - _openmp_mutex=5.1=1_gnu
  - _r-mutex=1.0.0=anacondar_1
  - binutils_impl_linux-64=2.38=h2a08ee3_1
  - binutils_linux-64=2.38.0=hc2dff05_0
  - biopython=1.78=py38h7f8727e_0
  - blas=1.0=openblas
  - blast=2.10.1=pl526he19e7b1_3
  - bottleneck=1.3.5=py38h7deecbd_0
  - brotli=1.0.9=h5eee18b_7
  - brotli-bin=1.0.9=h5eee18b_7
  - bwidget=1.9.11=1
  - bzip2=1.0.8=h7b6447c_0
  - c-ares=1.19.0=h5eee18b_0
  - ca-certificates=2023.05.30=h06a4308_0
  - cairo=1.16.0=hb05425b_4
  - cd-hit=4.8.1=h2e03b76_5
  - cialign=1.0.18=pyhb7b1952_0
  - configargparse=1.4=pyhd3eb1b0_0
  - contourpy=1.0.5=py38hdb19cb5_0
  - curl=7.88.1=h5eee18b_1
  - cycler=0.11.0=pyhd3eb1b0_0
  - dbus=1.13.18=hb2f20db_0
  - eigen=3.3.7=hd09550d_1
  - emboss=6.6.0=h6debe1e_0
  - entrez-direct=16.2=he881be0_1
  - expat=2.4.9=h6a678d5_0
  - ffmpeg=4.2.2=h20bf706_0
  - fontconfig=2.14.1=h52c9d5c_1
  - fonttools=4.25.0=pyhd3eb1b0_0
  - freetype=2.12.1=h4a9f257_0
  - fribidi=1.0.10=h7b6447c_0
  - gcc_impl_linux-64=11.2.0=h1234567_1
  - gcc_linux-64=11.2.0=h5c386dc_0
  - gfortran_impl_linux-64=11.2.0=h1234567_1
  - gfortran_linux-64=11.2.0=hc2dff05_0
  - giflib=5.2.1=h5eee18b_3
  - glib=2.69.1=he621ea3_2
  - gmp=6.2.1=h295c915_3
  - gnutls=3.6.15=he1e5248_0
  - graphite2=1.3.14=h295c915_1
  - gst-plugins-base=1.14.1=h6a678d5_1
  - gstreamer=1.14.1=h5eee18b_1
  - gxx_impl_linux-64=11.2.0=h1234567_1
  - gxx_linux-64=11.2.0=hc2dff05_0
  - harfbuzz=4.3.0=hf52aaf7_1
  - hdf5=1.10.6=h3ffc7dd_1
  - hmmer=3.3.2=h87f3376_2
  - icu=58.2=he6710b0_3
  - importlib_resources=5.2.0=pyhd3eb1b0_1
  - joblib=1.2.0=py38h06a4308_0
  - jpeg=9e=h5eee18b_1
  - kernel-headers_linux-64=2.6.32=he073ed8_15
  - kiwisolver=1.4.4=py38h6a678d5_0
  - krb5=1.20.1=hf590b9c_0
  - lame=3.100=h7b6447c_0
  - lcms2=2.12=h3be6417_0
  - ld_impl_linux-64=2.38=h1181459_1
  - lerc=3.0=h295c915_0
  - libblas=3.9.0=16_linux64_openblas
  - libbrotlicommon=1.0.9=h5eee18b_7
  - libbrotlidec=1.0.9=h5eee18b_7
  - libbrotlienc=1.0.9=h5eee18b_7
  - libcblas=3.9.0=16_linux64_openblas
  - libclang=10.0.1=default_hb85057a_2
  - libcurl=7.88.1=h251f7ec_1
  - libdeflate=1.17=h5eee18b_0
  - libedit=3.1.20221030=h5eee18b_0
  - libev=4.33=h7f8727e_1
  - libevent=2.1.12=hdbd6064_1
  - libffi=3.4.4=h6a678d5_0
  - libgcc-devel_linux-64=11.2.0=h1234567_1
  - libgcc-ng=11.2.0=h1234567_1
  - libgd=2.3.3=h6a678d5_2
  - libgfortran-ng=11.2.0=h00389a5_1
  - libgfortran5=11.2.0=h1234567_1
  - libgomp=11.2.0=h1234567_1
  - libidn2=2.3.4=h5eee18b_0
  - libllvm10=10.0.1=hbcb73fb_5
  - libnghttp2=1.52.0=h2d74bed_0
  - libopenblas=0.3.21=h043d6bf_0
  - libopus=1.3.1=h7b6447c_0
  - libpng=1.6.39=h5eee18b_0
  - libpq=12.15=h251f7ec_0
  - libprotobuf=3.20.3=he621ea3_0
  - libssh2=1.10.0=hdbd6064_1
  - libstdcxx-devel_linux-64=11.2.0=h1234567_1
  - libstdcxx-ng=11.2.0=h1234567_1
  - libtasn1=4.19.0=h5eee18b_0
  - libtiff=4.5.0=h6a678d5_2
  - libunistring=0.9.10=h27cfd23_0
  - libuuid=1.41.5=h5eee18b_0
  - libvpx=1.7.0=h439df22_0
  - libwebp=1.2.4=h11a3e52_1
  - libwebp-base=1.2.4=h5eee18b_1
  - libxcb=1.15=h7f8727e_0
  - libxkbcommon=1.0.1=hfa300c1_0
  - libxml2=2.9.14=h74e7548_0
  - libxslt=1.1.35=h4e12654_0
  - lz4-c=1.9.4=h6a678d5_0
  - mafft=7.505=hec16e2b_0
  - make=4.2.1=h1bed415_1
  - matplotlib=3.7.1=py38h578d9bd_0
  - matplotlib-base=3.7.1=py38h417a72b_1
  - munkres=1.1.4=py_0
  - ncurses=6.4=h6a678d5_0
  - nettle=3.7.3=hbbd107a_1
  - nspr=4.35=h6a678d5_0
  - nss=3.89.1=h6a678d5_0
  - numexpr=2.8.4=py38hd2a5715_1
  - numpy=1.24.3=py38hf838250_0
  - numpy-base=1.24.3=py38h1e6e340_0
  - opencv=4.6.0=py38h417a72b_3
  - openh264=2.1.1=h4ff587b_0
  - openjpeg=2.4.0=h3ad879b_0
  - openssl=3.0.9=h7f8727e_0
  - packaging=23.0=py38h06a4308_0
  - pandas=1.5.3=py38h417a72b_0
  - pango=1.50.7=h05da053_0
  - pcre=8.45=h295c915_0
  - pcre2=10.37=he7ceb23_1
  - pdf2image=1.16.3=pyhd8ed1ab_0
  - perl=5.26.2=h14c3975_0
  - perl-archive-tar=2.32=pl526_0
  - perl-carp=1.38=pl526_3
  - perl-common-sense=3.74=pl526_2
  - perl-compress-raw-bzip2=2.087=pl526he1b5a44_0
  - perl-compress-raw-zlib=2.087=pl526hc9558a2_0
  - perl-exporter=5.72=pl526_1
  - perl-exporter-tiny=1.002001=pl526_0
  - perl-extutils-makemaker=7.36=pl526_1
  - perl-io-compress=2.087=pl526he1b5a44_0
  - perl-io-zlib=1.10=pl526_2
  - perl-json=4.02=pl526_0
  - perl-json-xs=2.34=pl526h6bb024c_3
  - perl-list-moreutils=0.428=pl526_1
  - perl-list-moreutils-xs=0.428=pl526_0
  - perl-pathtools=3.75=pl526h14c3975_1
  - perl-scalar-list-utils=1.52=pl526h516909a_0
  - perl-types-serialiser=1.0=pl526_2
  - perl-xsloader=0.24=pl526_0
  - pillow=9.4.0=py38h6a678d5_0
  - pip=23.1.2=py38h06a4308_0
  - pixman=0.40.0=h7f8727e_1
  - ply=3.11=py38_0
  - psutil=5.9.0=py38h5eee18b_0
  - pyparsing=3.0.9=py38h06a4308_0
  - pyqt=5.15.7=py38h6a678d5_1
  - pyqt5-sip=12.11.0=py38h6a678d5_1
  - python=3.8.16=h955ad1f_4
  - python-dateutil=2.8.2=pyhd3eb1b0_0
  - python_abi=3.8=2_cp38
  - pytz=2022.7=py38h06a4308_0
  - qt-main=5.15.2=h327a75a_7
  - qt-webengine=5.15.9=hd2b0992_4
  - qtwebkit=5.212=h4eab89a_4
  - r=4.2=r42hd8ed1ab_1006
  - r-base=4.2.0=h1ae530e_0
  - r-boot=1.3_28.1=r42hc72bb7e_1
  - r-class=7.3_20=r42h76d94ec_0
  - r-cluster=2.1.3=r42h640688f_0
  - r-codetools=0.2_19=r42hc72bb7e_1
  - r-foreign=0.8_82=r42h76d94ec_0
  - r-kernsmooth=2.23_20=r42h640688f_0
  - r-lattice=0.20_45=r42h76d94ec_0
  - r-mass=7.3_57=r42h76d94ec_0
  - r-matrix=1.4_1=r42h76d94ec_0
  - r-mgcv=1.8_40=r42h76d94ec_0
  - r-nlme=3.1_157=r42h640688f_0
  - r-nnet=7.3_17=r42h76d94ec_0
  - r-recommended=4.2=r42hd8ed1ab_1005
  - r-rpart=4.1.16=r42h76d94ec_0
  - r-spatial=7.3_15=r42h76d94ec_0
  - r-survival=3.3_1=r42h76d94ec_0
  - readline=8.2=h5eee18b_0
  - scikit-learn=1.0.2=py38h1561384_0
  - scipy=1.9.3=py38h32ae08f_2
  - setuptools=67.8.0=py38h06a4308_0
  - sip=6.6.2=py38h6a678d5_0
  - six=1.16.0=pyhd3eb1b0_1
  - sqlite=3.41.2=h5eee18b_0
  - sysroot_linux-64=2.12=he073ed8_15
  - threadpoolctl=2.2.0=pyh0d69192_0
  - tk=8.6.12=h1ccaba5_0
  - tktable=2.10=h14c3975_0
  - toml=0.10.2=pyhd3eb1b0_0
  - tornado=6.2=py38h5eee18b_0
  - wget=1.21.4=h39e8969_0
  - wheel=0.38.4=py38h06a4308_0
  - x264=1!157.20191217=h7b6447c_0
  - xz=5.4.2=h5eee18b_0
  - zipp=3.11.0=py38h06a4308_0
  - zlib=1.2.13=h5eee18b_0
  - zstd=1.5.5=hc292b87_0
```
