## Supplemental Code for "MCHelper automatically curates transposable element libraries across eukaryotic species": MCHelper_Mac.yml

```
name: MCHelper_Mac
channels:
  - defaults
  - bioconda
  - anaconda
  - conda-forge
dependencies:
  - _libgcc_mutex=0.1=main
  - _r-mutex=1.0.1=anacondar_1
  - anyio=4.2.0=pyhd8ed1ab_0
  - aom=3.7.1=h93d8f39_0
  - appnope=0.1.3=pyhd8ed1ab_0
  - argon2-cffi=23.1.0=pyhd8ed1ab_0
  - argon2-cffi-bindings=21.2.0=py312h104f124_4
  - arrow=1.3.0=pyhd8ed1ab_0
  - asttokens=2.4.1=pyhd8ed1ab_0
  - async-lru=2.0.4=pyhd8ed1ab_0
  - attrs=23.2.0=pyh71513ae_0
  - babel=2.14.0=pyhd8ed1ab_0
  - beautifulsoup4=4.12.2=pyha770c72_0
  - binutils_impl_linux-64=2.40=h0b07ce5_0
  - biopython=1.81=py312h41838bb_1
  - blast=2.15.0=pl5321h91c44f7_1
  - bleach=6.1.0=pyhd8ed1ab_0
  - bottleneck=1.3.5=py312h32608ca_0
  - brotli=1.1.0=h0dc2134_1
  - brotli-bin=1.1.0=h0dc2134_1
  - brotli-python=1.1.0=py312heafc425_1
  - bwidget=1.9.14=h694c41f_1
  - bzip2=1.0.8=h10d778d_5
  - c-ares=1.23.0=h10d778d_0
  - ca-certificates=2024.2.2=h8857fd0_0
  - cached-property=1.5.2=hd8ed1ab_1
  - cached_property=1.5.2=pyha770c72_1
  - cairo=1.18.0=h99e66fa_0
  - cctools_osx-64=986=h031c385_0
  - cd-hit=4.8.1=h0100bc9_9
  - certifi=2024.2.2=pyhd8ed1ab_0
  - cffi=1.16.0=py312h38bf5a0_0
  - charset-normalizer=3.3.2=pyhd8ed1ab_0
  - cialign=1.1.0=pyhca03a8a_0
  - clang=17.0.6=default_h7151d67_4
  - clang-17=17.0.6=default_h7151d67_4
  - clang_impl_osx-64=17.0.6=h1af8efd_11
  - clang_osx-64=17.0.6=hb91bd55_11
  - clangxx=17.0.6=default_h7151d67_4
  - clangxx_impl_osx-64=17.0.6=hc3430b7_11
  - clangxx_osx-64=17.0.6=hb91bd55_11
  - comm=0.2.1=pyhd8ed1ab_0
  - compiler-rt=17.0.6=ha38d28d_1
  - compiler-rt_osx-64=17.0.6=ha38d28d_1
  - configargparse=1.7=pyhd8ed1ab_0
  - contourpy=1.2.0=py312hbf0bb39_0
  - curl=8.5.0=h726d00d_0
  - cycler=0.12.1=pyhd8ed1ab_0
  - dav1d=1.2.1=h0dc2134_0
  - debugpy=1.8.0=py312heafc425_1
  - decorator=5.1.1=pyhd8ed1ab_0
  - defusedxml=0.7.1=pyhd8ed1ab_0
  - emboss=6.6.0=h6debe1e_0
  - entrez-direct=16.2=h193322a_1
  - entrypoints=0.4=pyhd8ed1ab_0
  - exceptiongroup=1.2.0=pyhd8ed1ab_0
  - executing=2.0.1=pyhd8ed1ab_0
  - expat=2.5.0=hf0c8a7f_1
  - ffmpeg=6.1.0=gpl_hf5fe730_105
  - font-ttf-dejavu-sans-mono=2.37=hab24e00_0
  - font-ttf-inconsolata=3.000=h77eed37_0
  - font-ttf-source-code-pro=2.038=h77eed37_0
  - font-ttf-ubuntu=0.83=h77eed37_1
  - fontconfig=2.14.2=h5bb23bf_0
  - fonts-conda-ecosystem=1=0
  - fonts-conda-forge=1=0
  - fonttools=4.46.0=py312h41838bb_0
  - fqdn=1.5.1=pyhd8ed1ab_0
  - freetype=2.12.1=h60636b9_2
  - fribidi=1.0.10=hbcb3906_0
  - gawk=5.3.0=h2c496e9_0
  - gettext=0.21.1=h8a4c099_0
  - gfortran_impl_osx-64=12.3.0=hc328e78_3
  - gfortran_osx-64=12.3.0=h18f7dce_1
  - giflib=5.2.1=hb7f2c08_3
  - gmp=6.3.0=h93d8f39_0
  - gnutls=3.7.9=h1951705_0
  - graphite2=1.3.13=h2e338ed_1001
  - harfbuzz=8.3.0=hf45c392_0
  - hdf5=1.14.3=nompi_h691f4bf_100
  - hmmer=3.4=h7133b54_0
  - htop=3.2.2=h908806b_0
  - icu=73.2=hf5e326d_0
  - idna=3.6=pyhd8ed1ab_0
  - importlib-metadata=7.0.1=pyha770c72_0
  - importlib_metadata=7.0.1=hd8ed1ab_0
  - importlib_resources=6.1.1=pyhd8ed1ab_0
  - ipykernel=6.28.0=pyh3cd1d5f_0
  - ipython=8.20.0=pyh707e725_0
  - isl=0.26=imath32_h2e86a7b_101
  - isoduration=20.11.0=pyhd8ed1ab_0
  - jasper=4.1.0=h6ff19ee_0
  - jedi=0.19.1=pyhd8ed1ab_0
  - jinja2=3.1.2=pyhd8ed1ab_1
  - joblib=1.3.2=pyhd8ed1ab_0
  - json5=0.9.14=pyhd8ed1ab_0
  - jsonpointer=2.4=py312hb401068_3
  - jsonschema=4.20.0=pyhd8ed1ab_0
  - jsonschema-specifications=2023.12.1=pyhd8ed1ab_0
  - jsonschema-with-format-nongpl=4.20.0=pyhd8ed1ab_0
  - jupyter-lsp=2.2.1=pyhd8ed1ab_0
  - jupyter_client=8.6.0=pyhd8ed1ab_0
  - jupyter_core=5.5.0=py312hecd8cb5_0
  - jupyter_events=0.9.0=pyhd8ed1ab_0
  - jupyter_server=2.12.2=pyhd8ed1ab_0
  - jupyter_server_terminals=0.5.1=pyhd8ed1ab_0
  - jupyterlab=4.0.10=pyhd8ed1ab_0
  - jupyterlab_pygments=0.3.0=pyhd8ed1ab_0
  - jupyterlab_server=2.25.2=pyhd8ed1ab_0
  - kernel-headers_linux-64=2.6.32=he073ed8_16
  - kiwisolver=1.4.5=py312h49ebfd2_1
  - krb5=1.21.2=hb884880_0
  - lame=3.100=hb7f2c08_1003
  - lcms2=2.16=ha2f27b4_0
  - ld64_osx-64=711=hd3532be_0
  - ld_impl_linux-64=2.40=hc65bd9e_0
  - lerc=4.0.0=hb486fe8_0
  - libabseil=20230802.1=cxx17_h048a20a_0
  - libaec=1.1.2=he965462_1
  - libass=0.17.1=h80904bb_1
  - libblas=3.9.0=20_osx64_openblas
  - libbrotlicommon=1.1.0=h0dc2134_1
  - libbrotlidec=1.1.0=h0dc2134_1
  - libbrotlienc=1.1.0=h0dc2134_1
  - libcblas=3.9.0=20_osx64_openblas
  - libclang-cpp17=17.0.6=default_h7151d67_4
  - libcurl=8.5.0=h726d00d_0
  - libcxx=16.0.6=hd57cbcb_0
  - libdeflate=1.19=ha4e1b8e_0
  - libedit=3.1.20191231=h0678c8f_2
  - libev=4.33=h10d778d_2
  - libexpat=2.5.0=hf0c8a7f_1
  - libffi=3.4.2=h0d85af4_5
  - libgd=2.3.3=h0dceb68_9
  - libgfortran=5.0.0=13_2_0_h97931a8_1
  - libgfortran-devel_osx-64=12.3.0=h0b6f5ec_3
  - libgfortran-ng=3.0.1=h93005f0_2
  - libgfortran5=13.2.0=h2873a65_1
  - libglib=2.78.3=h198397b_0
  - libiconv=1.17=hd75f5a5_1
  - libidn2=2.3.4=hb7f2c08_0
  - libjpeg-turbo=3.0.0=h0dc2134_1
  - liblapack=3.9.0=20_osx64_openblas
  - liblapacke=3.9.0=20_osx64_openblas
  - libllvm17=17.0.6=hbedff68_1
  - libnghttp2=1.58.0=h64cf6d3_1
  - libopenblas=0.3.25=openmp_hfef2a42_0
  - libopencv=4.8.1=py312h989985c_5
  - libopenvino=2023.1.0=h93d8f39_2
  - libopenvino-auto-batch-plugin=2023.1.0=h93d8f39_2
  - libopenvino-auto-plugin=2023.1.0=h93d8f39_2
  - libopenvino-hetero-plugin=2023.1.0=h93d8f39_2
  - libopenvino-intel-cpu-plugin=2023.1.0=h93d8f39_2
  - libopenvino-ir-frontend=2023.1.0=h93d8f39_2
  - libopenvino-onnx-frontend=2023.1.0=h93d8f39_2
  - libopenvino-paddle-frontend=2023.1.0=h93d8f39_2
  - libopenvino-pytorch-frontend=2023.1.0=h93d8f39_2
  - libopenvino-tensorflow-frontend=2023.1.0=h93d8f39_2
  - libopenvino-tensorflow-lite-frontend=2023.1.0=h93d8f39_2
  - libopus=1.3.1=hc929b4f_1
  - libpng=1.6.39=ha978bb4_0
  - libprotobuf=4.24.4=h0ee05dc_0
  - libsodium=1.0.18=hbcb3906_1
  - libsqlite=3.44.2=h92b6c6a_0
  - libssh2=1.11.0=hd019ec5_0
  - libtasn1=4.19.0=hb7f2c08_0
  - libtiff=4.6.0=h684deea_2
  - libunistring=0.9.10=h0d85af4_0
  - libvpx=1.13.1=he965462_0
  - libwebp=1.3.2=h44782d1_1
  - libwebp-base=1.3.2=h0dc2134_0
  - libxcb=1.15=hb7f2c08_0
  - libxml2=2.12.3=hc0ae0f7_0
  - libzlib=1.2.13=h8a1eda9_5
  - llvm-openmp=17.0.6=hb6ac08f_0
  - llvm-tools=17.0.6=hbedff68_1
  - mafft=7.520=h2413b67_3
  - make=4.3=h22f3db7_1
  - markupsafe=2.1.3=py312h104f124_1
  - matplotlib=3.8.2=py312hb401068_0
  - matplotlib-base=3.8.2=py312h302682c_0
  - matplotlib-inline=0.1.6=pyhd8ed1ab_0
  - mistune=3.0.2=pyhd8ed1ab_0
  - mpc=1.3.1=h81bd1dd_0
  - mpfr=4.2.1=h0c69b56_0
  - munkres=1.1.4=pyh9f0ad1d_0
  - nbclient=0.8.0=pyhd8ed1ab_0
  - nbconvert-core=7.14.0=pyhd8ed1ab_0
  - nbformat=5.9.2=pyhd8ed1ab_0
  - ncbi-vdb=3.0.0=pl5321h9722bc1_0
  - ncurses=6.4=h93d8f39_2
  - nest-asyncio=1.5.8=pyhd8ed1ab_0
  - nettle=3.9.1=h8e11ae5_0
  - notebook-shim=0.2.3=pyhd8ed1ab_0
  - nspr=4.35=hea0b92c_0
  - nss=3.95=hfeb00ea_0
  - numexpr=2.8.7=py312haf8ecfc_4
  - numpy=1.26.2=py312hfd3bce2_0
  - opencv=4.8.1=py312hbdb8196_5
  - openh264=2.4.0=h93d8f39_0
  - openjpeg=2.5.0=ha4da562_3
  - openssl=3.2.1=hd75f5a5_1
  - overrides=7.4.0=pyhd8ed1ab_0
  - p11-kit=0.24.1=h65f8906_0
  - packaging=23.2=pyhd8ed1ab_0
  - pandas=2.1.1=py312he282a81_0
  - pandocfilters=1.5.0=pyhd8ed1ab_0
  - pango=1.50.14=h19c1c8a_2
  - parso=0.8.3=pyhd8ed1ab_0
  - pcre=8.45=he49afe7_0
  - pcre2=10.42=h0ad2156_0
  - pdf2image=1.16.3=pyhd8ed1ab_1
  - perl=5.32.1=4_h0dc2134_perl5
  - perl-archive-tar=2.40=pl5321hdfd78af_0
  - perl-carp=1.38=pl5321hdfd78af_4
  - perl-common-sense=3.75=pl5321hdfd78af_0
  - perl-compress-raw-bzip2=2.201=pl5321h9722bc1_0
  - perl-compress-raw-zlib=2.105=pl5321h9722bc1_0
  - perl-encode=3.19=pl5321ha5712d3_1
  - perl-exporter=5.72=pl5321hdfd78af_2
  - perl-exporter-tiny=1.002002=pl5321hdfd78af_0
  - perl-extutils-makemaker=7.70=pl5321hd8ed1ab_0
  - perl-io-compress=2.201=pl5321h7133b54_2
  - perl-io-zlib=1.14=pl5321hdfd78af_0
  - perl-json=4.10=pl5321hdfd78af_0
  - perl-json-xs=2.34=pl5321hcd10b59_5
  - perl-list-moreutils=0.430=pl5321hdfd78af_0
  - perl-list-moreutils-xs=0.430=pl5321ha5712d3_1
  - perl-parent=0.236=pl5321hdfd78af_2
  - perl-pathtools=3.75=pl5321ha5712d3_3
  - perl-scalar-list-utils=1.62=pl5321ha5712d3_0
  - perl-types-serialiser=1.01=pl5321hdfd78af_0
  - pexpect=4.8.0=pyh1a96a4e_2
  - pickleshare=0.7.5=py_1003
  - pillow=10.1.0=py312h0c70c2f_0
  - pip=23.3.1=pyhd8ed1ab_0
  - pixman=0.42.2=he965462_0
  - pkgutil-resolve-name=1.3.10=pyhd8ed1ab_1
  - platformdirs=4.1.0=pyhd8ed1ab_0
  - poppler=23.12.0=hdd5a5e8_0
  - poppler-data=0.4.12=hd8ed1ab_0
  - prometheus_client=0.19.0=pyhd8ed1ab_0
  - prompt-toolkit=3.0.42=pyha770c72_0
  - psutil=5.9.5=py312h104f124_1
  - pthread-stubs=0.4=hc929b4f_1001
  - ptyprocess=0.7.0=pyhd3deb0d_0
  - pugixml=1.14=he965462_0
  - pure_eval=0.2.2=pyhd8ed1ab_0
  - py-opencv=4.8.1=py312h128fb24_5
  - pycparser=2.21=pyhd8ed1ab_0
  - pygments=2.17.2=pyhd8ed1ab_0
  - pyobjc-core=10.1=py312h74abf1d_0
  - pyobjc-framework-cocoa=10.1=py312h74abf1d_0
  - pyparsing=3.1.1=pyhd8ed1ab_0
  - pysocks=1.7.1=pyha2e5f31_6
  - python=3.12.0=h30d4d87_0_cpython
  - python-dateutil=2.8.2=pyhd8ed1ab_0
  - python-fastjsonschema=2.19.1=pyhd8ed1ab_0
  - python-json-logger=2.0.7=pyhd8ed1ab_0
  - python-tzdata=2023.3=pyhd3eb1b0_0
  - python_abi=3.12=4_cp312
  - pytz=2023.3.post1=py312hecd8cb5_0
  - pyyaml=6.0.1=py312h104f124_1
  - pyzmq=25.1.2=py312hc789acb_0
  - r-base=4.3.2=h46d3641_2
  - readline=8.2=h9e318b2_1
  - referencing=0.32.1=pyhd8ed1ab_0
  - requests=2.31.0=pyhd8ed1ab_0
  - rfc3339-validator=0.1.4=pyhd8ed1ab_0
  - rfc3986-validator=0.1.1=pyh9f0ad1d_0
  - rpds-py=0.16.2=py312h6e28e45_0
  - scikit-learn=1.3.2=py312h5e37b4f_2
  - scipy=1.11.4=py312heccc6a5_0
  - send2trash=1.8.2=pyhd1c38e8_0
  - setuptools=68.2.2=pyhd8ed1ab_0
  - sigtool=0.1.3=h88f4db0_0
  - six=1.16.0=pyh6c4a22f_0
  - snappy=1.1.10=h225ccf5_0
  - sniffio=1.3.0=pyhd8ed1ab_0
  - soupsieve=2.5=pyhd8ed1ab_1
  - sqlite=3.44.2=h7461747_0
  - stack_data=0.6.2=pyhd8ed1ab_0
  - svt-av1=1.8.0=h93d8f39_0
  - sysroot_linux-64=2.12=he073ed8_16
  - tapi=1100.0.11=h9ce4665_0
  - tbb=2021.10.0=h1c7c39f_2
  - terminado=0.18.0=pyh31c8845_0
  - threadpoolctl=3.2.0=pyha21a80b_0
  - tinycss2=1.2.1=pyhd8ed1ab_0
  - tk=8.6.13=h1abcd95_1
  - tktable=2.10=ha166976_5
  - tomli=2.0.1=pyhd8ed1ab_0
  - tornado=6.3.3=py312h104f124_1
  - traitlets=5.14.1=pyhd8ed1ab_0
  - types-python-dateutil=2.8.19.20240106=pyhd8ed1ab_0
  - typing-extensions=4.9.0=hd8ed1ab_0
  - typing_extensions=4.9.0=pyha770c72_0
  - typing_utils=0.1.0=pyhd8ed1ab_0
  - tzdata=2023c=h71feb2d_0
  - uri-template=1.3.0=pyhd8ed1ab_0
  - urllib3=2.1.0=pyhd8ed1ab_0
  - wcwidth=0.2.13=pyhd8ed1ab_0
  - webcolors=1.13=pyhd8ed1ab_0
  - webencodings=0.5.1=pyhd8ed1ab_2
  - websocket-client=1.7.0=pyhd8ed1ab_0
  - wget=1.20.3=hd3787cc_1
  - wheel=0.42.0=pyhd8ed1ab_0
  - x264=1!164.3095=h775f41a_2
  - x265=3.5=hbb4e6a2_3
  - xorg-libxau=1.0.11=h0dc2134_0
  - xorg-libxdmcp=1.1.3=h35c211d_0
  - xz=5.2.6=h775f41a_0
  - yaml=0.2.5=h0d85af4_2
  - zeromq=4.3.5=h93d8f39_0
  - zipp=3.17.0=pyhd8ed1ab_0
  - zlib=1.2.13=h8a1eda9_5
  - zstd=1.5.5=h829000d_0
variables:
  CONDA_SUBDIR: osx-64
```
