## Supplemental Code for "MCHelper automatically curates transposable element libraries across eukaryotic species": README.md

# MCHelper
MCHelper: An automatic tool to curate transposable element libraries

## Table of Contents  
* [Introduction](#introduction)
* [Installation](#installation)
  * [Linux/Windows](#linuxwindows)
  * [MacOS](#MacOS)
* [Testing](#testing)
* [Usage](#usage)
* [Inputs](#inputs)
* [Outputs](#outputs)
* [Citation](#citation)


## Introduction
The number of species with high quality genome sequences continues to increase, in part due to scaling up of multiple large scale biodiversity sequencing projects. While the need to annotate genic sequences in these genomes is widely acknowledged, the parallel need to annotate transposable element sequences that have been shown to alter genome architecture, rewire gene regulatory networks, and contribute to the evolution of host traits is becoming ever more evident. However, accurate genome-wide annotation of transposable element sequences is still technically challenging. Several de novo transposable element identification tools are now available, but manual curation of the libraries produced by these tools is needed to generate high quality genome annotations. Manual curation is time-consuming, and thus impractical for largescale genomic studies, and lacks reproducibility. In this work, we present the Manual Curator Helper tool, MCHelper, which automates the TE library curation process. By leveraging MCHelper's fully automated mode with the outputs from two de novo transposable element identification tools, RepeatModeler2 and REPET, in fruit fly, rice, and zebrafish, we show a substantial improvement in the quality of the transposable element libraries and genome annotations. MCHelper libraries are less redundant, with up to 54% reduction in the number of consensus sequences, have up to 11.4% fewer false positive sequences, and also have up to ~45% fewer “unclassified/unknown” transposable element consensus sequences. Genome-wide transposable element annotations were also improved, including larger unfragmented insertions.

<p align="center">
  <img src="https://github.com/GonzalezLab/MCHelper/blob/main/MCHelper_Flow.jpg" alt="MCHelper Flowchart" />
</p>

## Installation
### Linux/Windows
For Windows Systems is necessarity to have a functional installation of Windows Subsystem for Linux (WSL) version 2, the Poppler package installed (sudo apt-get install poppler-utils), as well as the QT package (sudo apt-get install qtbase5-dev).

It is recommended to install the dependencies in an Anaconda environment. 

```
git clone https://github.com/gonzalezlab/MCHelper.git
```

Then, locate the MCHelper folder and find the file named "MCHelper.yml". Then, install the environment: 

```
conda env create -f MCHelper/MCHelper.yml
```

Now, unzip all the databases needed by MCHelper:

```
cd MCHelper/db
unzip '*.zip'
conda activate MCHelper
makeblastdb -in allDatabases.clustered_rename.fa -dbtype nucl
```

Then, download the pfam database released by REPET group and renamed it:

```
wget https://urgi.versailles.inrae.fr/download/repet/profiles/ProfilesBankForREPET_Pfam35.0_GypsyDB.hmm.tar.gz

tar xvf ProfilesBankForREPET_Pfam35.0_GypsyDB.hmm.tar.gz
mv ProfilesBankForREPET_Pfam35.0_GypsyDB.hmm Pfam35.0.hmm
```

And that's it. You have now installed MCHelper.

### MacOS 
These installation instructions have been tested on MacOS with M1/M2 architectures (Apple Silicon, arch: arm64). Therefore, these instructions are not compatible with MacOS with Intel Core processors).

Set up Rosetta. 
* Download and install the iTerm (or duplicate it if you have already installed, then rename it to, for example, iTerm_X86_64).
* Right click on the icon iTerm (or iTerm_X86_64 if you renamed it), and select the option Get Info, and check box: Open using Rosetta
* Open the new terminal iTerm (or iTerm_X86_64)
* Verify the architecture: uname -m. It should appear: x86_64

Install Mambaforge
Using the same iTerm (or iTerm_X86_64) we configured earlier, download the Mambaforge script and install it:
```
wget https://github.com/conda-forge/miniforge/releases/download/23.11.0-0/Mambaforge-23.11.0-0-MacOSX-x86_64.sh 
chmod +x Mambaforge-23.11.0-0-MacOSX-x86_64.sh
```
```
./Mambaforge-23.11.0-0-MacOSX-x86_64.sh
```
Follow the prompts, install, and initialize conda.

Install MCHelper conda environment using the special YML file for Mac (MCHelper_Mac.yml), using the Rosetta iTerm (or iTerm_X86_64):
```
git clone https://github.com/gonzalezlab/MCHelper.git
```
```
conda env create -f MCHelper/MCHelper_Mac.yml
```
```
conda activate MCHelper_Mac
```

Download and rename the TRF binary for MacOS:
```
cd MCHelper/tools
wget https://github.com/Benson-Genomics-Lab/TRF/releases/download/v4.09.1/trf409.macosx
```
```
rm -f trf409.linux64
mv trf409.macosx trf409.linux64
chmod +x trf409.linux64
cd -
```

Now, unzip all the databases needed by MCHelper:

```
cd MCHelper/db
unzip '*.zip'
makeblastdb -in allDatabases.clustered_rename.fa -dbtype nucl
```

Then, download the pfam database released by REPET group and renamed it:

```
wget https://urgi.versailles.inrae.fr/download/repet/profiles/ProfilesBankForREPET_Pfam35.0_GypsyDB.hmm.tar.gz

tar xvf ProfilesBankForREPET_Pfam35.0_GypsyDB.hmm.tar.gz
mv ProfilesBankForREPET_Pfam35.0_GypsyDB.hmm Pfam35.0.hmm
```

And that's it. You have now installed MCHelper.

## Testing
To test MCHelper, we provide some example inputs and also the expected results (located at Test_dir/) to allow you to compare with your own outputs. To check MCHelper is running properly, you can do:

First, activate the anaconda enviroment, if it isn't activated yet:

```
conda activate MCHelper
```

Then, be sure you are in the main folder (this one where MCHelper.py is located) and unzip the D. melanogaster genome:

```
unzip Test_dir/repet_input/Dmel_genome.zip -d Test_dir/repet_input/
```

Next step is download and format the host genes from BUSCO

```
wget https://busco-data.ezlab.org/v4/data/lineages/diptera_odb10.2020-08-05.tar.gz
mv diptera_odb10.2020-08-05.tar.gz Test_dir/repet_input/ 
cd Test_dir/repet_input/
tar xvf diptera_odb10.2020-08-05.tar.gz
cat diptera_odb10/hmms/*.hmm > diptera_odb10.hmm
cd -
```

Now, run the MCHelper script:

```
mkdir Test_dir/repet_output_own

python3 MCHelper.py -r A -t 8 -i Test_dir/repet_input/ -o Test_dir/repet_output_own -g Test_dir/repet_input/Dmel_genome.fasta --input_type repet -b Test_dir/repet_input/diptera_odb10.hmm -a F -n Dmel
```

This test will take the REPET's output and will do the curation automatically, using most of the parameters by default.
If you want to run the test for a toy library with less sequences, you can execute:

```
unzip Test_dir/fasta_input/Dmel_genome.zip -d Test_dir/fasta_input/

mkdir Test_dir/fasta_output_own

python3 MCHelper.py -r A -t 8 -l Test_dir/fasta_input/TE_lib_toy.fa -o Test_dir/fasta_output_own -g Test_dir/fasta_input/Dmel_genome.fna --input_type fasta -b Test_dir/repet_input/diptera_odb10.hmm -a F
```

## Usage
Be sure you have activated the anaconda environment:

```
conda activate MCHelper
```

Then, execute MCHelper with default parameters. For REPET input (see [Testing](#testing) for a practical example):

```
python3 MCHelper.py -i path/to/repet_output -o path/to/MCHelper_output -g path/to/genome -n repet_name_project --input_type repet -b path/to/reference_genes.hmm -a F
```

For fasta input:

```
python3 MCHelper.py -l path/to/TE_library_in_fasta -o path/to/MCHelper_output -g path/to/genome --input_type fasta -b path/to/reference_genes.hmm -a F
```

To see the full help documentation run:

```
python3 MCHelper.py --help
```

Full list of parameters include:
* -h, --help            show this help message and exit
* -r MODULE, --module MODULE:  module of curation [A, C, U, T, E, M]. Required*
* -i INPUT_DIR, --input INPUT_DIR:  Directory with the files required to do the curation (REPET output directory). Required*
* -g GENOME, --genome GENOME: Genome used to detect the TEs. Required*
* -o OUTPUTDIR, --output OUTPUTDIR: Path to the output directory. Required*
* --te_aid TE_AID:       Do you want to use TE-aid? [Y or N]. Default=Y.
* -a AUTOMATIC:          Level of automation: F: fully automated, S: semi-automated, M: fully manual?. Default=F.
* -n PROJ_NAME:          REPET project name. Required for repet input*
* -t CORES:              cores to execute some steps in parallel. Default=all available cores.
* -m REF_LIBRARY_UNCLASSIFIED_MODULE: Path to the sequences to be used as references in the unclassified module.
* -v VERBOSE            Verbose? [Y or N]. Default=N.
* --input_type INPUT_TYPE:  Input type: fasta or REPET.
* -l USER_LIBRARY:       User defined library to be used with input type fasta.
* -b BUSCO_LIBRARY:      Reference/BUSCO genes to filter out TEs (HMM format required).
* -z MINBLASTHITS       Minimum number of blast hits to process an element.
* -c MINFULLLFRAGMENTS:   Minimum number of full-length fragments to process an element. Default=1
* -s PERC_SSR:           Maximum length covered by single repetitions (in percentage between 0-100) allowed for a TE not to be removed. Default=60.
* -e EXT_NUCL           Number of nucleotides to extend each size of the element. Default=500.
* -x NUM_ITE            Number of iterations to extend the elements. Default=16.
* -k clust_algorithm            Clustering algorithm: cd-hit or meshclust. Default=cd-hit
* --version             show program's version number and exit.

MCHelper can be run in three different modes: Fully automatic (F), semi-automatic (S) and manual (M). The way you can control this is with the parameter **-a [F,S or M]**. Notice that the fully automatic mode will make all the decision by you and, at the end, will generate different outputs curated and non-curated sequences. In contrast, the semi-automatic mode runs the structural check and allows the user to inspect the consensus sequences that do not fit the structural requirements. The manual mode does not run the structural check and sends all the consensus sequences to manual inspection. 

MCHelper is a modular pipeline (see figure below), which can be run in a integrated way or module by module. You can control this with the -r or --module parameter, indicating which of the four modules you want to run. **If you want to run the whole pipeline, select -r A**. Otherwise, if you want just run one of them, select the letter corresponding to the module: consensus extension module=E (Figure A), Manual Inpection module=M (Figure B), and TE classification module=U (Figure C). You can also only run TE-Aid in parallel using the parameter -r T. 

<p align="center">
  <img src="https://github.com/GonzalezLab/MCHelper/blob/main/MCHelper_modules_Flow.jpg"/>
</p>

## Inputs
The input files required by MCHelper will depend of the tool you used to create the TE library. **If you used REPET**, then you will need the following files:

* the genome assembly 
* the library created by the TEdenovo pipeline. This library is named as projName_refTEs.fa, where projName is the name of your own REPET project. 
* the table with features created by PASTEC and is normally named projName_denovoLibTEs_PC.classif. Again, projName is the name of your own REPET project.
* a folder containing coverage plots created with the REPET tool "plotCoverage.py". This folder must be named "plotCoverage" and must be placed in the input folder specified in the -i parameter.
* a folder containing the gff files generated by the REPET tool "CreateGFF3sForClassifFeatures.py". This folder must be named "gff_reversed" and must be placed in the input folder specified in the -i parameter. 

*If you used any other tool that generates TE libraries in fasta format*, then you will need the following:
* the genome assembly 
* the library created by the tool. 

In the lastest case, MCHelper will find the information required to do the curation process. This information include: 
* How many full length copies and fragment has each consensus
* structural features such as terminal repeats, and coding domains
* BLASTn, BLASTx, and tBLASTx with TE databases

## Outputs
Outputs generated by MCHelper depend on the modules chosen to run. The tool will create an independent folder for the Classified and Unclassified modules. Inside, it will save some temporal as well as final files. The final processed sequences will be stored at the file named "curated_sequences_NR.fa" (Non Redundant version) and "curated_sequences_R.fa" (Redundant version).

The rest of the files are the following:
* ClassifiedModule folder
  * MSA_Plots: Folder containing the MSA graphs generated by CIAlign (F and S modes).
  * MSA_seeds: Folder containing the MSA files generated by CIAlign (F and S modes). Those files can be used to visualize the MSA and also to construct HMMs.
  * te_aid: Folder containing the TE+Aid plots generated. They are used by MCHelper in the manual inspection step (S and M modes), but also can be usefull for checking a specific TE by the user.
  * cons_flf.fa: File containing only the TEs with more than a certain threhold of full length fragments in the genome. This threshold is handle by the variable -c. A copy is considered as full length fragment when it covers at least 94% of the consensus length.
  * denovoLibTEs_PC.classif: Tabular file containing coding and structural information of each consensus.
  * fullLengthFrag.txt: Tabular file containing information about number of number of fragments and full length fragments.
  * input_to_unclassified_module_seqs.fa: Sequences that will be used in the Unclassified module (only when -r A is selected).
  * kept_seqs_classified_module_curated.fa: File containing the sequences considered that beloging a complete TEs.
  * kept_seqs_classified_module_non_curated.fa: File containing the sequences considered as incomplete, fragmented or that doesn't satisfy the structural conditions.
  * kept_seqs_classified_module.fa: File containing the sequences that have been kept in the module. It is a merge between the two previous described files (kept_seqs_classified_module_curated.fa and kept_seqs_classified_module_non_curated.fa). At the end, MCHelper will join this file with kept_seqs_unclassified_module.fa to create the curated_sequences_R.fa final file.

* UnclassifiedModule folder
  *  MSA_plots: Folder containing the MSA graphs generated by CIAlign.
  *  MSA_seeds: Folder containing the MSA files generated by CIAlign. Those files can be used to visualize the MSA and also to construct HMMs.
  *  cons_flf.fa: File containing only the TEs with more than a certain threhold of full length fragments in the genome. This threshold is handle by the variable -c. A copy is considered as full length fragment when it covers at least 94% of the consensus length.
  *  denovoLibTEs_PC.classif: Tabular file containing coding and structural information of each consensus.
  *  extended_cons.fa; File containing the extended TE.
  *  kept_seqs_unclassified_module.fa: File containing the sequences that have been kept in the module. At the end, MCHelper will join this file with kept_seqs_classified_module.fa to create the curated_sequences_R.fa final file.
  *  new_user_lib.fa: Intermidiate file containing fomated sequences needed to run with MCHelper.

## Citation
if you use this software, please cite us as following:
Orozco-Arias, S., Sierra, P., Durbin, R., Gonzalez, J. (2023). MCHelper automatically curates transposable element libraries across species. https://doi.org/10.1101/2023.10.17.562682. bioRxiv.
