## Supplemental Code for "MCHelper automatically curates transposable element libraries across eukaryotic species": blastndotplot.R

```
# self sequence db is being made by the shell script // or you need to make it to use in R
blastdotplot=function(query = NULL, db = NULL, blast = NULL, os = NULL, tables = NULL, output = NULL){
  
  # run the selfblast
  bl=read.table(text=system(paste("blastn -query", query, "-db", db, "-evalue 0.05 -outfmt 6 -word_size 11 -gapopen 5 -gapextend 2 -reward 2 -penalty -3 | cut -f 1,7-10 | sed 's/#/-/g'"),
                            intern = TRUE)
                )
  # order from left to right
  bl=bl[order(bl$V2, decreasing = F),]
  
  # test if there are orf detected; will later store in orf if TRUE
  test<-try(read.table(as.character(blast)), T)
  # test if there are TE prot detected; will later store in prot if TRUE
  #test2<-try(read.table(as.character(blast)), T)

  if(class(test) == "data.frame"){
     orfs=read.table(as.character(blast))
     } else {
     print("no orf to plot...")
     orfs <- suppressWarnings(as.data.frame(0))
     names(orfs)<-"V1"
   }

  # if(class(test2) == "data.frame"){
  #    orfs=read.table(as.character(getorf))
  #    }else{
  #    suppressWarnings(prot$V1 <- as.data.frame(c(0,0))[,1])
  #  }
  ###########################
  ## dot-plot (bottom left)##
  ###########################

  plot(x = 1, type = "n", xlim = c(0,bl$V3[1]),ylim = c(0,bl$V3[1]), col = "white",
       main = "TE consensus self dotplot (blastn)",
       ylab = paste(as.character(bl$V1[1]), "(bp)", sep = " "),
       xlab = paste(as.character(bl$V1[1]), "(bp)", sep = " ")
       )
    for(i in 1:length(bl$V1)){
      if(bl$V5[i] > bl$V4[i]){
        segments(x0 = bl$V2[i], x1 = bl$V3[i], y0 = bl$V4[i], y1 = bl$V5[i], col = "black", lwd = 1.5)
      } else {
        segments(x0 = bl$V2[i], x1 = bl$V3[i], y0 = bl$V4[i], y1 = bl$V5[i], col = "#009E73", lwd = 1.5)  
      }
        # if orientation
    } # for each segment end
  
  #####################################
  ## Annotation graph (bottom right) ##
  #####################################
  
  ## Arrows layer ##

  plot(x = 1, type = "n", xlim = c(0,bl$V3[1]),ylim = c(-max(length(orfs$V1),10),length(bl$V1)), col = "white", yaxt="n",
       main = "TE consensus structure and protein hits",
       xlab = paste(as.character(bl$V1[1]), "(bp)", sep = " "),
       ylab = ""
  )
  for(i in seq(1:length(bl$V1))){
    if(bl$V4[i] > bl$V2[i]){
        arrows(x0 = bl$V2[i], x1 = bl$V3[i], y0 = i, y1 = i, col = rainbow(length(bl$V1))[i], lwd = 3, length = 0.1)
        arrows(x0 = bl$V4[i], x1 = bl$V5[i], y0 = i, y1 = i, col = rainbow(length(bl$V1))[i], lwd = 3, length = 0.1)
    } # if to draw (filter)
  } # for each segment end


  ## Orfs layers ##
  if(class(test) != "data.frame"){ # if orf table is empty
     text(paste("no orf >",os," bp detected", sep=""), x=bl$V3[1]/2, y=-5, cex = 2)
    } else { # if ORF table, plot orfs
      for(i in seq(1:length(orfs$V1))){
        if(orfs$V1[i] < orfs$V2[i]){ # checking ORF orientation
          rect(xleft = orfs$V1[i], xright = orfs$V2[i], # draw a + ORF
               ybottom = -i-0.15, ytop = -i+0.15, lwd = 1, border = "black")
        } else {
          rect(xleft = orfs$V1[i], xright = orfs$V2[i], # draw a - ORF
               ybottom = -i-0.15, ytop = -i+0.15, lwd = 1, border = "red")  
        } # orientation

          ## TE protein hits (blastp) ##
          rect(xleft = orfs$V5[i], xright = orfs$V6[i], 
               ybottom = -i-0.15, ytop = -i+0.15, lwd = 1, col = as.character(paste("#",orfs$V8[i], sep="")), border = "white") # draw colored rectangle same way as orf
          text(paste(orfs$V3[i], orfs$V4[i]), x = (min(orfs$V1[i],orfs$V2[i])+max(orfs$V1[i],orfs$V2[i]))/2, y = -i+0.15, pos = 3) # print hit name
      
        } # for each segment
      #names(orfs)<-c("orf.start", "orf.end", "hit.TE.prot", "TE.Class", "hit.start", "hit.end", "strand", "color")
      #print(orfs)
  } # if orf present plot orfs and prot hits
    
  #print(tables)
  if(tables == "TRUE"){
    names(bl)<-c("TE", "from.1", "to.1", "from.2", "to.2")
    write.table(bl, file = paste(output, "/TE.self-blast.txt", sep = ""), quote = F, row.names = F)
  } # if tables to print
  
} # function end
```
