## Supplemental Code for "MCHelper automatically curates transposable element libraries across eukaryotic species": consensus2genome.R

```
#! /bin/Rscript
#######################################################################################
### consensus2genome - v2 - Clement Goubert (2020) -      ###
### ------------------------------------------------------------------------------- ###
### This R function blast a TE consensus against a reference genome and then plots  ###
### the genomic fragments found relative to the consensus sequence                  ###
### see https://github.com/clemgoub/consensus2genome for the full documentation     ###
### to use, copy and paste the following code into a R console                      ###
### USAGE: consensus2genome(query, db , ...)                                        ###
### query: path to query (fasta file)                                               ###
### db: path to blast db (blast formated nucleotide database)                       ###
#######################################################################################
# Changelog V2 --> V3 | 04.19.2021
# - Fit for shell wrapper, remove regions highlight from usused trim script
# Changelog V1 --> V2 | 03.13.2020
# - Add a second graph with suggested cut points
# 
# V2=alpha | not for release
# TO FIX: trace the coverage on the left graph
# TO DO: convert breakpoints to bed for getfasta


consensus2genome=function(query=NULL, db=NULL, evalue=10e-8, FL_thresh=0.9, alpha=0.3, full_alpha=1, auto_y=T, bins=NULL){
  if(is.null(query)){print('query not specified')}
  if(is.null(db)){print('db not specified')}
  #perform the blast
  blast=read.table(text=system(paste("blastn -query", query, "-db", db , "-evalue", evalue, "-outfmt 6 | sed 's/#/-/g'"), intern = TRUE))
  write.table(blast, file = "blastn.txt",  quote = F, row.names = F)
  #TE consensus size
  cons_len=as.numeric(system(paste(bins,"/getlength.sh ",query, sep = ""), intern = TRUE))
  print(paste("consensus length: ", cons_len, "bp", sep = " "))
  #list of almost full length fragments
  full=blast[abs(blast$V7-blast$V8) >= FL_thresh*as.numeric(cons_len),]
  #graph
  if(auto_y == T){
    #par(mar=c(2,2,2,2))
    plot(range(0, cons_len), range(0, max(100-blast$V3)), type = "n", main=paste("TE: ", as.character(blast[1,1]), "\n consensus size: ", as.character(cons_len), "bp; fragments: ", as.character(length(blast$V1)), "; full length: ", as.character(length(full$V1))," (>=",as.character(as.numeric(FL_thresh)*cons_len),"bp)", sep = ""), cex.main = 0.9, xlab = "TE consensus (bp)", ylab = "divergence to consensus (%)")
    for(i in 1:length(blast$V1)){
      segments(blast$V7[i], 100-blast$V3[i], blast$V8[i], 100-blast$V3[i], col=rgb(0,0,0,alpha=alpha))
    }
    for(i in 1:length(blast$V1)){
      segments(full$V7[i], 100-full$V3[i], full$V8[i], 100-full$V3[i], col=rgb(1,0,0, alpha=full_alpha), lwd=1.5)
    }}
  else{
    #par(mar=c(2,2,2,2))
    plot(range(0, cons_len), range(0, auto_y), type = "n", main=paste("TE: ", as.character(blast[1,1]), "\n consensus size: ", as.character(cons_len), "bp; fragments: ", as.character(length(blast$V1)), "; full length: ", as.character(length(full$V1))," (>=",as.character(as.numeric(FL_thresh)*cons_len),"bp)", sep = ""), cex.main = 0.9, xlab = "TE consensus (bp)", ylab = "divergence to consensus (%)")
    for(i in 1:length(blast$V1)){
      segments(blast$V7[i], 100-blast$V3[i], blast$V8[i], 100-blast$V3[i], col=rgb(0,0,0,alpha=alpha))
    }
    for(i in 1:length(blast$V1)){
      segments(full$V7[i], 100-full$V3[i], full$V8[i], 100-full$V3[i], col=rgb(1,0,0, alpha=full_alpha), lwd=1.5)
    }}
  
#make the coverage matrix and graph
coverage=matrix(rep(FALSE, length(blast$V1)*as.numeric(cons_len)), byrow = T, ncol = as.numeric(cons_len))
for(i in 1:length(blast$V1)){
    coverage[i,]<-c(rep(FALSE,blast$V7[i]-1),rep(TRUE,abs(blast$V8[i]-blast$V7[i])+1), rep(FALSE,as.numeric(cons_len)-blast$V8[i]))
}

    # TO FIX: trace the coverage on the left graph
    #points(colSums(coverage), type='l', axes = F, ylab = NA, xlab = NA, col=covcol, ylim = c(0, max(colSums(coverage))))
    #axis(side = 4)
    #mtext(side = 4, line = 3, 'consensus coverage (bp)')
    
    ## import removator
    
    removator<-function(covM){
      as.data.frame(covM)->covMT
      covMT$bp=rownames(covMT)
      plot(covM, type = "l", main = "TE consensus genomic coverage", xlab = "TE consensus (bp)", ylab = "coverage (bp)")
     } # removator function
    
    removator(colSums(coverage)) # makes the second graph
}# whole funtion
```
