## Supplemental Code for "MCHelper automatically curates transposable element libraries across eukaryotic species": consensus2genome_DEV.R

```
#! /bin/Rscript
#######################################################################################
### consensus2genome - v1.1 - Clement Goubert (2017) -    ###
### ------------------------------------------------------------------------------- ###
### This R function blast a TE consensus against a reference genome and then plots  ###
### the genomic fragments found relative to the consensus sequence                  ###
### see https://github.com/clemgoub/consensus2genome for the full documentation     ###
### to use, copy and paste the following code into a R console                      ###
### USAGE: consensus2genome(query, db , ...)                                        ###
### query: path to query (fasta file)                                               ###
### db: path to blast db (blast formated nucleotide database)                       ###
#######################################################################################

consensus2genome=function(query=NULL, db=NULL, evalue=10e-8, FL_thresh=0.9, alpha=0.3, full_alpha=1, auto_y=T, cover=T, covcol="blue"){
  if(is.null(query)){print('query not specified')}
  if(is.null(db)){print('db not specified')}
  #perform the blast
  blast=read.table(text=system(paste("blastn -query", query, "-db", db , "-evalue", evalue, "-outfmt 6 | sed 's/#/-/g'"), intern = TRUE))
  #TE consensus size
  cons_len=system(paste("grep -v '>' ",query ," | wc | awk '{print $3-$1}'"), intern = TRUE)
  #list of almost full length fragments
  full=blast[abs(blast$V7-blast$V8) >= FL_thresh*as.numeric(cons_len),]
  #graph
  if(auto_y == T){
    par(mar=c(5,5,5,5))
    plot(range(0, cons_len), range(0, max(100-blast$V3)), type = "n", main=paste("TE: ", as.character(blast[1,1]), "\n consensus size: ", as.character(cons_len), "bp; fragments: ", as.character(length(blast$V1)), "; full length: ", as.character(length(full$V1))," (>=",as.character(as.numeric(FL_thresh)*100),"%)", sep = ""), cex.main = 0.9, xlab = "TE consensus (bp)", ylab = "divergence to consensus (%)")
    for(i in 1:length(blast$V1)){
      segments(blast$V7[i], 100-blast$V3[i], blast$V8[i], 100-blast$V3[i], col=rgb(0,0,0,alpha=alpha))
    }
    for(i in 1:length(blast$V1)){
      segments(full$V7[i], 100-full$V3[i], full$V8[i], 100-full$V3[i], col=rgb(1,0,0, alpha=full_alpha), lwd=1.5)
    }}
  else{
    par(mar=c(5,5,5,5))
    plot(range(0, cons_len), range(0, auto_y), type = "n", main=paste("TE: ", as.character(blast[1,1]), "\n consensus size: ", as.character(cons_len), "bp; fragments: ", as.character(length(blast$V1)), "; full length: ", as.character(length(full$V1))," (>=",as.character(as.numeric(FL_thresh)*100),"%)", sep = ""), cex.main = 0.9, xlab = "TE consensus (bp)", ylab = "divergence to consensus (%)")
    for(i in 1:length(blast$V1)){
      segments(blast$V7[i], 100-blast$V3[i], blast$V8[i], 100-blast$V3[i], col=rgb(0,0,0,alpha=alpha))
    }
    for(i in 1:length(blast$V1)){
      segments(full$V7[i], 100-full$V3[i], full$V8[i], 100-full$V3[i], col=rgb(1,0,0, alpha=full_alpha), lwd=1.5)
    }}
  
  #make the coverage matrix and graph
  if(cover==T){coverage=matrix(rep(0, length(blast$V1)*as.numeric(cons_len)), byrow = T, ncol = as.numeric(cons_len))
  for(i in 1:length(blast$V1)){
    coverage[i,]<-c(rep(0,blast$V7[i]-1),rep(1,abs(blast$V8[i]-blast$V7[i])+1), rep(0,as.numeric(cons_len)-blast$V8[i]))
  }
  par(new=T)
  plot(colSums(coverage), type='l', axes = F, ylab = NA, xlab = NA, col=covcol, ylim = c(0, max(colSums(coverage))))
  axis(side = 4)
  mtext(side = 4, line = 3, 'consensus coverage (bp)')}
  print(colSums(coverage))
}
```
