## Supplemental Code for "MCHelper automatically curates transposable element libraries across eukaryotic species": getlength.sh

```
#! /bin/bash

awk 'BEGIN {OFS = "\n"}; /^>/ {print(substr(sequence_id, 2)" "sequence_length); sequence_length = 0; sequence_id = $0}; /^[^>]/ {sequence_length += length($0)}; END {print(substr(sequence_id, 2)" "sequence_length)}' $1 | awk 'NR > 1 {print $NF}'
```
