## Supplemental Code for "MCHelper automatically curates transposable element libraries across eukaryotic species": README.md

# TE+Aid [![status](https://img.shields.io/badge/status:-test-yellow)]() [![support](https://img.shields.io/badge/support:-yes-green)]()
<img src=https://i.imgur.com/pxxR3Ec.png width="500">

**TE-Aid** is a `shell`+`R` program aimed to help the manual curation of transposable elements (TE). It inputs a TE consensus sequence (fasta format) and requires a reference genome (in fasta as well). Using `R` and the `NCBI blast+ suite`, TE-Aid produces 4 figures reporting:
 1. (top left) the genomic hits with divergence to consensus
 2. (top right) the genomic coverage of the consensus
 3. (bottom left) a self dot-plot 
 4. (bottom right) a structure analysis including: TIR and LTR suggestions, open reading frames (ORFs) and TE protein hit annotation.

*include figure here* <img src=https://github.com/clemgoub/TE-Aid/blob/master/Example/TE1.jpeg width="900">

**Pipeline overview:**

- The TE (ideally, candidate consensus sequence) is searched against the provided reference genome with `blastn` 
	- Fig 1: genomic hits (horizontal lines) are represented relative to the query (TE consensus), the y axis represent the `blastn` divergence
	- Fig 2: pileup of the genomic hits relative to position along the query (TE consensus)
- The query is then blasted against itself in order to detect micro repeats and inversions (putative TIRs, LTRs)
	- Fig 3: self dot-plot and Fig 4 (top): TIR and LTR are suggested (colored arrows)
	- Bonus: a self dot-plot with `emboss dotmatcher` is also produced in an extra file
- Putative ORFs are searched with `emboss getorf` and the peptides queried against a TE protein database (distributed with [`RepeatMasker`](https://github.com/rmhubley/RepeatMasker))
	- Fig 4: ORFs (black rectangles: + orientation; red rectangles: - orientation), TE protein hits 

The consensus size, number of fragments (hits) and full length copies (according to user-defined threshold) are automatically printed on the graph.
If any ORFs and protein hits are found, their locations relative to the consensus are printed in the `stdout`


TE-Aid has been tested on MacOSX (shell, sh, zsh) and Linux (shell, sh)

**TE-Aid** comes from `consensus2genome` that is now deprecated

## Install

### Dependencies

- [R (Rscript)](https://cran.r-project.org/mirrors.html)
- [NCBI Blast+ suite](https://ftp.ncbi.nlm.nih.gov/blast/executables/blast+/LATEST/)
- [EMBOSS `getorf`](http://emboss.sourceforge.net/download/)

TE-Aid calls **NCBI blast** and **R** from the command line with `blastn`, `blastp`, `makeblastdb` and `Rscript` commands. All these executables must be accessible in the user path (usually the case following the default install). 
If not, you need to locate the executables' location and add them to your local path before using TE-Aid.
For instance: 
```
export PATH="/path/to/blast/bins/folder/:$PATH"` 
export PATH="/path/to/R/bins/folder/:$PATH"` 
```
These lines can be added to the user `~/.bashrc` (Linux) or `~/.zshrc` (macOS) to add these programs permanently to `$PATH`.

### Install **TE-Aid** from github
```
git clone https://github.com/clemgoub/TE-Aid.git
```

## Usage and options

### Minimal command line

```
<user-path>/TE-Aid [-q|--query <query.TE.fa>] [-g|--genome <genome.fa>] [options]
```
>**Note.** replace `<user-path>` with the path of the downloaded `TE-Aid` folder.

### Mendatory arguments:
```
    -q, --query                   TE consensus to blast (fasta file)
    -g, --genome                  Reference genome (fasta file)
```
### Optional arguments:

```
    -h, --help                    show this help message and exit
    
    -o, --output                  output folder (default "./")
    -t, --tables                  write features coordinates in tables (self dot-plot, ORFs and protein hits coordinates)
    -T, --all-Tables              same as -t plus write the genomic blastn table. 
                                  Warning: can be very large if your TE is highly repetitive!
    
    -e, --e-value                 genome blastn: e-value threshold to keep hit (default: 10e-8)
    -f, --full-length-threshold   genome blastn: min. proportion (hit_size)/(consensus_size) to be considered "full length" (0-1; default: 0.9)

    -m, --min-orf                 getorf: minimum ORF size (in bp)

    -a, --alpha                   graphical: transparency value for blastn hit (0-1; default 0.3)
    -F, --full-length-alpha       graphical: transparency value for full-length blastn hits (0-1; default 1)
    -y, --auto-y                  graphical: manual override for y lims (default: TRUE; otherwise: -y NUM)

    -D | --emboss-dotmatcher      Produce a dotplot with EMBOSS dotmatcher
```

## Tutorial

In this example we are going to analyze some transposable elements of *Drosophila melanogaster*. The consensus sequences for this tutorial are located in the `Example/` folder, and you will need to download the *D. melanogaster* reference genome (dm6). Let's go!

#### 1. Download the *D. melanogaster* genome

```shell
curl -o Example/dm6.fa.gz https://hgdownload.soe.ucsc.edu/goldenPath/dm6/bigZips/dm6.fa.gz
gunzip Example/dm6.fa.gz
```
*D. melanogaster* TE consensus are present in the folder `Examples`

#### 2. Analyze the TE consensus

Let's start with Jockey, a recent **LINE** element in the *D. melanogaster* genome

```shell
./TE-Aid -q Example/Jockey_DM.fasta -g Example/dm6.fa -o ../dm6example
```
<img src=https://github.com/clemgoub/TE-Aid/blob/master/Example/Jockey.TEaid.png width="1024">

Next is Gypsy-2, from the **LTR** lineage

```shell
./TE-Aid -q Example/Gypsy2_DM.fasta -g Example/dm6.fa -o ../dm6example
```
<img src=https://github.com/clemgoub/TE-Aid/blob/master/Example/Gypsy2.TEaid.png width="1024">
