## Supplemental Code for "MCHelper automatically curates transposable element libraries across eukaryotic species": Run-c2g.R

```
#! /usr/bin/Rscript

#######################################################################################
### Run-c2g.R - V0.1 - Clement Goubert (2020) -           ###
### ------------------------------------------------------------------------------- ###
### This script uses the R function consensus2genome to infer breakpoints           ###
### to split and/or trim TE                                                         ###
### consensi generated by tools like RepeatModeler2, EDTA, REPET,...                ###
### USAGE:  TE-slocer.sh <TE-header> <[>                                            ###                            ###
### query: path to query (fasta file)                                               ###
### db: path to blast db (blast formated nucleotide database)                       ###
#######################################################################################


Args 		=	commandArgs()
query		=	as.character(Args[6])
database	=	as.character(Args[7])
evalue		=	as.numeric(Args[8])
FLthresh 	=	as.numeric(Args[9])
alpha 		=	as.numeric(Args[10])
full_alpha  =	as.numeric(Args[11])
autoy		=	as.character(Args[12]) # TRUE or numeric value for y max
output		=	as.character(Args[13])
selfdb		=	as.character(Args[14]) # bool 
blastp      =   as.character(Args[15]) # includes orfs position
osize		=	as.numeric(Args[16])
wdir        =   as.character(Args[17]) # from the shell: path to running directory 
tables      =   as.character(Args[18])

source(paste(wdir, "/", "consensus2genome.R", sep = ""))
source(paste(wdir, "/", "blastndotplot.R", sep = ""))

pdf(width = 12, height = 12, file = paste(output, "/", tail(strsplit(as.character(query), "/")[[1]],1), ".c2g.pdf", sep=""))
#pdf(width = 16, height = 16, file = paste(tail(strsplit(as.character(query), "/")[[1]],1), ".c2g.pdf", sep=""))

par(mfrow=c(2,2))

print("R: ploting genome blastn results and computing coverage...")

consensus2genome(query 		=	query,
                 db 		=	database,
                 evalue 	=	evalue,
                 FL_thresh 	=	FLthresh,
                 alpha 		=	alpha,
                 full_alpha	=	full_alpha,
                 auto_y 	=	autoy,
                 bins       =   wdir
#                 ,
#                 cover		=	cover,
#                 cov_thresh	=	drops
#                 ,
                 )

print("R: ploting self dot-plot and orf/protein hits...")

blastdotplot(query  =  query,
             db     =  selfdb,
             blast  =  blastp,
             os     =  osize,
             tables =  tables,
             output =  output)

dev.off()
```
