## Supplementary figures and images for "MCHelper automatically curates transposable element libraries across eukaryotic species"

### cons2gen.jpeg

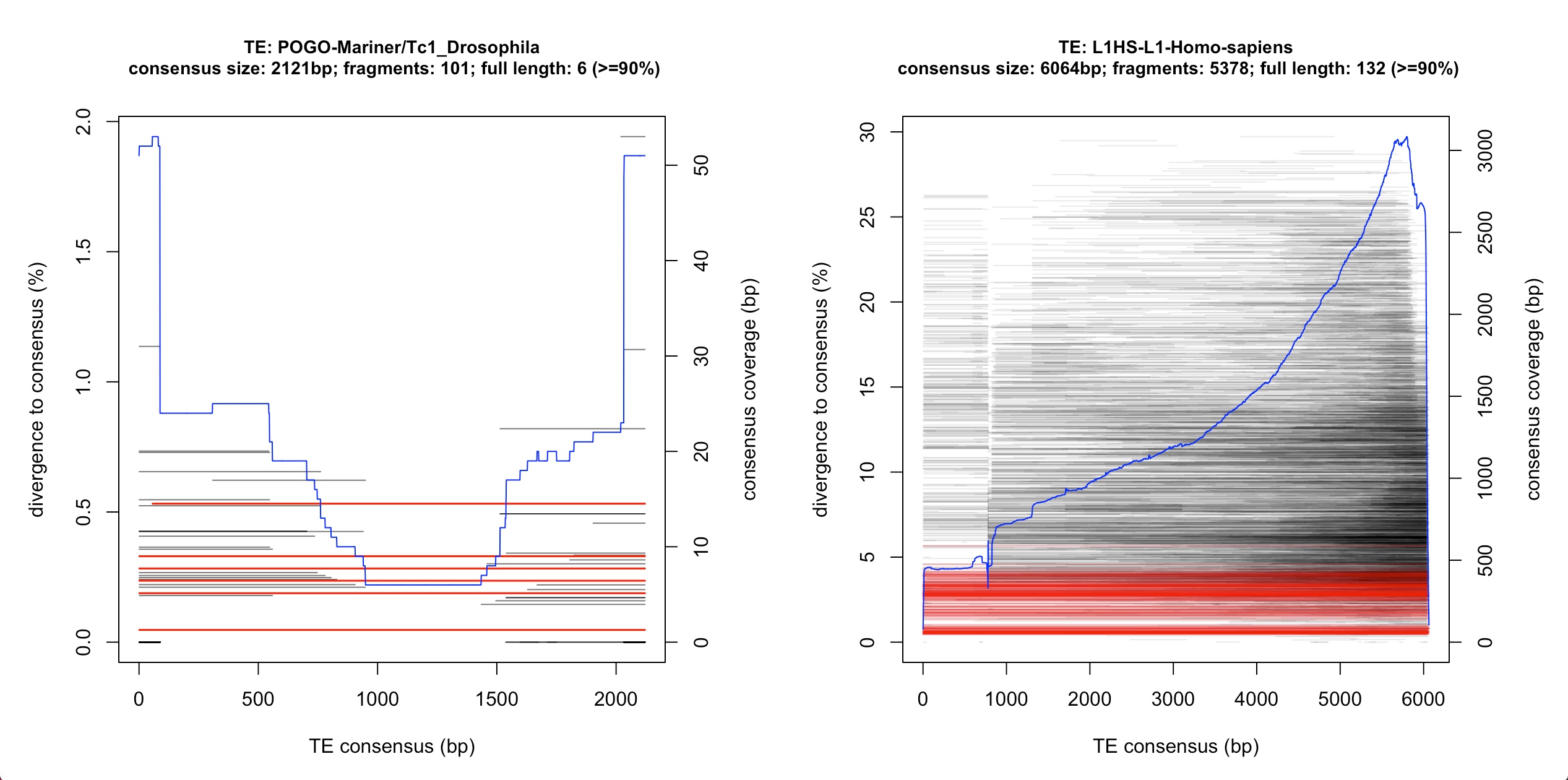

### Gypsy2.TEaid.png

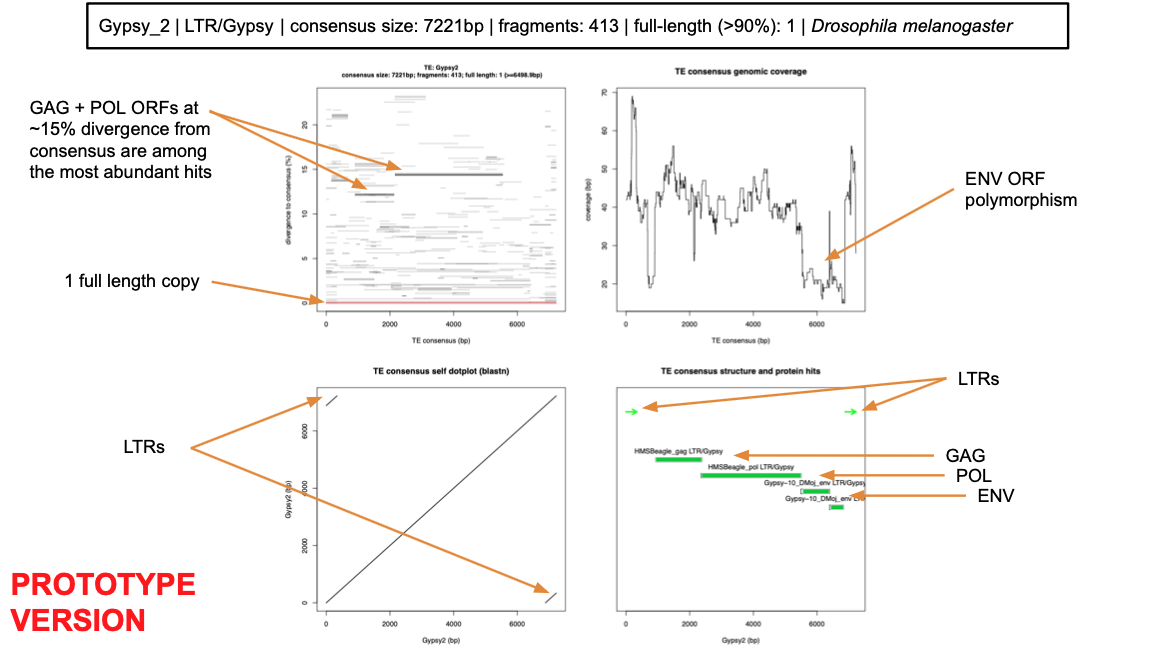

### Gypsy_example.jpeg

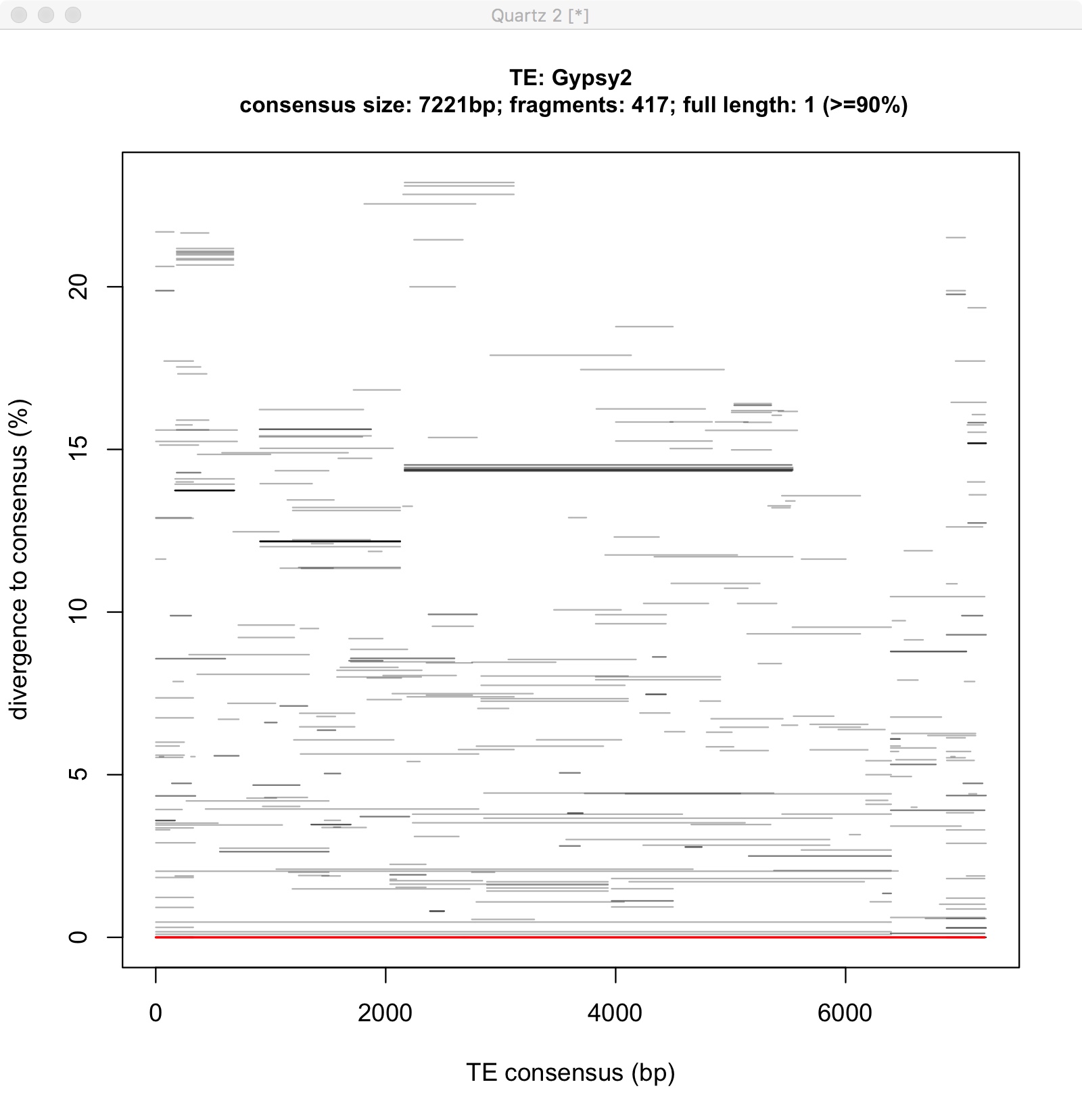

### Jockey-1.jpeg

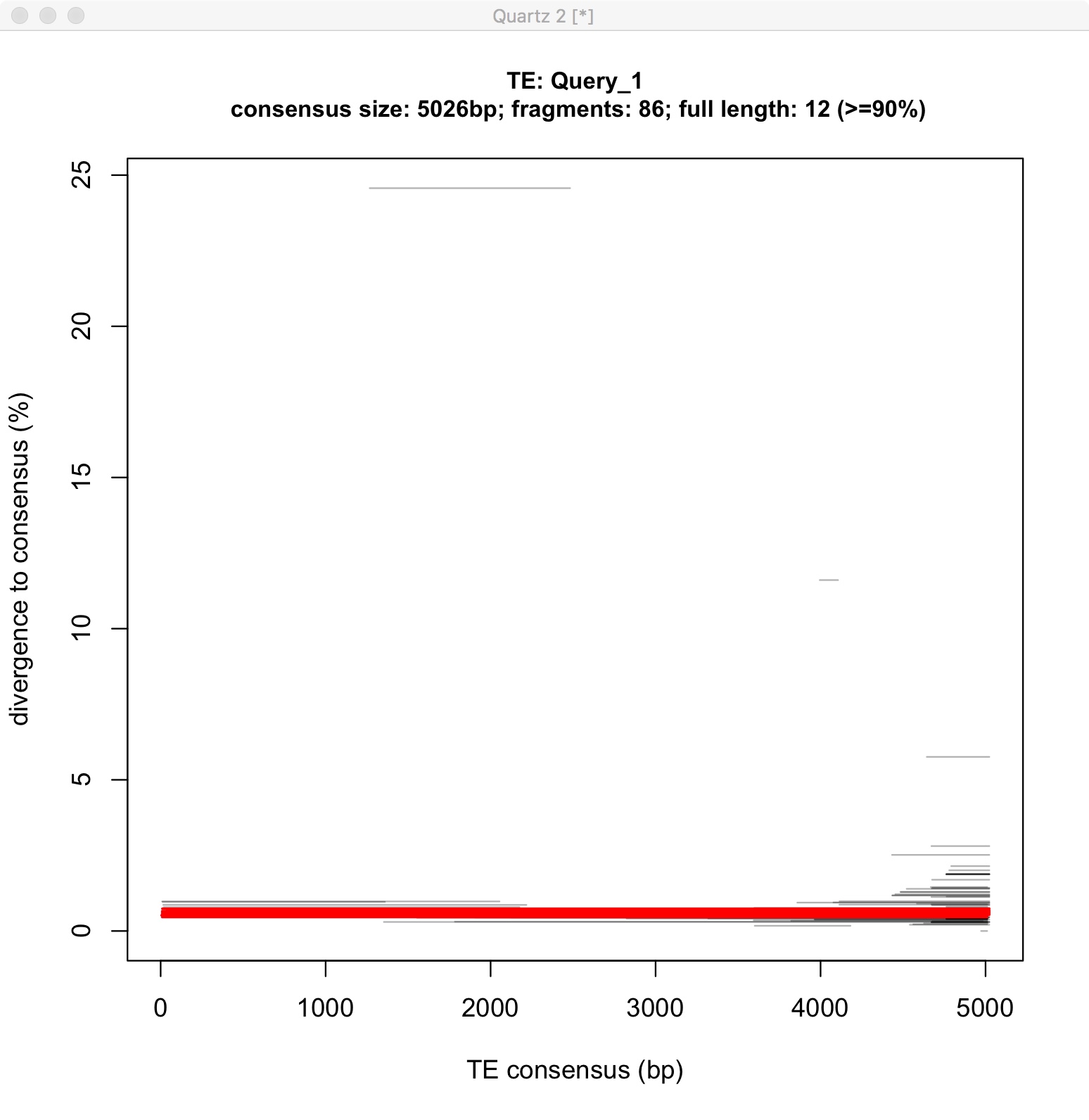

### Jockey-2.jpeg

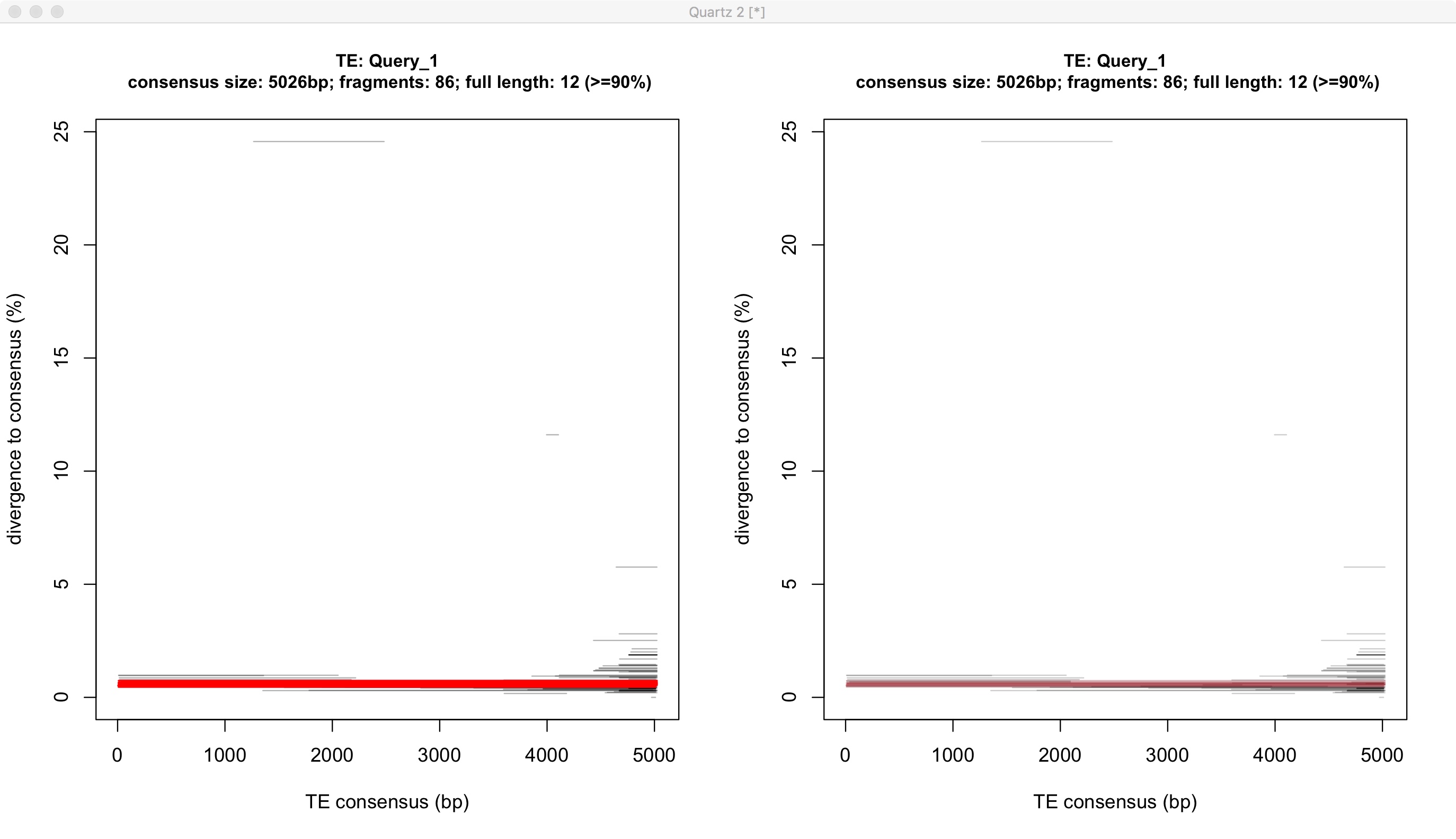

### Jockey-3.jpeg

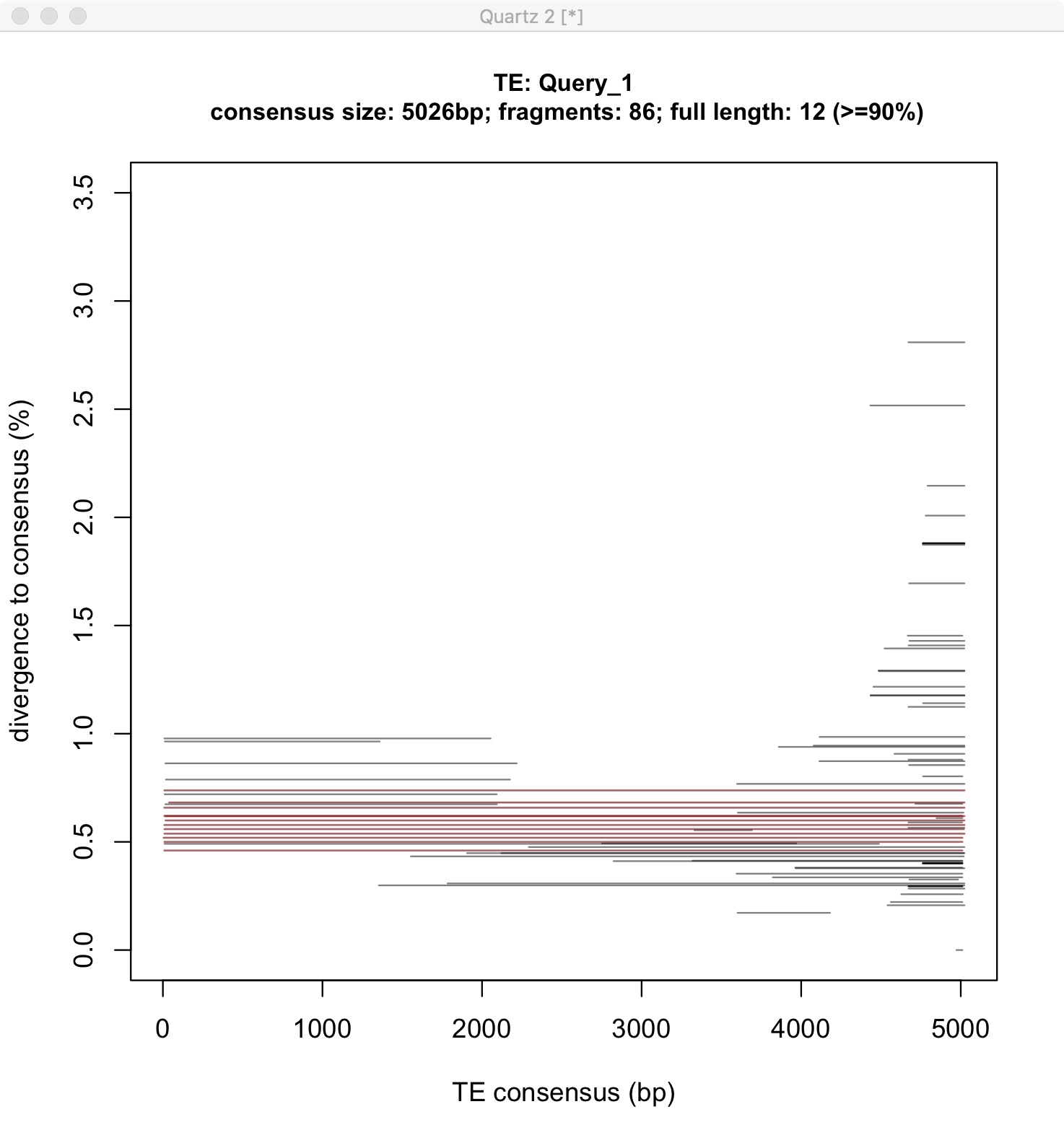

### Jockey.TEaid.png

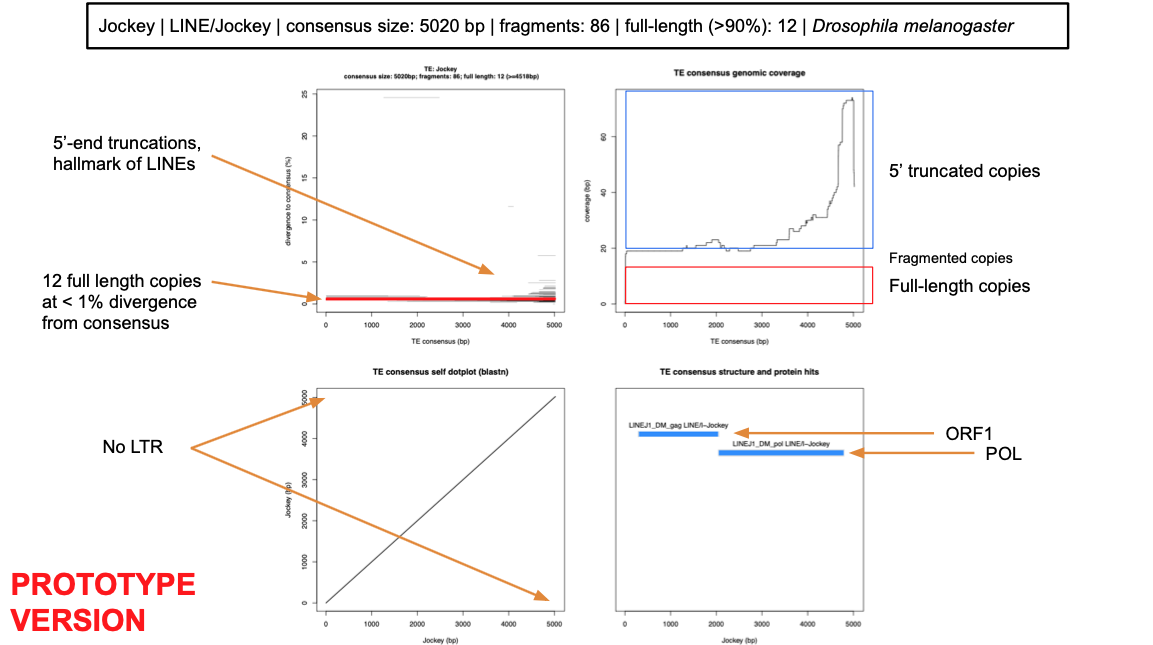

### Jockey_new.jpeg

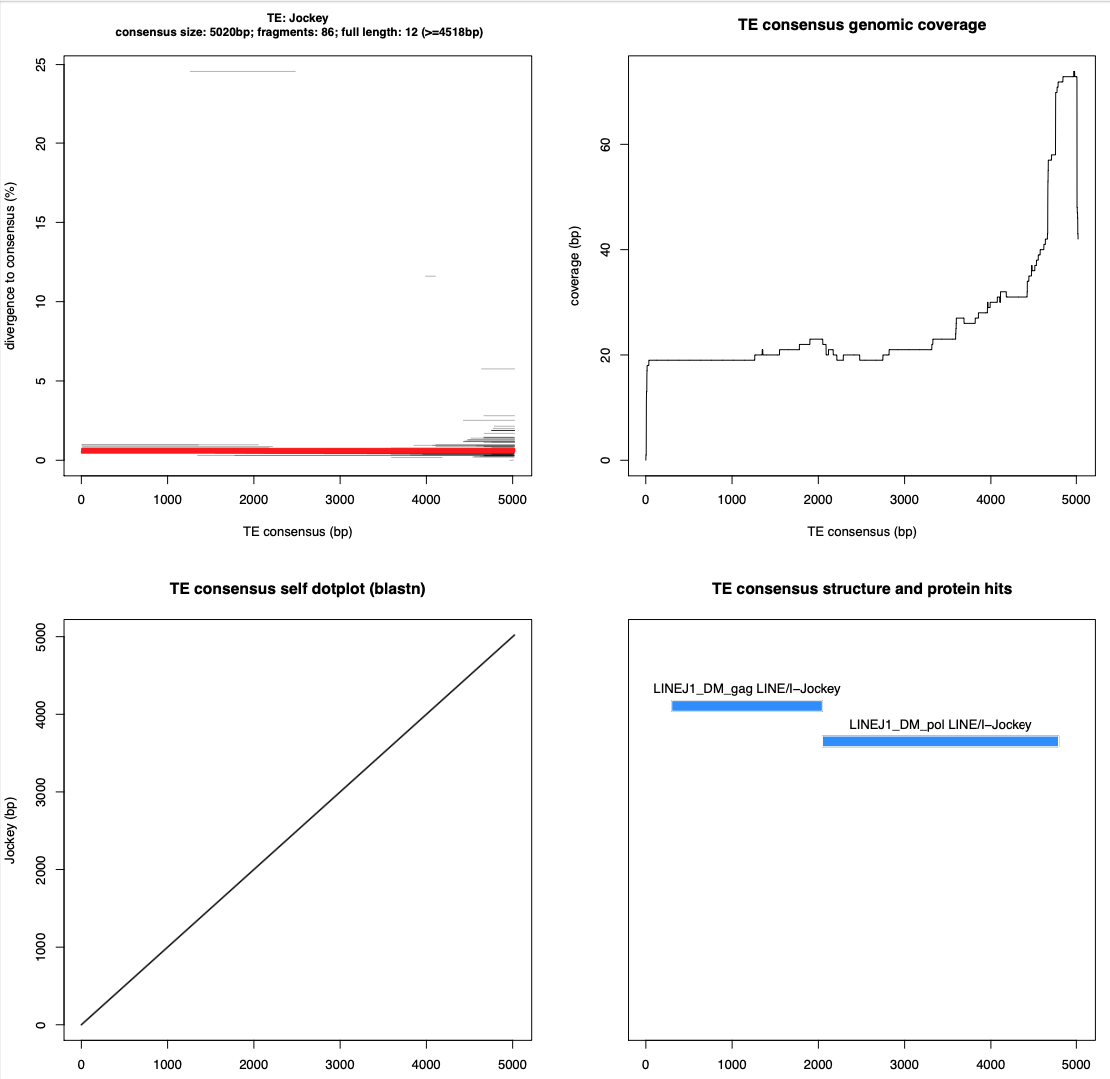

### LINE.1.fasta.c2g.jpg

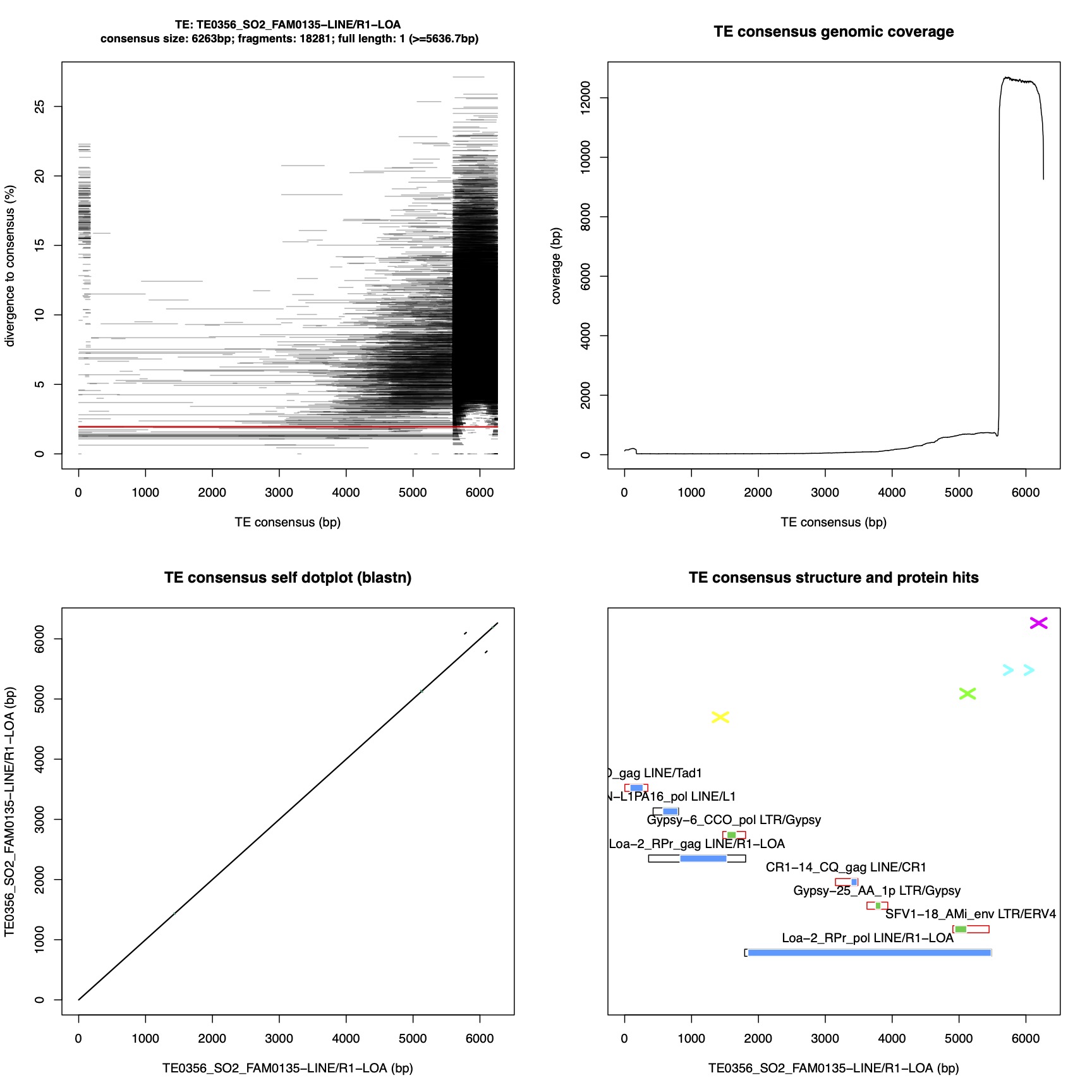

### MCHelper_Flow.jpg

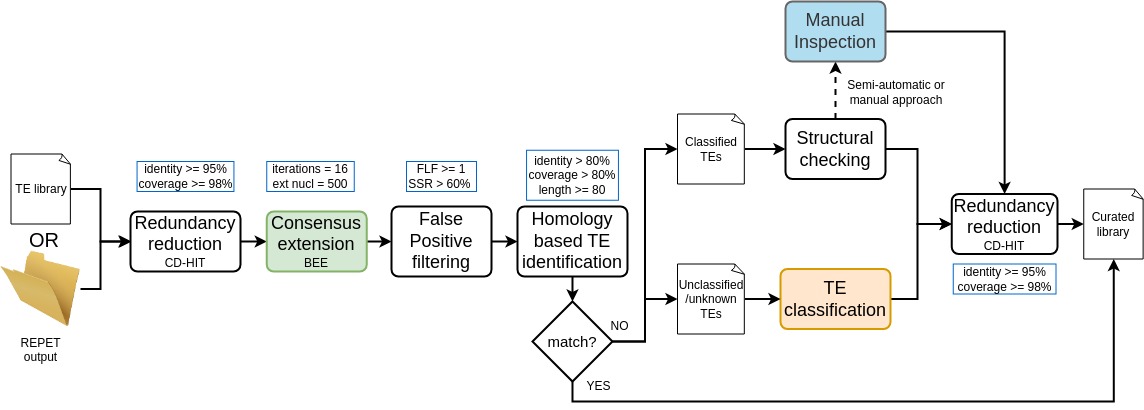

### MCHelper_modules_Flow.jpg

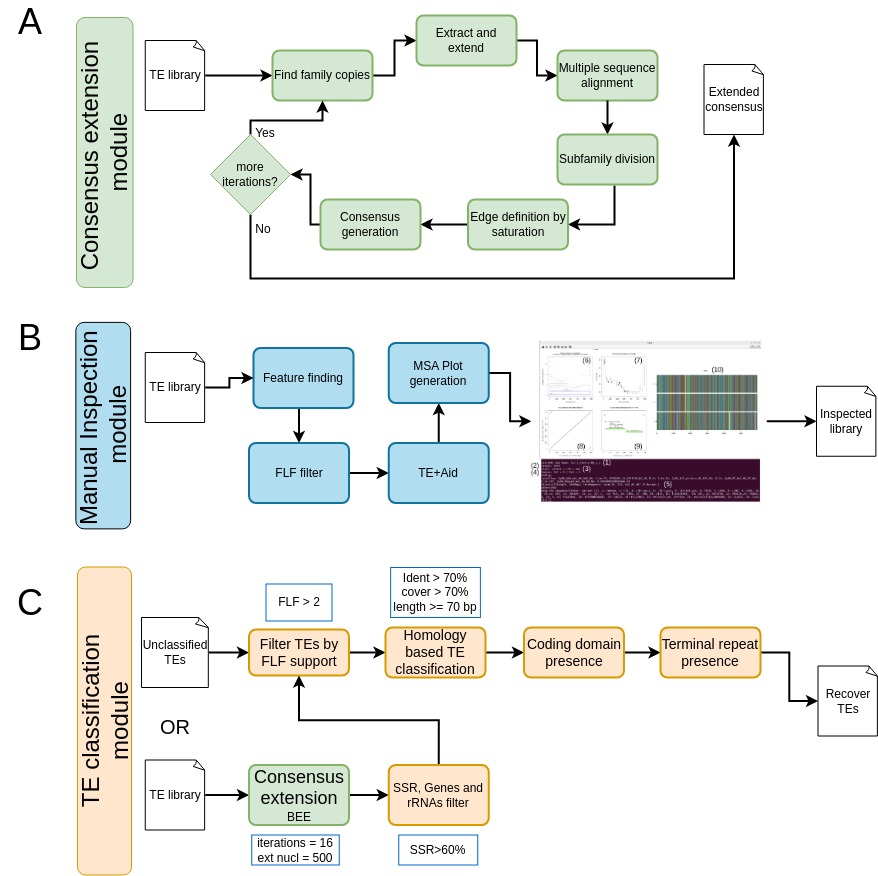

### TE1.jpeg

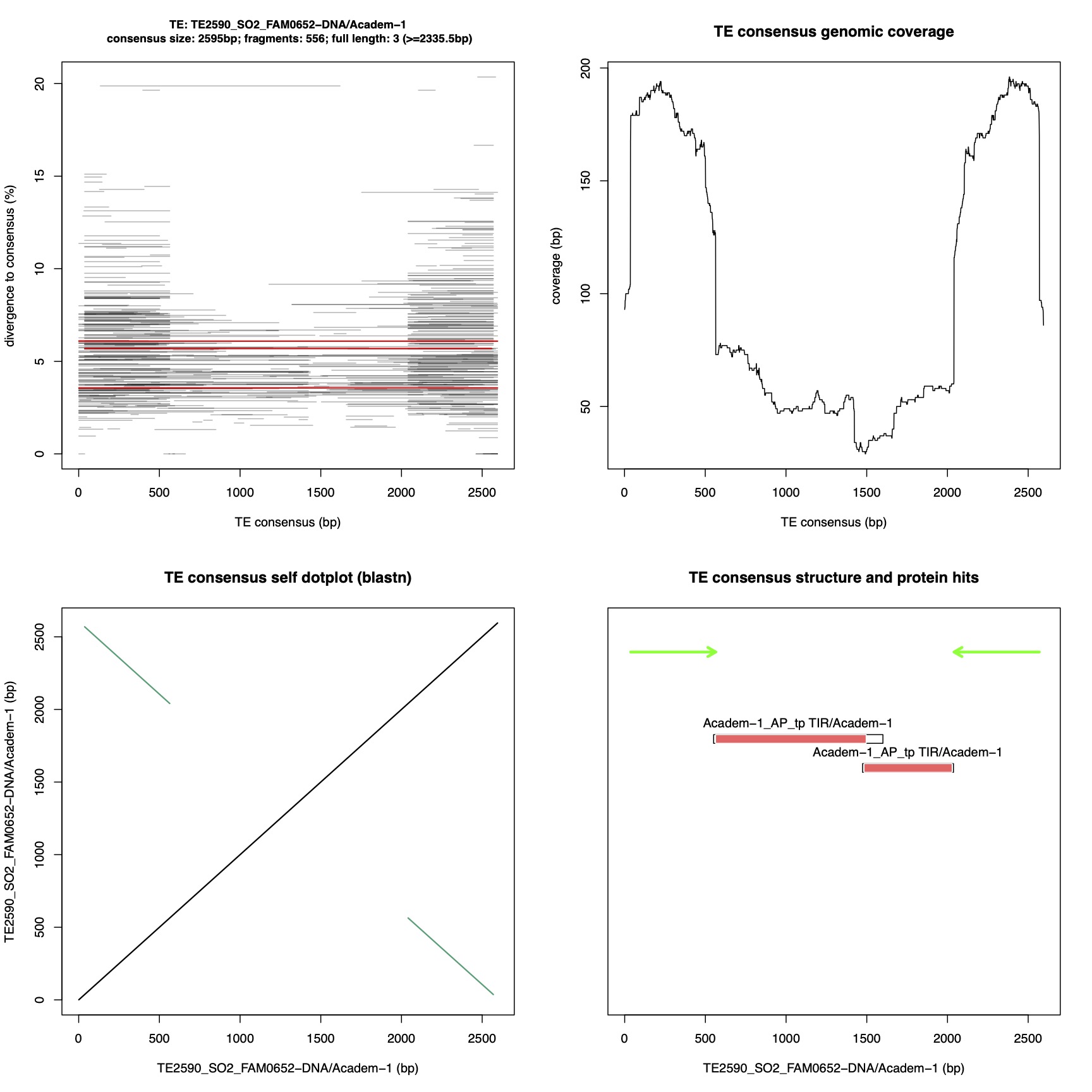
